# Small molecule inhibitors of the NorA multidrug efflux pump potentiate antibiotic activity by binding the outward-open conformation

**DOI:** 10.1101/2025.08.19.671038

**Authors:** Janine L. Gray, Elizabeth V. K. Ledger, Tiffany Suwatthee, Thomas J. Burden, Konstantina Arvaniti, Amber Sefton, Lydia E. Papagora, Thomas B. Clarke, Jennifer Riley, Erika G. Pinto, Fraser Cunningham, Ian H. Gilbert, David Gray, Da-Neng Wang, Kevin D. Read, Thomas Lanyon-Hogg, Nathaniel J. Traaseth, Andrew M. Edwards, Edward W. Tate

## Abstract

Antibiotic resistance is among the greatest threats of the modern era. Multidrug efflux pumps expel antibiotics from bacterial cells and present a particular challenge by conferring resistance to a broad range of antibiotic classes; however, there is currently a lack of potent and selective inhibitors. Here, we report the discovery of **IMP-2380**, a drug-like chemical probe for the multidrug efflux pump NorA that delivers low-nanomolar potentiation of ciprofloxacin activity *in vitro* and activity in an *in vivo S. aureus* infection model. A phenotypic high-throughput screen for inhibitors of the ciprofloxacin-activated SOS DNA repair pathway in methicillin-resistant *Staphylococcus aureus* (MRSA) identified hit compounds targeting NorA, and subsequent optimization established **IMP-2380** as the most potent NorA inhibitor discovered to date. The structure of NorA bound to **IMP-2380** was solved by cryo-electron microscopy at 2.52 Å resolution, revealing that the small molecule locks the pump in the ‘outward-open’ conformation. This closes the inner face and prevents antibiotics binding from the cytosol, providing an explanation for the exceptional potency of **IMP-2380** and structure-activity relationship across the series. **IMP-2380** represents an *in vivo* active NorA inhibitor, functioning via a structurally defined outward-open binding mode, and will enable future exploration of NorA as a druggable target to combat antibiotic resistance.

## Main

Antimicrobial resistance (AMR) is a global healthcare challenge, with drug-resistant infections causing 1.27 million deaths in 2019, expected to increase to 10 million per annum by 2050 if left unaddressed.^1–3^ The emergence and dissemination of multidrug resistant bacteria such as methicillin resistant *Staphylococcus aureus* (MRSA) has been accelerated by excessive prescription and misuse of antibiotics and there is now documented resistance against nearly all known antibiotics. Furthermore, of the ∼60 compounds in clinical development targeting World Health Organization (WHO) priority pathogens in 2023, only 38% were considered ‘innovative’ in terms of mechanism of action (MoA), target or chemical scaffold, greatly increasing the risk of rapid emergence of resistance.^4^ There is, therefore, a pressing unmet need to develop new therapeutics with novel MoA to treat or prevent bacterial infections.^5^ Antibiotic adjuvants are compounds that augment existing antibiotics through inhibition of resistance mechanisms or by modifying the host immune response, whilst lacking intrinsic antibacterial activity which might drive further resistance.^6,7^ This approach was first validated in the clinic by the β-lactamase inhibitor clavulanic acid,^8^ driving interest in targeting other important resistance mediators such as drug efflux pumps, which prevent intracellular antibiotic accumulation through active efflux coupled to proton or ion gradients, or ATP hydrolysis.^9–11^ Antibiotic efflux is a first line defense mechanism used by bacteria, with a number of structural studies revealing mechanistic insights that have guided the development of efflux pump inhibitors (EPIs).^12–14^ However, despite the presence of a number of reported EPIs in the literature, their low potency or toxicity has prevented progression into the clinic.^15,16^

MRSA employs multiple stress response and repair systems to survive in the hostile host environment, including the mutagenic SOS response, which is triggered by DNA damage caused by host immune cells or DNA-damaging antibiotics such as ciprofloxacin or co-trimoxazole.^17^ The SOS response drives homologous recombination to ensure DNA repair and bacterial survival, and induces expression of genes encoding low-fidelity DNA polymerases associated with mutagenesis, accelerating the emergence of mutants with decreased susceptibility to antibiotics and host defenses thus promoting persistent infections (Fig. S1a). Importantly, MRSA mutants defective for DNA repair and the SOS response are less able to sustain infection or survive in human blood.^17^ Due to the key role of the SOS response for bacterial survival and resistance generation, it has been proposed that adjuvants inhibiting this pathway might be promising therapeutics for MRSA in combination with existing antibiotics.^18–21^

Here, we report the discovery of **IMP-2380**, a drug-like low nanomolar inhibitor of the ciprofloxacin-induced SOS response in cells. Target deconvolution studies established efflux pump NorA as the direct target of **IMP-2380** and through cryo-electron microscopy (cryo-EM) we show that **IMP-2380** interacts with NorA extensively and locks the transporter in the ‘outward-open’ conformation explaining its potent inhibition of antibiotic efflux. We use **IMP-2380** to demonstrate robust potentiation with ciprofloxacin both *in vitro* and in an *in vivo* model of MRSA infection. **IMP-2380** is the first potent chemical probe for NorA with *in vivo* activity and a structurally defined binding mode, demonstrating the potential for selective efflux pump inhibition as a therapeutic strategy against antibiotic resistance.

## Results

### High throughput phenotypic screening identifies a novel inhibitor of the SOS response

To identify small molecule inhibitors of the SOS response, we used a cellular reporter assay in the MRSA strain USA300 JE2, where green fluorescent protein (GFP) expression was under the control of the *recA* promoter, which is induced by the SOS response (Fig. 1a, Fig. S1a).^19^ Assay conditions were optimized for high-throughput screen (HTS) compatibility using both a traditional ‘one factor at a time’ approach, as well as a ‘design of experiments’ multifactorial approach using a custom D Optimal design which generated comparable optimization results in a single experiment (Fig. S2).^22^ A HTS of 33,000 diverse compounds was performed at 30 μM compound with 48 μM ciprofloxacin to trigger DNA damage in a 384-well format (mean robust Z’ = 0.68 ± 0.11, Extended Data Fig. 1a), with hits classified as ≥60% inhibition of the maximal GFP signal normalized to bacterial growth (OD_600_) (Fig. S1b, Extended Data Fig. 1b). Following dose-response analysis, compounds were prioritized for an adjuvant mechanism of action by removing those that exhibited mammalian cytotoxicity or inhibition of bacterial cell growth in the absence of ciprofloxacin. Compound **1** (Fig. 1b) was subsequently selected for further investigation. **1** showed good potency in the SOS response assay (EC_50_ = 80 nM, Fig. 1c) and no intrinsic antibacterial activity (EC_50_ > 30 μM, Extended Data Fig. 2a), avoiding selection pressure for resistance-conferring mutations. **1** also had no impact on mammalian cell proliferation (EC_50_ > 30 μM, Extended Data Fig. 2b) or cytotoxicity (CC_50_ > 30 μM, Extended Data Fig. 2c). There are no associated assay data for **1** or similar scaffolds in the compound activity database ChEMBL, suggesting that this scaffold has not previously been reported as bioactive, and there are no documented off-targets in mammalian cells.^23,24^

**Figure 1.**
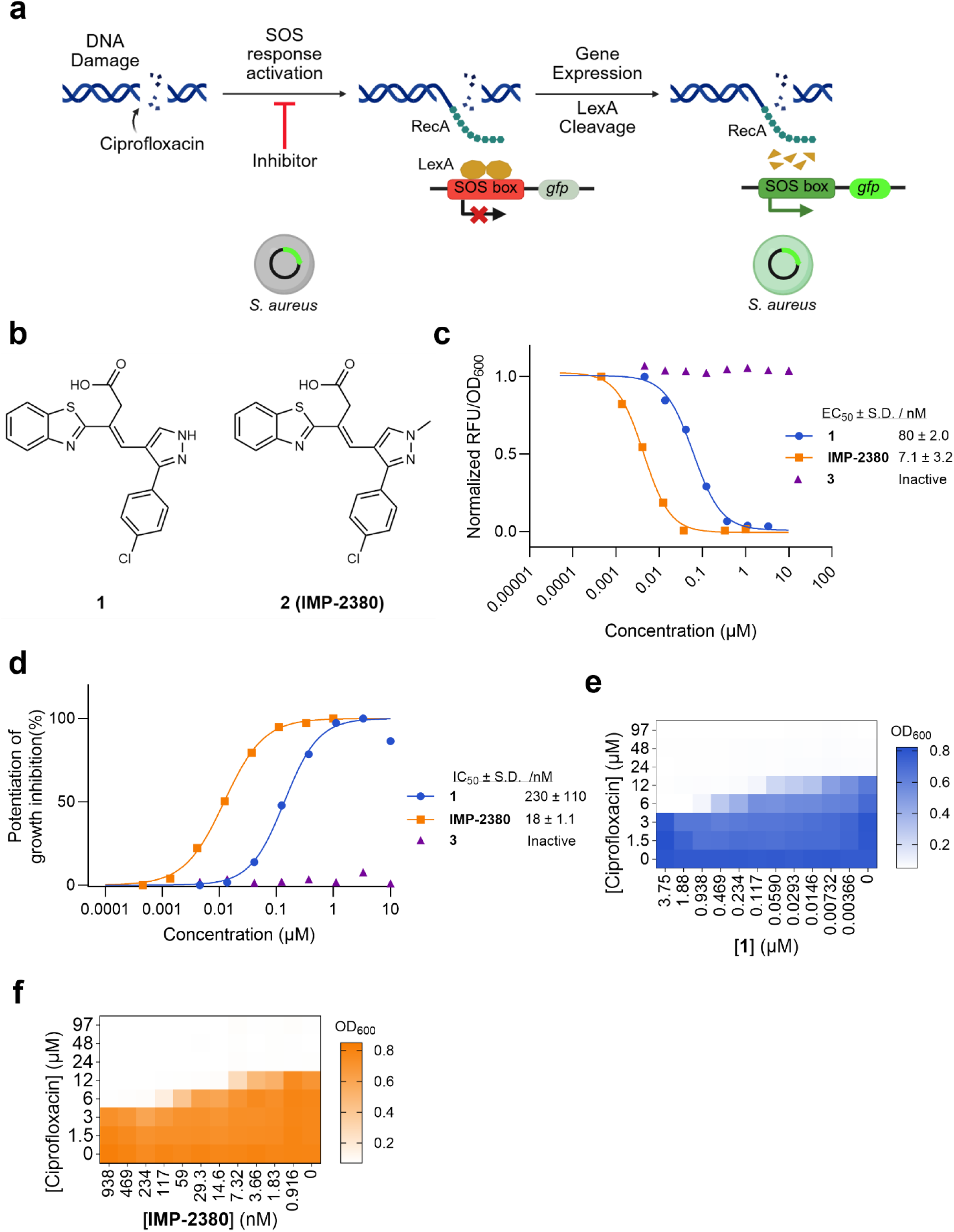
HTS identifies a potent inhibitor series for the SOS response in *S. aureus.* (**a**) The P*recA-gfp* reporter system in *S. aureus* USA300 JE2 measures induction of the SOS response. The SOS response can be activated by ciprofloxacin which causes DNA double strand breaks by interrupting DNA replication, resulting in DNA processing and RecA assembly into a nucleofilament in the region of DNA damage, which catalyzes autocleavage of transcriptional repressor LexA, resulting in expression of the SOS regulon, including *recA*. Using this reporter system, the activation of the SOS response by ciprofloxacin was measured as a function of GFP fluorescence. Created in BioRender. (**b**) Structures of hit compound **1** identified in SOS response inhibitor HTS (Fig. S1b), and lead compound **2** (**IMP-2380**) identified in initial SAR studies. (**c**) Dose-response curve for inhibition of SOS response by **1**, **IMP-2380** or inactive control **3** (Extended Data Fig. 3) in combination with 97 μM ciprofloxacin as measured using the P*recA-gfp* reporter system in *S. aureus* USA300 JE2. EC_50_ values are expressed as the geometric mean +/- SD. (n ≥ 3). Data shown are of a single replicate. (**d**) Dose-response curve for potentiation of ciprofloxacin-induced bacteria growth inhibition in the P*recA-gfp* reporter system in JE2 by **1**, **IMP-2380** or inactive control **3** in combination with 97 μM ciprofloxacin. EC_50_ values are expressed as the geometric mean +/- SD. (n ≥ 3). Data shown are of a single replicate. (**e-f**) Checkerboard MIC assays of **1** (**e**) or **IMP-2380** (**f**) and ciprofloxacin against the JE2 WT strain (n = 2).

We investigated the synergistic activity of **1** with antibiotics to understand its mechanism of action. **1** potentiated ciprofloxacin-induced inhibition of bacteria growth in the ciprofloxacin-resistant strain JE2 (IC_50_ = 230 nM, Fig. 1d) and potentiated ciprofloxacin activity in checkerboard minimum inhibitory concentration (MIC) assays (Fig. 1e), reaching a 4-fold decrease in ciprofloxacin MIC with increasing concentration of **1**. This activity was recapitulated in the ciprofloxacin-sensitive strain SH1000 (Extended Data Fig. 2d), and in clinical isolates of MRSA and methicillin-sensitive *S. aureus* (MSSA), demonstrating a 2-to 8-fold decrease in ciprofloxacin MIC at nanomolar concentrations (Table S1). In contrast, **1** had no significant effect on the susceptibility to nitrofurantoin, co-trimoxazole, daptomycin or cloxacillin (Extended Data Fig. 2e, Fig. S3). The considerable reduction of ciprofloxacin MIC at nanomolar concentrations of **1**, and the lack of potentiation of other classes of antibiotics, suggested that **1** inhibits the SOS response via a selective mechanism for fluoroquinolones.

### Initial structure-activity studies identify potent inhibitor IMP-2380

Initial structure-activity relationship (SAR) studies identified the importance of the carboxylic acid and pyrazole-phenyl motifs, alterations to which resulted in a significant or complete loss of activity (Extended Data Fig. 3). Replacement of the benzothiazole with alternative heterocycles, or the alkene with an amide, was also not tolerated. Compound **2** (hereafter named **IMP-2380**) in which the pyrazole ring in **1** is methylated exhibited a 10-fold improved EC_50_ of 7.1 nM in inhibiting the SOS response and IC_50_ of 18 nM in the potentiation of ciprofloxacin-induced bacterial growth inhibition assay (Fig. 1b-d). **IMP-2380** caused a similar fold-decrease in ciprofloxacin MIC as for **1**, but at ca. 10-fold lower concentration, suggesting a conserved target (Fig. 1f, Extended Data Fig. 2f). Compounds which did not inhibit the SOS response demonstrated no potentiation of ciprofloxacin activity (**3-10**, Fig. 1c-d, Extended Data Fig. 2g). Similarly to **1**, **IMP-2380** showed no mammalian cytotoxicity and no antibacterial activity in the absence of ciprofloxacin (Extended Fig. 2b-c, Extended Fig. 2h). Finally, we tested **IMP-2380** in MRSA and MSSA clinical isolates, where it potentiated ciprofloxacin activity to the same extent as **1**, but at a lower concentration (Table S1).

### IMP-2380 and 1 are potent and selective inhibitors of the *S. aureus* NorA drug efflux pump

To identify cellular targets of this series, two independent parallel transposon screens were undertaken in different genetic backgrounds (Fig. 2a). Firstly, we generated a saturated mutant library in the ciprofloxacin-susceptible *S. aureus* strain TM51 using a set of transposons with outward facing promoters,^25^ resulting in a pool of mutants with differential expression of specific genes relative to wild type (WT). By selecting for resistance to a normally growth inhibitory combination of 1.2 μM ciprofloxacin and 0.25 μM compound **1**, two independent mutants were isolated, both of which were found to have the transposon inserted upstream of the *norA* gene in an orientation that drives gene overexpression (Fig. 2b, Fig. S4). The *norA* gene encodes the NorA multidrug efflux pump present in all strains of *S. aureus* which is specific for certain quinolone antibiotics.^26^ Consistent with this finding, these isolated resistant mutants were also four-fold less susceptible to ciprofloxacin relative to WT bacteria and required more inhibitor to reduce the ciprofloxacin MIC than for the WT strain (Extended Data Fig. 4a, Fig. S4).

**Figure 2.**
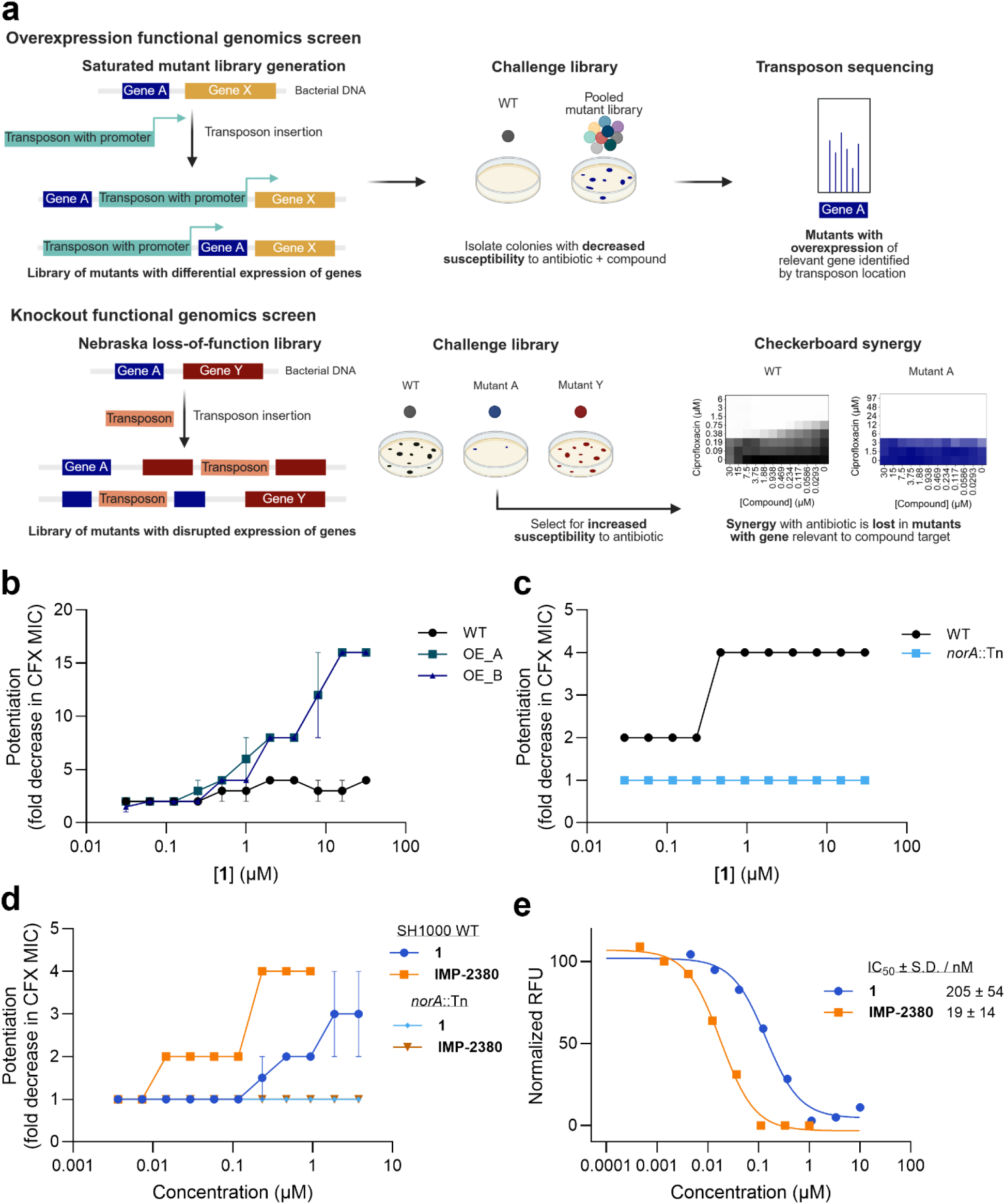
Identification of NorA as the cellular target of **IMP-2380** series. (**a**) Schematic of the transposon library strategies used to identify NorA as the cellular target. Overexpression functional genomics screen: Saturated mutant library was generated where genes were overexpressed. Pools of mutants were challenged with compound and ciprofloxacin. Colonies which survived were isolated and sequenced to identify the site and orientation of transposon insertion. Knockout functional genomics screen: Nebraska loss-of-function mutant library whereby genes have disrupted expression were challenged with ciprofloxacin. Mutants with increased susceptibility to ciprofloxacin as compared to wild type (WT) were taken forward for checkerboard MIC assays to identify mutants where the compound did not potentiate ciprofloxacin. Created in BioRender. (**b**) Fold change in ciprofloxacin (CFX) MIC in *S. aureus* TM51 WT or *norA* overexpression mutants (OE_A and OE_B) in response to varying concentrations of **1** (n = 2). Data represents median MIC ± 95% CI (**c**) Fold change in CFX MIC in *S. aureus* USA300 JE2 WT and *norA*::Tn strains in response to varying concentrations of **1** (n = 2). (**d**) Fold change in CFX MIC *S. aureus* SH1000 WT and *norA*::Tn strains in response to varying concentrations of **1** (n = 2) or **IMP-2380** (n = 1). Data represents median MIC ± 95% CI. (**e**) Dose-response curves for inhibition of NorA-catalyzed EtBr efflux in *S. aureus* JE2 WT induced by **1** or **IMP-2380**. IC_50_ values are expressed as the geometric mean +/- SD. (n ≥ 4). Data shown are of a single replicate.

Secondly, ciprofloxacin-susceptible mutants from the Nebraska loss-of-function transposon library were screened for loss of ciprofloxacin potentiation by **1**.^27^ This library was constructed in the ciprofloxacin-resistant JE2 background and each mutant carries a single transposon insertion in a defined gene, disrupting its expression (Fig. 2a). Mutants with increased susceptibility to ciprofloxacin were identified, and the ability of **1** to potentiate ciprofloxacin was tested in each of these mutants by checkerboard MIC assay. The only mutant in which compound **1** potentiation activity was lost contained an interrupted copy of *norA* (Fig. 2c, Fig. S5). This *norA*::Tn mutant exhibited four-fold increased susceptibility to ciprofloxacin as expected from loss of the NorA efflux pump and showed no further increase in susceptibility on addition of **1** (Extended Data Fig. 4b, Fig. S5). Additionally, we transduced the *norA*::Tn mutation into the ciprofloxacin-susceptible SH1000 strain, which resulted in a four-fold increase in susceptibility to ciprofloxacin compared to WT, and a loss of potentiation activity for both compound **1** and **IMP-2380**, consistent with NorA being the target of the series (Fig. 2d, Extended Data Fig. 4c).

Next, to understand whether these compounds directly interfered with NorA efflux activity, we used a previously established ethidium bromide (EtBr) accumulation assay.^28^ It was first demonstrated that EtBr efflux in *S. aureus* occurs selectively via NorA using a panel of mutants defective for various established and putative efflux systems (Extended Data Fig. 4d). **1** inhibited EtBr efflux with IC_50_ of 205 nM, whilst **IMP-2380** showed a 10-fold improved IC_50_ of 19 nM, fully consistent with the differential activity of these compounds in SOS and antibiotic potentiation assays (Fig. 2e). **IMP-2380** delivers a >400-fold improvement in potency over reserpine (IC_50_ = 8 μM), the most widely used reference compound for NorA inhibition.^29^ Finally, we evaluated ciprofloxacin potentiation by **IMP-2380** in a panel of different bacterial strains (Extended Data Fig. 4e, Fig. S6). As expected, **IMP-2380** showed no potentiation activity in *E. coli* or *S. agalactiae*, as *norA* is not present in these strains. At 15-fold the concentration required in *S. aureus*, **IMP-2380** caused a two-fold decrease in ciprofloxacin MIC in *E. faecalis*. This is most likely explained by the presence of *emeA* in *E. faecalis*, which encodes a member of the MFS superfamily of efflux pumps which is a close orthologue of *norA*.^30^

Taken together, these data are consistent with **1** and **IMP-2380** being selective inhibitors of the NorA drug efflux pump, with **IMP-2380** exhibiting weak activity on close *norA* orthologues at high concentrations.

### Cryo-EM structure reveals IMP-2380 binds the outward-open state of NorA

To establish the inhibitory mechanism of this series, we determined the cryo-EM structure of **IMP-2380** bound to NorA. Our approach involved incubating **IMP-2380** with a NorA construct where a BRIL domain was fused to the C-terminus of NorA (NorA-BRIL) and bound to multiple fiducial marks to increase the size of the complex for cryo-EM structure determination.^31^ Following sample preparation, cryo-EM data collection, and image processing, we obtained a high-quality cryo-EM map with a resolution of 2.52 Å that facilitated model building for all NorA side chain residues within TM segments (Extended Data Fig. 5-6, Table S2, PDB ID: 9VP0). Notably, we observed a continuous, non-protein density matching the structure of **IMP-2380** which could be modeled with high confidence (Fig. 3a).

**Figure 3.**
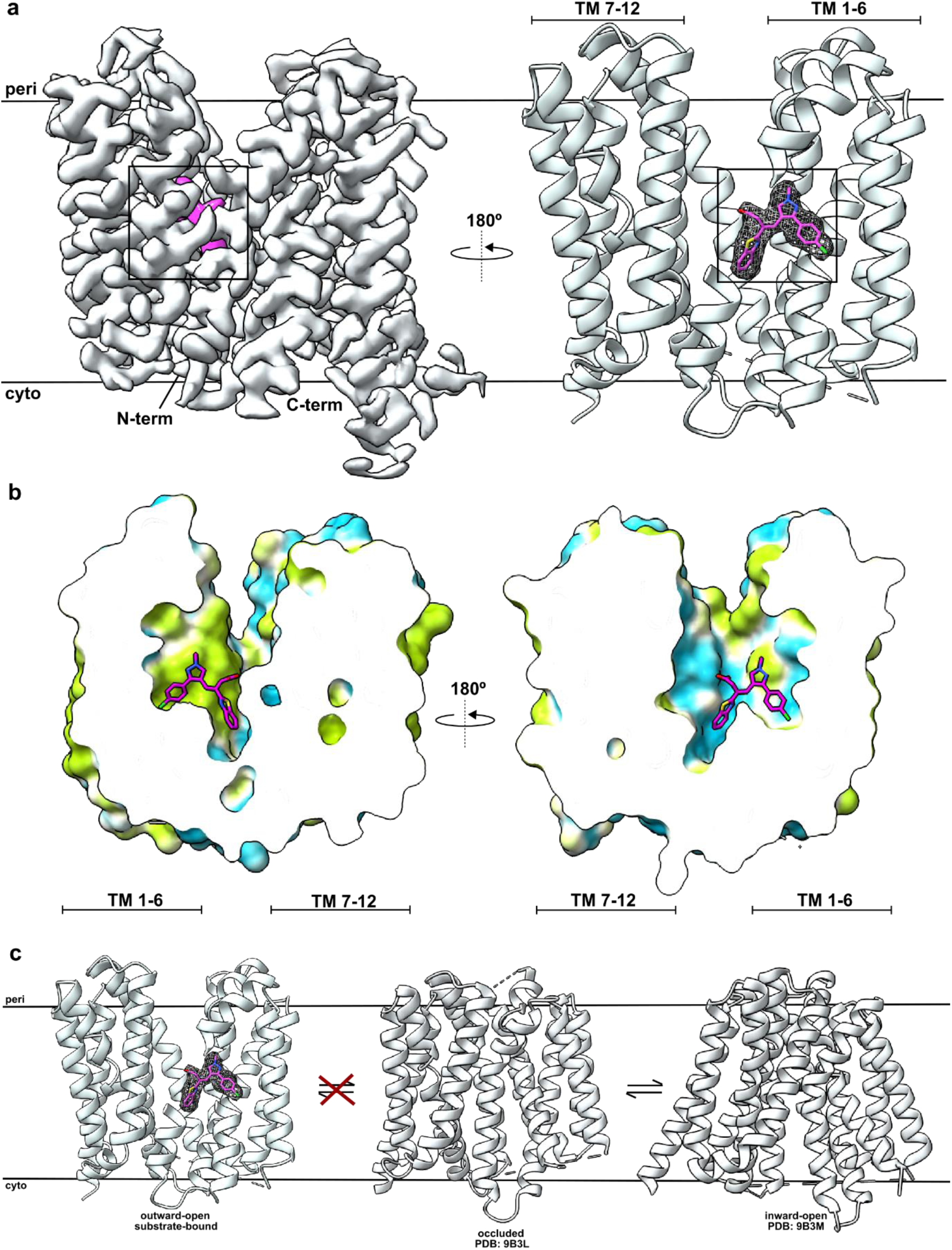
Cryo-EM structure of **IMP-2380** bound to NorA. (**a)** Cryo-EM map of NorA bound to **IMP-2380** complex; density of **IMP-2380** is highlighted in magenta (left) and outlined in a black box. Cartoon representation of NorA structure with cryo-EM density of **IMP-2380**. The density is contoured at 10σ and shown in dark grey mesh, with **IMP-2380** depicted as magenta sticks (right) and outlined in a black box. Parts of TM1-6 are hidden for visibility of **IMP-2380**. (**b**). Two views of cut open hydrophobic surface of NorA + **IMP-2380**. Hydrophobic surfaces are shown in light green and hydrophilic surfaces are shown in light blue. (**c**) Schematic of **IMP-2380**-dependent inhibition of NorA efflux. **IMP-2380** binding to the outward-open conformation of NorA prevents efflux and a conformational switch to the cytoplasmic substrate pocket in the inward-open conformation.

The structure of the complex revealed **IMP-2380** bound to NorA in the outward-open conformation. The small molecule **IMP-2380** adopts the shape of a saddle and inserts deeply within the bottom portion of the central cavity of NorA, defined as the region between the N-terminal domain (NTD, TM1-6) and C-terminal domain (CTD, TM7-12) (Fig. 3b). We propose that **IMP-2380** inhibits efflux of fluoroquinolones and EtBr by trapping the outward-open conformation (Fig. 3c) and preventing NorA from sampling other structural states in the catalytic cycle needed for proton and antibiotic transport, such as inward-facing conformations likely required for binding antibiotics.^31,32^ The quality of the cryo-EM map of NorA bound to **IMP-2380** indicates that the compound blocks such conformational rearrangements and renders the fluoroquinolone substrate pocket inaccessible from the cytosol, explaining the robust potentiation of ciprofloxacin observed. Furthermore, this binding mode to the outward-open conformation is accessible from the extracellular space, allowing compounds in this series to bind without crossing the bacterial membrane.

The cryo-EM structure of NorA in complex with **IMP-2380** is the first bound to a small molecule inhibitor or substrate and establishes a novel binding site in the transporter not previously observed. Whilst prior outward-open structures for NorA in complex with Fabs showed inhibitory interactions with essential NorA residues required for proton-coupled antibiotic efflux,^33^ **IMP-2380** does not interact with these residues and instead accesses two deeply buried cavities in the NTD of NorA, effectively pinning the base of the transporter in an inactive state (Extended Data Fig. 7a).

### Defining key structural features driving NorA binding in the outward-open conformation

The majority of intermolecular interactions between **IMP-2380** and NorA are formed with residues from the NTD. We categorized these interactions into four groups, supported by our preliminary SAR studies (Extended Data Fig. 3). First, the carboxylic acid motif and nitrogen of the benzothiazole ring of **IMP-2380** are positioned for hydrogen bonding with Asn340 and Gln51 of NorA, respectively (Fig. 4a, b). Second, the proximity of the benzene ring of the benzothiazole and the methyl of Thr336 from NorA suggest a C-H π interaction, which is likely strengthened by a hydrogen bond between the side chains of Thr336 and Arg310. Third, the *E*-configured double bond of **IMP-2380** positions the benzothiazole and phenyl substituents into two oppositely facing pockets in NorA (Fig. 3b). The phenyl ring binds within a distinct, hydrophobic cavity in the NTD formed by TM1 (Ile15, Ile19), TM3 (Phe78), and TM4 (Ala105, Val108, Met109) while the benzothiazole is oriented toward the CTD making primarily polar contacts (Fig. 4a, b). Fourth, the pyrazole ring is sandwiched between the walls of the cavity near Phe47 and the methylated nitrogen extends toward the outer face of the transporter. The substantial improvement in potency from **1** to **IMP-2380** may be due to a combination of factors, including van de Waals interactions with Phe47 and a desolvation effect from masking the free amine of the pyrazole. Previous studies have established a key role for deprotonation of Glu222 and Asp307 during coupling to substrate binding;^32^ notably, **IMP-2380** does not interact with these residues and is therefore unlikely to inhibit NorA via pH gradient modulation (Fig. 4a).

**Figure 4.**
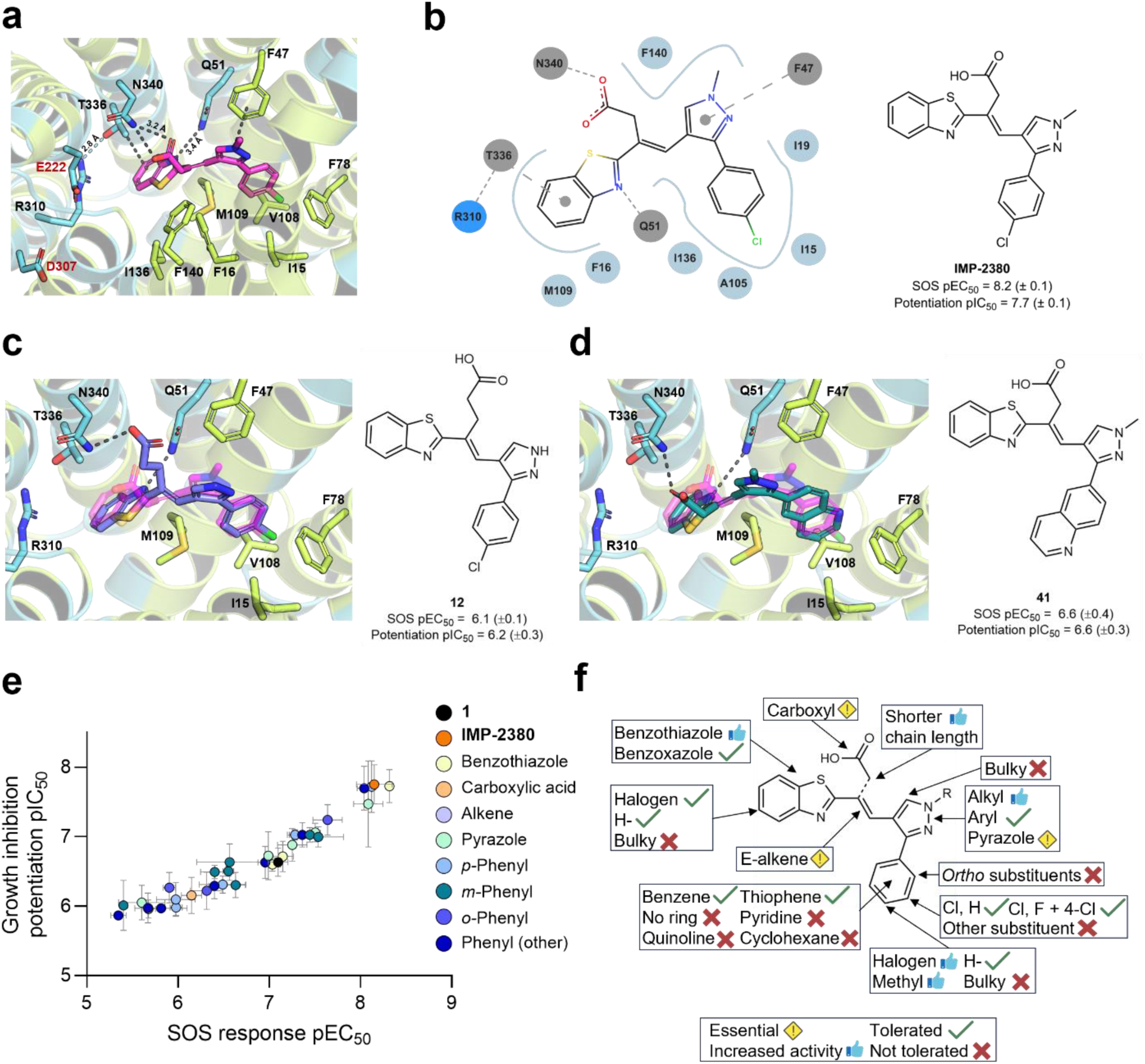
Expanded SAR series’ potency in SOS response and ciprofloxacin potentiation assay is corroborated by **IMP-2380**-NorA complex structure. (**a**) Cartoon representation of **IMP-2380** binding pocket. Key residues are depicted as sticks and labelled black. Residues involved in the protonation and energy coupling are labelled red. Hydrophobic residues/surfaces are shown in light green and hydrophilic residues/surfaces are shown in light blue. Parts of TM5 and TM7 are hidden for visibility of **IMP-2380**. (**b**) Key interactions between NorA and **IMP-2380**. Hydrogen bonds are indicated with dashed dark grey lines; π-π or C-H π interactions with dashed lines. Hydrophobic residues are shown as light green sticks, hydrophilic residues as light blue sticks. **IMP-2380** structure and activity provided for reference. (**c-d**) Overlay of docked analogues with **IMP-2380**-NorA structure. Docking was performed using Molsoft ICM-Pro. **IMP-2380** is shown as light pink sticks. Parts of TM5 and TM7 are hidden for visibility of compounds. (**c**) Overlay of **12** (blue sticks) in **IMP-2380**-NorA structure. The hydrogen bond interaction between the extended carboxylic acid chain and N340 is prioritized, causing the rest of the molecule to buckle out of position. (**d**) Overlay of **41** (green sticks) in **IMP-2380**-NorA structure. The larger quinoline ring causes the molecule to shift out of the small hydrophobic pocket. (**e**) Comparison of SOS response pEC_50_ to growth inhibition potentiation pIC_50_ (Pearson correlation r = 0.97, p <0.0001). Inactive compounds were excluded. Error bars show SD. (**f**) Summary of the conclusions from the SAR study.

We next expanded our SAR analysis to validate that structural interactions observed in the NorA-**IMP-2380** complex correlated with potentiation of ciprofloxacin-mediated growth inhibition and SOS response inhibition (Fig. 4a, Extended Data Fig. 8). We optimized two synthetic routes incorporating a key Knoevenagel condensation to install the *E*-alkene (Extended Data Fig. 9) and focused first on the core pyrazole and carboxylic acid motifs. A methyl ester analogue (**11**) of the carboxylic acid in **1** was not tolerated, whilst extending the spacer by one carbon (**12**) resulted in a 5-10 fold decrease in potency, consistent with the hypothesis that an optimal hydrogen bond interaction with Asn340 is crucial for binding to NorA (Fig. 4c). As predicted from the orientation of the pyrazole toward the outward-open face, an *N*-ethyl substituent (**13**) showed comparable activity to **IMP-2380**, with a bulkier phenyl (**14**) slightly less well accepted. Alterations at the 5-position of the pyrazole ring (**15**, **16**) resulted in a greater loss of activity, whilst a pyrazole regioisomer (**17**) which reduces cLogD by one unit was not tolerated, emphasizing the importance of this motif in a relatively hydrophobic pocket.

We next turned to the aromatic substituents which fill the two buried pockets in the NorA cavity. Exchanging the benzothiazole for a benzoxazole (**18**) caused a loss in activity, consistent with a weakened hydrogen bond to Gln51. Selectivity for 5-substituents on the benzothiazole ring showed a more subtle influence, with chloro (**19**) preferred over trifluoromethyl (**20**). The 4-chlorophenyl substituent in **IMP-2380** is positioned out of plane with the pyrazole, projecting into a relatively small hydrophobic pocket. The phenyl group is required for activity (**21**) and based on the cryo-EM structure we expected relatively few alterations to be tolerated. Removing the 4-chloro (**22**) caused a modest decrease in potency compared to **IMP-2380**, whilst small hydrophobic or electron withdrawing 4-substituents (**23**, **24**) showed comparable potency to **IMP-2380**. However, larger groups (**25-27**) and 2- or 3-substituents (**28-37**) were significantly less well tolerated. Replacement of the phenyl with pyridines (**38-40**), quinoline (**41,** Fig. 4d), or a cyclohexane (**42**) resulted in a significant loss of activity, with only a smaller thiophene substituent (**43-44**) moderately tolerated. Taken together, these results are consistent with the 4-chlorophenyl binding mode revealed in the cryo-EM structure, confirming the steric constraint in this part of the cavity (Fig. 4a, b). Furthermore, the broad correlation of SOS response data with the expected relative trends in activity predicted from the cryo-EM structure supports a link between pharmacological inhibition of NorA and inhibition of the SOS response (Fig. 4e, f, Extended Data Fig. 7b-c).

### NorA inhibition prevents induction of the ciprofloxacin-induced SOS response

Discovery of a NorA inhibitor series was unanticipated in the context of an HTS against the SOS response, leading us to further explore the connection between these phenomena. Activation of the SOS response induces auto-cleavage of transcriptional repressor LexA, instigating DNA repair, de-repression of genes in the SOS regulon and activation of prophages, resulting in horizontal gene transfer which can spread antibiotic resistance genes.^34^ **IMP-2380** reduced prophage induction upon activation of the SOS response in a ciprofloxacin-dependent manner (Fig. 5a), indicating that in this context NorA inhibition restricts activation of the physiological SOS response in bacteria.

**Figure 5.**
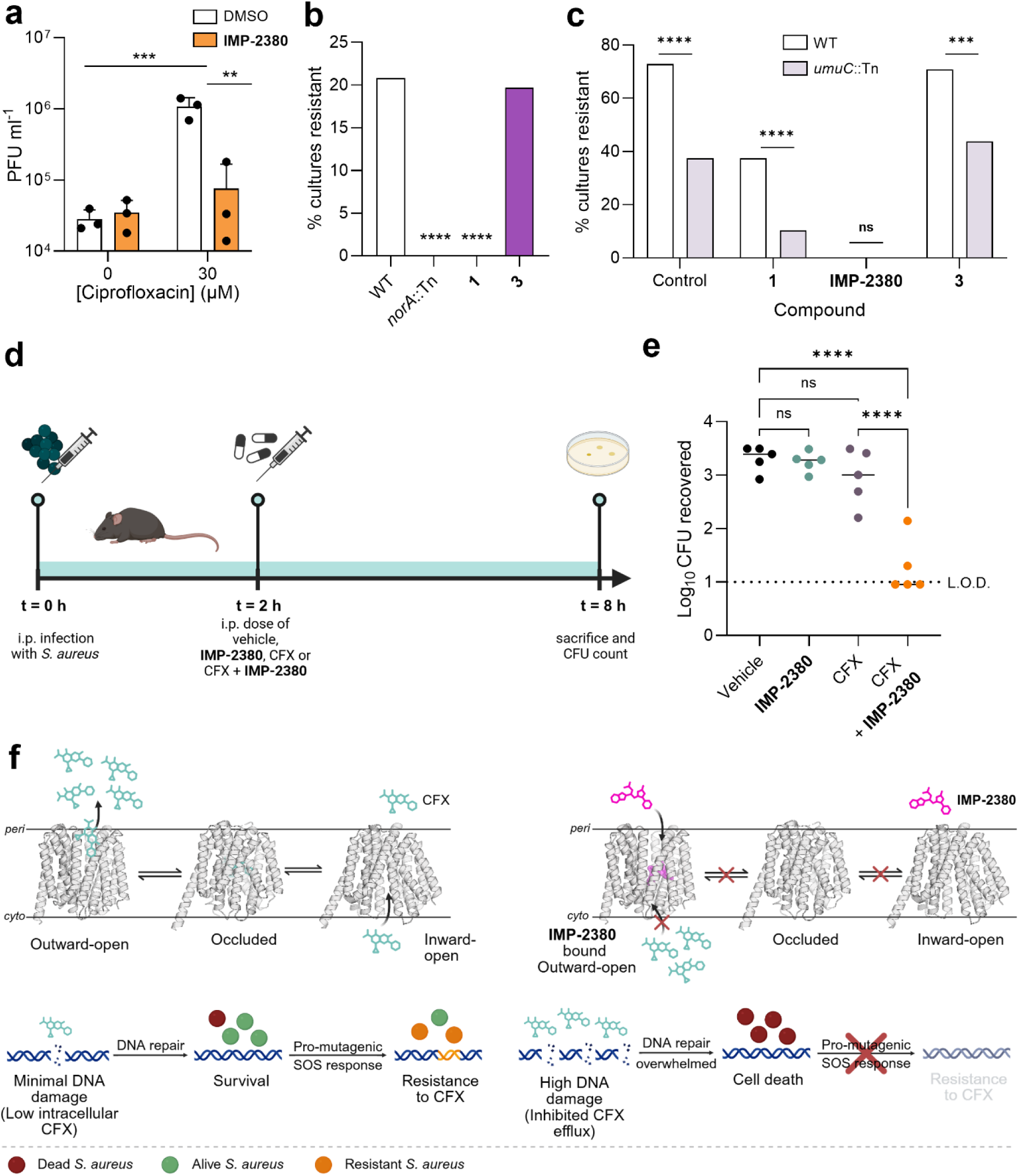
Chemical inhibition of NorA represses activation of the SOS response and potentiates CFX activity in MRSA infections *in vivo*. (**a**) Inhibition of prophage activation arising from induction of the SOS response. Plaque forming units (PFU) were determined following incubation *S. aureus* RN451 with 0-30 μM CFX ± 30 μM **IMP-2380** for 6 h. Data are shown as mean ± SD. (n = 3). Data were analyzed using a two-way ANOVA. (**b**) Resistance acquisition to CFX of *S. aureus* SH1000 WT, WT + **1**, WT + **3** or *norA*::Tn cultured with 3 μM CFX. Data were analyzed using a chi-squared test of independence. (**c**) Resistance acquisition to CFX of *S. aureus* SH1000 WT or *umuC*::Tn cultured with 3 µM CFX and either DMSO, **1**, **IMP-2380** or inactive control **3** (1 μM). Data were analyzed using a chi-squared test of independence. (**d**) Schematic for the *in vivo* MRSA infection model. (**e**) *S. aureus* USA300 JE2 recovered from the intraperitoneal cavity after 6 h treatment with vehicle (no treatment), 30 mg/kg CFX alone, 30 mg/kg **IMP-2380** or a combination of CFX and **IMP-2380**, both at 30 mg/kg (n = 5 in each group). Data were analyzed by a one-way ANOVA with Tukey’s post hoc test. Lines represent the median value of each group. (**f**) Left: NorA promotes the efflux of CFX from the cell, resulting in sub-inhibitory intracellular concentrations. CFX accesses the substrate pocket in the inward-open conformation, is cycled through the occluded state, before being extruded from the cell in the outward-open conformation. This means DNA damage is insufficient to cause cell death and allows sufficient time for the pro-mutagenic SOS response to be activated, resulting in the generation of CFX-resistant populations. Right: **IMP-2380** locks the pump in the outward-open conformation so that CFX cannot access the substrate pocket on the intracellular side of the pump. Thus, **IMP-2380** potentiates CFX activity by increasing its relative intracellular concentration, enhancing DNA damage and causing cell death. Cell death occurs before the SOS response can be activated, resulting in the suppression of resistance generation. Created in BioRender.

The mutagenic DNA repair program initiated by the SOS response occurs via the low-fidelity DNA polymerase UmuC and is known to increase the rate of spontaneous antibiotic resistance emergence.^17^ Whilst *norA* is not part of the SOS regulon, it has previously been reported that *norA* expression significantly promotes the rate of acquisition of quinolone resistance via chromosomal mutations,^35^ suggesting that NorA enables the cell to mount an effective SOS response by reducing antibiotic accumulation. We first confirmed that abrogation of NorA activity, either by transposon insertion or through chemical inhibition by **1,** blocked the emergence of ciprofloxacin resistance in WT *S. aureus* (Fig. 5b). We then compared the spontaneous emergence of ciprofloxacin resistance in WT *S. aureus* versus a *umuC:*:Tn mutant, which lacks the low-fidelity DNA polymerase responsible for mutagenic SOS DNA repair. The emergence of ciprofloxacin resistance was completely blocked by potent NorA inhibitor **IMP-2380** or significantly reduced by the less potent hit inhibitor **1**, whilst an inactive analogue had no effect (Fig. 5c). We also found that the SOS response makes a significant contribution to ciprofloxacin resistance, with ca. 50% lower frequency of resistance occurring with the *umuC*::Tn mutant relative to WT.

These data are consistent with a mechanism whereby the high intracellular ciprofloxacin concentration driven by NorA inhibition overwhelms and kills bacteria before the SOS response can be fully induced. NorA inhibition may therefore reduce ciprofloxacin resistance both by reducing the impact of resistance-conferring mutations through potentiation of ciprofloxacin, and by decreasing mutation rates by augmenting bacterial killing prior to activation of the SOS response.

### NorA inhibition by IMP-2380 potentiates ciprofloxacin *in vivo*

To investigate the potential of our compounds as *in vivo* tools to study the impact of NorA inhibition, we first evaluated the *in vitro* pharmacokinetic (PK) profiles of **1** and **IMP-2380**. Whilst both compounds exhibited good mouse microsomal stability and reasonable aqueous kinetic solubility (Extended Data Fig. 10a), **IMP-2380** exhibited significantly better permeability in a Madin-Darby canine kidney cell permeability assay. As **IMP-2380** showed superior potency and drug-like properties, it was advanced into an *in vivo* pharmacokinetic study in mice (Extended Data Fig. 10b). Following a single intraperitoneal (i.p.) dose of 10 mg/kg, after accounting for relatively high plasma protein binding (99.7%) the free drug concentration of **IMP-2380** in blood was estimated to be above the *in vitro* ciprofloxacin potentiation IC_50_ for 60 minutes (Extended Data Fig. 10c). Based on the assumption of PK dose linearity, we selected a single 30 mg/kg i.p. dose for an *in vivo* efficacy study, as the free drug concentration would be close to the IC_50_ for potentiation of CFX growth inhibition for up to 8 hours post-dosing.

We subsequently established a murine infection model of MRSA in which a combined intraperitoneal dose of **IMP-2380** and ciprofloxacin was compared to vehicle, **IMP-2380** or ciprofloxacin administered alone (Fig. 5d). The ciprofloxacin dose (30 mg/kg) was selected to mimic that used in humans.^36^ **IMP-2380** was well-tolerated, with no adverse events observed for any animals prior to termination of the study. Whilst **IMP-2380** or ciprofloxacin individually showed no significant decrease in bacterial burden compared to the vehicle, **IMP-2380** in combination with ciprofloxacin caused a significant 100-fold reduction in CFU count, with infections in three out of five animals being below the limit of detection of the assay (Fig. 5e). This demonstrates that **IMP-2380** is potent *in vivo*, and that the pharmacological inhibition of NorA is a viable therapeutic strategy in the treatment of fluoroquinolone-resistant infections.

## Discussion

Methicillin-resistant *S. aureus* (MRSA) is a major cause of mortality from drug-resistant infections, with routine surgeries and treatments increasingly threatened by nosocomial transmission.^1^ Drugs targeting antibiotic-resistance mechanisms or bacterial survival pathways may allow the use of existing drugs in combination therapy against multidrug resistant bacteria, thereby circumventing the challenge of identifying novel antibiotics.^37^ Efflux pumps expel antibiotics from bacterial cells and reduce intracellular concentrations to sub-inhibitory levels, representing a critical first line of defense in bacteria. Despite considerable interest in EPIs, none have so far progressed to clinical trials due to poor *in vivo* potency, toxicity, and lack of selectivity.^38^ NorA, a multidrug efflux pump of the major facilitator superfamily (MFS), is present in all MRSA strains.^26^ However, despite multiple studies, inhibitor development for NorA has been hindered by scaffolds with poor drug-like properties, off-target effects, and a lack of structural insight into inhibitor binding. Existing tool inhibitors include marketed drugs such as reserpine (a monoamine transporter inhibitor) and omeprazole (a proton pump inhibitor), which are weakly potent and show far greater activity against their human targets than against NorA. As such, current chemotypes are inadequate to validate NorA as a therapeutic target in MRSA combination therapy.^39,40^

In this study, we report the discovery of a novel EPI chemotype and binding mode for NorA. Starting from a phenotypic hit in a high-throughput screen for inhibitors of the ciprofloxacin-induced SOS response in MRSA, we used genetic, functional, and structural studies to unambiguously identify NorA as the target of this compound series. The lead compound, **IMP-2380**, is the most potent NorA inhibitor reported to date, exhibiting over 220-fold higher activity than reserpine and a lipophilic ligand efficiency (LLE) of 4.8 versus 2.4. Notably, NorA inhibition or *norA* knockout was found to block SOS induction and suppress both SOS-dependent and SOS-independent spontaneous resistance mutations to ciprofloxacin. **IMP-2380** potentiates ciprofloxacin activity potently against MRSA in a murine intraperitoneal infection model, demonstrating that selective NorA inhibition has the potential to deliver therapeutic benefit *in vivo*.

The first example of a small molecule inhibitor-bound structure of NorA identifies a novel mode of binding at atomic resolution, providing a new approach for structure-guided drug design against this target. This structure demonstrates that potent antibiotic potentiation can be achieved with a compound that binds on the periplasmic face of a single efflux pump. To our knowledge, this is the first example of a small molecule that can replicate the outward-open conformation previously only observed with Fabs.^10,11^ **IMP-2380** accesses a novel and more buried site that may be inaccessible to biologics yet achieves comparable potentiation of ciprofloxacin activity. Furthermore, the deep extracellular binding cavity exploited by these compounds is directly accessible from the external environment without requiring uptake into bacteria, thereby permitting physicochemical properties, such as a negatively charged carboxylic acid, that would typically hinder bacterial penetration but are essential for activity in this series. Together, these findings suggest a new paradigm for targeted inhibition of efflux pumps by small molecules.

Our findings establish a mechanistic link between efflux and bacterial stress responses. Beyond antibiotic sensitization, **IMP-2380** also prevents antibiotic-associated induction of the SOS response, a pathway implicated in the emergence of resistance. As SOS activation occurs in response to various antibiotic classes in both Gram-positive and Gram-negative bacteria, including *E. coli* and *P. aeruginosa*, this strategy may extend beyond *S. aureus*.^41^ Whilst it remains to be seen whether inhibition of other efflux pumps exerts similar effects on stress response pathways, our work suggests a previously unexplored therapeutic opportunity for antibiotic adjuvants: EPIs that lock an efflux pump in the outward-open conformation can resensitize bacteria to an approved antibiotic whilst preventing emergence of resistant strains without a stringent requirement to optimize bacterial membrane permeability, thereby circumventing one of the most challenging barriers to drug discovery in Gram-negative bacteria (Fig. 5f).

In summary, discovery of the first nanomolar, selective, *in vivo*-active, and drug-like chemical probe for NorA provides unprecedented structural insights into efflux pump inhibition and provides the first step towards validating NorA as a target for antibiotic adjuvant therapy, paving the way for a new class of adjuvants that restore antibiotic efficacy and suppress resistance emergence. We anticipate that **IMP-2380** will serve as a valuable tool compound for exploring the broader roles of NorA in antibiotic resistance and bacterial stress response.

## Methods

### Bacterial strains and culture conditions

The bacterial strains used in this paper are listed in Table S3. All *S*. *aureus* strains were grown in tryptic soy broth (TSB) at 37 °C with shaking at 180 rpm. Where appropriate, media were supplemented with kanamycin (154 µM) or erythromycin (14 µM). *Enterococcus faecalis* JH2-2, *Streptococcus agalactiae* COH1 were grown statically at 37 °C in 5% CO_2_ in Todd Hewitt broth with 2% yeast extract (THY) and *Escherichia coli* DH5α was grown in Luria Bertani broth (LB) at 37 °C with shaking at 180 rpm.

TSB (Becton-Dickenson 211822 or Millipore 22092). Antibiotics: Kanamycin (Sigma K1377 or 60615); erythromycin (Sigma E5389-5G); ciprofloxacin (Sigma 17850-5g-F).

### SOS HTS reporter assay general methods

*S. aureus* JE2 WT P*recA-gfp* (negative control for SOS inhibition) and *S. aureus* JE2 *rexB*::Tn P*recA-gfp* (positive control for SOS inhibition) were dispensed using a Multidrop Combi Liquid Dispenser (Thermo Scientific) with standard bore Multidrop tubing (Thermo Scientific, 24072670). Lines were washed with water, PBS, and TSB + kanamycin (154 µM) before priming with cultures, and washed with TSB + kanamycin (154 µM) between dispensing different strains. Cultures were diluted to specified OD_600_ dispensed into 384-well, black-walled, clear bottom plates (Greiner, 781091) and sealed with Breathe-Easy Gas Permeable Sealing Membranes (Diversified Biotech, BEM-1) prior to overnight growth at 37 °C with shaking. OD_600_ and fluorescence (excitation 485 nm; emission 525 nm) were recorded on a PHERAstar Plate Reader.

### HTS Assay

Test compounds (150 nL of 10 mM compound in DMSO) were dispensed into columns 1-22 using an ECHO 550 Acoustic Dispenser (Labcyte). DMSO (150 nL) was dispensed into high and low SOS control wells (columns 23 and 24, respectively). Overnight cultures were diluted to OD_600_ = 0.16 in TSB + kanamycin (154 µM) + CFX (48 µM). JE2 *rexB*::Tn P*recA-gfp* (50 µL) was inoculated into column 24, and JE2 WT P*recA-gfp* (50 µL) inoculated into columns 1-23. Plates were sealed, covered with plastic lids, secured in stacks of 4 plates and incubated at 37 °C with shaking (200 rpm) for 17 h before recording OD_600_ and fluorescence (excitation 485 nm; emission 525 nm).

The GFP/OD_600_ ratio was calculated and data analyzed using ActivityBase (IDBS). Robust Z’ values for each plate were calculated using the following equation:

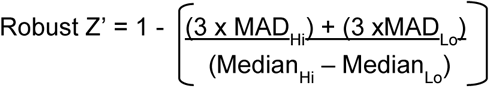

For quality control, data derived from assay plates with robust Z’ <0.5 or S/B <2 was rejected. The activity of each test compound was calculated as a percentage of the maximum effect, using the following equation:

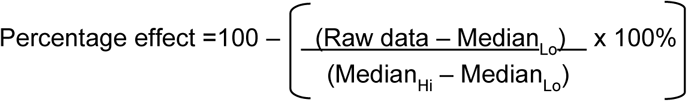

### Dose response assay

Overnight stationary phase cultures of *S. aureus* JE2 WT P*recA-gfp* were diluted to 10^8^ CFU mL^-1^ in TSB and dispensed into black-walled, flat-bottomed, 96-well plates (Grenier, 655090), with a final volume of 200 µL. Relevant wells were supplemented with CFX (97 µM) and either DMSO (0.1% v/v) or compounds in a dose-dependent manner. Plates were incubated in an Infinite M200-PRO or Tecan Spark Fusion microplate reader at 37 °C with shaking (700 rpm) for 17 h and fluorescence (excitation 475 nm; emission 525 nm) and OD_600_ determined every 15 min. For the SOS response inhibition EC_50_, the linear slope (t = 2.5-3.5 h) was calculated by plotting RFU/OD_600_ against time. The percentage inhibition of SOS response was determined by normalizing to the positive (CFX + DMSO) and negative (DMSO) control and then graphed against [compound]. EC_50_ values were determined using Prism 10 for each replicate, which were then used to calculate the geometric mean EC_50_ and associated errors.

For the potentiation of CFX growth inhibition IC_50_, the OD_600_ was plotted against time and area under the curve (AUC) for 0-8 h was calculated. Each concentration was normalized to the (CFX + DMSO) control, and data scaled from 0 to 100 using the 5-95% quantiles within each dose response. The percentage potentiation of CFX growth inhibition was then graphed against [compound] and the IC_50_ values determined using Prism 10 for each replicate. These were then used to calculate the geometric mean IC_50_ and associated errors.

### Checkerboard MIC assays

Synergy between compounds and antibiotics were determined using checkerboard MIC assays as described previously.^42^ A range of antibiotic or compound concentrations was generated in the relevant media (200 µL) by making 2-fold serial dilutions in the wells of 96-well clear bottom plates (VWR 734-2781). To assess potentiation, one plate was used to generate a concentration range of antibiotic across the plate and another range with a test compound was generated down another plate. Then, 100 µL of the diluted antibiotic from one plate was combined with the same volume of diluted compound from the second plate to produce a matrix of different concentrations of each compound. In the case of daptomycin, media were supplemented with 1.25 mM CaCl_2_.

The relevant bacteria (*S. aureus*, *E. faecalis*, *S. agalactiae* or *E. coli*) were inoculated into the wells to a final inoculum of 5 × 10^5^ CFU mL^-1^ before incubation statically at 37°C for 18 h. Bacterial growth after 18 h incubation was measured by obtaining OD_595_ measurements using a Bio-Rad iMark microplate absorbance reader (Bio-Rad Laboratories, USA). The MIC was defined as the lowest concentration at which there was no detectable growth by OD_600_ (mean of background OD_600_ ± 2 SD.).

To measure the ability of compound **1** and **IMP-2380** to potentiate ciprofloxacin in clinical isolates, two-fold serial dilutions of ciprofloxacin were prepared in 96-well plates in a final volume of 200 µL. Where appropriate, wells also contained 2 µM of compound **1** (n = 1) or 1 µM of **IMP-2380** (n = 2). Bacterial strains were inoculated to 5 x 10^5^ CFU/mL. Plates were incubated statically at 37 °C for 17 h and the MIC was determined as the lowest concentration that inhibited bacterial growth.

### Transposon mutagenesis to identify mechanism of action

Transposon mutagenesis was performed as described in Wang et al.^43^ Transposon libraries were plated onto TSA + 7 µM erythromycin, 1.2 µM ciprofloxacin and 0.25 µM compound **1** and incubated for 3 days at 37 °C. Two single colonies were picked, grown overnight and 2 mL cultures used for genomic DNA extraction (eluted in 200 µL water). 20 µL gDNA was digested with aciI for 3 h at 37 °C. Restriction enzyme was inactivated at 65 °C for 20 min. 18 µL digest reaction was ligated with T4 ligase overnight at room temperature in a final volume of 100 µL. DNA was purified using the QIAGEN PCR purification kit, DNA molecules containing the transposon were amplified by PCR (30 cycles, annealing temp 63 °C, extension time 2 min) using Martn_F and Martn_R^44^ and sequenced by Eurofins genomics using Martn_F.

### Transduction of *norA*::Tn into SH1000

The *norA*::Tn insertion was transduced from JE2 *norA*::Tn (from the Nebraska Transposon Mutant Library^45^) into SH1000 by phage transduction with φ11. TSB (5 mL) supplemented with 5 mM CaCl_2_ was inoculated with 100 µL overnight culture of the JE2 *norA*::Tn mutant and incubated for 3 h at 37 °C with shaking. 10-fold serial dilutions of φ11 lysate were prepared in TMG buffer (10 mM Tris-HCl (pH 7.5), 10 mM MgSO_4_, 0.1 % gelatin). φ11 lysate (100 µL) was mixed with 500 µL bacteria and incubated at room temperature for 30 min before 5 mL top agar (0.8% agar, 0.8% NaCl) was added and poured over agar plates and incubated overnight before phage were harvested in TMG buffer. Overnight cultures of SH1000 were concentrated 10-fold in TSB supplemented with 5 mM CaCl_2_ before 250 µL bacteria was mixed with 200 µL phage lysate and incubated for 20 min at 37 °C. Phage infection was stopped by three washes in cold 20 mM sodium citrate before plating onto TSA supplemented with 20 mM sodium citrate and 10 µg/mL erythromycin. Successful disruption of *norA* with the transposon was confirmed by PCR (Applied Biosciences 2720 Thermal Cycler).

### Ethidium bromide efflux assay

100 µL bacterial overnight culture was inoculated into 10 mL TSB and incubated for 2 h at 37 °C with shaking (180 rpm). Bacteria were incubated for 20 min with 25 µM ethidium bromide (Merck E1510) at 41 µM reserpine (Sigma) at 37 °C with shaking (180 rpm). Bacteria were washed twice in PBS and concentrated 10-fold before 20 µL was inoculated into 200 µL TSB + various concentrations of compound. Fluorescence (excitation 525 nm; emission 605 nm) was measured every min (Infinite M200-PRO microplate reader (Tecan)) for 20 min. Background was subtracted at t = 0, and the data scaled from 0 to 100 within each dose response. Normalized RFU was plotted against [compound], and IC_50_ values were determined in Prism 10 for each replicate. These were then used to calculate the geometric mean IC_50_ and associated errors.

### BAG2 Fab expression and purification

BAG2, a Fab developed to bind the BRIL domain, was expressed and purified as previously described.^31^ Briefly, BAG2 was expressed in the 55244 strain of *E. coli* (ATCC) in TBG medium for 22 h at 30 °C. Cell pellets were collected by centrifugation and resuspended in running buffer (20 mM sodium phosphate pH 7.0) supplemented with 1 mg/mL chicken egg white lysozyme. After passing through an EmulsiFlex-C3 high pressure homogenizer five times, cell debris was removed by centrifuging for 1.5 h at 15,000 rpm using a Beckman centrifuge equipped with a JA 25.50 fixed-angle rotor. The supernatant was loaded onto a protein G column (GE Healthcare) equilibrated in running buffer. BAG2 was eluted from the column with 100 mM glycine (pH 2.7) and immediately neutralized with 2 M Tris buffer (pH 8). Peak fractions were pooled and used immediately for cryo-EM.

### Anti-kappa VHH-domain expression and purification

The gene encoding the anti-kappa VHH-domain (VHH) with a hexahistidine tag at the N-terminus was purchased (Twist Bioscience) and subcloned into a pET29b(+) expression vector.^46^ VHH was expressed in *E. coli* BL21 (DE3). Cells were grown at 37 °C in ZY medium supplemented with 100 µg/mL kanamycin. At an OD_600_ of 0.7, cells were induced with 1 mM isopropyl β-D-1-thiogalactopyranoside (IPTG) and the temperature was reduced to 20 °C and grown for an additional 18 to 20 h. The culture was harvested by centrifugation and stored at -80 °C.

Cell pellets were resuspended in 20 mM Tris pH 7.5, 150 mM NaCl, and the resultant lysate was centrifuged for 20 min at 24,500 rpm using a Beckman centrifuge equipped with a JA 25.50 fixed-angle rotor. VHH was purified using immobilized metal-affinity chromatography using a Ni-NTA affinity resin (ThermoFisher Scientific) and successively washed with chromatography buffer (20 mM Tris pH 7.5, 150 mM NaCl) containing 10 mM imidazole. VHH was eluted using a 25 to 50% gradient of the chromatography buffer containing 600 mM imidazole. VHH elution fractions were dialyzed into SEC buffer (20 mM Tris pH 7.5 and 150 mM NaCl) and subsequently treated with TEV protease to remove the N-terminal hexahistidine tag. VHH was then purified using a Superdex 75 10/300 column (Cytiva) equilibrated in SEC buffer. Peak fractions were pooled, concentrated with a 10 kDa centrifugal concentrator (Millipore), and used immediately.

### NorA-BRIL^3A^ expression and purification

NorA-BRIL^3A^ expression and purification was performed as previously described.^33^ Briefly, NorA-BRIL^3A^ was expressed in *E. coli* C43 (DE3) using autoinduction by growing bacteria at 32 °C in ZYP-5052 medium supplemented with 1 mM MgSO_4_.^47^ At an OD_600_ of 0.5, the temperature was reduced to 20 °C, and cultures were grown for an additional 18 to 20 h. Cells were collected by centrifugation and stored at -80 °C.

Cell pellets were resuspended in 40 mM Tris pH 8.0, 400 mM NaCl, and 10% glycerol and passed through an EmulsiFlex-C3 high pressure homogenize five times. The resultant lysate was centrifuged for 30 min at 13,000 x g using a Beckman centrifuge equipped with a JA 25.50 fixed-angle rotor. Next, the supernatant was centrifuged for 2.5 h at 40,000 rpm using a Beckman ultracentrifuge equipped with a Type 45 Ti fixed-angle rotor to isolate the membrane fraction. The pellet was resuspended in 20 mM Tris pH 8.0, 300 mM NaCl, 10% glycerol, 10 mM imidazole, and 1% (w/v) lauryl maltose neopentyl glycol (LMNG, Anatrace). The membrane fraction was vigorously stirred for 2.5 h at 4 °C. Next, the sample was centrifuged for 30 min at 40,000 rpm using a Beckman ultracentrifuge equipped with a Type 45 Ti fixed-angle rotor to remove the insoluble fraction. The supernatant containing NorA-BRIL^3A^ was passed over a Ni-NTA affinity resin (ThermoFisher Scientific) and subsequently washed with buffer (20 mM Tris pH 8.0, 200 mM NaCl, 10% glycerol, and 0.2% (w/v) LMNG) containing 25 mM imidazole and eluted with the same buffer containing 400 mM imidazole. NorA-BRIL^3A^ elution fractions were dialyzed into SEC buffer (20 mM Tris pH 7.5, 10 mM NaCl) in the presence of TEV protease to remove the carboxy-terminal decahistidine tag.

For cryo-EM, cleaved NorA-BRIL^3A^ was incubated with PMAL-C8 amphipol (Anatrace) at a ratio of 1:5 NorA-BRIL^3A^:amphipol (w/w) overnight at 4 °C. Following the overnight treatment, Bio-beads (Bio-Rad) were added to the sample at a ratio of 100/1 (w/w) relative to LMNG for 24 h at 4 °C. Finally, the sample was concentrated and purified on a Superdex 200 10/300 column equilibrated in SEC buffer (20 mM Na_2_HPO_4_ pH 5, 150 mM NaCl). Peak fractions were pooled, concentrated with a 10 kDa centrifugal concentrator (Millipore), and used immediately for cryo-EM.

### Cryo-EM sample preparation and data collection

Amphipol-purified NorA-BRIL^3A^ was incubated with a 3-fold molar excess of BAG2 for 30 min at 4 °C. Following incubation, the NorA-BRIL^3A^-BAG2 complex was purified using a Superdex 200 10/300 column in SEC buffer. The most concentrated SEC fractions were treated with 2-fold molar excesses of **IMP-2380** and VHH relative to the NorA-BRIL^3A^-BAG2 complex immediately before cryo-EM grid freezing.

Cryo-EM grids were prepared by applying 4 µL of the sample (50 μM) to glow-discharged UltrAuFoil 300-mesh R1.2/1.3 grid (Quantifoil). The sample was blotted for 3.5 s under 100% humidity at 16 °C before plunge-freezing into liquid ethane using a Mark IV Vitrobot (ThermoFisher Scientific).

Cryo-EM data for NorA-BRIL^3A^ in complex with BAG2, VHH, and **IMP-2380** was acquired on a Titan Krios microscope (ThermoFisher Scientific) at 300 kV equipped with a K3 direct electron detector (Gatan) using a GIF-Quantum energy filter with a 15-eV slit width. Leginon 3.6 was used for automated data collection of squares with optimal ice thickness.^48–50^ Movies were collected at a nominal magnification of 105,000x in SuperRes mode with a physical pixel size of 0.4125 Å and dose fractioned over 40 frames. The NorA-BRIL^3A^- BAG2-VHH-**IMP-2380** complex dataset received an accumulated dose of 47.12 e^-^/Å^−2^. A total of 13,288 movies were collected.

### Cryo-EM image processing and model building

Movies of the NorA^3A^-BRIL-BAG2-VHH-**IMP-2380** complex were processed in cryoSPARC v4.4.1.^51^ Imported movies were motion corrected, and the contrast transfer function estimated. Initial 2D class averages were generated using particles picked from pretrained Topaz model (ResNet8 – 32 units). Particles in 2D classes representing NorA^3A^-BRIL-BAG2-VHH-**IMP-2380** complex were used as templates for Topaz picking.^52,53^ A total of 8.28 million particles were selected. Particles underlying well-resolved 2D classes were used for initial *ab initio* model building, and all picked particles were used for subsequent heterogeneous 3D refinement. Particles from classes corresponding to NorA^3A^-BRIL-BAG2-VHH-**IMP-2380** complex or complexes apparently lacking BAG2 and VHH domain were retained for subsequent rounds of *ab initio* model building and heterogeneous 3D classification. After multiple rounds, a non-uniform 3D refinement step was used to generate final 2.47 Å resolution map for NorA^3A^-BRIL-BAG2-VHH-**IMP-2380** complex from 634,207 particles, as assessed using the gold standard Fourier shell correlation (FSC). The map was further processed using local refinement with a mask for NorA. The final local refinement map was obtained with a resolution of 2.52 Å. The final cryo-EM map was sharpened using Phenix.auto_sharpen or 3D model building.^54^

The structural models were constructed using Namdinator^55^ and Coot^56^, and refined using Phenix^54^ and Coot. Structures of NorA (PDB ID: 7LO8) was used as reference for model building.

### Prophage activation

Overnight cultures of *S. aureus* RN451^57^ were diluted 10-fold in TSB containing 30 µM ciprofloxacin ± 30 µM **IMP-2380** and incubated at 37 °C with shaking (180 rpm) for 6 h. Samples were centrifuged at 13,000 x g for 1 min and the supernatant filtered through a 0.2 µm filter. Plaque forming units (PFU) per mL were determined by spotting serial dilutions of the supernatant onto 0.8% agar plates containing RN4220. Data was analyzed by a two-way ANOVA in Prism 10.

### Spontaneous CFX resistance acquisition

TSB (200 µL) containing 2X SH1000 WT MIC ciprofloxacin (3 µM) and compound **1**, **IMP-2380** or inactive control compound **3** (1 µM) was added to the wells of a 96-well plate (VWR, 734-2781) and inoculated with 5 x 10^5^ CFU/mL bacteria. After 72 h incubation at 37 °C, the number of wells with visible growth (indicative of resistance emergence) were enumerated and calculated as a percentage of total wells inoculated. Data were analyzed using a chi-squared test of independence in Prism 10.

### *In vivo* efficacy

Animal work was conducted in accordance with the Animals (Scientific Procedures) Act 1986 and approved by the Imperial College Animal Welfare and Ethical Review Body (AWERB) (PPL70/7969, PF93C158E).

Female C57BL/6 mice (6-8 weeks, Envigo) were infected via the intraperitoneal route with 2.5 × 10^7^ CFU *S. aureus* USA300 JE2. Two hours post infection, mice were treated with vehicle alone (no treatment), a 30 mg/kg dose of **IMP-2380** (200 µL, 3 mg/mL), a 30 mg/kg dose of ciprofloxacin (200 µL, 3 mg/mL) or both ciprofloxacin and **IMP-2380** combined, each at 30 mg/kg. The dose of ciprofloxacin used was chosen to mimic that used in humans.^58^ Treatments were administered in a mixture of 5% DMSO, 40% PEG400 and 55% sterile deionised water via the intraperitoneal route. Eight hours post infection, animals were humanely killed by cervical dislocation, and death was confirmed by severing the femoral artery. The peritoneal cavity was washed with PBS, and CFU counts were determined on tryptic soy agar plates.

The size of the groups (n = 5) used was based on power analysis of *in vitro* data. Mice were randomly assigned to treatment groups and the investigators conducting the experiment were blinded to the treatments administered.

## Supplementary information

Supplementary Figs. 1-6, Tables 1-3, additional biological methods (mammalian cell toxicity, docking, *in vitro* and *in vivo* pharmacokinetics), and synthetic procedures and characterization.

## Acknowledgements

We would like to thank José Penadés (Imperial College London) and Timothy Meredith (Merck) for providing bacterial strains, and Sandra O’Neil and Barbara Forte (University of Dundee) and Ravi Singh (Imperial College London) for assistance and discussions. We acknowledge the provision of strains by the Network on Antimicrobial Resistance in *Staphylococcus aureus* (NARSA) Program: under NIAID/NIH Contract No. HHSN272200700055C. We thank A. T. Sahin and U. Zachariae (University of Dundee) for their preliminary contributions to modelling. Cryo-EM datasets were collected at the Cryo-Electron Microscopy Facility of the NYU School of Medicine. Schematic figures were generated using BioRender.com. Funding sources included the following: Rosetrees Trust (ID2020\100014) to A.M.E and E.W.T; EPSRC Impact Acceleration Award (EP/R511547/1) to A.M.E and E.W.T; MRC Impact Acceleration Award (MR/X502959/1) to A.M.E and E.W.T; National Institutes of Health (NIH) grant (R01 AI165782) to N.J.T.

## Contributions

A.M.E and E.W.T carried out conceptualization. J.L.G, E.V.K.L, T.S., T.L-H, T.J.B, K.A, L.E.P, T.B.C, J.R and E.G.P. performed experiments. J.L.G, E.V.K.L, T.J.B, A.S, F.C, I.H.G., D.G, D.N.W, K.D.R, N.J.T, A.M.E and E.W.T supervised the research. J.L.G, E.V.K.L, T.L-H, D.N.W, N.J.T, A.M.E and E.W.T secured funding. J.L.G and E.W.T were responsible for writing the original draft. All authors participated in the writing, review and editing.

## Ethics

J.L.G, E.V.K.L, T.J.B, A.M.E and E.W.T are inventors on a patent related to this work (GB2319181.0) filed by Imperial College Innovations Ltd on 14^th^ December 2023. E.W.T. is a founder and shareholder in Myricx Bio and Siftr Bio. Other authors declare no conflict of interest.

## Extended Data Figures

**Extended Data Figure 1.**
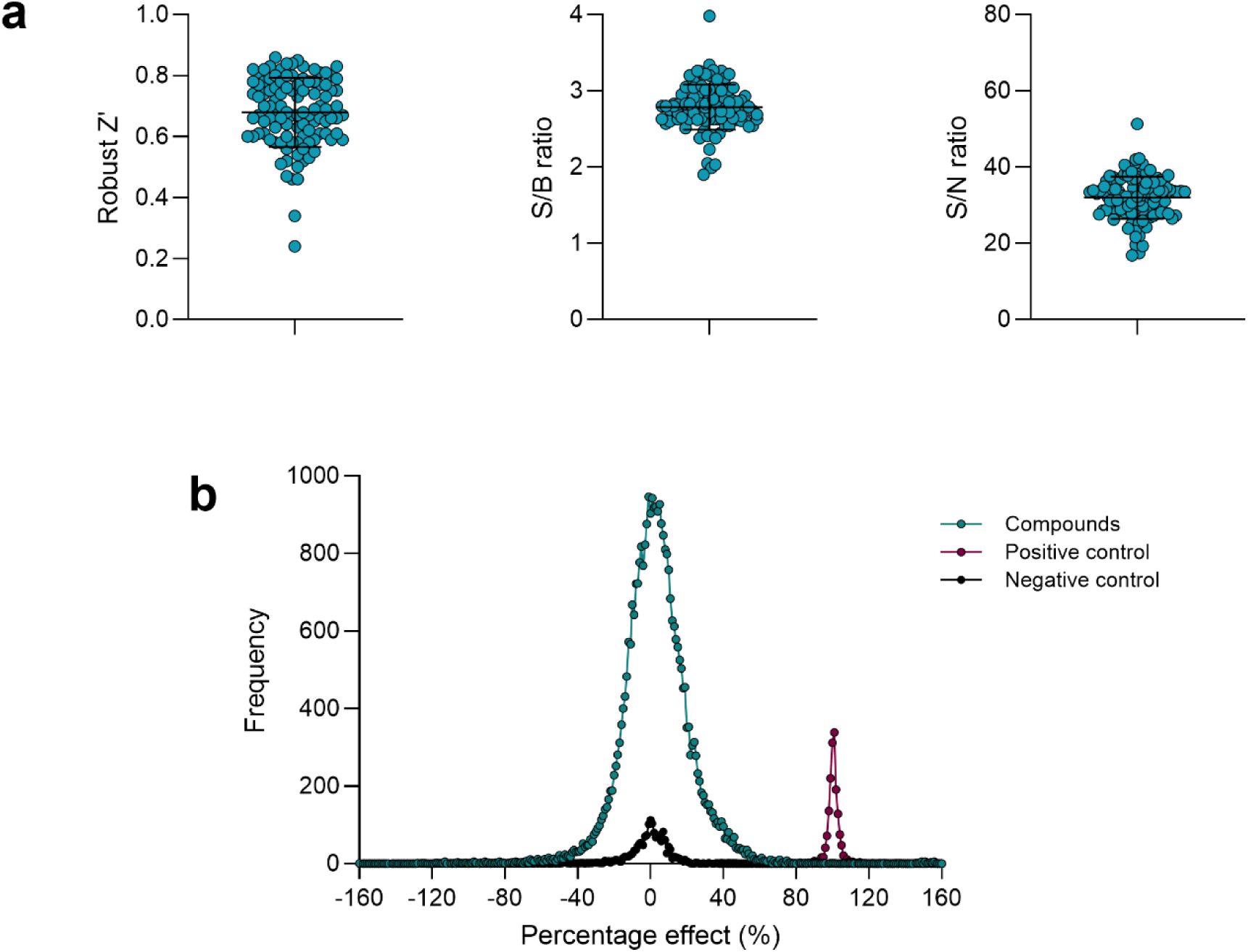
HTS analysis. (**a**) Assay metrics for SOS response HTS. Error bars represent mean ± SD. Data are based on screen of 106 384-well plates. Robust Z’, S/B ratio and S/N ratio calculated using positive control (RFU/OD_600_ signal for *S.aureus* USA300 JE2 *rexB*::Tn + 48 μM CFX + 0.3% DMSO), negative control (RFU/OD_600_ signal for *S.aureus* USA300 JE2 + 48 μM CFX + 0.3% DMSO) and blank (media only). Data derived from assay plates with a robust Z’ < 0.5 or S:B < 2 was rejected. (**b**) Frequency distribution of compound activities compared to positive and negative controls in the HTS. The positive control was RFU/OD_600_ signal for *S.aureus* USA300 JE2 *rexB*::Tn + 48 μM CFX + 0.3% DMSO, whereby the SOS response is blocked due to lack of double strand break processing. The negative control was the maximal SOS response activation (RFU/OD_600_) signal for *S.aureus* USA300 JE2 + 48 μM CFX + 0.3% DMSO. Hits were classified as inducing > 60% percent reduction in signal compared to the maximal effect from the positive control.

**Extended Data Figure 2.**
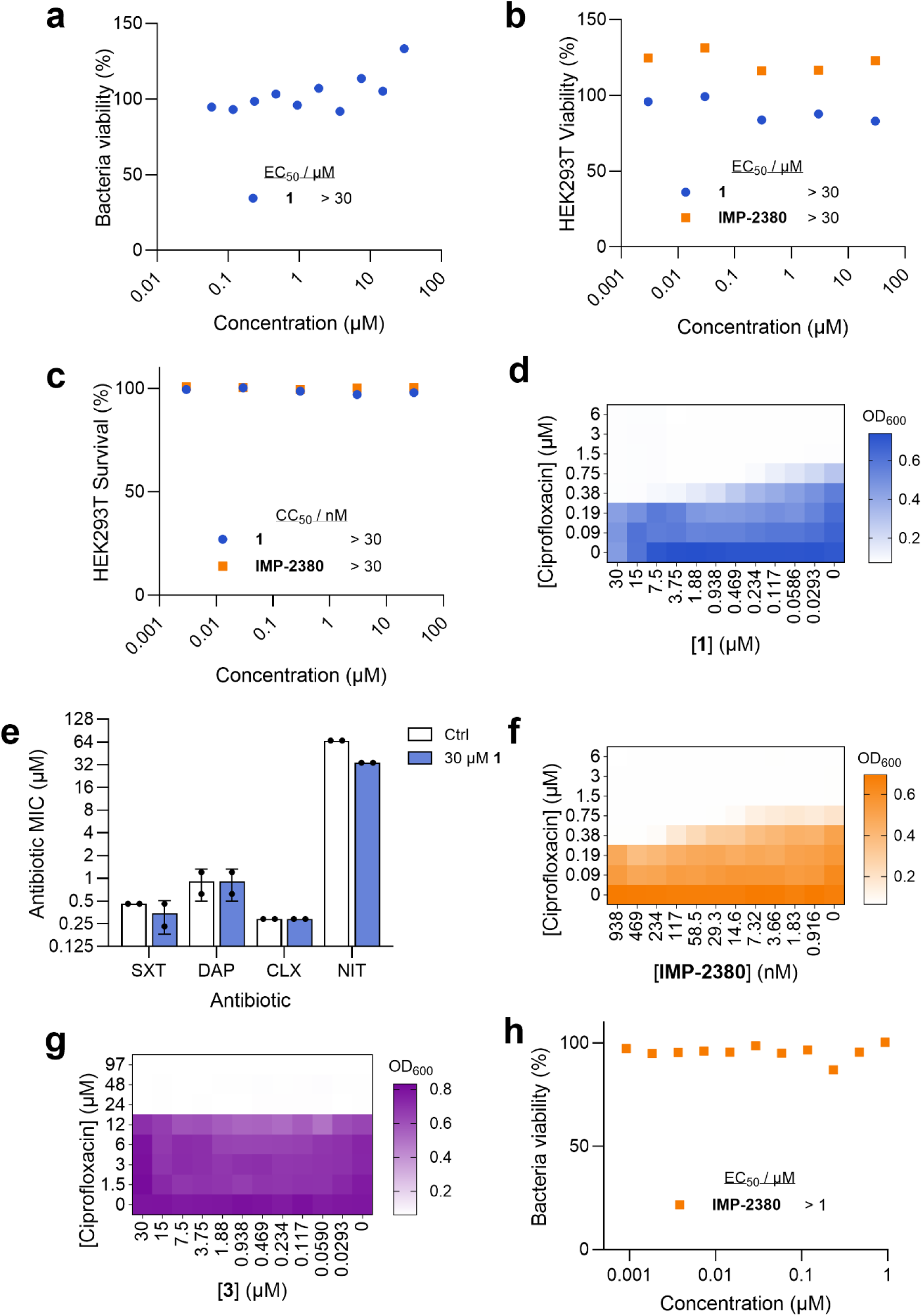
Validation of **1** and **IMP-2380** as selective SOS response inhibitors. (**a**) Dose-response curve for inhibition of *S. aureus* USA300 JE2 growth induced by **1** in the absence of CFX (n = 2). Data shown are of a single replicate. (**b**) Dose-response curve for inhibition of HEK293T proliferation by **1** or **IMP-2380** (n = 3). Data shown are of a single replicate. (**c**) Dose-response curve for HEK293T cytotoxicity in response to **1** or **IMP-2380** (n = 3). Data shown are of a single replicate. (**d**) Checkerboard MIC assay in *S. aureus* SH1000 of **1** and ciprofloxacin (n = 2). (**e**) Co-trimoxazole (SXT, n = 2), daptomycin (DAP, n = 2), cloxacillin (CLX, n = 2 and nitrofurantoin (NIT, n = 2) MIC in *S. aureus* USA300 JE2 ± 30 μM **1**. The full checkerboard MIC assay is shown in Fig. S3. (**f**) Checkerboard MIC assay in *S. aureus* SH1000 of ciprofloxacin and **IMP-2380** (n = 1). (**g**) Checkerboard MIC assay in *S. aureus* USA300 of ciprofloxacin and inactive control **3** (n = 2). (**h**) Dose-response curve for inhibition of *S. aureus* USA300 JE2 growth induced by **IMP-2380** in the absence of CFX (n = 3). Data shown are of a single replicate.

**Extended Data Figure 3.**
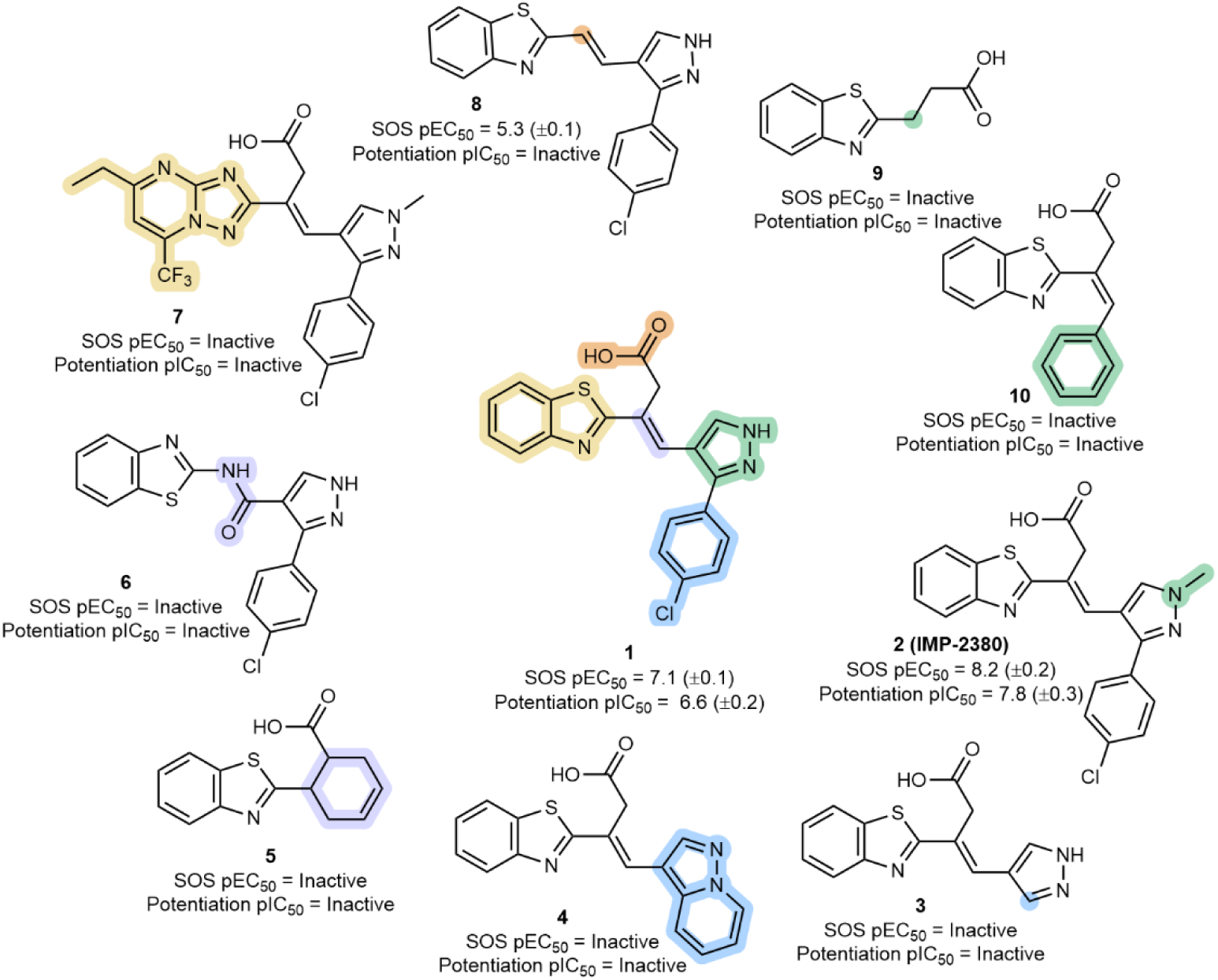
Initial SAR analysis yielded potent lead **IMP-2380**. Functional groups benzothiazole, carboxylic acid, alkene, pyrazole and 4-chlorophenyl that are altered in analogues are highlighted in yellow, orange, purple, green and blue, respectively. The exact stereochemistry of analogues **3**, **4**, **8** and **10** was not determined.

**Extended Data Figure 4.**
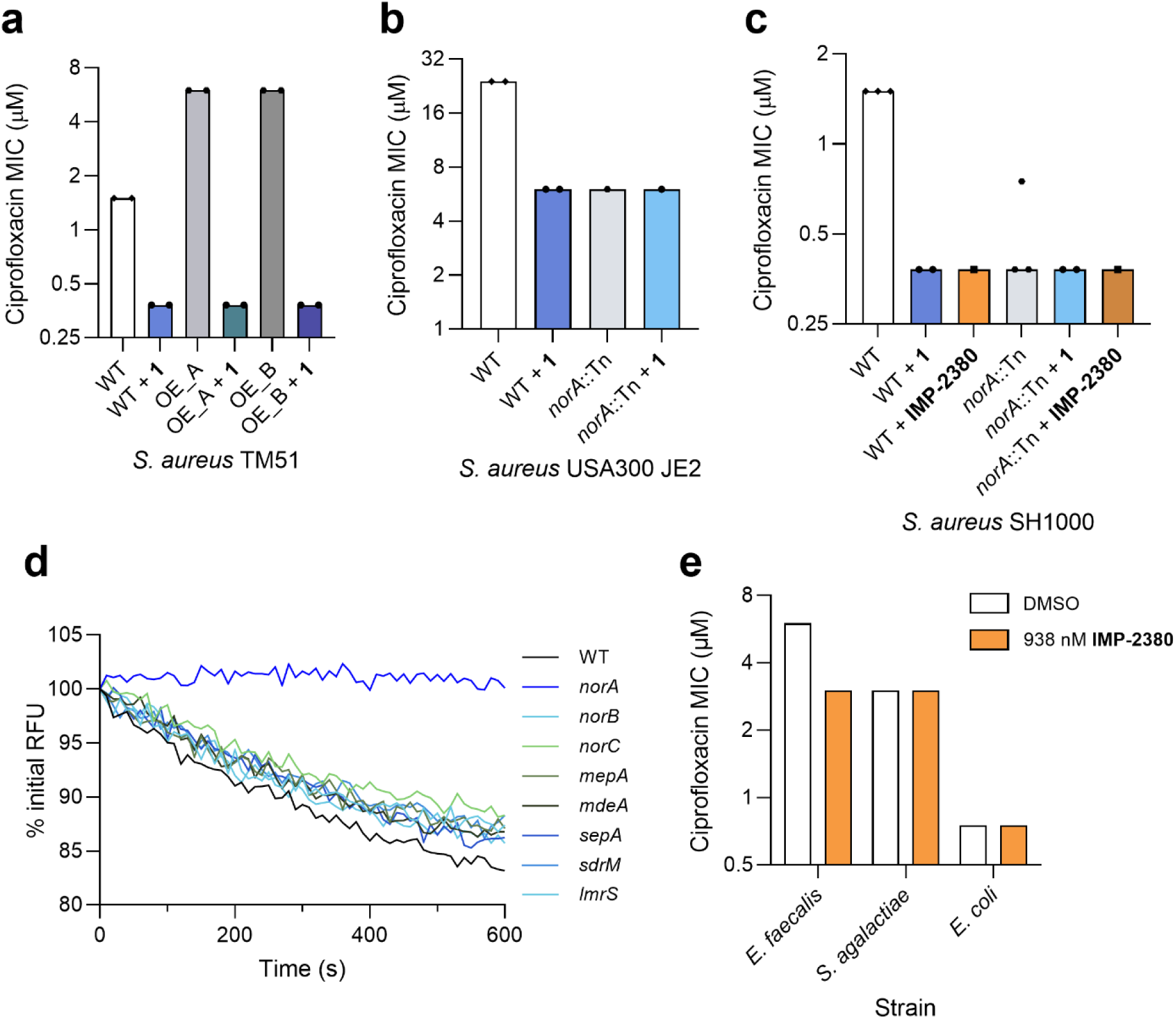
Additional validation of NorA as the cellular target of **1** and **IMP-2380**. (**a**) Ciprofloxacin MIC in *S. aureus* TM51 WT and two independent *norA* overexpression mutants OE_A and OE_B ± 32 μM **1** (n = 2). The data represent the median MIC. (**b**) Ciprofloxacin MIC in *S. aureus* USA300 JE2 WT (n = 2) and *norA*::Tn mutant ± 30 μM **1** (n =1). The data represent the median MIC. (**c**) Ciprofloxacin MIC in *S. aureus* SH1000 WT and *norA*::Tn mutant ± 3.75 μM **1** (n = 2) or 0.938 μM **IMP-2380** (n = 1). The data represent the median MIC. (**d**) EtBr accumulation assay in *S. aureus* JE2 WT and transposon mutants each defective for a single efflux pump. (**e**) Ciprofloxacin MIC in *Enterococcus faecalis* JH2-2 (n = 1), *Streptococcus agalactiae* COH1 (n = 1) and *Escherichia coli* DH5α (n = 1) in response to 0.938 μM **IMP-2380**. The full checkerboard MIC assays are shown in Fig. S4-6.

**Extended Data Figure 5.**
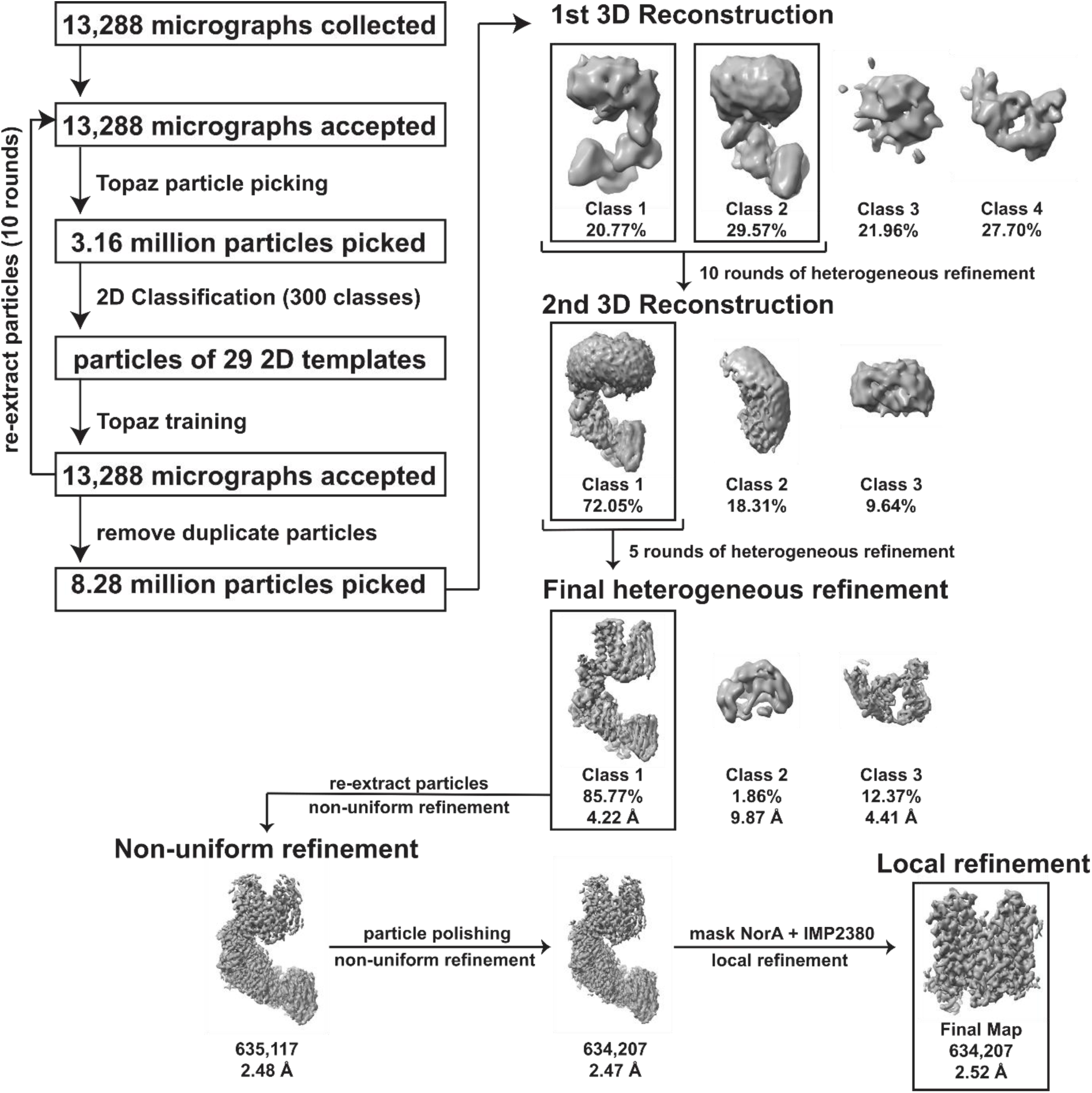
This schematic outlines the key steps involved in the cryo-EM data processing workflow utilized to determine the Coulomb potential map of the NorA^3A^-BRIL-BAG2-VHH-**IMP2380** at pH 7.5. Solid boxes are employed to highlight the intact NorA^3A^-BRIL-BAG2-VHH-**IMP2380**. Each 3D class is annotated with the number of constituent particles and the map resolution. The percentages indicated in the figure represent the fraction of particles chosen for subsequent refinement stages in relation to the initial total particle count.

**Extended Data Figure 6.**
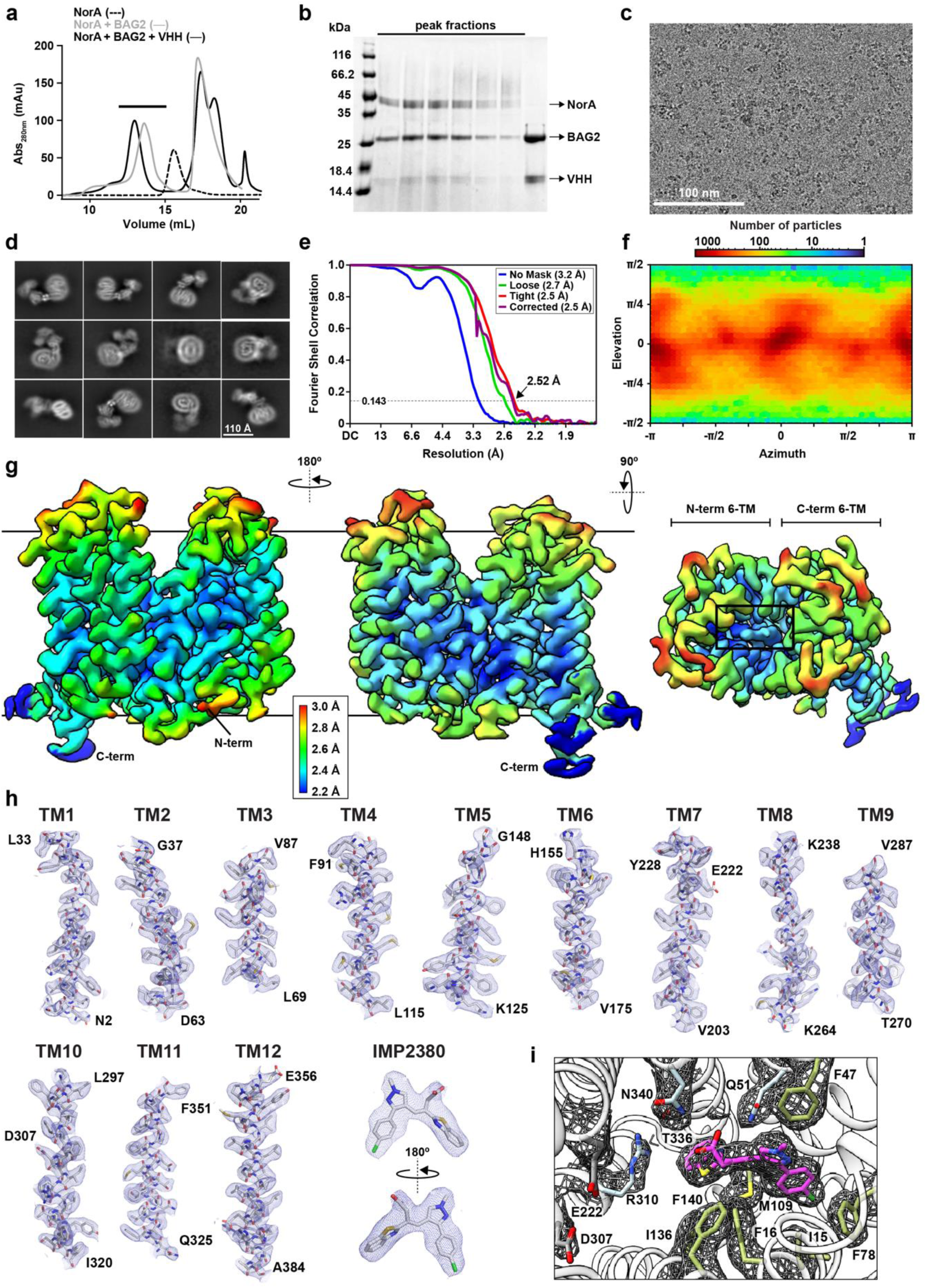
(**a**) SEC chromatogram depicting NorA reconstituted in PMAL-C8 amphipol alone (black), NorA in the presence of three-fold molar excess of BAG2 (gray), and NorA in the presence of three-fold molar excess of BAG2 and two-fold molar excess of anti-kappa VHH domain (blue). (**b**) Coomassie-stained SDS PAGE gel displaying peak fractions obtained from the corresponding SEC chromatogram (a). (**c**) Representative cryo-EM micrograph for the NorA^3A^-BRIL-BAG2-VHH-**IMP2380** complex. (**d**) Exemplary cryo-EM 2D classes of the NorA^3A^-BRIL-BAG2-VHH-**IMP-2380** complex. **(e)** Fourier shell correlation (FSC) curves for the final NorA-**IMP-2380** reconstruction. An arrow indicates the map resolution at the gold standard FSC value of 0.143. **(f)** Orientation distribution heat map for NorA^3A^-BRIL-BAG2-VHH-**IMP-2380** particles. **(g)** Three views of the local resolution cryo-EM map of NorA^3A^-BRIL-BAG2-VHH-**IMP-2380** complex (left and middle) with a top-facing view of **IMP-2380** bound to NorA (right). **IMP-2380** electron density is indicated by the black rectangle. **(h)** The quality of the NorA-**IMP-2380** structural model is depicted through their agreement with the cryo-EM maps (in a blue mesh). The model-to-map fitting is displayed for each transmembrane (TM) helix within NorA and **IMP-2380**. The map contour levels were set to 7s for each TM helix and IMP2380 using the isomesh command in PyMOL. Each TM helix is defined by the following residues: TM1, 2-33; TM2, 37-63; TM3, 69-87; TM4, 91-115; TM5, 125-148; TM6, 155-175; TM7, 203-228; TM8, 238-264; TM9, 270-287; TM10, 297-320; TM11, 325-351; TM12, 356-384. **(i)** Assessment of model-to-map fitting quality showing a cartoon representation of **IMP-2380** in the binding pocket of NorA. Indicated residues are depicted as sticks and labelled black with the superimposed cryo-EM map in a black mesh. A similar structural view is displayed in Fig. 4a of the main text.

**Extended Data Figure 7.**
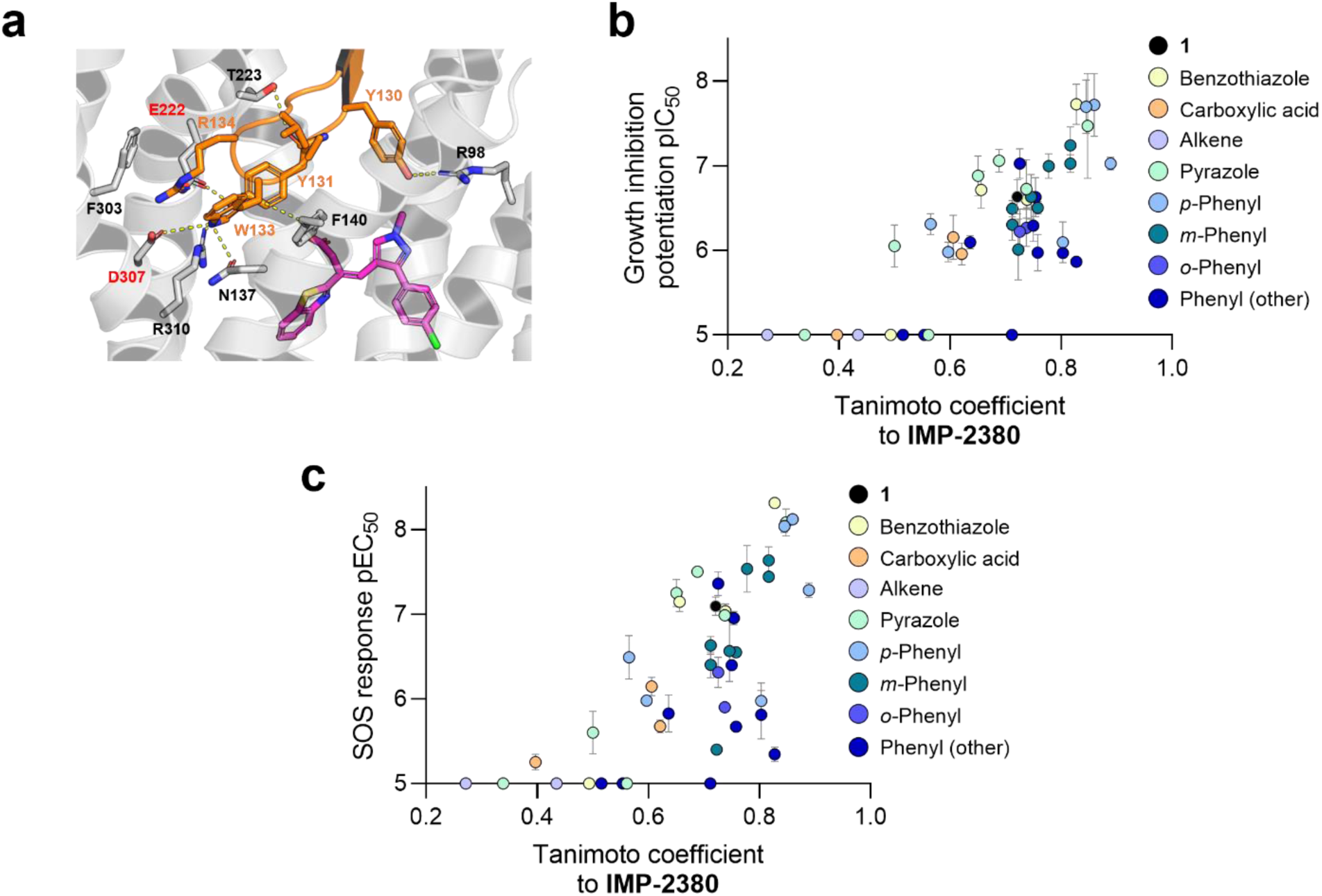
(**a**) Comparison of **IMP-2380** binding site compared to Fab36-bound NorA (PDB: 7LO8). Fab36 (orange cartoon and sticks) forms interaction at the top of the pocket in the outward-open conformation, including with residues involved in the proton-coupled transport (labelled in red). **IMP-2380** accesses a deeper portion of the pocket with different residues. (**b**) Correlation analysis between similarity to **IMP-2380** (Tanimoto coefficient) and growth inhibition potentiation pIC_50_ (Spearman correlation r = 0.69, p <0.0001). Inactive compounds set as pIC_50_ = 5, the maximum concentration tested in the assay. (**c**) Correlation analysis between similarity to **IMP-2380** (Tanimoto coefficient) and SOS response pEC_50_ (Spearman correlation r = 0.68, p <0.0001). Inactive compounds set as pIC_50_ = 5, the maximum concentration tested in the assay.

**Extended Data Figure 8.**
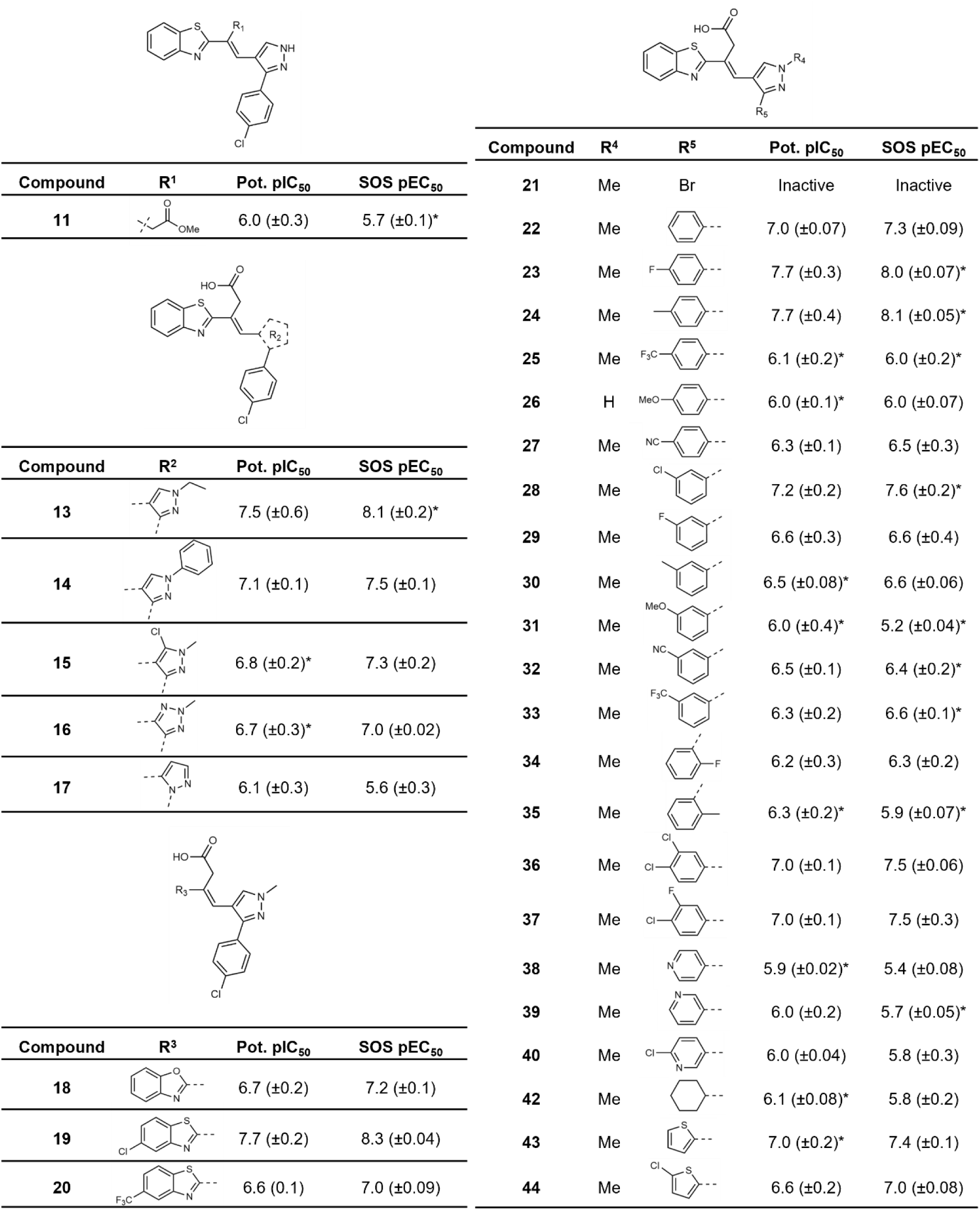
Second analogue library generation. pIC50s (± SD) for potentiation of growth inhibition by ciprofloxacin (Pot. pIC_50_) and pEC_50_s (± SD) for inhibition of SOS response (SOS pEC_50_) in the P*recA-gfp* reporter system in *S. aureus* USA300 JE2 are provided. * represents (n = 2), all other measurements represent (n = 3).

**Extended Data Figure 9.**
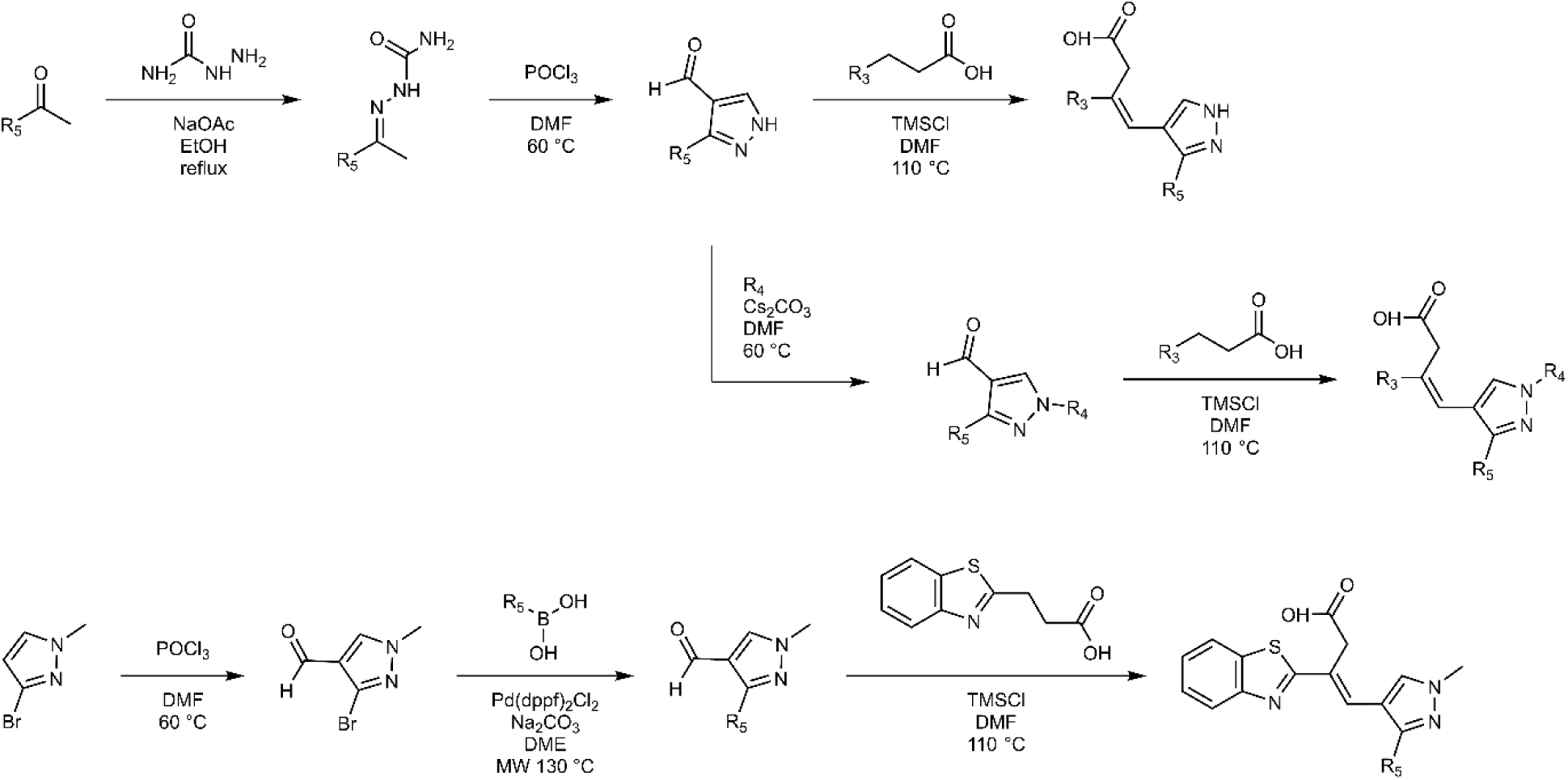
General synthetic routes (**a**) and (**b**) to analogues of **1** and **IMP-2380**.

**Extended Data Figure 10.**
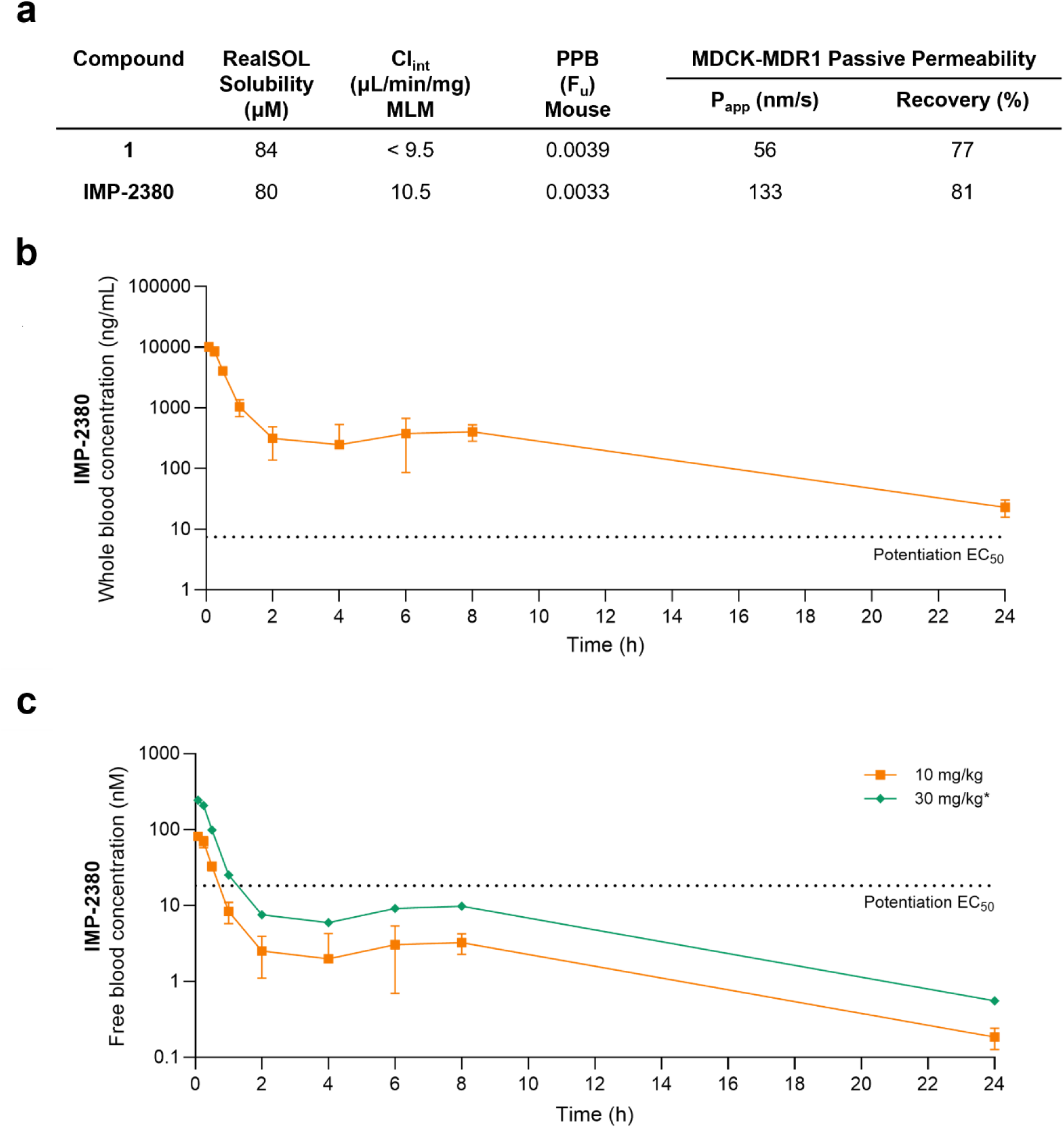
*In vitro* and *in vivo* pharmacokinetic studies of **1** and **IMP-2380**. (**a**) *In vitro* pharmacokinetic parameters for hit compounds **1** and **IMP-2380**. Cl_int_ = intrinsic clearance. MLM = mouse liver microsomes. PPB = Plasma protein binding. (**b**) *In vivo* pharmacokinetic study. Female C57BL/6J mice (n = 3) were given a single intraperitoneal (i.p.) dose of 10 mg/kg **IMP-2380** (5% (v/v) dimethyl sulfoxide (DMSO), 40% polyethylene glycol 400 and 55% sterile deionized water). The whole blood concentration of **IMP-2380** is shown as a function of time. Data are shown as mean ± SD. (**c**) Calculated *in vivo* free blood concentration (C_free_) of **IMP-2380**. The single 10 mg/kg intraperitoneal dose are shown as mean ± SD. C_free_ values for 10 mpk dose were calculated using the equation C_free_ = C_whole_ x F_u_, where F_u_ is the fraction unbound in plasma determined using equilibrium dialysis. *The data for single 30 mg/kg intraperitoneal dose was extrapolated from the mean of the 10 mg/kg data and the free concentration was assumed to correlate linearly with increasing dose.

