## Supplementary file for "Small molecule inhibitors of the NorA multidrug efflux pump potentiate antibiotic activity by binding the outward-open conformation"

### Supplemental figures

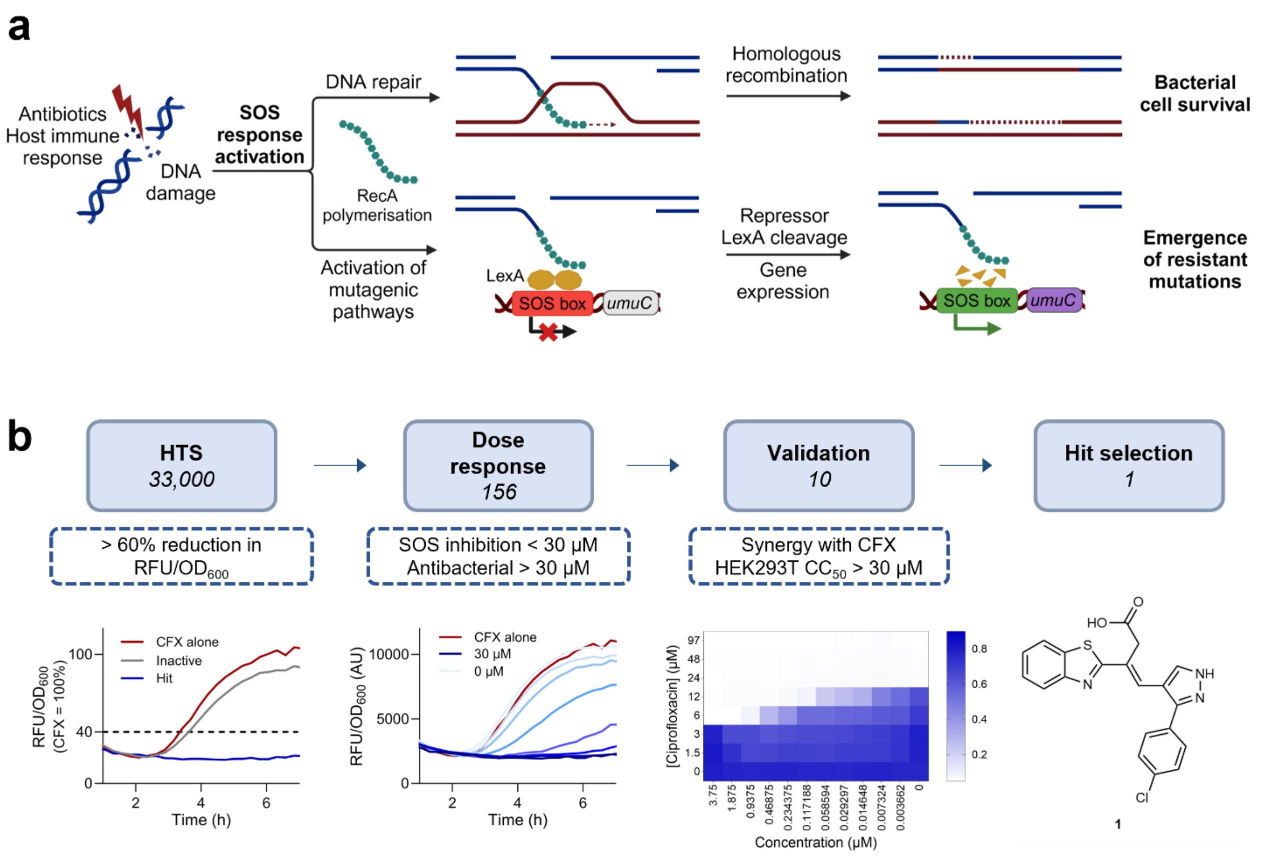


**Figure S1**. High-throughput screen to discover inhibitors of SOS response in *S. aureus.* (**a**) Schematic of SOS activation following DNA damage caused by antibiotics or ROS from the host immune response. RecA forms a nucleofilament and binds in the region of DNA damage. Double-strand breaks are repaired by homologous recombination, allowing for bacterial cell survival. Cleavage of the transcriptional repressor LexA causes expression of the SOS regulon, including genes such as *umuC*, which encodes an error-prone polymerase associated with spontaneous emergence of resistance-bearing mutations. (**b**) Workflow for the high-throughput screen.

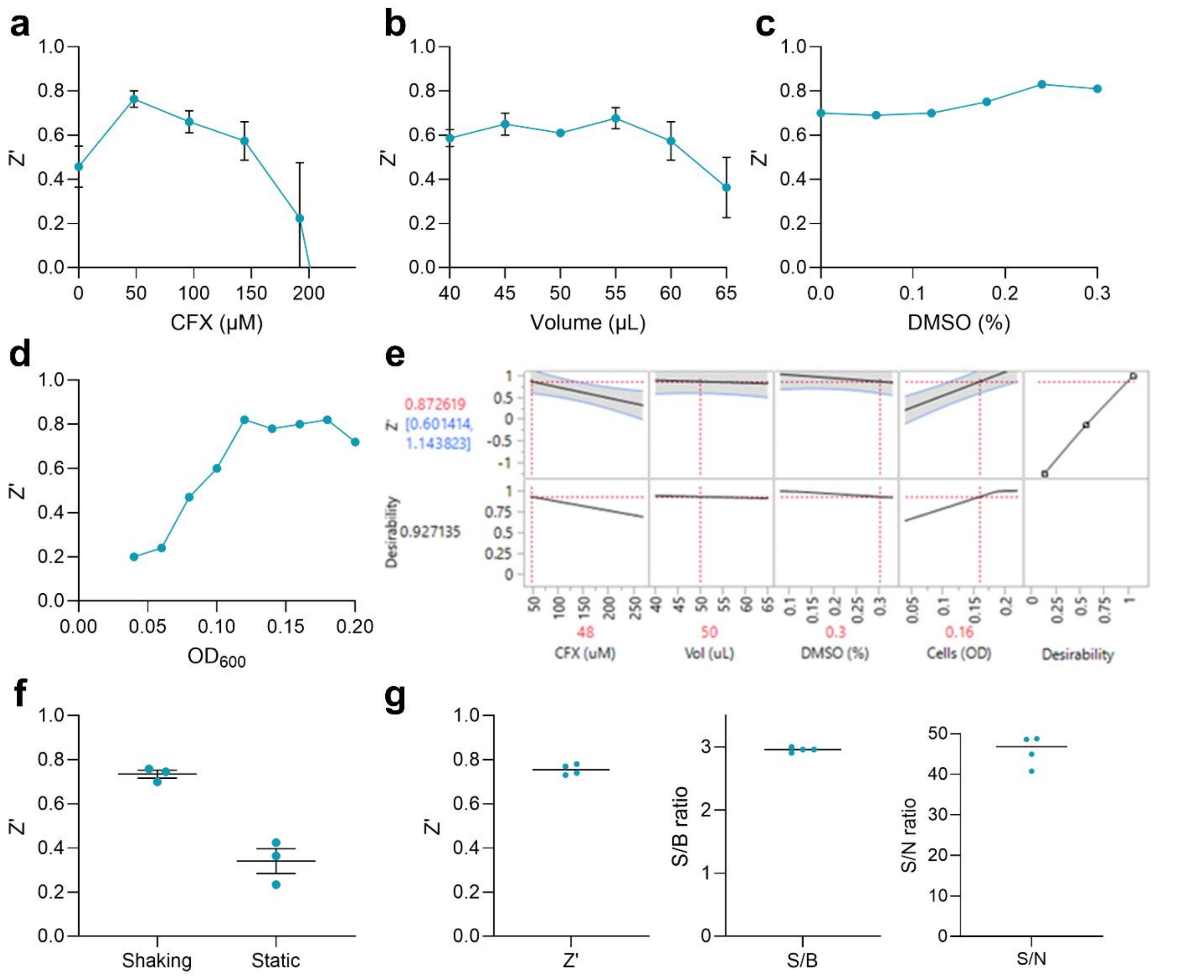


**Figure S2.** HTS development and validation. For HTS optimization, overnight cultures of *S. aureus* JE2 WT *PrecA-gfp* (negative control for SOS inhibition) and *S. aureus* JE2 *rexB*::Tn *PrecA-gfp* (positive control for SOS inhibition)[1] were diluted to the required starting $OD_{600}$ in TSB + kanamycin (90 µg/mL), with the required concentrations of CFX and DMSO, at a defined final volume in a 384-well plate and incubated at 37 °C, 200 rpm overnight before recording $OD_{600}$ and GFP fluorescence. (**a**) 'One-factor at a time' (OFAT) optimization of CFX concentration, showing decreased Z' with increased CFX >48 µM. 1:10 dilution of overnight culture starting density, 50 µL final assay volume, 0% DMSO. Z' was calculated using 16 replicates per condition, data represent mean and SEM (n = 3). (**b**) OFAT optimization of assay volume, showing minimal effect on Z', with a potential local Z' maxima at 45-55 µL and decreased Z' >55 µL final assay volume. 1:10 dilution of overnight culture starting density, 48 µM CFX, 0% DMSO. Z' was calculated using 16 replicates per condition, data represent mean and SEM (n = 3). (**c**) OFAT optimization of DMSO concentration, showing DMSO concentration 0-0.3% (v/v) had minimal effect on Z'. 1:10 dilution of overnight culture starting density, 48 µM CFX, 50 µL final assay volume. Z' was calculated using 16 replicates per condition (n = 1). (**d**) OFAT optimization of seeding density, showing increased Z' with increased $OD_{600}$, reaching a potential local Z' maxima at 0.12-0.18 $OD_{600}$. 48 µM CFX, 50 µL final assay volume, 0.3% DMSO. Z' was calculated using 16 replicates per condition (n = 1). (**e**) Design of experiments (DoE) multifactorial analysis. CFX concentration, DMSO concentration, seeding density and assay volume were varied simultaneously in one experiment following a custom D-optimal design of 24 conditions (see Materials & Methods), and the effect of each input factor deconvoluted from the Z' response using JMP Student Edition software. Representative screenshot of the interactive Prediction Profiler for data exploration, showing how the Z' response changed with variation of each input factor level, at the selected combination of levels. Maximum Desirability (1) was set to Z' = 1. Image shows assay response with input factor levels set to final HTS conditions. Consistent with OFAT optimization: Z' decreased with CFX concentration >48 μM; Z' varied minimally with assay volume 40-65 μL; Z' varied minimally with DMSO concentration 0-0.3% (v/v); Z' increased with increased starting $OD_{600}$ up to a local maxima in desirability at $OD_{600}$ = 0.18. Prediction Profiler shaded areas represent 95% confidence intervals for the simulated means on the curve, Z' was calculated using 8 replicates per condition (n = 1). Final HTS conditions were selected to maximize test compound concentration (0.3% DMSO = 30 μM test compound), and maximize Z' (CFX = 48 μM, volume = 50 μL) whilst using lower cell seeding density ($OD_{600}$ = 0.16) to increase the likelihood of effects from test compounds being observed. (**f**) Optimization of growth conditions for stacked plates to increase throughput. Following OFAT and DoE optimization, overnight cultures of negative and positive control strains were diluted 1:10 in TSB + kanamycin (90 μg/mL) + CFX (48 μM) + DMSO (0.3%), and 50 μL inoculated into a 384-well plate and plates stacks incubated at 37 °C with or without shaking (200 rpm) overnight. Z' was calculated using 196 replicates per condition (n = 3). (**g**) Optimized HTS assay protocol metrics. Overnight cultures of negative and positive control strains were diluted $OD_{600}$ = 0.16 in TSB + kanamycin (90 μg/mL) + CFX (48 μM) + DMSO (0.3%), 50 μL inoculated into a 384-well plate and plate stacks incubated at 37 °C, 200 rpm for 17 h. Z', S/B, and S/N were calculated using 196 replicates per condition (n = 4).

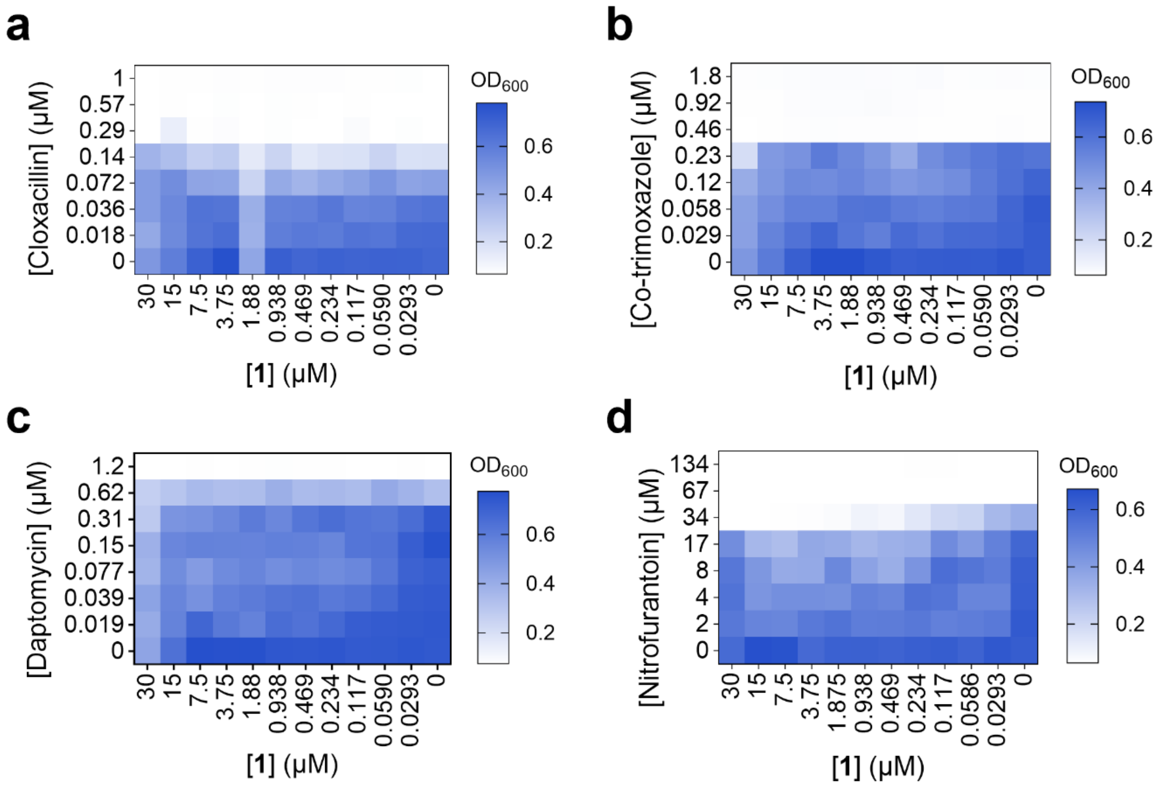


**Figure S3.** Checkerboard MIC assays in *S. aureus* USA300 JE2 with cloxacillin (n = 2) (**a**), co-trimoxazole (n = 2) (**b**), daptomycin (n = 2) (**c**) or nitrofurantoin (n = 2) (**d**) and compound **1**. These figures were used to calculate MIC values in Extended Data Fig. 2e.

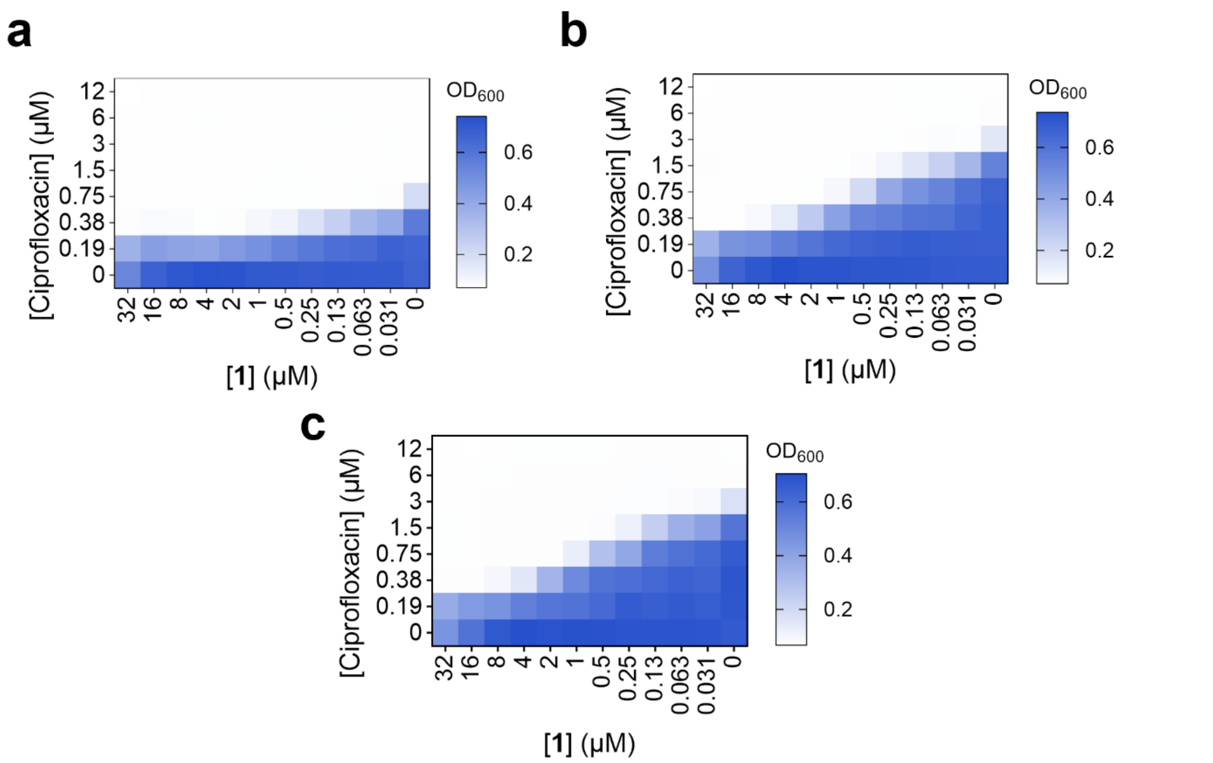


**Figure S4.** Checkerboard MIC assays in *S. aureus* TM51 WT (n = 2) (**a**) 0.25_1 (OE_A) (n = 2) (**b**) or 0.25_1 (OE_B) (n = 2) (**c**) with ciprofloxacin and compound **1**. These figures were used to calculate MIC values used in Fig. 2b and Extended Data Fig. 4a.

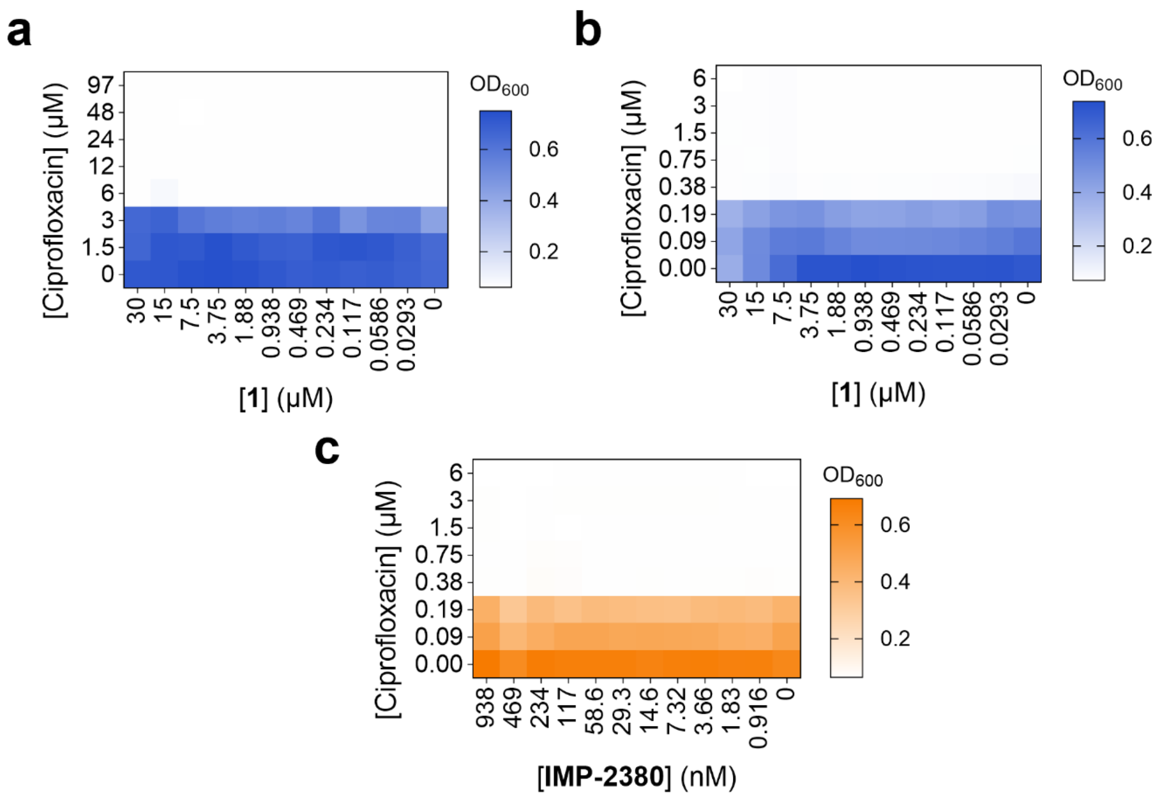

**Figure S5.** Checkerboard MIC assay with ciprofloxacin and compound **1** in *S. aureus* USA300 JE2 *norA*::Tn (**a**) and in SH1000 *norA*::Tn (**b**) or with ciprofloxacin and **IMP-2380** in SH1000 *norA*::Tn (**c**). These figures were used to calculate MIC values used in Fig. 2c and Extended Data Fig. 4b-c.

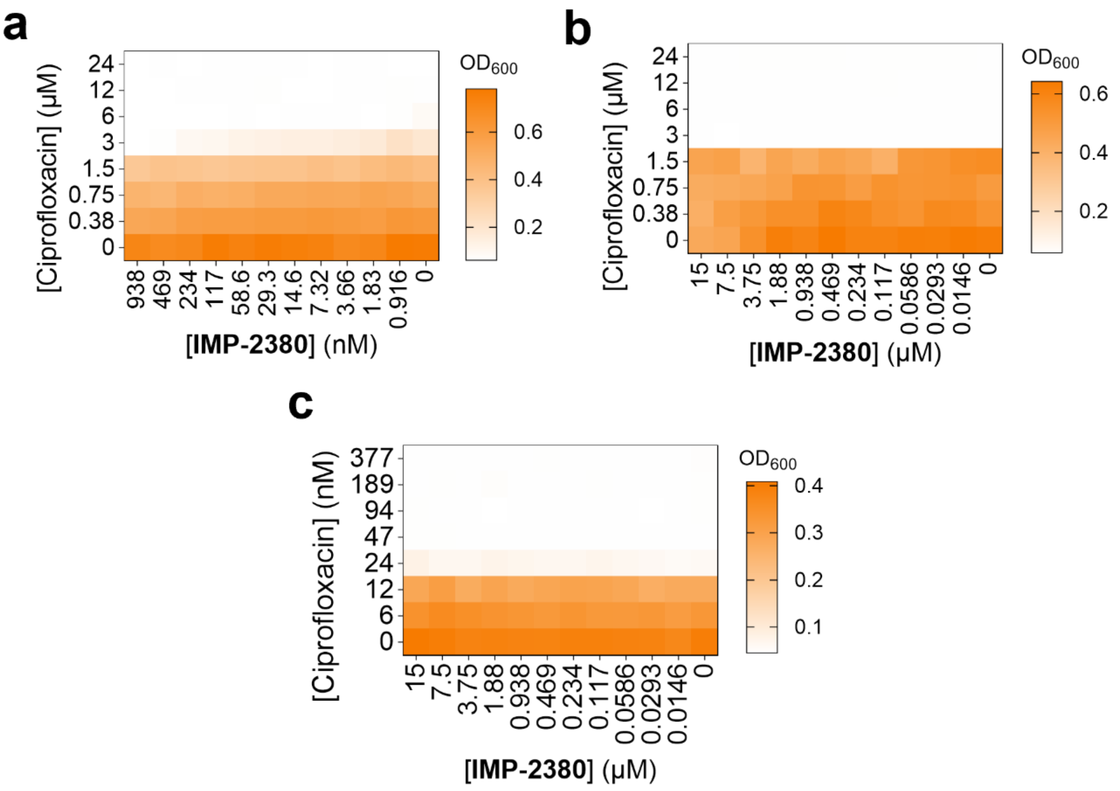


**Figure S6.** Checkerboard MIC assays in *E. faecalis* (**a**), *S. agalactiae* (**b**) and *E. coli* (**c**) with ciprofloxacin and **IMP-2380**. These figures were used to calculate MIC values in Extended Data Fig. 4e.

**Supplementary Tables**

**Table S1.** Ciprofloxacin MIC of MRSA and methicillin sensitive *S. aureus* (MSSA) clinical isolates in combination with **1** (2 μM, n = 1) or **IMP-2380** (1 μM, n = 2).

| | | Ciprofloxacin MIC (μM) | | | | |
|---|---|---|---|---|---|---|
| **Strain** | **Characteristics** | **DMSO (n = 3) Median** | **DMSO (n = 3) Range** | **+ 1 (n = 1)** | **+ IMP-2380 (n = 2) Median** | **+ IMP-2380 (n = 2) Range** |
| 1115 | MRSA | 386 | 386 | 97 | 97 | 97 |
| 1098 | | 386 | 386 | 48 | 97 | 97 |
| 1642 | MRSA | 386 | 386 | 97 | 97 | 97 |
| 1643 | | 386 | 386 | 97 | 97 | 97 |
| 244 | MRSA | 193 | 193 | 97 | 97 | 97 |
| 255 | | 193 | 193 | 97 | 97 | 97 |
| 1810 | MRSA | 193 | 193 | 48 | 97 | 97 |
| 1811 | | 193 | 193 | 48 | 97 | 97 |
| 260 | MRSA | 193 | 193 | 48 | 72.5 | 48-97 |
| 261 | | 193 | 193 | 48 | 72.5 | 48-97 |
| 1659 | MRSA | 97 | 48-97 | 24 | 48 | 48 |
| 1658 | | 48 | 48-97 | 24 | 24 | 24 |
| 655 | MRSA | 48 | 48 | 12 | 18 | 12-24 |
| 654 | | 48 | 48 | 12 | 18 | 12-24 |
| 1980 | MRSA | 48 | 48 | 12 | 12 | 12 |
| 1981 | | 48 | 48 | 12 | 12 | 12 |
| 2004 | MSSA | 3 | 3 | 0.75 | 0.75 | 0.75 |
| 1963 | | 3 | 3 | 0.75 | 0.75 | 0.75 |
| 1745 | MSSA | 1.5 | 1.5 | 0.38 | 0.38 | 0.38 |
| 1688 | | 1.5 | 1.5 | 0.38 | 0.38 | 0.38 |
| 611 | MSSA | 1.5 | 1.5 | 0.38 | 0.38 | 0.38 |
| 408 | | 1.5 | 1.5 | 0.38 | 0.38 | 0.38 |
| 287 | MSSA | 1.5 | 1.5 | 0.38 | 0.285 | 0.19-0.38 |
| 288 | | 1.5 | 1.5 | 0.38 | 0.285 | 0.19-0.38 |

**Table S2. Cryo-EM data collection/processing and structure refinement statistics.**

| | NorA bound to IMP-2380 |
|---|---|
| **Conformation** | Outward-open |
| **PDB ID** | 9PV0 |
| **EMDB ID** | EMD-71880 |
| **Data collection and processing** | |
| Magnification (x) | 105,000 |
| Voltage (kV) | 300 |
| Electron dose ($e^{-}/Å^2$) | 47.12 |
| Defocus range (μm) | -0.4 to -2.5 |
| Collection mode | Super-resolution |
| Pixel size (Å) | 0.4125 |
| Symmetry imposed | C1 |
| Initial number of particles | 8,259,459 |
| Final number of particles | 634,207 |
| Map sharpening B factor ($Å^2$) | 112.1 |
| Map resolution (Å)* | 2.52 |
| **Refinement** | |
| Non-hydrogen atoms | 2,783 |
| Protein residues | 361 |
| Mean B factor | |
| Protein ($Å^2$) | 89.34 |
| **IMP-2380** | 56.05 |
| RMS deviations | |
| Bond lengths (Å) | 0.010 |
| Bond angles (°) | 1.382 |
| MolProbity score | 1.30 |
| Clash score | 0.00 |
| Rotamer outliers (%) | 5.00 |
| Ramachandran plot | |
| Favored (%) | 96.92 |
| Allowed (%) | 3.03 |
| Outliers (%) | 0.00 |
| Model resolution (Å)+ | 2.94 |

*Resolution determined by a FSC value of 0.143.

+Resolution determined between the model and the sharpened map by the FSC value of 0.5.

**Supplemental Methods**

**Table S3. Bacterial Strains**

| Bacterial Strain | Details |
|---|---|
| *S. aureus* USA300 JE2 | LAC derivative strain of the USA300 CA-MRSA lineage, cured of plasmids, resistant to CFX |
| *S. aureus* USA300 JE2 P*recA-gfp*[2] | JE2 strain transformed with pCN34 plasmid with *gfp* gene under the control of *recA* promoter $Kan^R$ |
| *S. aureus* USA300 JE2 *rexB::Tn* P*recA-gfp* | JE2 strain with a *bursa aurealis* transposon insertion in *rexB* and transformed with pCN34 plasmid with *gfp* gene under the control of *recA* promoter $Erm^R$, $Kan^R$ |
| JE2 *norA*::Tn[3] | JE2 strain with a *bursa aurealis* transposon insertion in *norA*. $Erm^R$ |
| JE2 *norB*::Tn[3] | JE2 strain with a *bursa aurealis* transposon insertion in *norB*. $Erm^R$ |
| JE2 *norC*::Tn[3] | JE2 strain with a *bursa aurealis* transposon insertion in *norC*. $Erm^R$ |
| JE2 *mepA*::Tn[3] | JE2 strain with a *bursa aurealis* transposon insertion in *mepA*. $Erm^R$ |
| JE2 *mdeA*::Tn[3] | JE2 strain with a *bursa aurealis* transposon insertion in *mdeA*. $Erm^R$ |
| JE2 *sepA*::Tn[3] | JE2 strain with a *bursa aurealis* transposon insertion in *sepA*. $Erm^R$ |
| JE2 *sdrM*::Tn[3] | JE2 strain with a *bursa aurealis* transposon insertion in *srdM*. $Erm^R$ |
| JE2 *lmrS*::Tn[3] | JE2 strain with a *bursa aurealis* transposon insertion in *lmrS*. $Erm^R$ |
| MRSA and MSSA Clinical isolates[4] | Recovered from patient samples |
| *S. aureus* SH1000[5] | |
| SH1000 *norA*::Tn | SH1000 with a *bursa aurealis* transposon insertion in *norA*. $Erm^R$ |
| SH1000 *umuC*::Tn | SH1000 with a *bursa aurealis* transposon insertion in *umuC*. $Erm^R$ |
| *S. aureus* RN451[6] | RN450 lysogenized with prophage ϕ11 |
| *S. aureus* RN4220[7] | A ciprofloxacin susceptible strain of *S. aureus* lacking any prophages |
| *S. aureus* TM51 | RN4220 ΔattBφ11::Orf5 (pTM378) KanR ; Tnp+ recipient strain |
| *S. aureus* TM51 0.25_1 (OE_A) | *norA*-overexpressing strain selected for with 0.25 µM compound |
| *S. aureus* TM51 0.25_2 (OE_B) | *norA*-overexpressing strain selected for with 0.25 µM compound |
| *Enterococcus faecalis* JH2-2[8] | $Fus^R$ $Rif^R$ strain derived from non-haemolytic clinical isolate JH2 |
| *Streptococcus agalactiae* COH1[9] | Serotype III clinical isolate derived from human bloodstream infection |
| *E. coli* DH5α | Competent cloning strain of *E. coli* with T1 phage resistance and *endA* deficiency |
| *E. coli* C43 (DE3) | Competent strain of *E. coli* derived from BL21 (DE3) with mutations conferring tolerance to the expression of toxic proteins |
| *E. coli* 55244 | Competent strain of *E. coli* for recombinant protein expression |
| *E. coli* BL21 (DE3) | Competent strain of *E. coli* for recombinant protein expression |

**HTS 'One Factor at A Time' (OFAT) optimization:** Overnight cultures of *S. aureus* JE2 WT P*recA-gfp* and *S. aureus* JE2 *rexB*::Tn P*recA-gfp* were diluted to the required starting $OD_{600}$ in TSB + kanamycin (90 µg/mL), with the required concentrations of CFX and DMSO, at a defined final volume, with variation of individual factors as described in Figure S2.

**HTS 'Design of Experiments' (DoE) optimization:** Overnight cultures of *S. aureus* JE2 WT pCN34 *PrecA-gfp* and *S. aureus* JE2 *rexB*::Tn pCN34 *PrecA-gfp* were diluted to the required starting $OD_{600}$ in TSB + kanamycin (90 µg/mL), with the required concentrations of CFX and DMSO, at a defined final volume. Input factor levels were generated using a Custom D Optimal Design in JMP Student Edition (JMP Statistical Discovery LLC) within ranges defined by those explored in OFAT optimization (Figure S2), with the following values:

| Column | Input factor level | | | | Response |
|---|---|---|---|---|---|
| | DMSO (%) | CFX (μM) | $OD_{600}$ | Vol (μL) | Z' |
| 1 | 0.3 | 60 | 0.15 | 50 | 0.87 |
| 2 | 0.13 | 227 | 0.06 | 40 | 0.04 |
| 3 | 0.08 | 46 | 0.08 | 65 | 0.57 |
| 4 | 0.25 | 227 | 0.19 | 40 | 0.64 |
| 5 | 0.11 | 268 | 0.22 | 45 | 0.72 |
| 6 | 0.33 | 50 | 0.04 | 60 | 0.09 |
| 7 | 0.23 | 232 | 0.04 | 65 | -1.17 |
| 8 | 0.25 | 227 | 0.19 | 40 | 0.69 |
| 9 | 0.27 | 220 | 0.09 | 55 | 0.40 |
| 10 | 0.15 | 46 | 0.15 | 65 | 0.88 |
| 11 | 0.17 | 201 | 0.08 | 60 | 0.50 |
| 12 | 0.25 | 151 | 0.21 | 60 | 0.80 |
| 13 | 0.08 | 139 | 0.04 | 65 | 0.44 |
| 14 | 0.13 | 151 | 0.06 | 40 | 0.22 |
| 15 | 0.22 | 67 | 0.06 | 45 | 0.53 |
| 16 | 0.08 | 252 | 0.17 | 60 | 0.88 |
| 17 | 0.25 | 76 | 0.06 | 40 | 0.61 |
| 18 | 0.33 | 134 | 0.17 | 45 | 0.74 |
| 19 | 0.31 | 186 | 0.19 | 65 | 0.75 |
| 20 | 0.13 | 76 | 0.19 | 40 | 0.85 |
| 21 | 0.18 | 55 | 0.18 | 55 | 0.84 |
| 22 | 0.11 | 134 | 0.06 | 45 | -0.21 |
| 23 | 0.1 | 181 | 0.05 | 50 | 0.24 |
| 24 | 0.3 | 60 | 0.15 | 50 | 0.82 |

The Z' response calculated from $OD_{600}$ and GFP fluorescence measurements for positive and negative controls was analyzed in JMP Student Edition using a Standard Least Squares model with emphasis on Effect Screening. Data was explored using the interactive Prediction Profiler functionality in JMP for comparison to OFAT optimization

**Mammalian cell toxicity**

HEK293T cells were cultured in Dulbecco's modified Eagle medium (DMEM) supplemented with GlutaMAX (Gibco, 61965026) and 10% v/v Fetal Bovine Serum (FBS, Gibco, A5256701) in a 37 °C, 5% $CO_2$ incubator.

HEK293T cells were seeded in a sterile 96-well plate (Grenier, 655180) at a density of 2000 cells/well and incubated at 37 °C overnight. The next day, the cells were treated with SYTOX™ Green Nucleic Acid Stain (Invitrogen, S7020, 250 nM) and compounds in a dose-dependent manner. For controls, cells were treated with 0.1% DMSO or 4.2 μM puromycin (Sigma, P8833-25MG). The plate was placed into an IncuCyte® S3 Live Cell Analysis System (Sartorius) and readings for phase and green fluorescence were taken every 4 hours for 120 h, with 4 images per well. The images were analyzed with IncuCyte® 2021C software.

For cell proliferation, the confluence area (%) was exported, the background at t = 0 h subtracted and the slopes of the linear portion (56-80 h) calculated by plotting against time. The percent viability was determined by normalizing to the positive (DMSO) and negative (puromycin) control and graphed against [compound]. $EC_{50}$ values were determined using Prism 10 for each replicate.

For cytotoxicity, the ratio of green fluorescence area to phase-contrast area (confluence) was exported. The background at t = 0 h was subtracted and the total area under the curve (AUC) was calculated. The percent cytotoxicity was determined by normalizing to the positive (puromycin) and negative (DMSO) control and graphed against [compound]. $CC_{50}$ values were determined using Prism 10 for each replicate.

**Docking of compounds into the Cryo-EM Structure of NorA-IMP-2380**

Flexible non-covalent docking of the compounds was performed using ICM-Pro software (version 3.9-5/Windows, Molsoft LLC). The chemical structures of docked compounds were added to a combined .sdf file for docking in batch. The cryo-EM structure of NorA with **IMP-2380** bound (PDB ID: 9VP0) was converted to an ICM object for the docking process. In preparation, the energetically favorable degree of protonation was performed on side chains residues of His, Pro, Asn, Gln, and Cys prior to docking. The location of **IMP-2380** was used to identify the binding pocket for batch docking. Three independent docking runs were performed for the set of ligands, varying the thoroughness between 1-5, with the top 10 docking poses compared. **IMP-2380** was re-docked into the same binding site as a control. Ligand positions were ranked according to their ICM scores (unitless), which ranks position based on ligand-receptor energy parameters and interactions. Final ligand poses were then presented for final figures and indication of interactions in Pymol Version 2.5.4.

**Intrinsic clearance (Cli) experiments**

Compounds **1** or **IMP-2380** (0.5 µM) were incubated with female CD1 mouse liver microsomes (Xenotech; 0.5 mg/mL 50 mM potassium phosphate buffer, pH 7.4) and the reaction started with addition of excess NADPH (8 mg/mL 50 mM potassium phosphate buffer, pH 7.4). Immediately, at t = 0, then at 3, 6, 9, 15 and 30 mins a 50 µL aliquot of the incubation mixture was removed and mixed with Acetonitrile (100 µL) to stop the reaction. Internal standard was added to all samples, the samples centrifuged to sediment precipitated protein and the plates then sealed prior to UPLC-MSMS analysis (Xevo TQ-S Micro, Waters ™).

XLfit (IDBS, UK) was used to calculate the exponential decay and consequently the rate constant (k) from the ratio of peak area of test compound to internal standard at each timepoint. The rate of intrinsic clearance (CLi) of each test compound was then calculated using the following calculation:

$$CLi(mL/min/mg\ protein) = k \times V$$

Where V (mL/mg protein) = incubation volume (0.5 mL) / mg protein added (0.25 mg protein). Verapamil (0.5 µM) was used as a positive control to confirm acceptable assay performance

**"RealSOL" method for solubility**

5 µL of 10 mM **1** or **IMP-2380** DMSO stock solutions were added to 195 µL of phosphate buffered saline, pH 7.4 (Sigma-Aldrich, #P4417) in duplicate. This solution was then mixed for 24 h (900 rpm, 25 °C) excluding light. After mixing, the solubility test samples were filtered under vacuum to remove any undissolved material (Millipore Multiscreen HTS filter, 96-well format, #MSHVN4B10). The filtrate was analyzed for dissolved drug compound using a truncated UHPLC methodology. A Shimadzu Nexera X2 UHPLC system was used, with a reversed-phase column and a formic acid gradient elution. The Shimadzu analytical column was a Hypersil Gold C18 2.1x50mm with a flow rate of 0.6 mL/min. The Shimadzu system solvent gradients started at 2% MeCN in water ending at 98% MeCN in water after 2 minutes with a 0.05% formic acid additive.

A calibration solution was prepared in the following way: The same 10 mM solution used to prepare the solubility test sample was diluted in DMSO to give a 500 µM solution. This solution was then again diluted with

50:50 MeCN:$H_2O$ to give a 50 µM solution (one replicate). Aliquots (0.2, 2.0 and 5.0 µL) of this 50 µM solution were then injected onto the UHPLC system and the areas of the resultant peaks integrated to produce a calibration line. Aliquots of the test sample filtrate (0.4 and 5.0 µL) were then injected onto the UHPLC system and the resultant peak areas for any peaks corresponding to the test compound determined and quantified using the calibration line.

**Plasma protein binding experiments**

In brief, a 96-well equilibrium dialysis apparatus was used to determine the free fraction in plasma for each compound (HT Dialysis LLC, Gales Ferry). Membranes (12-14 kDA cut-off) were conditioned in deionized water for 60 mins, followed by conditioning in 80:20 deionized $H_2O$:EtOH for 20 mins, and then rinsed in isotonic buffer before use. Freshly thawed female CD1 mouse plasma (BioIVT) was then centrifuged (Allegra X12-R, Beckman Coulter), spiked with test compound (final concentration 10 µg/mL), and 150 µL aliquots (n = 6) loaded into the 96-well equilibrium dialysis plate. Dialysis vs isotonic buffer (150 µL) was carried out for 5 h in a temperature-controlled incubator at 37 ºC (Barworld scientific Ltd) using an orbital microplate shaker at 100 revolutions/minute (Barworld scientific Ltd). After incubation, 50 µL aliquots of plasma or buffer were transferred to micronic tubes (Micronic B.V.) and the composition in each tube balanced with control fluid (50 µL), such that the volume of buffer to plasma is the same. Sample extraction was performed by the addition of 200 µL of MeCN containing an appropriate internal standard. Samples were allowed to mix for 1 min and then centrifuged at 3000 rpm for 15 mins (Allegra X12-R, Beckman Coulter) after which 150 µL of supernatant was removed to 50 µL of water. All samples were analyzed by UPLC-MS/MS. The unbound fraction was determined as the ratio of the peak area in buffer to that in plasma.

**MDCK Passive Permeability**

MDCK-MDR1 cells (Netherlands Cancer Institute) were maintained in culture (DMEM, Gibco 61965-026 supplemented with 1% penicillin/streptomycin, 10% FCS) until required.

MDCK-MDR1 cells were seeded onto individual transwell 'Thincerts' (Greiner, 662610) at a density of 35,000 cells/well. Cells were grown at 37 °C, 5% $CO_2$ for 3 days. On day 4, media was replaced with fresh media and incubated for 1 h. Media was removed and replaced with Dulbecco's PBS (Gibco, 14287-080) and cell inserts incubated for a further 1 h. Dosing solutions containing 3 µM compound **1** or **IMP-2380** and 10 µM Lucifer Yellow (1% DMSO) were prepared. 1.2 mL of PBS (1% DMSO) was added to wells of a 24-well cell culture plate (Corning, 353504). 0.35 mL of dosing solution was added in duplicate to the apical side of the transwell and transwells transferred into the receiver plate solutions. Transwell plates were then incubated for 1 h after which inserts were removed to an empty plate to prevent any further permeation of compound. 100 mL of solution from donor, receiver wells were removed to a 96 well plate alongside 100 mL of dosing solution. 150 µL of MeCN containing internal standard (eg 100 ng/mL Sulfadimethoxine) was then added to all samples prior to analysis by LC-MS/MS. Bupropion (positive controls) and Atenolol (negative control) were run alongside test compounds. To confirm monolayer integrity, a further 100 µL from each compartment was added of the 96 well F-bottomed microtiter plate containing the Lucifer Yellow standard curve for fluorescence determination of Lucifer Yellow concentrations. Papp (apparent permeability) values were calculated using the following equation:

$$\text{Papp (nm/sec)} = \frac{(\text{Volume receiver/ A}) * (\text{Response receiver/ Response donor})}{\text{Incubation time (sec)}}$$

### *In vivo* pharmacokinetics

All regulated procedures, at the University of Dundee, on living animals were carried out under the authority of a project license approved by the local Animal Welfare and Ethical Review Board (reference WEC2019_08), and issued by the Home Office under the Animals (Scientific Procedures) Act 1986, as amended in 2012 (and in compliance with EU Directive EU/2010/63).

Female C57BL/6J mice of approximately 6 weeks of age were obtained from Charles River Laboratories, UK. Animals were maintained under a 12-hour light / 12-hour dark cycle. Temperature and relative humidity were maintained between 20-24°C and 45-65% respectively. Food and water were supplied ad-libitum throughout. All animals received a minimum of 10 days acclimatization and were randomized prior to the start of study. Animals were not blinded as all animals in the pharmacokinetic study received the same dose, and results are based on objective blood concentration measurements.

The dose formulation was prepared on the day of dosing. **IMP-2380** was dosed intraperitoneally to n = 3 mice as a solution at 1 mg free base / kg (dose volume 10 mL/kg; dose vehicle, 5% (v/v) dimethyl sulfoxide (DMSO), 40% polyethylene glycol 400 and 55% sterile deionized water). Blood samples (10 µL) were taken from each mouse prior to dose, 0.05, 0.25, 0.5, 1, 2, 4, 6, 8, and 24 h post dose and mixed with distilled water (90 µL). All samples were stored at -20 °C until analysis. After suitable sample preparation, the concentration of test compound in blood was determined by UPLC-MS/MS. Pharmacokinetic parameters were derived from the blood concentration time curve using Phoenix WinNonLin software (Certara, USA).

### General Chemical Information

All chemicals, reagents and solvents were purchased from Sigma-Aldrich, Fluorochem or Enamine and used without further purification. Reactions monitored by TLC used UV light (254 nm) in visualization of analytical TLC plates. Microwave reactions were conducted in a Biotage Initiator+. Reactions monitored and intermediates confirmed by LCMS used an Agilent 1260 Infinity II MSD/XT Single Quad system. The Agilent analytical column was a *Poroshell* HPH-C18 3.0x50mm with a flow rate of 0.5 mL/min and an injection volume of 2.00 µL. The Agilent system solvent gradients started at 5% MeCN in water ending at 95% MeCN in water after 7 minutes with a 0.1% formic acid additive. Flash silica column chromatography for purification was performed using Biotage Selekt.

Nuclear magnetic resonance (NMR) spectra were in all cases consistent with the proposed structures. $^{1}$H, $^{19}$F, and $^{13}$C NMR spectra were recorded at room temperature (RT) at 600, 500, and 400 ($^{1}$H), 377 ($^{19}$F), 150 and 101 ($^{13}$C) MHz, respectively. Characteristic chemical shifts (δ) are given in parts-per-million downfield from tetramethylsilane (for $^{1}$H-NMR) using conventional abbreviations for designation of major peaks: e.g. s, singlet; d, doublet; t, triplet; q, quartet; m, multiplet; br, broad. The following abbreviations have been used for common solvents: MeOD, deuteromethanol $CDCl_3$, deuterochloroform and DMSO-$d_6$, deuterodimethylsulfoxide. High resolution mass spectrometry (HRMS) was performed using electrospray ionization (ESI) and time-of-flight (TOF) mass analysis.

**Purchased Compounds**

| Compound | Supplier | Purity by HPLC |
|---|---|---|
| **1** | Enamine | >99% |
| **3** | Enamine | >99% |
| **4** | Enamine | >99% |
| **5** | Enamine | >99% |
| **6** | Enamine | >99% |
| **8** | Enamine | >99% |
| **9** | Fluorochem | >99% |
| **10** | Enamine | >99% |
| **12** | Enamine | >99% |

**General Synthetic Procedures**

**Method A: Pyrazole *N*-alkylation**

The appropriate pyrazole-4-carbaldehyde (1 equiv) was dissolved in anhydrous DMF. Cesium carbonate (2 equiv) and the appropriate alkyl halide (1.5 equiv) were added and the reaction stirred at 60 °C for 16 h. The reaction was poured onto water, stirred at r.t. for 1 h and subsequently extracted with EtOAc. The combined organic phases were washed with 10% LiCl, brine, dried using $Na_2SO_4$ and solvent removed *in vacuo*. The crude product was purified using flash column chromatography.

**Method B: Knoevenagel condensation**

The appropriate propionic acid (0.876-1 equiv) and pyrazole-4-carbaldehyde (1 equiv) were dissolved in anhydrous DMF and sealed in a microwave vial. TMSCl (4.7-6 equiv) was added dropwise before heating the reaction to 110-135 °C for 9-24 h. The crude compound was precipitated in water, the solid collected and triturated in MeOH and the solid collected by vacuum filtration. The solid was further washed with cold MeOH and water and dried *in vacuo.*

**Method C: Benzothiazole cyclisation**

The appropriate amino-benzenethiol (1 equiv) was dissolved in toluene. Succinic anhydride (1 equiv) was added, and the reaction stirred at r.t. for 4 h, followed by refluxing for a further 2 h. The reaction solvent was removed *in vacuo*. The crude solid was dissolved in minimal EtOAc and washed three times with sat. aq. $NaHCO_3$. The combined aqueous phases were then neutralized to pH = 7 with 37% HCl. The precipitated product was then isolated through vacuum filtration, washed three times with cold water and dried *in vacuo*.

**Method D: Suzuki coupling**

[1,1′-bis(diphenylphosphine)ferrocene]dichloropalladium(II)·DCM (0.1 equiv) was added to 3-bromo-1-methyl-1*H*-pyrazole-4-carbaldehyde (1 equiv), boronic acid (1-2 equiv) and 2 M sodium carbonate (1.5 equiv) in DME and heated in a microwave at 130 °C for 45 minutes. The reaction mixture was diluted with EtOAc, washed with water, brine, dried using $Na_2SO_4$ and solvent removed *in vacuo*. The crude product was resuspended in DCM, filtered through a pad of Celite, and purified using flash column chromatography.

**(*E*)-3-(benzo[*d*]thiazol-2-yl)-4-(3-(4-chlorophenyl)-1-methyl-1*H*-pyrazol-4-yl)but-3-enoic acid (IMP-2380)**

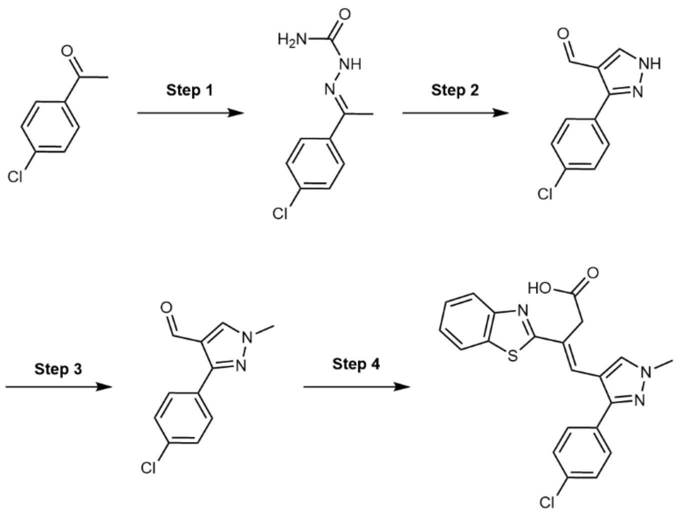

**Step 1**

Semicarbazide hydrochloride (765 mg, 9.68 mmol) and water (172 mL) were added in small portions to a stirred solution of 4'-chloroacetophenone (1.50 g, 9.22 mmol) and sodium acetate (1.11 g, 12.9 mmol) in EtOH (172 mL). The reaction was refluxed for 16 h and then stirred at r.t. for 6 h. EtOH was removed *in vacuo* and the precipitate collected by vacuum filtration. (*E*)-2-(1-(4-chlorophenyl)ethylidene)hydrazine-1-carboxamide (1.76 g, 8.31 mmol, 90%) was obtained as a colorless solid. $^1$H NMR (400 MHz, $CDCl_3$) δ 8.33 (s, 1H), 7.65-7.60 (m, 2H), 7.37-7.33 (m, 2H), 2.22 (s, 3H). LCMS: $[M-H]^+$ = 212.6.

**Step 2**

(*E*)-2-(1-(4-chlorophenyl)ethylidene)hydrazine-1-carboxamide (1.05 g, 4.96 mmol) was dissolved in anhydrous DMF (50 mL), cooled to 0 °C and $POCl_3$ (6 mL) added dropwise. The reaction was allowed to heat to r.t., then heated at 60 °C for 16 h, before cooling to r.t. and stirred for 18 h. The reaction was quenched by pouring into ice-cold water and neutralized with 10% NaOH. The mixture was cooled to 0 °C and extracted twice with DCM. The combined organic phases were washed with 10% LiCl, brine, dried using $Na_2SO_4$ and the solvent removed *in vacuo*. The crude product was purified using flash column chromatography eluting with a gradient of 0-45% EtOAc in n-Hex. 3-(4-chlorophenyl)-1*H*-pyrazole-4-carbaldehyde (645 mg, 3.12 mmol, 63%) was obtained as an off-white solid. $^1$H NMR (400 MHz, DMSO-$d_6$) δ 13.81 (s, 1H), 9.89 (s, 1H), 8.58 (s, 1H), 7.90 (d, *J* = 8.0 Hz, 2H), 7.55 (d, *J* = 8.2 Hz, 2H). LCMS: $[M-H]^+$ = 207.0.

**Step 3**

According to general method A, 3-(4-chlorophenyl)-1*H*-pyrazole-4-carbaldehyde (479 mg, 2.32 mmol) was reacted with cesium carbonate (1.57 g, 4.64 mmol) and iodomethane (228 µL, 3.48 mmol) in anhydrous DMF (20 mL). The crude product was purified using flash column chromatography eluting with a gradient of 15-40% EtOAc in Hex. 3-(4-chlorophenyl)-1-methyl-1*H*-pyrazole-4-carbaldehyde (221 mg, 1.39 mmol, 60%) was obtained as a light yellow solid. $^1$H NMR (400 MHz, $CDCl_3$) δ 9.90 (s, 1H), 7.99 (s, 1H), 7.72-7.68 (m, 2H), 7.44-7.40 (m, 2H), 3.98 (s, 3H). LCMS: $[M-H]^+$ = 221.6.

**Step 4**

According to general method B, 3-(1,3-benzothiazol-2-yl)propanoic acid (216 mg, 1.04 mmol) and 3-(4-chlorophenyl)-1-methyl-1*H*-pyrazole-4-carbaldehyde (247 mg, 1.10 mmol) were dissolved in anhydrous DMF (1.35 mL) and sealed in a microwave vial. TMSCl (709 µL, 5.48 mmol) was added dropwise before heating the reaction to 135 °C for 9 h. (E)-3-(benzo[*d*]thiazol-2-yl)-4-(3-(4-chlorophenyl)-1-methyl-1*H*-pyrazol-4-yl)but-3-enoic acid (70.0 mg, 0.171 mmol, 16%) was obtained as a yellow solid. $^{1}$H NMR (400 MHz, DMSO-$d_6$) δ 12.50 (s, 1H), 8.11 (s, 1H), 8.07-8.00 (m, 1H), 7.98-7.91 (m, 1H), 7.63-7.53 (m, 4H), 7.49 (ddd, *J* = 8.2, 7.2, 1.4 Hz, 1H), 7.41 (ddd, *J* = 8.3, 7.2, 1.3 Hz, 1H), 7.33 (s, 1H), 3.97 (s, 3H), 3.89 (s, 2H). $^{13}$C NMR (101 MHz, DMSO-$d_6$) δ 172.0, 170.3, 153.7, 149.9, 134.3, 133.3, 132.1, 132.0, 130.3, 129.3, 128.1, 127.7, 126.9, 125.9, 123.1, 122.5, 114.1, 35.7.(One $^{13}$C NMR environment hidden behind DMSO peak). HRMS (ESI) $C_{21}H_{17}N_3O_2SCl$ $[M+H]^+$ m/z found 410.0722 cald 410.0730

**(*E*)-4-(3-(4-chlorophenyl)-1-methyl-1*H*-pyrazol-4-yl)-3-(5-ethyl-7-(trifluoromethyl)-[1,2,4]triazolo[1,5-a]pyrimidin-2-yl)but-3-enoic acid (7)**

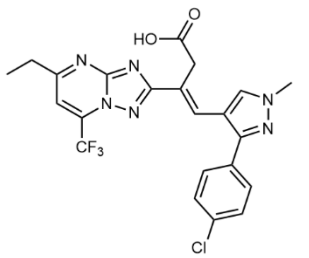


According to general method B, 3-(5-ethyl-7-(trifluoromethyl)-[1,2,4]triazolo[1,5-a]pyrimidin-2-yl)propanoic acid (39.0 mg, 0.136 mmol) and 3-(4-chlorophenyl)-1-methyl-1*H*-pyrazole-4-carbaldehyde (31.6 mg, 0.143 mmol) were dissolved in anhydrous DMF (0.8 mL) and sealed in a microwave vial. TMSCl (86.3 µL, 0.680 mmol) was added dropwise before heating the reaction to 120 °C for 10 h. (*E*)-4-(3-(4-chlorophenyl)-1-methyl-1*H*-pyrazol-4-yl)-3-(5-ethyl-7-(trifluoromethyl)-[1,2,4]triazolo[1,5-a]pyrimidin-2-yl)but-3-enoic acid (46.1 mg, 93.9 µmol, 69%) was obtained as a pale orange solid. $^{1}$H NMR (400 MHz, DMSO-$d_6$) δ 12.25 (s, 1H), 8.26 (s, 1H), 8.05 (s, 1H), 7.67 (s, 1H), 7.62 (d, *J* = 8.5 Hz, 2H), 7.52 (d, *J* = 8.6 Hz, 2H), 3.97 (s, 3H), 3.10 (t, *J* = 7.2 Hz, 2H), 2.79 (t, *J* = 7.2 Hz, 2H), 2.35 (s, 3H). $^{19}$F NMR (377 MHz, DMSO-$d_6$) δ -67.17 (s). $^{13}$C[$^{19}$F] NMR (101 MHz, DMSO-$d_6$) δ 173.8, 169.2, 163.8, 156.1, 145.0, 139.2, 133.2, 133.1, 132.3, 132.1, 130.6, 130.2, 129.2, 129.0, 128.1, 115.4, 106.3, 31.6, 24.4, 15.7. HRMS (ESI) calculated for $C_{22}H_{19}N_6O_2SClF_3$ $[M+H]^+$: 491.1210, found: 491.1216.

**Methyl (*E*)-3-(benzo[*d*]thiazol-2-yl)-4-(3-(4-chlorophenyl)-1*H*-pyrazol-4-yl)but-3-enoate (11)**

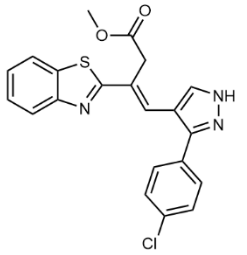

(*E*)-3-(benzo[*d*]thiazol-2-yl)-4-(3-(4-chlorophenyl)-1*H*-pyrazol-4-yl)but-3-enoic acid (53.0 mg, 0.134 mmol) was suspended in MeOH (1.5 mL) and cooled to 0 °C. Thionyl chloride (19.0 µL, 0.254 mmol) was added and the reaction was left to stir at room temperature for 1 h. Additional thionyl chloride (10.0 µL, 0.134 mmol) was added and the reaction was stirred at r.t. for 16 h. The reaction was subsequently quenched with water and extracted three times with EtOAc. The combined organic phases were washed with brine, dried using $MgSO_4$ and solvent removed *in vacuo*. The crude product was purified using flash column chromatography eluting with a gradient of 0-70% EtOAc in n-Hex. Methyl (*E*)-3-(benzo[*d*]thiazol-2-yl)-4-(3-(4-chlorophenyl)-1*H*-pyrazol-4-yl)but-3-enoate (14.3 mg, 34.9 µmol, 24%) was obtained as a yellow solid. $^1$H NMR (400 MHz, $CDCl_3$) δ 7.98 (d, *J* = 8.1 Hz, 1H), 7.93 (s, 1H), 7.84-7.78 (m, 1H), 7.53 (d, *J* = 8.6 Hz, 2H), 7.47-7.42 (m, 3H), 7.39-7.33 (m, 2H), 4.06 (s, 2H), 3.73 (s, 3H). $^{13}$C NMR (101 MHz, $CDCl_3$) δ 171.6, 169.6, 153.9, 135.1, 134.7, 129.8, 129.6, 129.4, 128.7, 127.5, 126.3, 125.5, 123.4, 121.5, 52.5, 35.8. HRMS (ESI) calculated for $C_{21}H_{17}N_3O_2SCl$ $[M+H]^+$: 410.0730, found: 410.0732.

**(*E*)-3-(benzo[*d*]thiazol-2-yl)-4-(3-(4-chlorophenyl)-1-ethyl-1*H*-pyrazol-4-yl)but-3-enoic acid (13)**

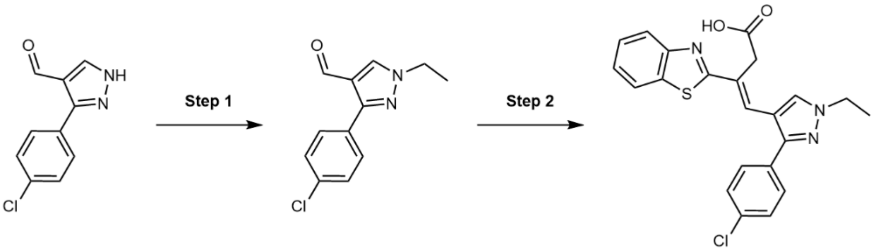


**Step 1**

According to general method A, 3-(4-chlorophenyl)-1*H*-pyrazole-4-carbaldehyde (301 mg, 1.46 mmol) was reacted with cesium carbonate (989 mg, 2.91 mmol) and iodoethane (184 µL, 2.19 mmol) in anhydrous DMF (13 mL). The crude product was purified using flash column chromatography eluting with a gradient of 10-40% EtOAc in n-Hex. 3-(4-chlorophenyl)-1-ethyl-1*H*-pyrazole-4-carbaldehyde (175 mg, 0.746 mmol, 51%) was obtained as a yellow solid. $^1$H NMR (400 MHz, $CDCl_3$) δ 9.90 (s, 1H), 8.02 (s, 1H), 7.75-7.67 (m, 2H), 7.46-7.38 (m, 2H), 4.23 (q, *J* = 7.3 Hz, 2H), 1.55 (t, *J* = 7.3 Hz, 3H). LCMS: $[M-H]^+$ = 235.7.

**Step 2**

According to general method B, 3-(1,3-benzothiazol-2-yl)propanoic acid (148 mg, 0.714 mmol) and 3-(4-chlorophenyl)-1-ethyl-1*H*-pyrazole-4-carbaldehyde (175 mg, 0.731 mmol) were dissolved in anhydrous DMF

(1.10 mL) and sealed in a microwave vial. TMSCl (473 µL, 3.66 mmol) was added dropwise before heating the reaction to 100 °C for 14 h and then 120 °C for 2 h. (*E*)-3-(benzo[*d*]thiazol-2-yl)-4-(3-(4-chlorophenyl)-1-ethyl-1*H*-pyrazol-4-yl)but-3-enoic acid (16.9 mg, 39.9 µmol, 5%) was obtained as a yellow solid. $^{1}$H NMR (400 MHz, DMSO-$d_6$) δ 12.53 (s, 1H), 8.16 (s, 1H), 8.04 (dd, *J* = 8.0, 1.3 Hz, 1H), 7.98-7.91 (m, 1H), 7.63-7.53 (m, 4H), 7.49 (ddd, *J* = 8.2, 7.2, 1.3 Hz, 1H), 7.41 (td, *J* = 7.6, 1.3 Hz, 1H), 7.34 (s, 1H), 4.26 (q, *J* = 7.2 Hz, 2H), 3.90 (s, 2H), 1.45 (t, *J* = 7.2 Hz, 3H). $^{13}$C NMR (101 MHz, DMSO-$d_6$) δ 171.6, 169.9, 153.2, 149.4, 133.8, 132.8, 131.6, 130.3, 129.8, 128.8, 127.6, 127.3, 126.4, 125.5, 122.6, 122.0, 113.4, 46.8, 35.3, 15.3. HRMS (ESI) $C_{22}H_{19}ClN_3O_2S$ $[M+H]^+$ m/z found 424.0886 cald 424.0887.

**(*E*)-3-(benzo[*d*]thiazol-2-yl)-4-(1,3-diphenyl-1*H*-pyrazol-4-yl)but-3-enoic acid (14)**

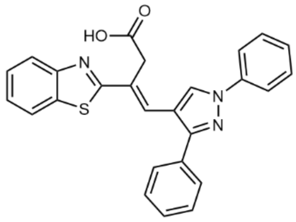


According to general method B, 3-(1,3-benzothiazol-2-yl)propanoic acid (175 mg, 0.844 mmol) and 1,3-diphenyl-1*H*-pyrazole-4-carbaldehyde (222 mg, 0.896 mmol) were dissolved in anhydrous DMF (1 mL) and sealed in a microwave vial. TMSCl (551 µL, 4.25 mmol) was added dropwise before heating the reaction to 110 °C for 16 h. (*E*)-3-(benzo[*d*]thiazol-2-yl)-4-(1,3-diphenyl-1*H*-pyrazol-4-yl)but-3-enoic acid (192 mg, 0.438 mmol, 52%) was obtained as a brown solid. $^{1}$H NMR (400 MHz, DMSO-$d_6$) δ 12.52 (s, 1H), 8.83 (s, 1H), 8.09-8.03 (m, 1H), 7.97 (d, *J* = 8.1 Hz, 3H), 7.75-7.68 (m, 2H), 7.61-7.54 (m, 4H), 7.50 (td, *J* = 7.3, 1.5 Hz, 2H), 7.45-7.36 (m, 3H), 4.06 (s, 2H). $^{13}$C NMR (101 MHz, DMSO-$d_6$) δ 171.7, 169.8, 153.2, 152.5, 139.2, 133.9, 132.1, 129.7, 128.9, 128.7, 128.4, 128.0, 127.1, 126.9, 126.5, 125.6, 122.7, 122.1, 119.0, 115.9, 35.3. HRMS (ESI) $C_{26}H_{20}N_3O_2S$ $[M+H]^+$ m/z found 438.1269 cald 438.1276

**(*E*)-3-(benzo[*d*]thiazol-2-yl)-4-(5-chloro-1-methyl-3-phenyl-1*H*-pyrazol-4-yl)but-3-enoic acid (15)**

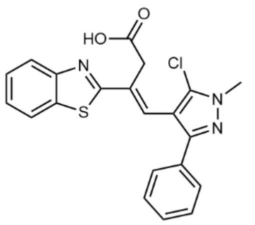


According to general method B, 3-(1,3-benzothiazol-2-yl)propanoic acid (129 mg, 0.624 mmol) and 5-chloro-1-methyl-3-phenyl-1*H*-pyrazole-4-carbaldehyde (152 mg, 0.689 mmol) were dissolved in anhydrous DMF (0.8 mL) and sealed in a microwave vial. TMSCl (419 µL, 3.29 mmol) was added dropwise before heating the reaction to 110 °C for 9 h. (E)-3-(benzo[*d*]thiazol-2-yl)-4-(5-chloro-1-methyl-3-phenyl-1*H*-pyrazol-4-yl)but-3-enoic acid (114 mg, 0.278 mmol, 43%) was obtained as a light brown solid. $^{1}$H NMR (400 MHz, DMSO-$d_6$) δ 12.34 (s, 1H), 8.09 (dd, *J* = 7.9, 1.4 Hz, 1H), 8.02-7.92 (m, 1H), 7.72-7.65 (m, 2H), 7.52 (ddd, *J* = 8.1, 7.2, 1.4 Hz, 1H), 7.48-7.40 (m, 3H), 7.40-7.34 (m, 1H), 7.30 (s, 1H), 3.93 (s, 3H), 3.61 (s, 2H). $^{13}$C NMR (101 MHz,

DMSO-$d_6$) δ 171.1, 168.7, 153.0, 148.3, 134.0, 133.0, 132.6, 128.8, 128.3, 126.9, 126.6, 126.3, 126.3, 125.8, 122.9, 122.2, 110.9, 36.9, 35.6. HRMS (ESI) $C_{21}H_{17}ClN_3O_2S$ $[M+H]^+$ m/z found 410.0714 cald 410.0730.

**(*E*)-3-(benzo[*d*]thiazol-2-yl)-4-(5-(4-chlorophenyl)-2-methyl-2*H*-1,2,3-triazol-4-yl)but-3-enoic acid (16)**

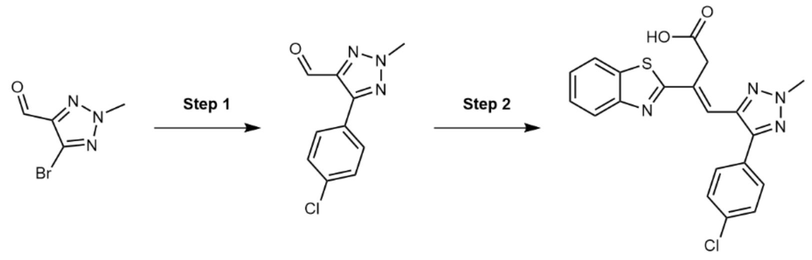


**Step 1**

According to general method D, [1,1′-bis(diphenylphosphino)ferrocene]dichloropalladium(II)·DCM (51.9 mg, 63.5 µmol) was added to 5-bromo-2-methyl-2*H*-1,2,3-triazole-4-carbaldehyde (158 mg, 0.832 mmol), 4-chlorophenylboronic acid (221.1 mg, 1.6 mmol) and 2 M sodium carbonate (480 µL, 0.952 mmol) in DME (1.85 mL). The crude product was purified using flash column chromatography eluting with a gradient of 0-15% EtOAc in n-Hex. 5-(4-chlorophenyl)-2-methyl-2*H*-1,2,3-triazole-4-carbaldehyde (110.8 mg, 0.501 mmol, 60%) was obtained as a colorless solid. $^1H$ NMR (400 MHz, $CDCl_3$) δ 10.18 (s, 1H), 8.04 (d, *J* = 8.4 Hz, 2H), 7.48 – 7.40 (m, 2H), 4.32 (d, *J* = 1.6 Hz, 3H). $^{13}C$ NMR (101 MHz, $CDCl_3$) δ 184.3, 148.1, 143.0, 135.9, 130.0, 129.0, 127.4, 42.7. LCMS: $[M-H]^+$ = 222.0.

**Step 2**

According to general method B, 3-(benzo[*d*]thiazol-2-yl)propanoic acid (68.1 mg, 0.329 mmol) and 5-bromo-2-methyl-2*H*-1,2,3-triazole-4-carbaldehyde (76.5 mg, 0.346 mmol) were dissolved in anhydrous DMF (0.9 mL) and sealed in a microwave vial. TMSCl (260 µL, 2.00 mmol) was added dropwise before heating the reaction to 135 °C for 48 h. (*E*)-3-(benzo[*d*]thiazol-2-yl)-4-(5-(4-chlorophenyl)-2-methyl-2*H*-1,2,3-triazol-4-yl)but-3-enoic acid (128 mg, 0.312 mmol, 95%) was obtained as a grey solid. $^1H$ NMR (400 MHz, DMSO-$d_6$) δ 12.40 (s, 1H), 8.12 – 8.07 (m, 1H), 8.00 (d, *J* = 8.0 Hz, 1H), 7.68 (d, *J* = 8.8 Hz, 2H), 7.65 (d, *J* = 8.9 Hz, 2H), 7.56 – 7.50 (m, 1H), 7.49 – 7.44 (m, 1H), 7.41 (s, 1H), 4.29 (s, 3H), 4.25 (s, 2H). $^{13}C$ NMR (101 MHz, DMSO-$d_6$) δ 171.3, 169.2, 153.2, 146.2, 139.6, 134.1, 133.8, 132.2, 129.9, 129.3, 128.6, 126.7, 125.9, 123.0, 122.2, 122.1, 42.3, 35.5. HRMS (ESI) $C_{20}H_{16}ClN_4O_2S$ $[M+H]^+$ m/z found 411.0682 cald 411.0683.

**(*E*)-3-(benzo[*d*]thiazol-2-yl)-4-(1-phenyl-1*H*-pyrazol-5-yl)but-3-enoic acid (17)**

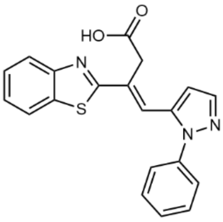

According to general method B, 3-(1,3-benzothiazol-2-yl)propanoic acid (109.3 mg, 0.527 mmol) and 1-phenyl-1*H*-pyrazole-5-carbaldehyde (100 mg, 0.552 mmol) were dissolved in anhydrous DMF (0.514 mL) and sealed in a microwave vial. TMSCl (357 µL, 2.76 mmol) was added dropwise before heating the reaction to 110 °C for 9 h. (*E*)-3-(benzo[*d*]thiazol-2-yl)-4-(1-phenyl-1*H*-pyrazol-5-yl)but-3-enoic acid (50.9 mg, 0.141 mmol, 27%) was obtained as a light pink solid. $^{1}$H NMR (400 MHz, DMSO-$d_6$) δ 12.74 (s, 1H), 8.06 (dd, *J* = 8.0, 1.3 Hz, 1H), 8.02-7.97 (m, 1H), 7.86 (d, *J* = 1.9 Hz, 1H), 7.63-7.54 (m, 4H), 7.54-7.48 (m, 2H), 7.44 (td, *J* = 7.6, 1.3 Hz, 1H), 7.27 (s, 1H), 6.78 (d, *J* = 2.0 Hz, 1H), 4.02 (s, 2H). $^{13}$C NMR (101 MHz, DMSO-$d_6$) δ 171.2, 168.8, 153.1, 140.6, 139.0, 136.8, 134.1, 131.1, 129.5, 128.4, 126.7, 126.0, 125.0, 123.3, 123.0, 122.3, 108.4, 35.5. HRMS (ESI) $C_{20}H_{16}N_3O_2S$ [M+H]$^+$ m/z found 362.0962 cald 362.0963

**(*E*)-3-(benzo[*d*]oxazol-2-yl)-4-(3-(4-chlorophenyl)-1-methyl-1*H*-pyrazol-4-yl)but-3-enoic acid (18)**

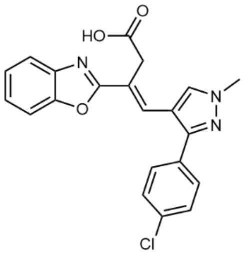

According to general method B, 3-(1,3-benzo[*d*]oxazol-2-yl)propanoic acid (49.7 mg, 0.260 mmol) and 3-(4-chlorophenyl)-1-methyl-1*H*-pyrazole-4-carbaldehyde (59.6 mg, 0.270 mmol) were dissolved in anhydrous DMF (0.9 mL) and sealed in a microwave vial. TMSCl (315 µL, 2.48 mmol) was added dropwise before heating the reaction to 135 °C for 9 h. (*E*)-3-(benzo[*d*]oxazol-2-yl)-4-(3-(4-chlorophenyl)-1-methyl-1*H*-pyrazol-4-yl)but-3-enoic acid (69.1 mg, 0.176 mmol, 68%) was obtained as a colorless solid. $^{1}$H NMR (400 MHz, DMSO-$d_6$) δ 12.44 (s, 1H), 8.15 (s, 1H), 7.78 – 7.67 (m, 2H), 7.61 (s, 1H), 7.59 (s, 4H), 7.43 – 7.30 (m, 2H), 3.98 (s, 3H), 3.83 (s, 2H). $^{13}$C NMR (151 MHz, DMSO) δ 171.5, 164.0, 150.0, 149.1, 141.5, 132.9, 131.9, 131.5, 129.9, 128.9, 127.4, 125.3, 124.7, 119.7, 119.5, 113.4, 110.7, 40.1, 34.2. HRMS (ESI) $C_{21}H_{17}$ $N_3O_3Cl$ [M+H]$^+$ m/z found 394.0951 cald 394.0958.

**(*E*)-3-(5-chlorobenzo[*d*]thiazol-2-yl)-4-(3-(4-chlorophenyl)-1-methyl-1*H*-pyrazol-4-yl)but-3-enoic acid (19)**

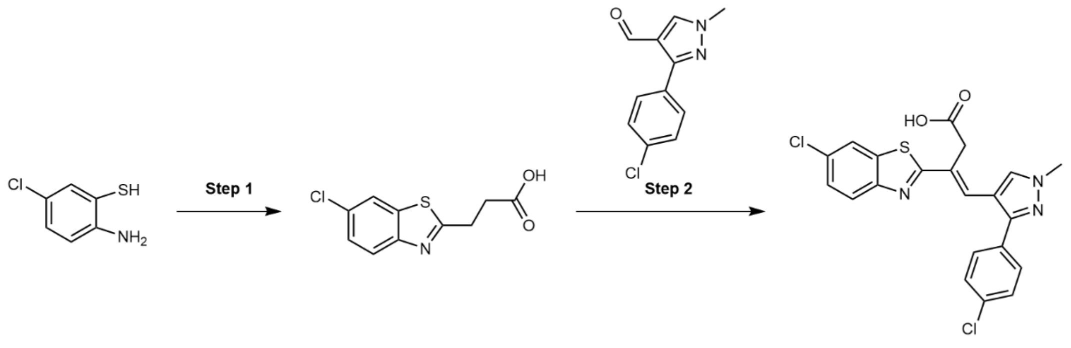


**Step 1**

According to general method C, 2-amino-4-chlorobenzenethiol (320 mg, 2.00 mmol) was dissolved in toluene (20 mL) and succinic anhydride (200 mg, 2.00 mmol) added. 3-(5-chlorobenzo[*d*]thiazol-2-yl)propanoic acid (377 mg, 1.56 mmol, 78%) was obtained as a colorless solid. $^{1}$H NMR (400 MHz, $CDCl_3$) δ 7.97 (d, *J* = 2.0 Hz, 1H), 7.75 (d, *J* = 8.5 Hz, 1H), 7.36 (dd, *J* = 8.5, 2.0 Hz, 1H), 3.44 (t, *J* = 7.1 Hz, 2H), 3.02 (t, *J* = 7.1 Hz, 2H). LCMS: $[M-H]^+$ = 242.0.

**Step 2**

According to general method B, 3-(5-chlorobenzo[*d*]thiazol-2-yl)propanoic acid (61.0 mg, 0.251 mmol) and 3-(4-chlorophenyl)-1-methyl-1*H*-pyrazole-4-carbaldehyde (54.0 mg, 0.265 mmol) were dissolved in anhydrous DMF (0.8 mL) and sealed in a microwave vial. TMSCl (169 µL, 1.33 mmol) was added dropwise before heating the reaction to 120 °C for 10 h. (*E*)-3-(5-chlorobenzo[*d*]thiazol-2-yl)-4-(3-(4-chlorophenyl)-1-methyl-1*H*-pyrazol-4-yl)but-3-enoic acid (66.8 mg, 0.151 mmol, 60%) was obtained as a colorless solid. $^{1}$H NMR (400 MHz, DMSO-*d*$_6$) δ 12.58 (s, 1H), 8.14 (s, 1H), 8.09 (d, *J* = 8.6 Hz, 1H), 8.04 (d, J = 2.0 Hz, 1H), 7.59 (s, 4H), 7.48 (dd, *J* = 8.6, 2.1 Hz, 1H), 7.37 (s, 1H), 3.98 (s, 3H), 3.89 (s, 2H). $^{13}$C NMR (101 MHz, DMSO-*d*$_6$) δ 172.3, 171.5, 154.2, 149.7, 132.9, 132.6, 131.8, 131.5, 131.2, 129.9, 128.9, 128.0, 127.2, 125.5, 123.5, 122.0, 113.6, 35.2. HRMS (ESI) calculated for $C_{21}H_{16}N_3O_2SCl_2$ $[M+H]^+$: 444.0340, found: 444.0327.

**(*E*)-4-(3-(4-chlorophenyl)-1-methyl-1*H*-pyrazol-4-yl)-3-(5-(trifluoromethyl)benzo[*d*]thiazol-2-yl)but-3-enoic acid (20)**

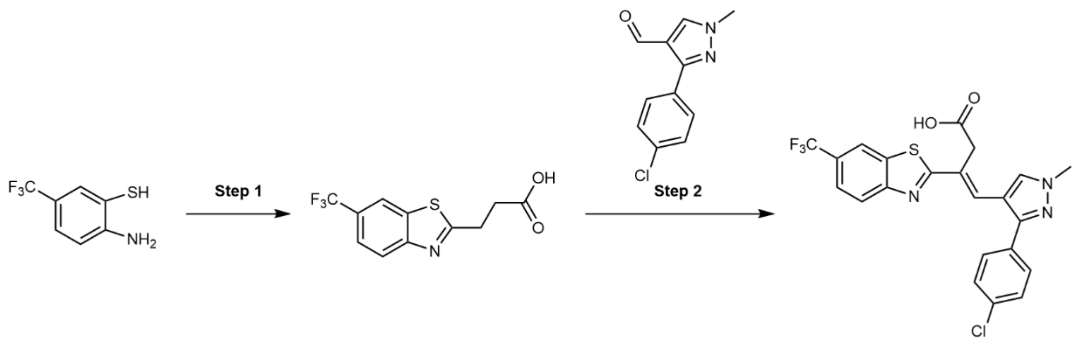


**Step 1**

According to general method C, 2-amino-4-trifluoromethylbenzenethiol (386 mg, 2.00 mmol) was dissolved in toluene (20 mL) and succinic anhydride (200 mg, 2.00 mmol) added. 3-(5-(trifluoromethyl)benzo[*d*]thiazol-2-yl)propanoic acid (176 mg, 0.639 mmol, 32%) was obtained as a colorless solid. $^1$H NMR (400 MHz, $CDCl_3$) δ 8.28 (s, 1H), 7.99 (d, *J* = 8.5 Hz, 1H), 7.65 (d, *J* = 9.0 Hz, 1H), 3.51 (t, *J* = 7.0 Hz, 2H), 3.08 (t, *J* = 7.0 Hz, 2H). LCMS: $[M]^+$ = 275.7.

**Step 2**

According to general method B, 3-(5-(trifluoromethyl)benzo[*d*]thiazol-2-yl)propanoic acid (60.7 mg, 0.221 mmol) and 3-(4-chlorophenyl)-1-methyl-1*H*-pyrazole-4-carbaldehyde (60.2 mg, 0.282 mmol) were dissolved in anhydrous DMF (0.8 mL) and sealed in a microwave vial. TMSCl (169 µL, 1.33 mmol) was added dropwise before heating the reaction to 120 °C for 10 h. (*E*)-4-(3-(4-chlorophenyl)-1-methyl-1*H*-pyrazol-4-yl)-3-(5-(trifluoromethyl)benzo[*d*]thiazol-2-yl)but-3-enoic acid (44.4 mg, 93.1 µmol, 42%) was obtained as a pale orange solid. $^1$H NMR (400 MHz, DMSO-*d*$_6$) δ 12.58 (s, 1H), 8.35-8.28 (m, 2H), 8.17 (s, 1H), 7.75 (dd, *J* = 8.9, 1.5 Hz, 1H), 7.60 (s, 4H), 7.42 (s, 1H), 3.99 (s, 3H), 3.92 (s, 2H). $^{19}$F NMR (377 MHz, DMSO-*d*$_6$) δ -60.11 (s). $^{13}$C[$^{19}$F] NMR (101 MHz, DMSO-*d*$_6$) δ 184.3, 172.6, 171.5, 152.9, 149.7, 138.1, 132.9, 131.9, 131.4, 130.1, 129.9, 128.9, 128.5, 128.4, 127.2, 123.5, 121.4, 119.2, 113.5, 35.4. HRMS (ESI) calculated for $C_{22}H_{16}N_3O_2SClF_3$ $[M+H]^+$: 478.0604, found: 478.0599.

**(*E*)-3-(benzo[*d*]thiazol-2-yl)-4-(3-bromo-1-methyl-1*H*-pyrazol-4-yl)but-3-enoic acid (21)**

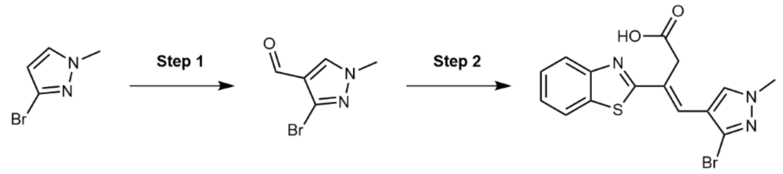


**Step 1**

Anhydrous DMF (3 mL) was cooled to 0 °C and $POCl_3$ (3 mL) was added dropwise. The mixture was allowed to warm to r.t. and stirred for 1 h. 3-bromo-1-methylpyrazole (1.00 g, 6.21 mmol) was added dropwise before the reaction was heated to 95 °C and stirred for 3 h. The reaction was then allowed to cool and quenched by pouring into cold water. The mixture was neutralized with 2M NaOH and the resultant precipitate was collected *in vacuo*, washed with cold water and dried in vacuo. 3-bromo-1-methyl-1*H*-pyrazole-4-carbaldehyde (929 mg, 4.92 mmol, 79%) was obtained as a brown solid. $^1$H NMR (400 MHz, DMSO-$d_6$) δ 9.69 (s, 1H), 8.46 (s, 1H), 3.89 (s, 3H). LCMS: $[M-H]^+$ = 189.0.

**Step 2**

According to general method B, 3-(1,3-benzothiazol-2-yl)propanoic acid (97.2 mg, 0.469 mmol) and 3-bromo-1-methyl-1*H*-pyrazole-4-carbaldehyde (100 mg, 0.529 mmol) were dissolved in anhydrous DMF (0.640 mL) and sealed in a microwave vial. TMSCl (337 µL, 2.60 mmol) was added dropwise before heating the reaction to 110 °C for 9 h. (*E*)-3-(benzo[*d*]thiazol-2-yl)-4-(3-bromo-1-methyl-1*H*-pyrazol-4-yl)but-3-enoic acid (89.0 mg, 0.235 mmol, 50%) was obtained as a yellow solid. $^1$H NMR (400 MHz, MeOD) δ 7.99-7.91 (m, 3H), 7.51-7.46 (m, 1H), 7.41 (td, *J* = 7.6, 1.2 Hz, 1H), 7.32 (s, 1H), 3.97 (s, 2H), 3.94 (s, 3H). $^{13}$C NMR (126 MHz, MeOD) δ 174.0, 171.4, 154.9, 135.7, 132.9, 129.8, 129.3, 127.5, 127.0, 126.7, 123.8, 122.7, 118.0, 40.0, 36.6. HRMS (ESI) $C_{15}H_{13}BrN_3O_2S$ $[M+H]^+$ m/z found 377.9919 cald 377.9912.

**(*E*)-3-(benzo[*d*]thiazol-2-yl)-4-(1-methyl-3-phenyl-1*H*-pyrazol-4-yl)but-3-enoic acid (22)**

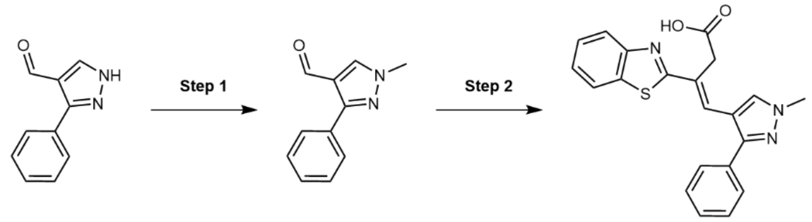


**Step 1**

According to general method A, 3-phenyl-1*H*-pyrazole-4-carbaldehyde (300 mg, 1.74 mmol) was reacted with cesium carbonate (1.18 g, 3.48 mmol) and iodomethane (171 µL, 2.61 mmol) in anhydrous DMF (14 mL). The crude product was purified using flash column chromatography eluting with a gradient of 0-45% EtOAc in n-Hex. 1-methyl-3-phenyl-1*H*-pyrazole-4-carbaldehyde (154 mg, 0.826 mmol, 48%) was obtained as a yellow oil. $^1$H NMR (400 MHz, $CDCl_3$) δ 9.89 (d, *J* = 1.3 Hz, 1H), 7.96 (s, 1H), 7.74-7.64 (m, 2H), 7.48-7.33 (m, 3H), 3.92 (s, 3H). LCMS: $[M-H]^+$ = 187.1.

**Step 2**

According to general method B, 3-(1,3-benzothiazol-2-yl)propanoic acid (151 mg, 0.726 mmol) and 1-methyl-3-phenyl-1*H*-pyrazole-4-carbaldehyde (154 mg, 0.826 mmol) were dissolved in anhydrous DMF (0.966 mL) and sealed in a microwave vial. TMSCl (508 µL, 3.92 mmol) was added dropwise before heating the reaction to 120 °C for 9 h. (E)-3-(benzo[*d*]thiazol-2-yl)-4-(1-methyl-3-phenyl-1*H*-pyrazol-4-yl)but-3-enoic acid (142 mg, 0.378 mmol, 52%) was obtained as a brown solid. $^1$H NMR (400 MHz, DMSO-$d_6$) δ 12.52 (s, 1H), 8.11 (s, 1H), 8.03 (dd, *J* = 8.0, 1.3 Hz, 1H), 7.97-7.90 (m, 1H), 7.57 (dt, *J* = 6.4, 1.5 Hz, 2H), 7.54-7.38 (m, 5H), 7.36 (s, 1H), 3.97 (s, 3H), 3.91 (s, 2H). $^{13}$C NMR (101 MHz, DMSO-$d_6$) δ 171.6, 170.0, 153.3, 150.8, 133.8, 132.7, 131.5, 128.8, 128.2, 128.1, 127.6, 127.1, 126.5, 125.4, 122.6, 122.0, 113.5, 39.1, 35.2. HRMS (ESI) $C_{21}H_{18}N_3O_2S$ $[M+H]^+$ m/z found 376.1122 cald 376.1120.

**(*E*)-3-(benzo[*d*]thiazol-2-yl)-4-(3-(4-fluorophenyl)-1-methyl-1*H*-pyrazol-4-yl)but-3-enoic acid (23)**

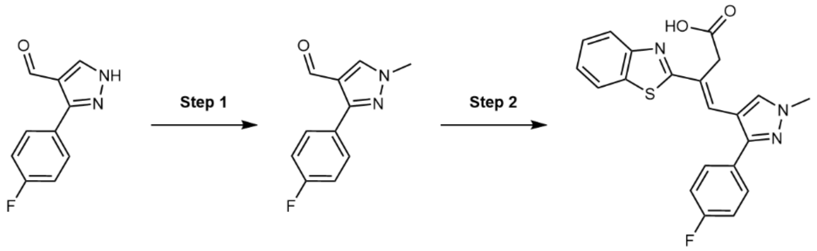

**Step 1**

According to general method A, 3-(4-fluorophenyl)-1*H*-pyrazole-4-carbaldehyde (300 mg, 1.58 mmol) was reacted with cesium carbonate (1.07 g, 3.16 mmol) and iodomethane (155 µL, 2.37 mmol) in anhydrous DMF (12 mL). The crude product was purified using flash column chromatography eluting with a gradient of 0-50% EtOAc in n-Hex. 3-(4-fluorophenyl)-1-methyl-1*H*-pyrazole-4-carbaldehyde (111 mg, 0.543 mmol, 34%) was obtained as a colorless solid. $^{1}$H NMR (400 MHz, $CDCl_3$) δ 9.89 (s, 1H), 7.98 (s, 1H), 7.77-7.68 (m, 2H), 7.13 (t, *J* = 8.8 Hz, 2H), 3.97 (s, 3H). LCMS: $[M-H]^+$ = 205.2.

**Step 2**

According to general method B, 3-(1,3-benzothiazol-2-yl)propanoic acid (99 mg, 0.478 mmol) and 3-(4-fluorophenyl)-1-methyl-1*H*-pyrazole-4-carbaldehyde (111 mg, 0.543 mmol) were dissolved in anhydrous DMF (0.634 mL) and sealed in a microwave vial. TMSCl (334 µL, 2.58 mmol) was added dropwise before heating the reaction to 120 °C for 9 h. (*E*)-3-(benzo[*d*]thiazol-2-yl)-4-(3-(4-fluorophenyl)-1-methyl-1*H*-pyrazol-4-yl)but-3-enoic acid was obtained as a colorless solid. $^{1}$H NMR (400 MHz, DMSO-*d*$_6$) δ 12.53 (s, 1H), 8.11 (s, 1H), 8.03 (dd, *J* = 7.9, 1.3 Hz, 1H), 7.97- 7.89 (m, 1H), 7.64- 7.56 (m, 2H), 7.48 (ddd, *J* = 8.3, 7.2, 1.4 Hz, 1H), 7.41 (ddd, *J* = 8.4, 7.3, 1.3 Hz, 1H), 7.38- 7.29 (m, 3H), 3.96 (s, 3H), 3.90 (s, 2H). $^{19}$F NMR (377 MHz, DMSO-*d*$_6$) δ 108.99-109.24 (m). $^{13}$C NMR (101 MHz, DMSO-*d*$_6$) δ 171.6, 169.9, 162.0 (d, J = 244.9 Hz), 153.3, 149.8, 133.8, 131.5, 130.2 (d, *J* = 8.2 Hz), 129.2, 127.4, 126.5, 125.5, 122.6, 122.0, 115.7 (d, *J* = 21.5 Hz), 113.5, 39.1, 35.2. HRMS (ESI) $[M+H]^+$ $C_{21}H_{17}FN_3O_2S$ m/z found 394.1027 cald 394.1026.

**(*E*)-3-(benzo[*d*]thiazol-2-yl)-4-(1-methyl-3-(*p*-tolyl)-1*H*-pyrazol-4-yl)but-3-enoic acid (24)**

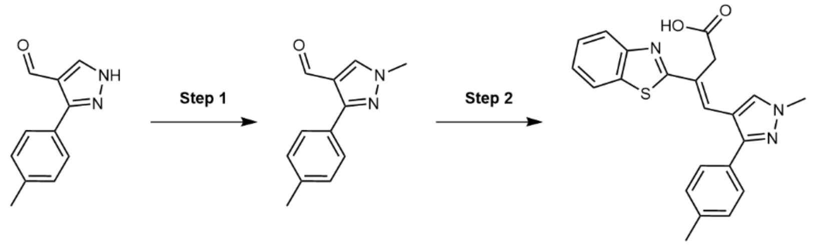

**Step 1**

According to general method A, 3-(*p*-tolyl)-1*H*-pyrazole-4-carbaldehyde (300 mg, 1.61 mmol) was reacted with cesium carbonate (1.09 g, 3.22 mmol) and iodomethane (158 µL, 2.47 mmol) in anhydrous DMF (13 mL). The crude product was purified using flash column chromatography eluting with a gradient of 0-50% EtOAc in n-Hex. 1-methyl-3-(*p*-tolyl)-1*H*-pyrazole-4-carbaldehyde (150 mg, 0.750 mmol, 47%) was obtained as a colorless solid. $^1$H NMR (400 MHz, $CDCl_3$) δ 9.90 (s, 1H), 7.96 (s, 1H), 7.61-7.54 (m, 2H), 7.25 (d, J = 8.0 Hz, 2H), 3.95 (s, 3H), 2.38 (s, 3H). LCMS: $[M-H]^+$ = 201.1.

**Step 2**

According to general method B, 3-(1,3-benzothiazol-2-yl)propanoic acid (136 mg, 0.657 mmol) and 1-methyl-3-(p-tolyl)-1*H*-pyrazole-4-carbaldehyde (150 mg, 0.750 mmol) were dissolved in anhydrous DMF (0.8 mL) and sealed in a microwave vial. TMSCl (461 µL, 3.56 mmol) was added dropwise before heating the reaction to 120 °C for 12 h. (*E*)-3-(benzo[*d*]thiazol-2-yl)-4-(1-methyl-3-(*p*-tolyl)-1*H*-pyrazol-4-yl)but-3-enoic acid (15.3 mg, 37.3 µmol, 6%) was obtained as a yellow solid. $^1$H NMR (400 MHz, DMSO-*d*$_6$) δ 12.52 (s, 1H), 8.09 (s, 1H), 8.03 (d, *J* = 7.9 Hz, 1H), 7.93 (d, *J* = 8.1 Hz, 1H), 7.52-7.43 (m, 3H), 7.43-7.38 (m, 1H), 7.36-7.29 (m, 3H), 3.96 (s, 3H), 3.90 (s, 2H), 2.37 (s, 3H). $^{13}$C NMR (126 MHz, DMSO-*d*$_6$) δ 171.6, 170.0, 153.3, 150.9, 137.5, 133.8, 131.4, 129.3, 128.1, 127.7, 126.8, 126.4, 125.4, 122.5, 122.0, 113.4, 39.0, 35.2, 20.8. HRMS (ESI) $C_{22}H_{20}N_3O_2S$ $[M+H]^+$ m/z found 390.1275 cald 390.1276.

**(*E*)-3-(benzo[*d*]thiazol-2-yl)-4-(1-methyl-3-(4-(trifluoromethyl)phenyl)-1*H*-pyrazol-4-yl)but-3-enoic acid (25)**

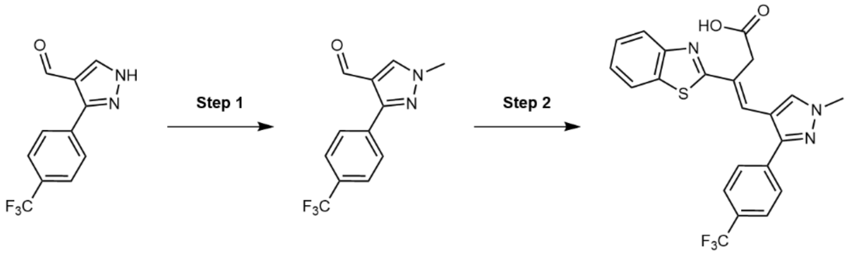


**Step 1**

According to general method A, 3-(4-(trifluoromethyl)phenyl)-1*H*-pyrazole-4-carbaldehyde (348 mg, 1.45 mmol) was reacted with cesium carbonate (945 mg, 2.90 mmol) and iodomethane (135 µL, 2.18 mmol) in anhydrous DMF (10 mL). The crude product was purified using flash column chromatography eluting with a gradient of 15-40% EtOAc in n-Hex. 1-methyl-3-(4-(trifluoromethyl)phenyl)-1*H*-pyrazole-4-carbaldehyde (151 mg, 0.594 mmol, 41%) was obtained as a colorless crystalline solid. $^1$H NMR (400 MHz, $CDCl_3$) δ 9.93 (s, 1H), 8.02 (s, 1H), 7.91 (d, *J* = 8.2 Hz, 2H), 7.71 (d, *J* = 8.2 Hz, 2H), 4.01 (s, 3H). LCMS: $[M-H]^+$ = 255.2.

**Step 2**

According to general method B, 3-(1,3-benzothiazol-2-yl)propanoic acid (54.6 mg, 0.263 mmol) and 1-methyl-3-(4-(trifluoromethyl)phenyl)-1*H*-pyrazole-4-carbaldehyde (70.6 mg, 0.277 mmol) were dissolved in anhydrous DMF (1 mL) and sealed in a microwave vial. TMSCl (174 µL, 1.39 mmol) was added dropwise before heating the reaction to 135 °C for 18 h. (*E*)-3-(benzo[*d*]thiazol-2-yl)-4-(1-methyl-3-(4-(trifluoromethyl)phenyl)-1*H*-pyrazol-4-yl)but-3-enoic acid (82.9 mg, 0.187 mmol, 71%) was obtained as an off-white solid. $^1$H NMR (400 MHz, DMSO-*d*$_6$) δ 12.55 (s, 1H), 8.15 (s, 1H), 8.06 (d, *J* = 7.8 Hz, 1H), 7.96 (d, *J* = 8.1 Hz, 1H), 7.89 (d, *J* = 8.2 Hz, 2H), 7.82 (d, *J* = 8.2 Hz, 2H), 7.54-7.46 (m, 1H), 7.45-7.41 (m, 1H), 7.39 (s, 1H), 4.01 (s, 3H), 3.92 (s, 2H). $^{19}$F NMR (377 MHz, DMSO-*d*$_6$) δ -60.99 (s). $^{13}$C NMR (400 MHz, DMSO-*d*$_6$) δ 171.5, 169.8, 153.2, 148.9,

136.7, 133.9, 131.9, 128.6, 128.52 – 128.34 (m), 128.2, 127.0, 126.5, 125.7, 125.5, 123.0, 122.6, 122.1, 114.0, 39.2, 35.3. HRMS (ESI) calculated for $C_{22}H_{17}N_3O_2SF_3$ $[M+H]^+$: 444.0994, found: 444.0996.

**(*E*)-3-(benzo[*d*]thiazol-2-yl)-4-(3-(4-methoxyphenyl)-1*H*-pyrazol-4-yl)but-3-enoic acid (26)**

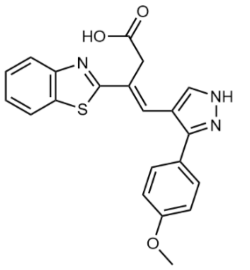

According to general method B, 3-(1,3-benzothiazol-2-yl)propanoic acid (103 mg, 0.485 mmol) and 3-(4-Methoxyphenyl)-1*H*-pyrazole-4-carbaldehyde (100 mg, 0.485 mmol) were dissolved in anhydrous DMF (0.5 mL) and sealed in a microwave vial. TMSCl (314 µL, 2.42 mmol) was added dropwise before heating the reaction to 135 °C for 6 h. (*E*)-3-(benzo[*d*]thiazol-2-yl)-4-(3-(4-methoxyphenyl)-1*H*-pyrazol-4-yl)but-3-enoic acid 16.8 mg, 42.9 µmol, 8%) was obtained as a yellow solid. $^1$H NMR (400 MHz, DMSO-$d_6$) δ 12.51 (s, 1H), 8.03 (d, $J$ = 7.9 Hz, 1H), 7.96-7.89 (m, 2H), 7.53-7.46 (m, 3H), 7.43-7.35 (m, 2H), 7.12 (d, $J$ = 8.4 Hz, 2H), 3.94 (s, 2H), 3.83 (s, 3H). $^{13}$C NMR (101 MHz, DMSO-$d_6$) δ 171.8, 170.2, 159.5, 153.3, 133.8, 129.6, 127.9, 126.7, 126.4, 125.4, 122.5, 122.0, 114.4, 112.9, 55.3, 35.3. HRMS (ESI) $C_{21}H_{18}N_3O_3S$ $[M+H]^+$ m/z found 392.1065 cald 392.1069.

**(*E*)-3-(benzo[*d*]thiazol-2-yl)-4-(3-(4-cyanophenyl)-1-methyl-1*H*-pyrazol-4-yl)but-3-enoic acid (27)**

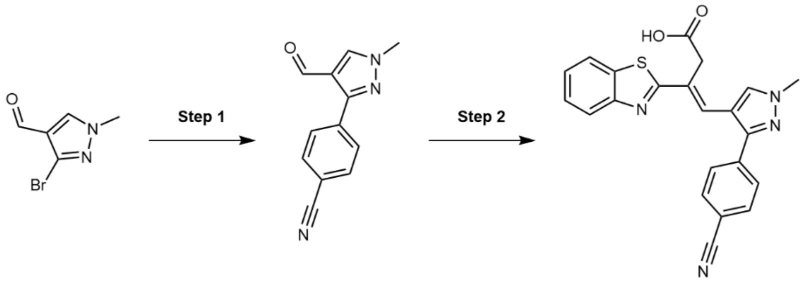


**Step 1**

According to general method D, [1,1′-bis(diphenylphosphino)ferrocene]dichloropalladium(II)·DCM (51.9 mg, 63.5 µmol) was added to 3-bromo-1-methyl-1*H*-pyrazole-4-carbaldehyde (178 mg, 0.940 mmol), 4-cyanophenylboronic acid (179.9 mg, 1.22 mmol) and 2 M sodium carbonate (480 µL, 0.952 mmol) in DME (1.85 mL). The crude product was purified using flash column chromatography eluting with a gradient of 0-40% EtOAc in n-Hex. 4-(4-formyl-1-methyl-1*H*-pyrazol-3-yl)benzonitrile (100 mg, 0.474 mmol, 50%) was obtained as a colorless solid. $^1$H NMR (400 MHz, DMSO-$d_6$) δ 9.88 (s, 1H), 8.61 (s, 1H), 8.15 – 8.04 (m, 2H), 7.97 – 7.89 (m, 2H), 3.98 (s, 3H). $^{13}$C NMR (101 MHz, $CDCl_3$) δ 183.6, 151.1, 136.8, 136.1, 132.4, 129.3, 121.5, 118.7, 112.5, 39.8, 29.7. LCMS: $[M-H]^+$ = 212.0.

**Step 2**

According to general method B, 3-(benzo[*d*]thiazol-2-yl)propanoic acid (72.5 mg, 0.350 mmol) and 4-(4-formyl-1-methyl-1*H*-pyrazol-3-yl)benzonitrile (77.0 mg, 0.365 mmol) were dissolved in anhydrous DMF (0.9 mL) and sealed in a microwave vial. TMSCl (280 µL, 2.2 mmol) was added dropwise before heating the reaction to 135 °C for 48 h. (*E*)-3-(benzo[*d*]thiazol-2-yl)-4-(3-(4-cyanophenyl)-1-methyl-1*H*-pyrazol-4-yl)but-3-enoic acid (77.0 mg, 0.192 mmol, 55%) was obtained as an off-white solid. $^1$H NMR (400 MHz, DMSO-$d_6$) δ 8.14 (s, 1H), 8.05 (d, *J* = 7.8 Hz, 1H), 7.98 (s, 1H), 7.95 (d, *J* = 6.5 Hz, 3H), 7.78 (d, *J* = 8.2 Hz, 2H), 7.50 (t, *J* = 7.7 Hz, 1H), 7.43 (t, *J* = 7.5 Hz, 1H), 7.36 (s, 1H), 3.99 (s, 3H), 3.88 (s, 2H). $^{13}$C NMR (101 MHz, DMSO-$d_6$) δ 171.9, 170.1, 153.6, 148.9, 137.7, 134.3, 133.2, 132.5, 129.0, 128.9, 127.4, 126.9, 126.0, 123.1, 122.5, 119.2, 114.6, 110.9, 39.6, 35.7. HRMS (ESI) $C_{22}H_{17}N_4O_2S$ [M+H]$^+$ m/z found 401.1076 cald 401.1072.

**(*E*)-3-(benzo[*d*]thiazol-2-yl)-4-(3-(3-chlorophenyl)-1-methyl-1*H*-pyrazol-4-yl)but-3-enoic acid (28)**

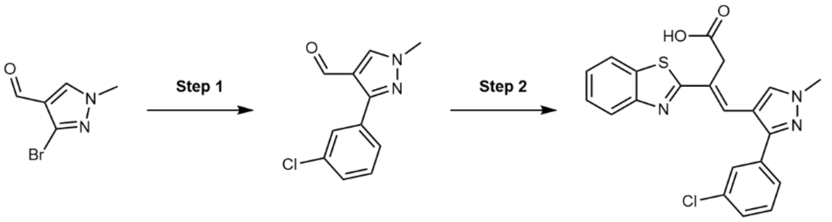


**Step 1**

According to general method D, [1,1′-bis(diphenylphosphino)ferrocene]dichloropalladium(II)·DCM (51.9 mg, 63.5 µmol) was added to 3-bromo-1-methyl-1*H*-pyrazole-4-carbaldehyde (120 mg, 0.635 mmol), 3-chlorophenylboronic acid (199 mg, 1.27 mmol), and 2 M sodium carbonate (480 µL, 0.952 mmol) in DME (1.85 mL). The crude product was purified using flash column chromatography eluting with a gradient of 0-50% EtOAc in n-Hex. 3-(3-chlorophenyl)-1-methyl-1*H*-pyrazole-4-carbaldehyde (97.0 mg, 0.440 mmol, 69%) was obtained as a yellow oil. $^1$H NMR (400 MHz, $CDCl_3$) δ 9.90 (s, 1H), 7.99 (s, 1H), 7.75 (dt, *J* = 1.7, 1.1 Hz, 1H), 7.67-7.57 (m, 1H), 7.43-7.35 (m, 2H), 3.98 (d, *J* = 8.9 Hz, 3H). LCMS: [M-H]$^+$ = 221.0.

**Step 2**

According to general method B, 3-(1,3-benzothiazol-2-yl)propanoic acid (80.2 mg, 0.387 mmol) and 3-(3-chlorophenyl)-1-methyl-1*H*-pyrazole-4-carbaldehyde (97.0 mg, 0.440 mmol) were dissolved in anhydrous DMF (0.514 mL) and sealed in a microwave vial. TMSCl (271 µL, 2.09 mmol) was added dropwise before heating the reaction to 110 °C for 9 h. (*E*)-3-(benzo[*d*]thiazol-2-yl)-4-(3-(3-chlorophenyl)-1-methyl-1*H*-pyrazol-4-yl)but-3-enoic acid (21.9 mg, 53.4 µmol, 14%) was obtained as a dark green solid. $^1$H NMR (400 MHz, DMSO-$d_6$) δ 12.51 (s, 1H), 8.13 (d, *J* = 6.1 Hz, 1H), 8.05 (dd, *J* = 7.9, 1.3 Hz, 1H), 7.95 (d, *J* = 8.1 Hz, 1H), 7.62-7.58 (m, 1H), 7.55-7.46 (m, 4H), 7.44-7.38 (m, 1H), 7.35 (s, 1H), 3.97 (s, 3H), 3.89 (s, 2H). $^{13}$C NMR (101 MHz, DMSO-$d_6$) δ 171.6, 169.8, 153.2, 148.9, 134.7, 133.8, 133.5, 131.8, 130.7, 127.9, 127.6, 127.2, 126.7, 126.5, 125.5, 122.6, 122.1, 113.7, 39.2, 35.2. HRMS (ESI) $C_{21}H_{17}ClN_3O_2S$ [M+H]$^+$ m/z found 410.0734 cald 410.0730.

**(*E*)-3-(benzo[*d*]thiazol-2-yl)-4-(3-(3-fluorophenyl)-1-methyl-1*H*-pyrazol-4-yl)but-3-enoic acid (29)**

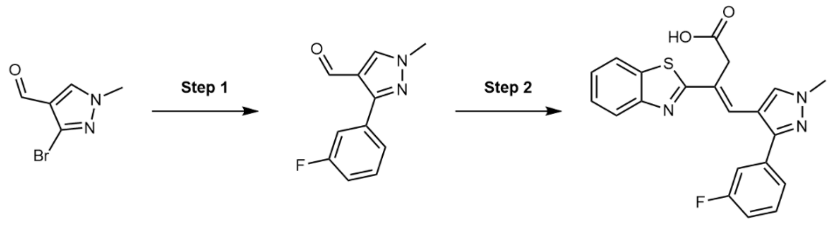


**Step 1**

According to general method D, [1,1′-bis(diphenylphosphino)ferrocene]dichloropalladium(II)·DCM (51.9 mg, 63.5 µmol) was added to 3-bromo-1-methyl-1*H*-pyrazole-4-carbaldehyde (178 mg, 0.940 mmol), 3-fluorophenylboronic acid (224 mg, 1.60 mmol) and 2 M sodium carbonate (480 µL, 0.952 mmol) in DME (1.85 mL). The crude product was purified using flash column chromatography eluting with a gradient of 0-70% EtOAc in n-Hex. 3-(3-fluorophenyl)-1-methyl-1*H*-pyrazole-4-carbaldehyde (97.8 mg, 0.479 mmol, 51%) was obtained as a colorless solid. $^{1}$H NMR (400 MHz, DMSO-$d_6$) δ 9.86 (s, 1H), 8.56 (s, 1H), 7.72 (s, 1H), 7.70 (s, 1H), 7.50 (q, *J* = 7.4 Hz, 1H), 7.26 (dt, *J* = 9.2, 5.0 Hz, 1H), 3.96 (s, 3H). $^{13}$C NMR (101 MHz, DMSO-$d_6$) δ 184.2, 163.3, 160.9, 149.9, 139.0, 134.0 (d, *J* = 8.8 Hz), 130.4 (d, *J* = 8.4 Hz), 124.4, 120.5, 115.2 (dd, *J* = 58.1, 22.0 Hz), 39.2. $^{19}$F NMR (377 MHz, DMSO-$d_6$) δ -113.08 (q, *J* = 9.2 Hz). LCMS: $[M-H]^+$ = 205.0.

**Step 2**

According to general method B, 3-(benzo[*d*]thiazol-2-yl)propanoic acid (90.6 mg, 0.429 mmol) and 3-(3-fluorophenyl)-1-methyl-1*H*-pyrazole-4-carbaldehyde (94.0 mg, 0.451 mmol) were dissolved in anhydrous DMF (0.9 mL) and sealed in a microwave vial. TMSCl (350 µL, 1.9 mmol) was added dropwise before heating the reaction to 135 °C for 48 h. (*E*)-3-(benzo[*d*]thiazol-2-yl)-4-(3-(3-fluorophenyl)-1-methyl-1*H*-pyrazol-4-yl)but-3-enoic acid (88.5 mg, 0.225 mmol, 52%) was obtained as a brown solid. $^{1}$H NMR (400 MHz, DMSO-$d_6$) δ 12.52 (s, 1H), 8.12 (s, 1H), 8.06 (d, *J* = 7.8 Hz, 1H), 7.94 (t, *J* = 8.3 Hz, 1H), 7.60 – 7.54 (m, 1H), 7.55 – 7.49 (m, 1H), 7.49 – 7.45 (m, 1H), 7.42 (s, 1H), 7.40 (t, *J* = 1.3 Hz, 1H), 7.38 (d, *J* = 4.8 Hz, 1H), 7.33 – 7.24 (m, 1H), 3.98 (s, 3H), 3.90 (s, 2H). $^{13}$C NMR (101 MHz, DMSO-$d_6$) δ 171.5, 169.8, 162.3 (d, *J* = 243.8 Hz), 153.2, 149.2, 135.0 (d, *J* = 8.4 Hz), 133.8, 131.71, 130.8 (d, *J* = 8.8 Hz), 129.6, 127.8, 127.2, 126.5, 126.0 – 124.8 (m), 125.5, 124.2, 122.6, 114.7 (dd, *J* = 35.8, 21.7 Hz), 114.0, 113.7, 39.1. $^{19}$F NMR (377 MHz, DMSO-$d_6$) δ -112.5 – -112.7 (m). HRMS (ESI) $C_{21}H_{17}N_3O_2FS$ $[M+H]^+$ m/z found 394.1020 cald 394.1026.

**(*E*)-3-(benzo[*d*]thiazol-2-yl)-4-(1-methyl-3-(m-tolyl)-1*H*-pyrazol-4-yl)but-3-enoic acid (30)**

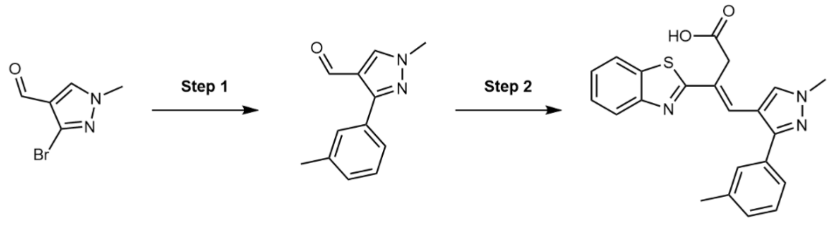


**Step 1**

According to general method D, [1,1′-bis(diphenylphosphino)ferrocene]dichloropalladium(II)·DCM (112 mg, 0.138 mmol) was added to 3-bromo-1-methyl-1*H*-pyrazole-4-carbaldehyde (260 mg, 1.38 mmol), m-tolylboronic acid (374 mg, 2.75 mmol), and 2 M sodium carbonate (1.03 mL, 2.06 mmol) in DME (4 mL). The crude product purified using flash column chromatography eluting with a gradient of 0-50% EtOAc in n-Hex. 1-methyl-3-(*m*-tolyl)-1*H*-pyrazole-4-carbaldehyde (199 mg, 0.995 mmol, 72%) was obtained as a yellow solid. $^{1}H$ NMR (400 MHz, $CDCl_3$) δ 9.92 (s, 1H), 8.00 (s, 1H), 7.54-7.45 (m, 2H), 7.35 (t, *J* = 7.6 Hz, 1H), 7.28-7.22 (m, 1H), 3.97 (s, 3H), 2.41 (s, 3H). LCMS: $[M-H]^+$ = 201.1.

**Step 2**

According to general method B, 3-(1,3-benzothiazol-2-yl)propanoic acid (200 mg, 0.965 mmol) and 1-methyl-3-(*m*-tolyl)-1*H*-pyrazole-4-carbaldehyde (199 mg, 0.995 mmol) were dissolved in anhydrous DMF (1.2 mL) and sealed in a microwave vial. TMSCl (612 µL, 4.73 mmol) was added dropwise before heating the reaction to 110 °C for 24 h. (*E*)-3-(benzo[*d*]thiazol-2-yl)-4-(1-methyl-3-(*m*-tolyl)-1*H*-pyrazol-4-yl)but-3-enoic acid (177 mg, 0.455 mmol, 47%) was obtained as a green solid. $^{1}H$ NMR (400 MHz, DMSO-$d_6$) δ 12.46 (s, 1H), 8.10 (s, 1H), 8.06-8.02 (m, 1H), 7.92 (t, *J* = 8.4 Hz, 1H), 7.48 (ddd, *J* = 8.3, 7.2, 1.4 Hz, 1H), 7.43-7.32 (m, 5H), 7.25 (d, *J* = 7.3 Hz, 1H), 3.96 (s, 3H), 3.90 (s, 2H), 2.38 (s, 3H). $^{13}C$ NMR (101 MHz, DMSO-$d_6$) δ 172.1, 170.5, 153.7, 151.3, 138.4, 134.2, 133.0, 131.9, 129.3, 129.2, 129.1, 128.2, 127.4, 126.9, 125.8, 123.0, 122.5, 113.9, 39.5, 35.7, 32.7, 21.6. HRMS (ESI) $C_{22}H_{20}N_3O_2S$ $[M+H]^+$ m/z found 390.1272 cald 390.1276.

**(*E*)-3-(benzo[*d*]thiazol-2-yl)-4-(3-(3-methoxyphenyl)-1-methyl-1*H*-pyrazol-4-yl)but-3-enoic acid (31)**

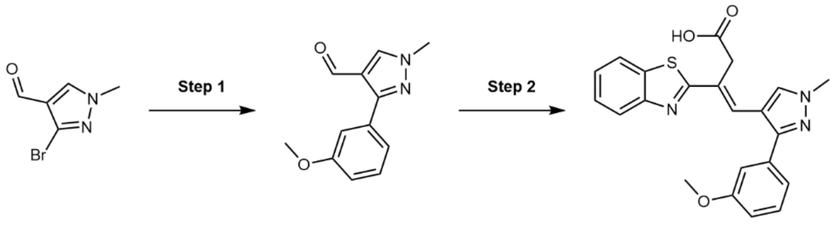


**Step 1**

According to general method D, [1,1′-bis(diphenylphosphino)ferrocene]dichloropalladium(II)·DCM (51.9 mg, 63.5 µmol) was added to 3-bromo-1-methyl-1*H*-pyrazole-4-carbaldehyde (178 mg, 0.940 mmol), 3-methoxyphenylboronic acid (186 mg, 1.22 mmol) and 2 M sodium carbonate (480 µL, 0.952 mmol) in DME (1.85 mL). The crude product was purified using flash column chromatography eluting with a gradient of 0-60% EtOAc in n-Hex. 3-(3-methoxyphenyl)-1-methyl-1*H*-pyrazole-4-carbaldehyde (102 mg, 0.472 mmol, 50%)

was obtained as a colorless solid. $^1$H NMR (400 MHz, DMSO-$d_6$) δ 9.85 (s, 1H), 8.51 (s, 1H), 7.39 (q, *J* = 1.2 Hz, 1H), 7.38 (d, *J* = 2.0 Hz, 1H), 7.35 (d, *J* = 7.9 Hz, 1H), 6.99 (dt, *J* = 6.8, 2.5 Hz, 1H), 3.94 (s, 3H), 3.80 (s, 3H). $^{13}$C NMR (101 MHz, DMSO-$d_6$) δ 184.4, 159.3, 151.5, 137.9, 133.0, 129.6, 120.8, 120.4, 114.6, 113.6, 55.1, 39.1. LCMS: $[M-H]^+$ = 217.0.

**Step 2**

According to general method B, 3-(benzo[*d*]thiazol-2-yl)propanoic acid (66.4 mg, 0.320 mmol) and 3-(3-methoxyphenyl)-1-methyl-1*H*-pyrazole-4-carbaldehyde (73.0 mg, 0.331 mmol) were dissolved in anhydrous DMF (0.9 mL) and sealed in a microwave vial. TMSCl (250 µL, 1.9 mmol) was added dropwise before heating the reaction to 135 °C for 48 h. (*E*)-3-(benzo[*d*]thiazol-2-yl)-4-(3-(3-methoxyphenyl)-1-methyl-1*H*-pyrazol-4-yl)but-3-enoic acid (73.5 mg, 0.181 mmol, 57%) was obtained as a brown solid. $^1$H NMR (400 MHz, DMSO-$d_6$) δ 12.52 (s, 1H), 8.12 (s, 1H), 8.05 (d, *J* = 7.8 Hz, 1H), 7.98 – 7.90 (m, 1H), 7.49 (t, *J* = 7.5 Hz, 1H), 7.45 – 7.35 (m, 3H), 7.18 – 7.11 (m, 2H), 7.01 (d, *J* = 8.4 Hz, 1H), 3.98 (s, 3H), 3.89 (s, 2H), 3.80 (s, 3H). $^{13}$C NMR (101 MHz, DMSO-$d_6$) δ 171.6, 170.0, 159.4, 153.3, 150.5, 134.0, 133.8, 131.5, 129.8, 127.5, 127.4, 126.4, 125.4, 124.8, 122.5, 122.0, 120.5, 114.0, 113.6, 113.1, 55.1, 39.0. HRMS (ESI) $C_{22}H_{20}N_3O_3S$ $[M+H]^+$ m/z found 406.1234 cald 406.1225.

**(*E*)-3-(benzo[*d*]thiazol-2-yl)-4-(3-(3-cyanophenyl)-1-methyl-1*H*-pyrazol-4-yl)but-3-enoic acid (32)**

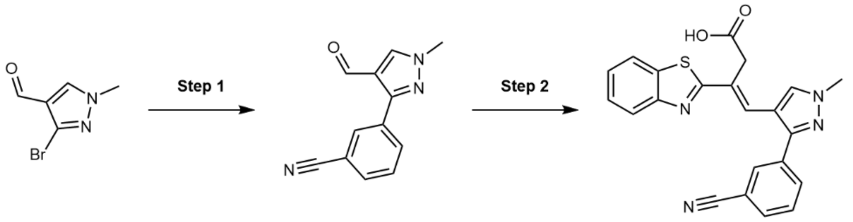


**Step 1**

According to general method D, [1,1′-bis(diphenylphosphino)ferrocene]dichloropalladium(II)·DCM (51.9 mg, 63.5 µmol) was added to 3-bromo-1-methyl-1*H*-pyrazole-4-carbaldehyde (120 mg, 0.635 mmol), 3-Cyanophenylboronic acid (187 mg, 1.27 mmol) and 2 M sodium carbonate (480 µL, 0.952 mmol) in DME (1.85 mL). The crude product was purified using flash column chromatography eluting with a gradient of 0-50% EtOAc in n-Hex. 3-(4-formyl-1-methyl-1*H*-pyrazol-3-yl)benzonitrile (57.2 mg, 0.271 mmol, 43%) was obtained as a colorless solid. $^1$H NMR (400 MHz, $CDCl_3$) δ 9.91 (s, 1H), 8.14 (t, *J* = 1.8 Hz, 1H), 8.08 (dt, *J* = 7.8, 1.5 Hz, 1H), 8.02 (s, 1H), 7.68 (dt, *J* = 7.8, 1.4 Hz, 1H), 7.55 (t, *J* = 7.8 Hz, 1H), 4.00 (s, 3H). LCMS: $[M-H]^+$ = 212.2.

**Step 2**

According to general method B, 3-(1,3-benzothiazol-2-yl)propanoic acid (49.7 mg, 0.240 mmol) and 3-(4-formyl-1-methyl-1*H*-pyrazol-3-yl)benzonitrile (57.2 mg, 0.271 mmol) were dissolved in anhydrous DMF (0.317 mL) and sealed in a microwave vial. TMSCl (167 µL, 1.29 mmol) was added dropwise before heating the reaction to 110 °C for 9 h. (*E*)-3-(benzo[*d*]thiazol-2-yl)-4-(3-(3-cyanophenyl)-1-methyl-1*H*-pyrazol-4-yl)but-3-enoic acid (11.2 mg, 28.0 µmol, 12%) was obtained as a light brown solid. $^1$H NMR (400 MHz, DMSO-$d_6$) δ 12.53 (s, 1H), 8.13 (s, 1H), 8.06 (dd, J = 8.0, 1.3 Hz, 1H), 8.00-7.86 (m, 4H), 7.73 (t, *J* = 7.8 Hz, 1H), 7.49

(ddd, *J* = 8.3, 7.2, 1.3 Hz, 1H), 7.42 (td, *J* = 7.6, 1.2 Hz, 1H), 7.34 (s, 1H), 3.99 (s, 3H), 3.89 (s, 2H). $^{13}$C NMR (101 MHz, DMSO-$d_6$) δ 171.5, 169.7, 153.2, 148.4, 133.9, 133.9, 132.7, 131.9, 131.7, 131.2, 130.2, 128.4, 126.9, 126.5, 125.6, 122.7, 122.1, 118.6, 113.9, 112.0, 39.2, 35.3. HRMS (ESI) $C_{22}H_{17}N_4O_2S$ $[M+H]^+$ m/z found 401.1068 cald 401.1072

**(*E*)-3-(benzo[*d*]thiazol-2-yl)-4-(1-methyl-3-(3-(trifluoromethyl)phenyl)-1*H*-pyrazol-4-yl)but-3-enoic acid (33)**

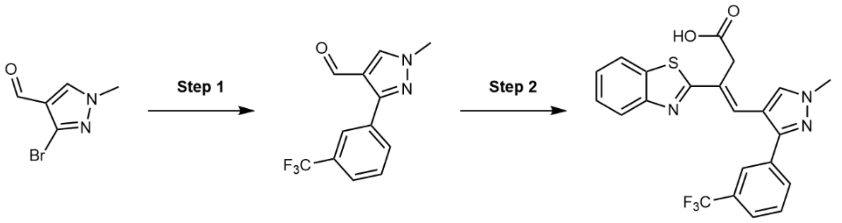


**Step 1**

According to general method D, [1,1′-bis(diphenylphosphino)ferrocene]dichloropalladium(II)·DCM (51.9 mg, 63.5 μmol) was added to 3-bromo-1-methyl-1*H*-pyrazole-4-carbaldehyde (178 mg, 0.940 mmol), 3-trifluoromethylphenylboronic acid (186.0 mg, 1.22 mmol) and 2 M sodium carbonate (480 μL, 0.952 mmol) in DME (1.85 mL). The crude product was purified using flash column chromatography eluting with a gradient of 0-40% EtOAc in n-Hex. 4-(4-formyl-1-methyl-1*H*-pyrazol-3-yl)benzonitrile (120 mg, 0.472 mmol, 50%) was obtained as a colorless solid. $^1$H NMR (400 MHz, DMSO-$d_6$) δ 9.88 (s, 1H), 8.61 (s, 1H), 8.26 (s, 1H), 8.20 (d, *J* = 7.8 Hz, 1H), 7.80 (d, *J* = 7.8 Hz, 1H), 7.71 (t, *J* = 7.8 Hz, 1H), 3.99 (s, 3H). $^{19}$F NMR (377 MHz, DMSO-$d_6$) δ -61.1. $^{13}$C NMR (101 MHz, DMSO-$d_6$) δ 184.6, 149.7, 140.2, 133.1, 132.6, 130.0, 129.8, 129.5 – 128.7 (m), 125.7, 125.0, 120.9, 39.6. LCMS: $[M-H]^+$ = 255.0.

**Step 2**

According to general method B, 3-(benzo[*d*]thiazol-2-yl)propanoic acid (54.2 mg, 0.256 mmol) and 3-(3,4-dichlorophenyl)-1-methyl-1*H*-pyrazole-4-carbaldehyde (70.0 mg, 0.270 mmol) were dissolved in anhydrous DMF (0.9 mL) and sealed in a microwave vial. TMSCl (210 μL, 1.6 mmol) was added dropwise before heating the reaction to 135 °C for 48 h. (*E*)-3-(benzo[*d*]thiazol-2-yl)-4-(1-methyl-3-(3-(trifluoromethyl)phenyl)-1*H*-pyrazol-4-yl)but-3-enoic acid (79.3 mg, 0.179 mmol, 70%) was obtained as a brown solid. $^1$H NMR (400 MHz, DMSO-$d_6$) δ 8.16 (s, 1H), 8.06 (d, *J* = 8.0 Hz, 1H), 8.00 – 7.92 (m, 1H), 7.91 – 7.84 (m, 2H), 7.84 – 7.72 (m, 2H), 7.59 – 7.46 (m, 1H), 7.43 (t, *J* = 7.4 Hz, 1H), 7.38 (s, 1H), 4.00 (s, 3H), 3.90 (s, 2H).$^{19}$F NMR (377 MHz, DMSO-$d_6$) δ -61.2. $^{13}$C NMR (151 MHz, DMSO-$d_6$) δ 171.6, 169.7, 153.2, 148.8, 133.9, 133.7, 131.9, 131.8, 130.0, 129.6 (q, *J* = 31.7 Hz), 128.2, 127.0, 126.5, 125.5, 124.6 (d, *J* = 3.8 Hz), 124.2 (d, *J* = 3.9 Hz), 124.1 (q, *J* = 272.5, 271.9 Hz), 122.7, 122.0, 113.8, 39.2, 35.3. HRMS (ESI) $C_{22}H_{17}N_3O_2F_3S$ $[M+H]^+$ m/z found 444.1001 cald 444.0994.

**(*E*)-3-(benzo[*d*]thiazol-2-yl)-4-(3-(2-fluorophenyl)-1-methyl-1*H*-pyrazol-4-yl)but-3-enoic acid (34)**

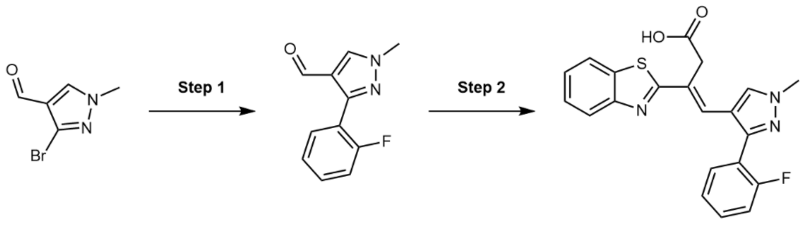

**Step 1**

According to general method D, [1,1′-bis(diphenylphosphino)ferrocene]dichloropalladium(II)·DCM (51.9 mg, 63.5 μmol) was added to 3-bromo-1-methyl-1*H*-pyrazole-4-carbaldehyde (178 mg, 0.940 mmol), 2-fluorophenylboronic acid (224 mg, 1.1 mmol) and 2 M sodium carbonate (480 μL, 0.952 mmol) in DME (1.85 mL). The crude product was purified using flash column chromatography eluting with a gradient of 0-70% EtOAc in n-Hex. 3-(2-fluorophenyl)-1-methyl-1*H*-pyrazole-4-carbaldehyde (80.0 mg, 0.392 mmol, 42%) was obtained as a colorless solid. $^{1}$H NMR (400 MHz, DMSO-$d_6$) δ 9.71 (d, *J* = 2.3 Hz, 1H), 8.52 (s, 1H), 7.59 – 7.47 (m, 2H), 7.37 – 7.26 (m, 2H), 3.96 (s, 3H). $^{13}$C NMR (101 MHz, DMSO-$d_6$) δ 184.3, 159.5 (d, *J* = 247.0 Hz), 146.3, 136.1, 131.4 (d, *J* = 3.0 Hz), 131.0 (d, *J* = 8.2 Hz), 124.51 (d, *J* = 3.5 Hz), 121.2, 119.9 (d, *J* = 14.4 Hz), 115.8 (d, *J* = 22.0 Hz), 39.2. $^{19}$F NMR (377 MHz, DMSO $d_6$) δ -115.43 – -115.87. LCMS: $[M-H]^+$ = 205.2.

**Step 2**

According to general method B, 3-(benzo[*d*]thiazol-2-yl)propanoic acid (92.2 mg, 0.445 mmol) and 3-(2-fluorophenyl)-1-methyl-1*H*-pyrazole-4-carbaldehyde (95.6 mg, 0.468 mmol) were dissolved in anhydrous DMF (0.9 mL) and sealed in a microwave vial. TMSCl (350 μL, 2.75 mmol) was added dropwise before heating the reaction to 135 °C for 48 h. (*E*)-3-(benzo[*d*]thiazol-2-yl)-4-(1-methyl-3-(quinolin-6-yl)-1*H*-pyrazol-4-yl)but-3-enoic acid (88.0 mg, 0.224 mmol, 50%) was obtained as a brown solid. $^{1}$H NMR (400 MHz, DMSO-$d_6$) δ 8.20 (s, 1H), 8.02 (d, *J* = 7.7 Hz, 1H), 7.92 (d, *J* = 8.1 Hz, 1H), 7.52 – 7.45 (m, 3H), 7.41 – 7.34 (m, 3H), 7.14 (s, 1H), 3.98 (d, *J* = 3.4 Hz, 3H), 3.90 (s, 2H). $^{13}$C NMR (101 MHz, DMSO-$d_6$) δ 171.4, 170.0, 159.3 (d, *J* = 246.7 Hz), 153.5, 146.4, 133.7, 132.0 (d, *J* = 3.5 Hz), 131.01– 130.8 (m), 127.0, 126.5 (d, *J* = 7.4 Hz), 126.0, 125.4, 124.9 – 124.8 (m), 122.5, 122.2, 122.0, 120.2, 120.1, 116.07 (d, *J* = 21.7 Hz), 115.0, 39.1. $^{19}$F NMR (377 MHz, DMSO-$d_6$) δ -114.9. HRMS (ESI) $C_{21}H_{14}ClN_3O_2S$ $[M+H]^+$ m/z found 394.1020 cald 394.1026.

**(*E*)-3-(benzo[*d*]thiazol-2-yl)-4-(1-methyl-3-(*o*-tolyl)-1*H*-pyrazol-4-yl)but-3-enoic acid (35)**

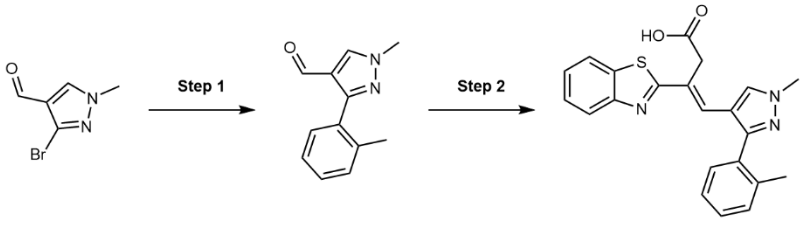

**Step 1**

According to general method D, [1,1′-bis(diphenylphosphino)ferrocene]dichloropalladium(II)·DCM (51.9 mg, 63.5 μmol) was added to 3-bromo-1-methyl-1*H*-pyrazole-4-carbaldehyde (120 mg, 0.635 mmol), *o*-tolylboronic acid (173 mg, 1.27 mmol) and 2 M sodium carbonate (480 µL, 0.952 mmol) in DME (1.85 mL). The crude product was purified using flash column chromatography eluting with a gradient of 0-50% EtOAc in n-Hex. 1-methyl-3-(*o*-tolyl)-1*H*-pyrazole-4-carbaldehyde (72.4 mg, 0.362 mmol, 57%) was obtained as a yellow oil. $^{1}$H NMR (400 MHz, $CDCl_3$) δ 9.60 (s, 1H), 8.01 (s, 1H), 7.39-7.18 (m, 4H), 3.99 (s, 3H), 2.30 (s, 3H). LCMS: $[M-H]^+$ = 201.1.

**Step 2**

According to general method B, 3-(1,3-benzothiazol-2-yl)propanoic acid (67.0 mg, 0.323 mmol) and 1-methyl-3-(*o*-tolyl)-1*H*-pyrazole-4-carbaldehyde (72.4 mg, 0.362 mmol) were dissolved in anhydrous DMF (0.424 mL) and sealed in a microwave vial. TMSCl (224 µL, 1.73 mmol) was added dropwise before heating the reaction to 110 °C for 9 h. (*E*)-3-(benzo[*d*]thiazol-2-yl)-4-(1-methyl-3-(*o*-tolyl)-1*H*-pyrazol-4-yl)but-3-enoic acid (31.2 mg, 80.1 µmol, 25%) was obtained as a light brown solid. $^{1}$H NMR (400 MHz, DMSO-$d_6$) δ 12.50 (s, 1H), 8.20 (s, 1H), 7.98 (dd, *J* = 7.9, 1.3 Hz, 1H), 7.91-7.87 (m, 1H), 7.45 (ddd, *J* = 8.3, 7.2, 1.4 Hz, 1H), 7.40-7.35 (m, 3H), 7.33-7.26 (m, 1H), 7.24 (d, *J* = 7.4 Hz, 1H), 6.97 (s, 1H), 3.97 (s, 3H), 3.91 (s, 2H), 2.23 (s, 3H). $^{13}$C NMR (101 MHz, DMSO-$d_6$) δ 171.4, 170.1, 153.3, 151.9, 136.8, 133.7, 131.8, 130.7, 130.6, 130.4, 128.5, 127.4, 126.4, 125.9, 125.7, 125.3, 122.5, 122.0, 39.0, 35.3, 32.3, 19.9. HRMS (ESI) $C_{22}H_{20}N_3O_2S$ $[M+H]^+$ m/z found 390.1270 cald 390.1276.

**(*E*)-3-(benzo[*d*]thiazol-2-yl)-4-(3-(3,4-dichlorophenyl)-1-methyl-1*H*-pyrazol-4-yl)but-3-enoic acid (36)**

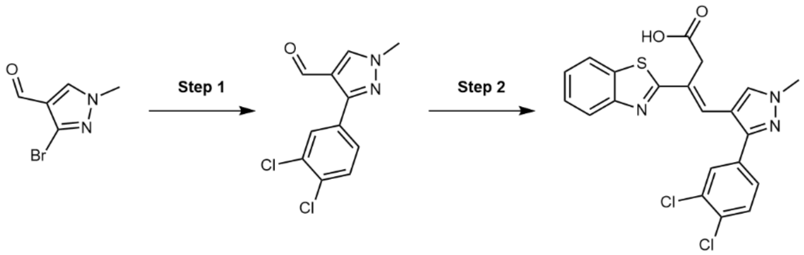


**Step 1**

According to general method D, [1,1′-bis(diphenylphosphino)ferrocene]dichloropalladium(II)·DCM (51.9 mg, 63.5 µmol) was added to 3-bromo-1-methyl-1*H*-pyrazole-4-carbaldehyde (120 mg, 0.635 mmol), 3,4-dichlorophenylboronic acid (242 mg, 1.27 mmol) and 2 M sodium carbonate (480 µL, 0.952 mmol) in DME (1.85 mL). The crude product was purified using flash column chromatography eluting with a gradient of 0-40% EtOAc in n-Hex. 3-(3,4-dichlorophenyl)-1-methyl-1*H*-pyrazole-4-carbaldehyde (79.3 mg, 0.311 mmol, 49%) was obtained as a colorless solid. $^{1}$H NMR (400 MHz, DMSO-$d_6$) δ 9.85 (s, 1H), 8.58 (s, 1H), 8.17 (d, *J* = 2.1 Hz, 1H), 7.89 (dd, *J* = 8.4, 2.1 Hz, 1H), 7.71 (d, *J* = 8.4 Hz, 1H), 3.96 (s, 3H). $^{13}$C NMR (101 MHz, DMSO-$d_6$) δ 184.2, 148.3, 139.8, 132.3, 131.3, 131.2, 130.7, 129.8, 128.3, 120.5, 39.2. LCMS: $[M-H]^+$ = 256.0.

**Step 2**

According to general method B, 3-(benzo[*d*]thiazol-2-yl)propanoic acid (50.9 mg, 0.246 mmol) and 3-(3,4-dichlorophenyl)-1-methyl-1*H*-pyrazole-4-carbaldehyde (66.0 mg, 0.258 mmol) were dissolved in anhydrous DMF (0.9 mL) and sealed in a microwave vial. TMSCl (190 µL, 1.6 mmol) was added dropwise before heating the reaction to 135 °C for 48 h. (*E*)-3-(benzo[*d*]thiazol-2-yl)-4-(3-(3,4-dichlorophenyl)-1-methyl-1*H*-pyrazol-4-

yl)but-3-enoic acid (64.0 mg, 0.144 mmol, 56%) was obtained as an off-white solid. $^{1}H$ NMR (400 MHz, DMSO-$d_6$) δ 12.51 (s, 1H), 8.58 (s, 1H), 8.11 (s, 1H), 8.04 (d, *J* = 1.2 Hz, 1H), 7.94 (s, 1H), 7.76 (s, 1H), 7.56 (d, *J* = 2.1 Hz, 1H), 7.49 (d, *J* = 1.1 Hz, 1H), 7.42 (s, 1H), 7.34 (s, 1H), 3.96 (s, 3H), 3.90 (d, *J* = 1.7 Hz, 2H). $^{13}C$ NMR (101 MHz, DMSO-$d_6$) δ 171.5, 169.7, 153.2, 147.9, 133.9, 133.3, 131.9, 131.6, 131.1, 130.7, 129.4, 128.3, 128.0, 126.9, 126.5, 125.6, 122.7, 122.1, 113.8, 39.2, 35.2. HRMS (ESI) $C_{21}H_{16}N_3O_2Cl_2$ $[M+H]^+$ m/z found 444.0359 cald 444.0340.

**(*E*)-3-(benzo[*d*]thiazol-2-yl)-4-(3-(4-chloro-3-fluorophenyl)-1-methyl-1*H*-pyrazol-4-yl)but-3-enoic acid (37)**

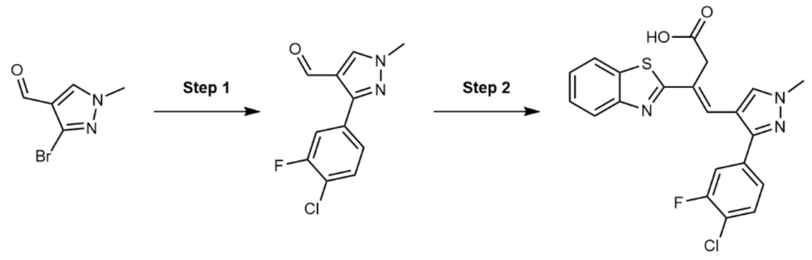


**Step 1**

According to general method D, [1,1′-bis(diphenylphosphino)ferrocene]dichloropalladium(II)·DCM (51.9 mg, 63.5 μmol) was added to 3-bromo-1-methyl-1H-pyrazole-4-carbaldehyde (178 mg, 0.940 mmol), (4-chloro-3-fluorophenyl)boronic acid (279.1 mg, 1.6 mmol), and 2 M sodium carbonate (480 μL, 0.952 mmol) in DME (1.85 mL) . The crude product was purified using flash column chromatography eluting with a gradient of 0-50% EtOAc in n-Hex. 3-(4-chloro-3-fluorophenyl)-1-methyl-1*H*-pyrazole-4-carbaldehyde (143 mg, 0.602 mmol, 64%) was obtained as a colorless solid. $^{1}H$ NMR (400 MHz, DMSO-$d_6$) δ 9.85 (s, 1H), 8.59 (s, 1H), 7.97 (dd, *J* = 10.9, 2.0 Hz, 1H), 7.78 (dd, *J* = 8.4, 2.0 Hz, 1H), 7.67 (t, *J* = 8.1 Hz, 1H), 3.96 (s, 3H). $^{13}C$ NMR (101 MHz, DMSO-$d_6$) δ 184.2, 152.2 (d, *J* = 729.0 Hz), 148.6, 139.7, 132.8, 132.7, 130.7, 129.95, 125.3, 120.5, 116.3 (d, *J* = 22.8 Hz), 39.2. $^{19}F$ NMR (377 MHz, DMSO-$d_6$) δ -116.03 – -116.16 (m). LCMS: $[M-H]^+$ = 239.0.

**Step 2**

According to general method B, 3-(benzo[*d*]thiazol-2-yl)propanoic acid (92.2 mg, 0.445 mmol) and 3-(2-fluorophenyl)-1-methyl-1*H*-pyrazole-4-carbaldehyde (110 mg, 0.461 mmol) were dissolved in anhydrous DMF (0.9 mL) and sealed in a microwave vial. TMSCl (350 μL, 2.75 mmol) was added dropwise before heating the reaction to 135 °C for 48 h. (*E*)-3-(benzo[*d*]thiazol-2-yl)-4-(1-methyl-3-(quinolin-6-yl)-1*H*-pyrazol-4-yl)but-3-enoic acid (87.0 mg, 0.203 mmol, 45%) was obtained as a brown solid. $^{1}H$ NMR (400 MHz, DMSO-$d_6$) δ 12.54 (s, 1H), 8.12 (s, 1H), 8.06 (d, *J* = 7.6 Hz, 1H), 7.95 (d, *J* = 8.0 Hz, 1H), 7.74 (t, *J* = 8.1 Hz, 1H), 7.57 (dd, *J* = 10.4, 1.9 Hz, 1H), 7.49 (d, *J* = 7.6 Hz, 1H), 7.44 (s, 1H), 7.41 (d, *J* = 6.9 Hz, 1H), 7.35 (s, 1H), 3.97 (d, *J* = 4.3 Hz, 3H), 3.88 (s, 2H). $^{19}F$ NMR (377 MHz, DMSO-$d_6$) δ -115.48 – -115.61 (m). HRMS (ESI) $C_{21}H_{15}ClFN_3O_2S$ $[M+H]^+$ m/z found 428.0636. cald 428.0645.

**(*E*)-3-(benzo[*d*]thiazol-2-yl)-4-(1-methyl-3-(pyridine-4-yl)-1*H*-pyrazol-4-yl)but-3-enoic acid (38)**

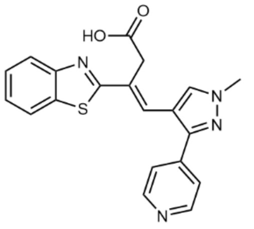

According to general method B, 3-(1,3-benzothiazol-2-yl)propanoic acid (101 mg, 0.489 mmol) and 1-methyl-3-(pyridine-4-yl)-1*H*-pyrazole-4-carbaldehyde (100 mg, 0.508 mmol) were dissolved in anhydrous DMF (0.625 mL) and sealed in a microwave vial. TMSCl (329 µL, 2.54 mmol) was added dropwise before heating the reaction to 110 °C for 9 h. (*E*)-3-(benzo[*d*]thiazol-2-yl)-4-(1-methyl-3-(pyridine-4-yl)-1*H*-pyrazol-4-yl)but-3-enoic acid (34.5 mg, 91.7 µmol, 19%) was obtained as a yellow solid. $^1$H NMR (400 MHz, DMSO-$d_6$) δ 12.56 (s, 1H), 8.91-8.85 (m, 2H), 8.17 (s, 1H), 8.11-8.07 (m, 1H), 8.06-8.02 (m, 2H), 8.01-7.96 (m, 1H), 7.54-7.49 (m, 2H), 7.45 (td, *J* = 7.6, 1.3 Hz, 1H), 4.04 (s, 3H), 3.87 (s, 2H). $^{13}$C NMR (101 MHz, DMSO-$d_6$) δ 171.5, 169.3, 153.2, 146.4, 145.1, 144.2, 134.1, 133.0, 130.4, 126.6, 126.3, 125.8, 123.5, 122.8, 122.1, 116.0, 39.6, 35.3. HRMS (ESI) $C_{20}H_{17}N_4O_2S$ $[M+H]^+$ m/z found 377.1062 cald 377.1072.

**(*E*)-3-(benzo[*d*]thiazol-2-yl)-4-(1-methyl-3-(pyridine-3-yl)-1*H*-pyrazol-4-yl)but-3-enoic acid (39)**

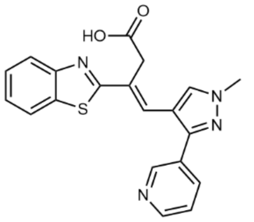

3-(1,3-benzothiazol-2-yl)propanoic acid (101 mg, 0.489 mmol) and 1-methyl-3-(pyridine-3-yl)-1*H*-pyrazole-4-carbaldehyde (100 mg, 0.508 mmol) were dissolved in anhydrous DMF (0.625 mL) and sealed in a microwave vial. TMSCl (329 µL, 2.54 mmol) was added dropwise before heating the reaction to 110 °C for 9 h. The reaction was quenched by pouring into water. The crude mixture was concentrated in vacuo and purified using reverse phase flash column chromatography eluting with a gradient of 5-100% ACN in $H_2O$. Pure fractions were combined and the organic solvent removed in vacuo, resulting in precipitation of product out of the aqueous phase. The precipitated product was collected by vacuum filtration, washed with cold water and dried *in vacuo*. (*E*)-3-(benzo[*d*]thiazol-2-yl)-4-(1-methyl-3-(pyridine-3-yl)-1*H*-pyrazol-4-yl)but-3-enoic acid (45.3 mg, 0.120 mmol, 25%) was obtained as a yellow solid. $^1$H NMR (400 MHz, DMSO-$d_6$) δ 12.54 (s, 1H), 8.78 (d, J = 2.2 Hz, 1H), 8.63 (dd, *J* = 4.8, 1.6 Hz, 1H), 8.16 (s, 1H), 8.04 (d, *J* = 7.5 Hz, 1H), 8.00-7.92 (m, 2H), 7.54 (dd, *J* = 8.0, 4.8 Hz, 1H), 7.49 (ddd, *J* = 8.2, 7.2, 1.4 Hz, 1H), 7.41 (td, *J* = 7.6, 1.2 Hz, 1H), 7.35 (s, 1H), 3.99 (s, 3H), 3.90 (s, 2H). $^{13}$C NMR (101 MHz, DMSO-$d_6$) δ 171.5, 169.8, 153.2, 149.1, 148.6, 147.7, 135.4, 133.8, 131.8, 128.6, 128.0, 126.9, 126.5, 125.5, 123.9, 122.6, 122.1, 114.0, 39.2, 35.3. HRMS (ESI) $C_{20}H_{17}N_4O_2S$ $[M+H]^+$ m/z found 377.1073 cald 377.1072

**(*E*)-3-(benzo[*d*]thiazol-2-yl)-4-(3-(6-chloropyridin-3-yl)-1-methyl-1*H*-pyrazol-4-yl)but-3-enoic acid (40)**

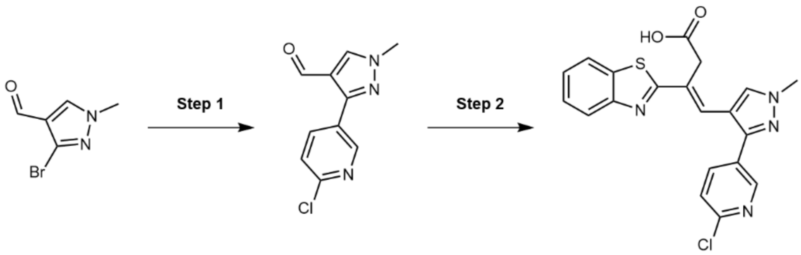

**Step 1**

According to general method D, [1,1′-bis(diphenylphosphino)ferrocene]dichloropalladium(II)·DCM (51.9 mg, 63.5 µmol) was added to 3-bromo-1-methyl-1*H*-pyrazole-4-carbaldehyde (120 mg, 0.635 mmol), (6-chloropyridin-3-yl)boronic acid (200 mg, 1.27 mmol) and 2 M sodium carbonate (480 µL, 0.952 mmol) in DME (1.85 mL) . The crude product was purified using flash column chromatography eluting with a gradient of 0-60% EtOAc in n-Hex. 3-(6-chloropyridin-3-yl)-1-methyl-1*H*-pyrazole-4-carbaldehyde (97.1 mg, 0.438 mmol, 69%) was obtained as a colorless solid. $^{1}$H NMR (400 MHz, DMSO-$d_6$) δ 9.87 (s, 1H), 8.87 (d, *J* = 2.5 Hz, 1H), 8.63 (s, 1H), 8.32 (dd, *J* = 8.4, 2.5 Hz, 1H), 7.64 (d, *J* = 8.4 Hz, 1H), 3.99 (s, 3H). $^{13}$C NMR (101 MHz, DMSO-$d_6$) δ 184.7, 150.7, 149.4, 147.2, 140.1, 139.6, 127.7, 121.2. (Methyl environment obscured by solvent residual peak). LCMS: $[M-H]^+$ = 222.6.

**Step 2**

According to general method B, 3-(benzo[*d*]thiazol-2-yl)propanoic acid (23.7 mg, 0.114 mmol) and 3-(6-chloropyridin-3-yl)-1-methyl-1*H*-pyrazole-4-carbaldehyde (26.7 mg, 0.120 mmol) were dissolved in anhydrous DMF (0.9 mL) and sealed in a microwave vial. TMSCl (160 µL, 1.3 mmol) was added dropwise before heating the reaction to 135 °C for 48 h. (*E*)-3-(benzo[*d*]thiazol-2-yl)-4-(3-(6-chloropyridin-3-yl)-1-methyl-1*H*-pyrazol-4-yl)but-3-enoic acid (20.0 mg, 48.6 µmol, 40%) was obtained as a brown solid. $^{1}$H NMR (400 MHz, DMSO-$d_6$) δ 12.52 (s, 1H), 8.60 (d, *J* = 2.5 Hz, 1H), 8.16 (s, 1H), 8.11 – 7.97 (m, 2H), 7.95 (d, *J* = 8.1 Hz, 1H), 7.68 (d, *J* = 8.3 Hz, 1H), 7.54 – 7.45 (m, 1H), 7.35 (s, 1H), 3.99 (d, *J* = 4.8 Hz, 3H), 3.89 (s, 2H). $^{13}$C NMR (101 MHz, DMSO-$d_6$) δ 171.5, 169.7, 153.2, 149.6, 148.5, 146.3, 138.8, 133.9, 131.9, 128.5, 128.1, 126.6, 125.5, 124.6, 123.6, 122.6, 122.1, 114.2, 39.2, 35.2. HRMS (ESI) $C_{20}H_{15}ClN_4O_2S$ $[M+H]^+$ m/z found 411.0683 cald 411.0683.

**(*E*)-3-(benzo[*d*]thiazol-2-yl)-4-(1-methyl-3-(quinolin-6-yl)-1*H*-pyrazol-4-yl)but-3-enoic acid (41)**

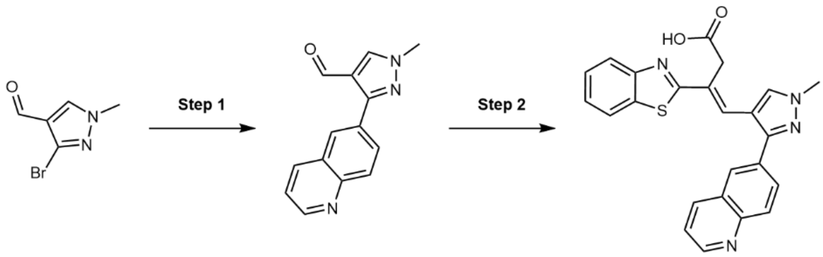

**Step 1**

According to general method D, [1,1′-bis(diphenylphosphino)ferrocene]dichloropalladium(II)·DCM (51.9 mg, 63.5 µmol) was added to 3-bromo-1-methyl-1*H*-pyrazole-4-carbaldehyde (178 mg, 0.940 mmol), quinolin-6-

ylboronic acid (280 mg, 1.62 mmol). and 2 M sodium carbonate (480 µL, 0.952 mmol) in DME (1.85 mL). The crude product was purified using flash column chromatography eluting with a gradient of 0-70% EtOAc in n-Hex. 1-methyl-3-(quinolin-6-yl)-1*H*-pyrazole-4-carbaldehyde (156 mg, 0.658 mmol, 70%) was obtained as a colorless solid. $^{1}$H NMR (400 MHz, $CDCl_3$) δ 10.02 (s, 1H), 8.95 (dd, *J* = 4.3, 1.7 Hz, 1H), 8.28 (d, *J* = 1.9 Hz, 1H), 8.24 (dd, *J* = 8.2, 1.7 Hz, 1H), 8.19 (d, *J* = 8.8 Hz, 1H), 8.13 (dd, *J* = 8.8, 1.9 Hz, 1H), 8.05 (s, 1H), 7.45 (dd, *J* = 8.3, 4.2 Hz, 1H), 4.04 (s, 3H). $^{13}$C NMR (101 MHz, $CDCl_3$) δ 184.5, 153.1, 151.2, 150.8, 148.5, 136.7, 135.7, 130.34, 130.0, 129.3, 128.2, 126.2, 121.6, 39.8. LCMS: $[M-H]^+$ = 238.2.

**Step 2**

According to general method B, 3-(benzo[*d*]thiazol-2-yl)propanoic acid (23.4 mg, 0.113 mmol) and 1-methyl-3-(quinolin-6-yl)-1*H*-pyrazole-4-carbaldehyde (33.0 mg, 0.139 mmol) were dissolved in anhydrous DMF (0.9 mL) and sealed in a microwave vial. TMSCl (150 µL, 2.26 mmol) was added dropwise before heating the reaction to 135 °C for 48 h. (*E*)-3-(benzo[*d*]thiazol-2-yl)-4-(1-methyl-3-(quinolin-6-yl)-1*H*-pyrazol-4-yl)but-3-enoic acid (37.2 mg, 87.1 µmol, 77%) was obtained as a grey solid. $^{1}$H NMR (400 MHz, DMSO-*d*$_6$) δ 8.07 (s, 1H), 8.04 (d, *J* = 5.1 Hz, 1H), 7.94 (d, *J* = 7.7 Hz, 1H), 7.89 (d, *J* = 19.2 Hz, 1H), 7.70 (dd, *J* = 4.9, 3.0 Hz, 1H), 7.64 (s, 1H), 7.48 (s, 1H), 7.40 (d, *J* = 6.0 Hz, 1H), 7.39 (s, 1H), 3.94 (d, *J* = 3.6 Hz, 3H), 3.86 (s, 2H). $^{13}$C NMR (101 MHz, DMSO-*d*$_6$) δ 173.1, 171.6, 170.0, 169.0, 153.3, 146.8, 133.9, 133.6, 131.2, 129.1, 127.4, 127.0, 126.4, 125.4, 123.4, 122.6, 122.0, 113.6, 39.0. (Methyl environment obscured by solvent residual peak). HRMS (ESI) $C_{24}H_{19}N_4O_2S$ $[M+H]^+$ m/z found 427.1222 cald 427.1229.

**(E)-3-(benzo[*d*]thiazol-2-yl)-4-(3-cyclohexyl-1-methyl-1*H*-pyrazol-4-yl)but-3-enoic acid (42)**

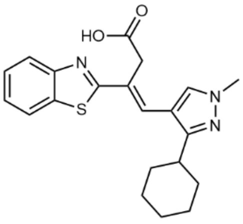


According to general method B, 3-(1,3-benzothiazol-2-yl)propanoic acid (101 mg, 0.488 mmol) and 3-cyclohexyl-1-methyl-1*H*-pyrazole-4-carbaldehyde (100 mg, 0.494 mmol) were dissolved in anhydrous DMF (0.625 mL) and sealed in a microwave vial. TMSCl (320 µL, 2.47 mmol) was added dropwise before heating the reaction to 110 °C for 9 h. (*E*)-3-(benzo[*d*]thiazol-2-yl)-4-(3-cyclohexyl-1-methyl-1*H*-pyrazol-4-yl)but-3-enoic acid (32 mg, 83.4 µmol, 17%) was obtained as a yellow solid. $^{1}$H NMR (400 MHz, DMSO-*d*$_6$) δ 12.48 (s, 1H), 8.05 (d, J = 7.5 Hz, 1H), 7.96-7.88 (m, 2H), 7.48 (ddd, *J* = 8.2, 7.2, 1.3 Hz, 1H), 7.40 (td, *J* = 7.6, 1.3 Hz, 1H), 7.36 (s, 1H), 3.84 (s, 5H), 2.85-2.70 (m, 1H), 1.87-1.75 (m, 4H), 1.74-1.67 (m, 1H), 1.58-1.32 (m, 4H), 1.24 (tt, *J* = 12.3, 3.4 Hz, 1H). $^{13}$C NMR (101 MHz, DMSO-*d*$_6$) δ 171.5, 170.3, 156.6, 153.3, 133.8, 130.4, 126.8, 126.4, 125.4, 125.3, 122.4, 122.0, 112.7, 38.7, 35.6, 35.4, 32.7, 26.1, 25.8. HRMS (ESI) $C_{21}H_{24}N_3O_2S$ $[M+H]^+$ m/z found 382.1589 cald 382.1589.

**(*E*)-3-(benzo[*d*]thiazol-2-yl)-4-(1-methyl-3-(thiophen-2-yl)-1*H*-pyrazol-4-yl)but-3-enoic acid (43)**

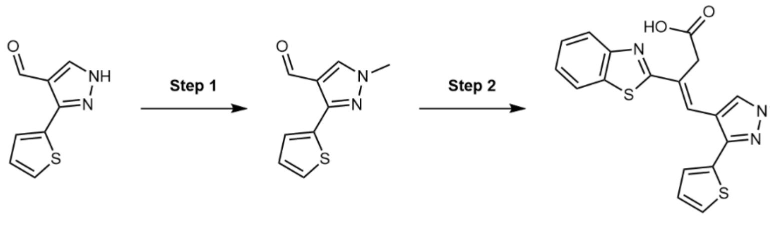


**Step 1**

According to general method A, 3-(thiophen-2-yl)-1*H*-pyrazole-4-carbaldehyde (300 mg, 1.68 mmol) was reacted with cesium carbonate (1.14 g, 3.37 mmol) and iodomethane (165 µL, 2.53 mmol) in anhydrous DMF (13 mL). The crude product was purified using flash column chromatography eluting with a gradient of 0-50% EtOAc in n-Hex. 1-methyl-3-(thiophen-2-yl)-1*H*-pyrazole-4-carbaldehyde (203 mg, 1.06 mmol, 63%) was obtained as a colorless solid. $^{1}$H NMR (400 MHz, $CDCl_3$) δ 9.99 (s, 1H), 7.92 (s, 1H), 7.79 (dd, *J* = 3.7, 1.2 Hz, 1H), 7.36 (dd, *J* = 5.1, 1.2 Hz, 1H), 7.10 (dd, *J* = 5.1, 3.7 Hz, 1H), 3.92 (s, 3H). LCMS: $[M\text{-}H]^+$ = 193.0.

**Step 2**

According to general method B, 3-(1,3-benzothiazol-2-yl)propanoic acid (217 mg, 1.05 mmol) and 1-methyl-3-(thiophen-2-yl)-1*H*-pyrazole-4-carbaldehyde (203 mg, 1.06 mmol) were dissolved in anhydrous DMF (1.34 mL) and sealed in a microwave vial. TMSCl (684 µL, 5.28 mmol) was added dropwise before heating the reaction to 110 °C for 24 h. (*E*)-3-(benzo[*d*]thiazol-2-yl)-4-(1-methyl-3-(thiophen-2-yl)-1*H*-pyrazol-4-yl)but-3-enoic acid (128 mg, 0.336 mmol, 32%) was obtained as a brown solid. $^{1}$H NMR (400 MHz, DMSO-$d_6$) δ 12.52 (s, 1H), 8.10 (s, 1H), 8.07 (ddd, *J* = 7.9, 1.3, 0.6 Hz, 1H), 7.97-7.94 (m, 1H), 7.64 (dd, *J* = 5.1, 1.1 Hz, 1H), 7.53 (s, 1H), 7.50 (ddd, *J* = 8.2, 7.2, 1.3 Hz, 1H), 7.42 (ddd, *J* = 8.4, 7.3, 1.3 Hz, 1H), 7.28 (dd, *J* = 3.6, 1.1 Hz, 1H), 7.21 (dd, *J* = 5.1, 3.6 Hz, 1H), 3.95 (s, 3H), 3.89 (s, 2H). $^{13}$C NMR (101 MHz, DMSO-$d_6$) δ 171.5, 169.8, 153.3, 145.0, 134.6, 133.8, 131.7, 128.0, 127.8, 126.9, 126.5, 126.5, 125.8, 125.5, 122.6, 122.1, 113.2, 39.0, 35.3. HRMS (ESI) $C_{19}H_{16}N_3O_2S_2$ $[M+H]^+$ m/z found 382.0666 cald 382.0684.

**(*E*)-3-(benzo[*d*]thiazol-2-yl)-4-(3-(5-chlorothiophen-2-yl)-1-methyl-1*H*-pyrazol-4-yl)but-3-enoic acid (44)**

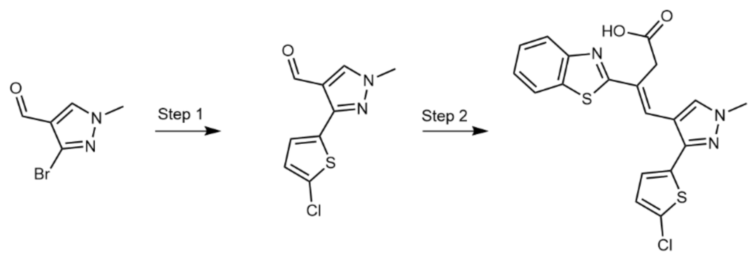


**Step 1**

According to general method D, [1,1′-bis(diphenylphosphino)ferrocene]dichloropalladium(II)·DCM (51.9 mg, 63.5 µmol) was added to 3-bromo-1-methyl-1*H*-pyrazole-4-carbaldehyde (120 mg, 0.635 mmol), (5-chlorothiophen-2-yl)boronic acid (206.2 mg, 1.27 mmol) and 2 M sodium carbonate (480 µL, 0.952 mmol) in DME (1.85 mL). The crude product was purified using flash column chromatography eluting with a gradient of

0-40% EtOAc in n-Hex. 3-(5-chlorothiophen-2-yl)-1-methyl-1*H*-pyrazole-4-carbaldehyde (61.9 mg, 0.273 mmol, 43%) was obtained as a colorless solid. $^{1}H$ NMR (400 MHz, DMSO-*d*$_6$) δ 9.89 (s, 1H), 8.59 (s, 1H), 8.00 (d, *J* = 4.0 Hz, 1H), 7.18 (d, *J* = 4.0 Hz, 1H), 3.93 (s, 3H). $^{13}C$ NMR (101 MHz, DMSO-*d*$_6$) δ 183.9, 144.2, 140.3, 133.3, 129.0, 127.8, 127.6, 119.6, 39.1. LCMS: $[M-H]^+$ = 227.7.

**Step 2**

According to general method B, 3-(benzo[*d*]thiazol-2-yl)propanoic acid (42.8 mg, 0.206 mmol) and 3-(5-chlorothiophen-2-yl)-1-methyl-1*H*-pyrazole-4-carbaldehyde (49.2 mg, 0.217 mmol) were dissolved in anhydrous DMF (0.9 mL) and sealed in a microwave vial. TMSCl (160 µL, 1.3 mmol) was added dropwise before heating the reaction to 135 °C for 48 h. (*E*)-3-(benzo[*d*]thiazol-2-yl)-4-(3-(5-chlorothiophen-2-yl)-1-methyl-1*H*-pyrazol-4-yl)but-3-enoic acid (50.0 mg, 0.120 mmol, 58%) was obtained as a brown solid. $^{1}H$ NMR (400 MHz, DMSO-*d*$_6$) δ 8.58 (s, 1H), 8.11 – 8.04 (m, 1H), 7.98 (s, 1H), 7.90 (s, 1H), 7.52 (s, 1H), 7.41 (s, 1H), 7.20 (s, 1H), 7.10 (s, 1H), 3.91 (s, 3H), 3.87 (s, 2H).$^{13}C$ NMR (101 MHz, DMSO-*d*$_6$) δ 171.5, 169.6, 153.2, 143.8, 133.9, 133.9, 132.6, 131.1, 128.5, 127.9, 126.5, 126.3, 125.6, 125.3, 122.7, 122.1, 113.3, 39.1, 35.3. HRMS (ESI) $C_{19}H_{14}ClN_3O_2S_2$ $[M+H]^+$ m/z found 416.0284 cald 416.0294.

**$^{1}H$, $^{19}F$, and $^{13}C$ NMR Spectra**

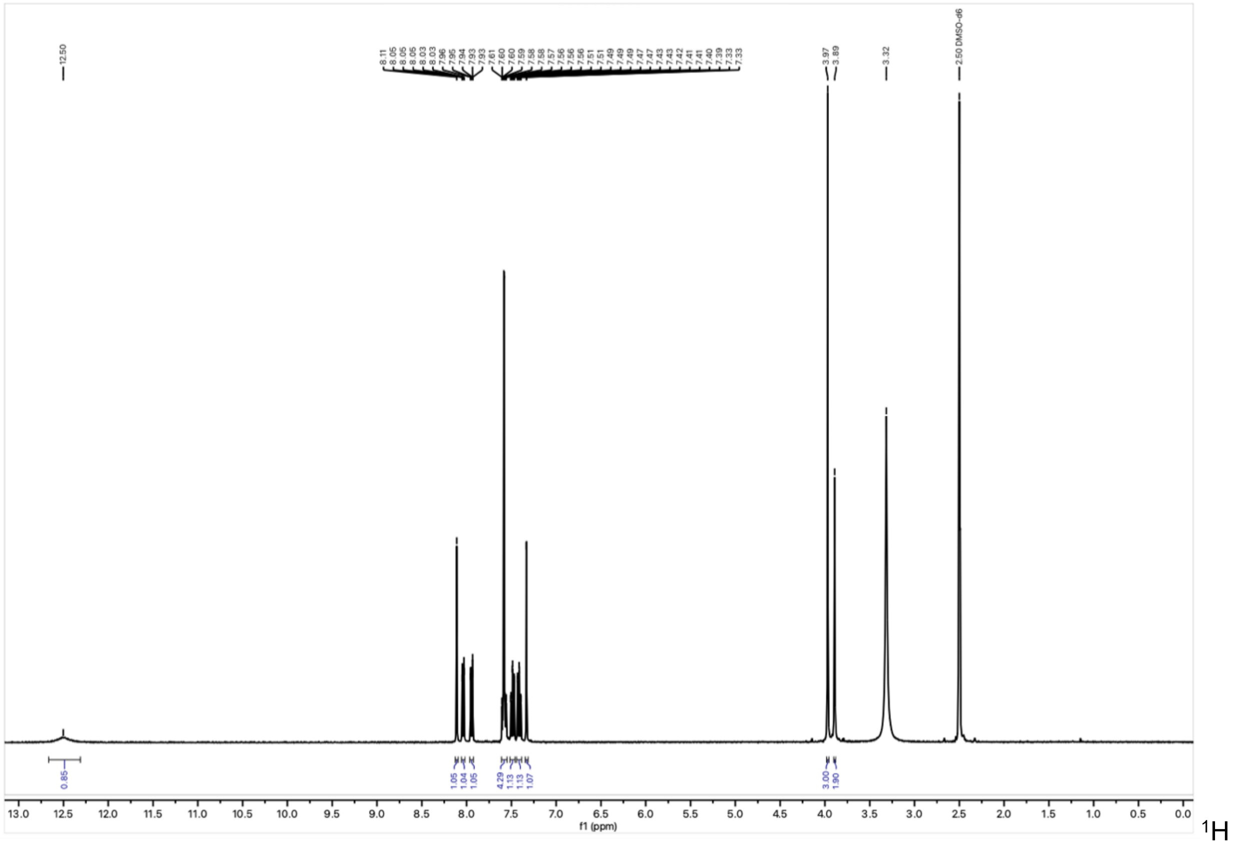

NMR spectrum of **IMP-2380** in DMSO-$d_6$ at 400 MHz.

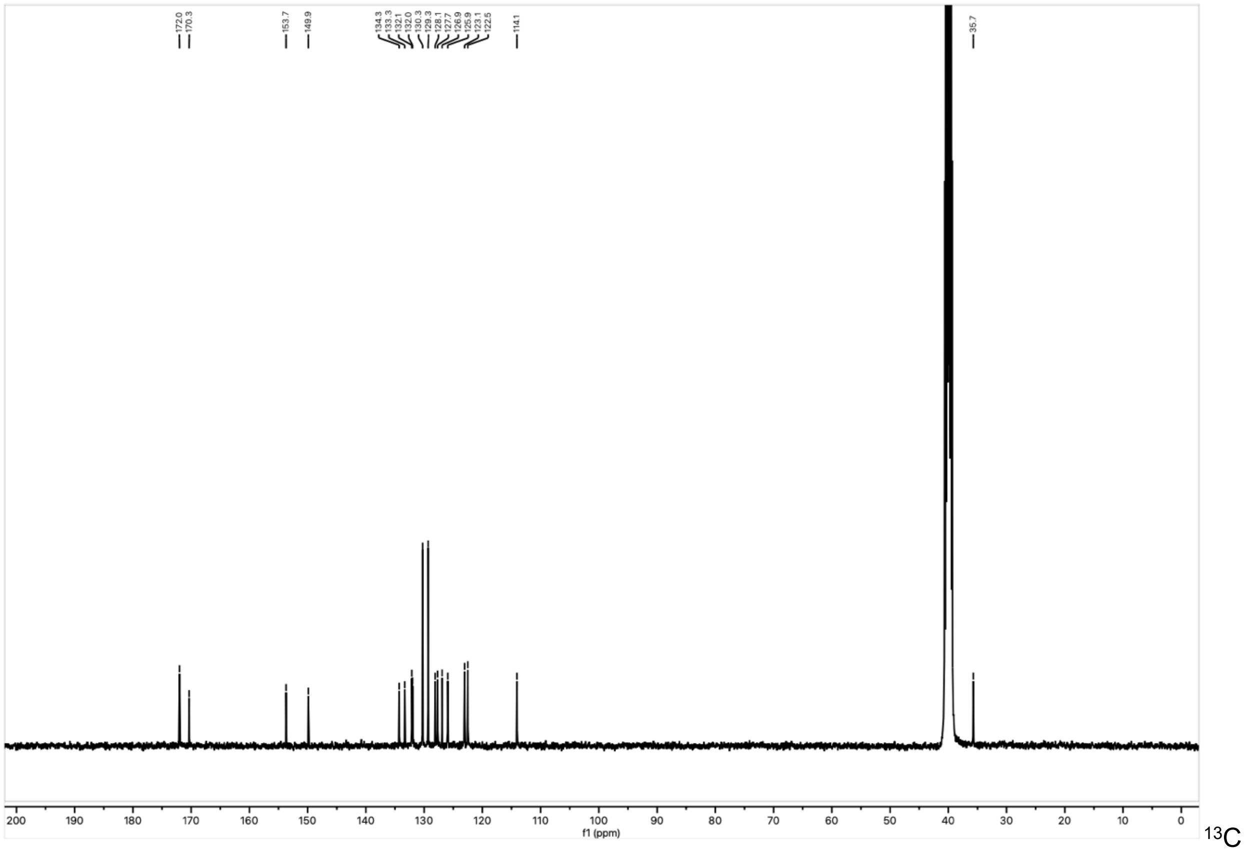

NMR spectrum of **IMP-2380** in DMSO-$d_6$ at 101 MHz.

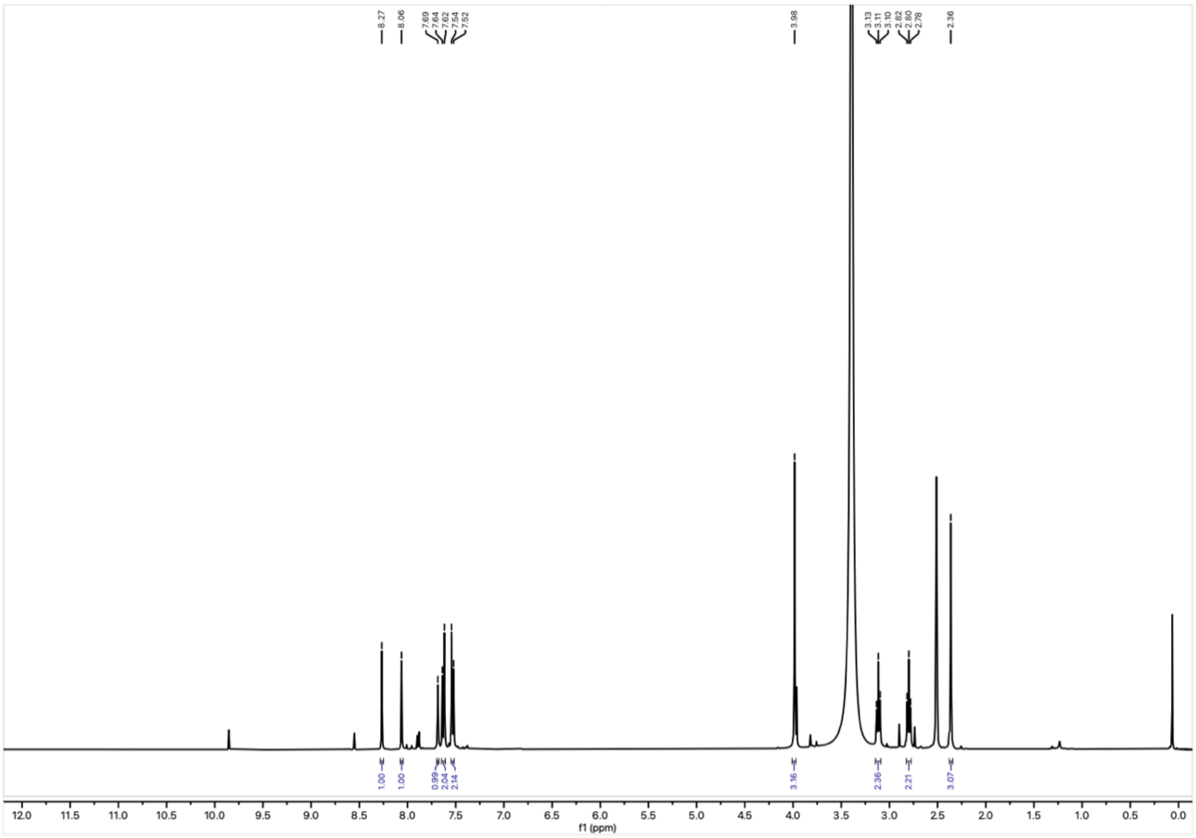

$^{1}H$ NMR spectrum of **7** in DMSO-$d_6$ at 400 MHz.

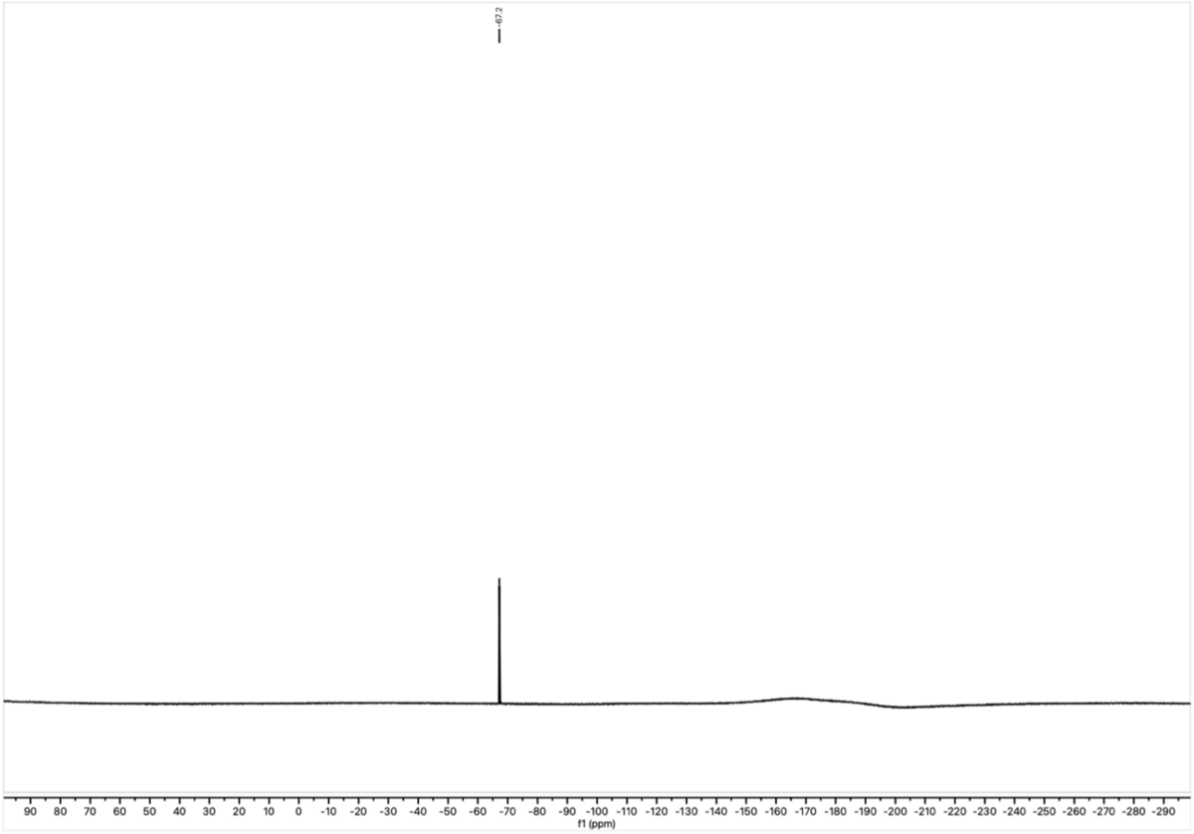

$^{19}F$ NMR spectrum of **7** in DMSO-$d_6$ at 377 MHz.

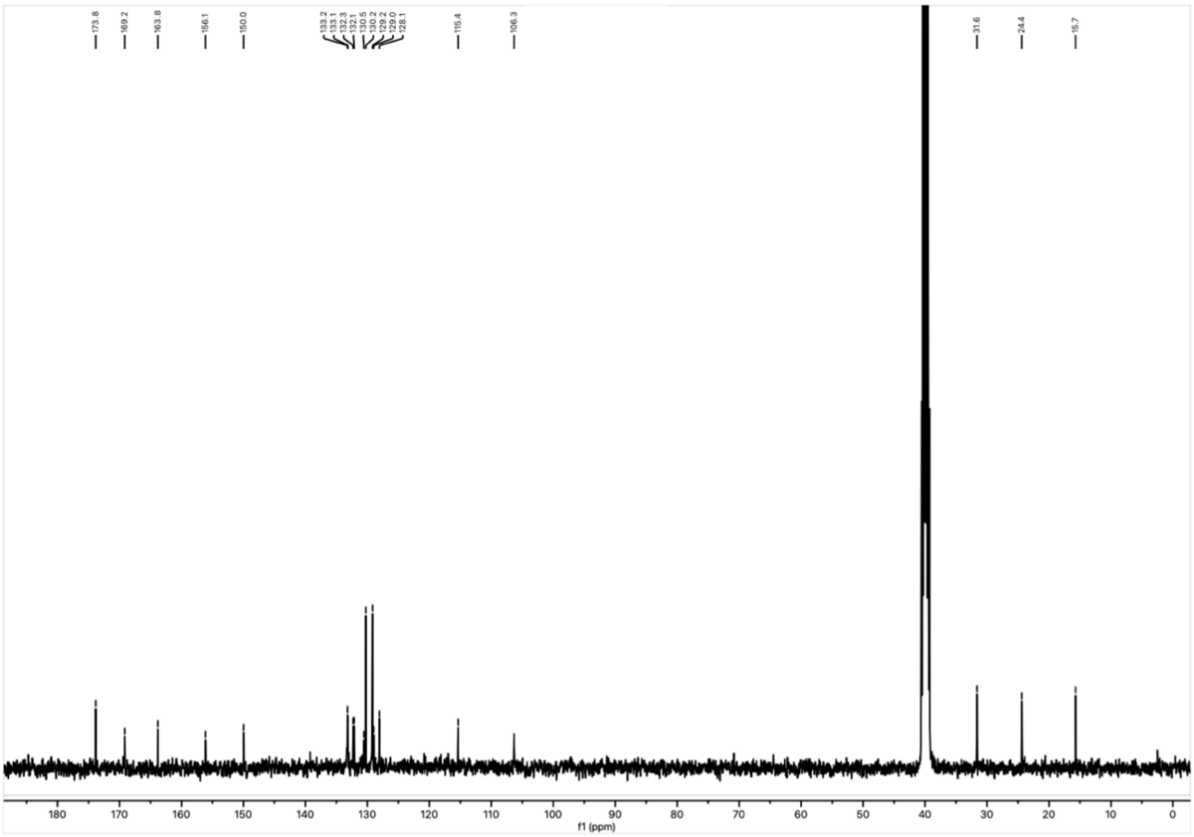


$^{13}$C NMR spectrum of **7** in DMSO-$d_6$ at 101 MHz.

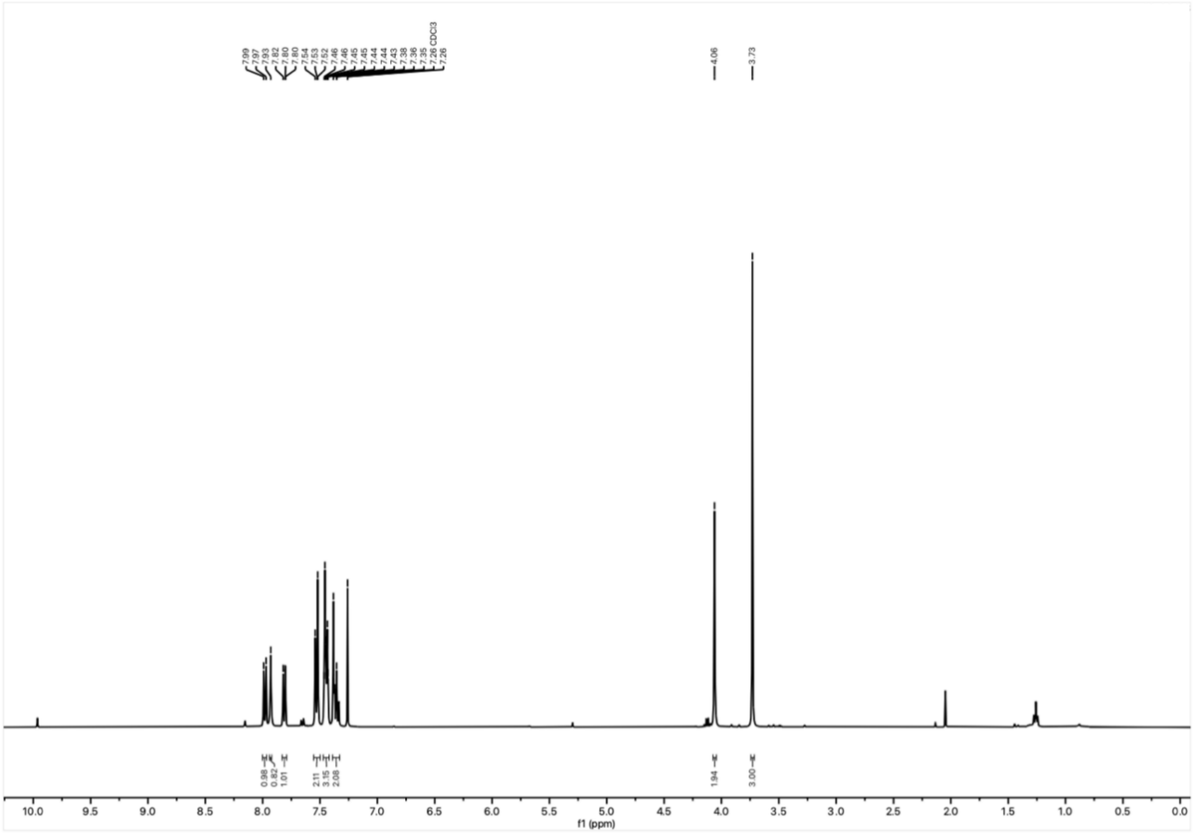


$^1$H NMR spectrum of **11** in Chloroform-*d* at 400 MHz.

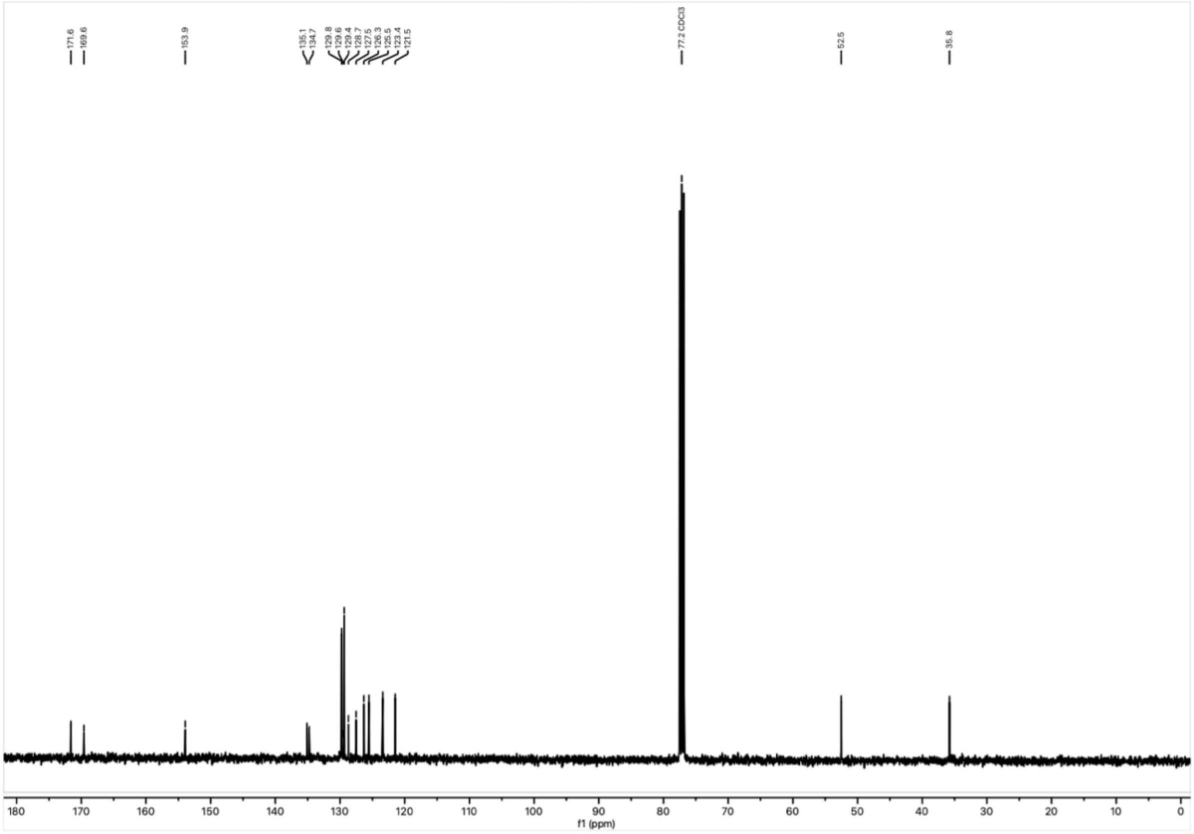


$^{13}C$ NMR spectrum of **11** in Chloroform-*d* at 101 MHz.

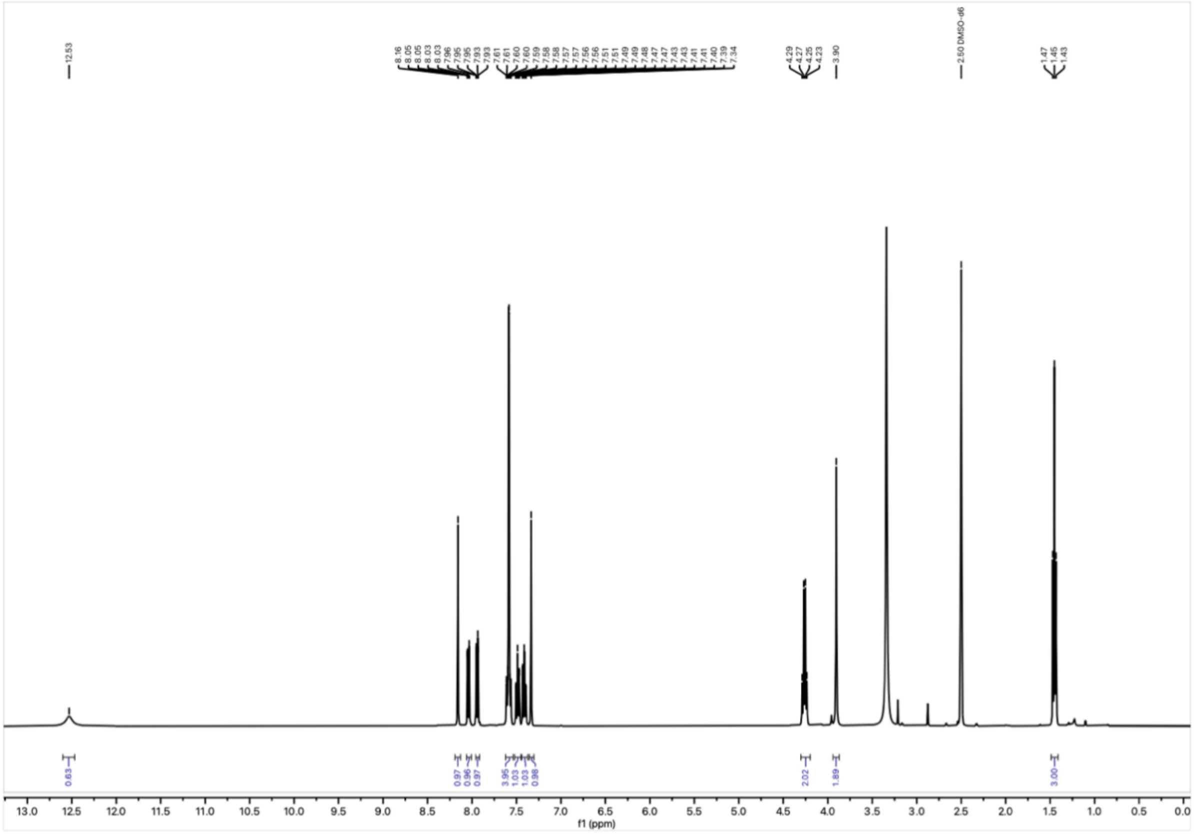


$^{1}H$ NMR spectrum of **13** in DMSO-$d_6$ at 400 MHz.

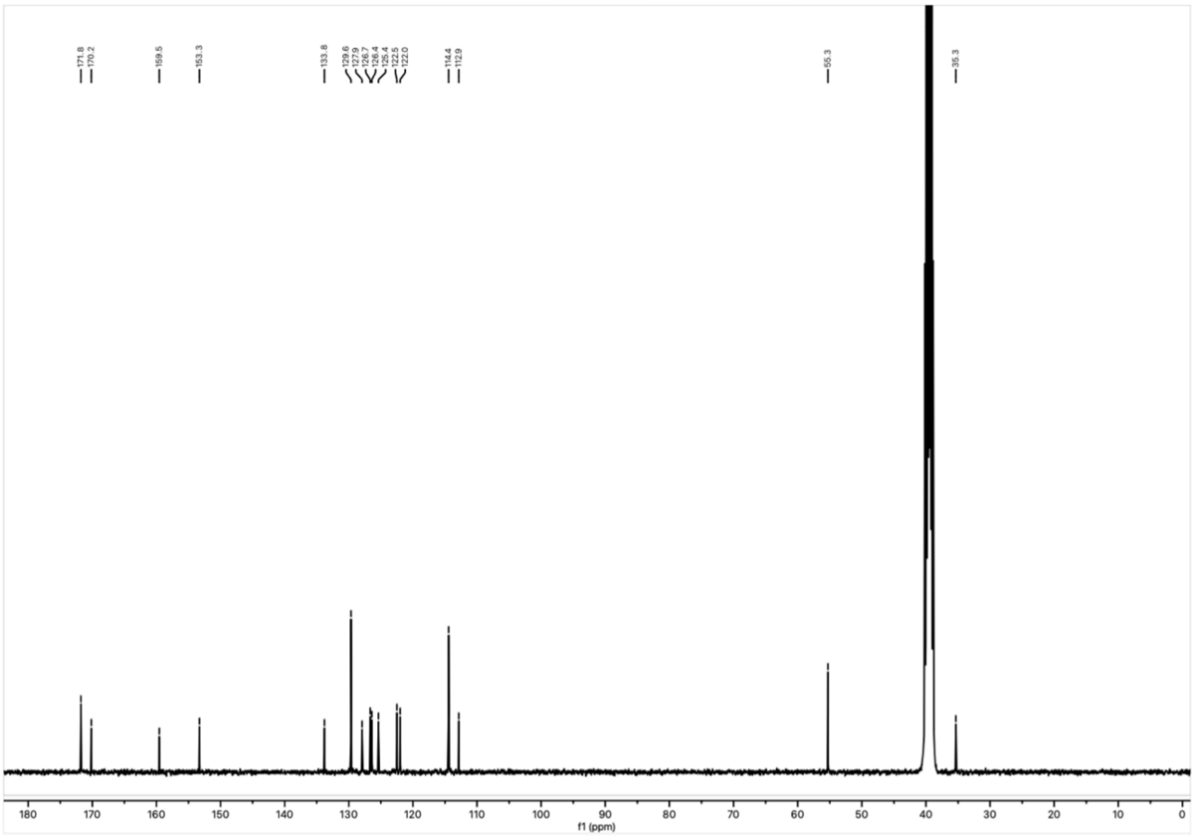

$^{13}C$ NMR spectrum of **13** in DMSO-$d_6$ at 101 MHz.

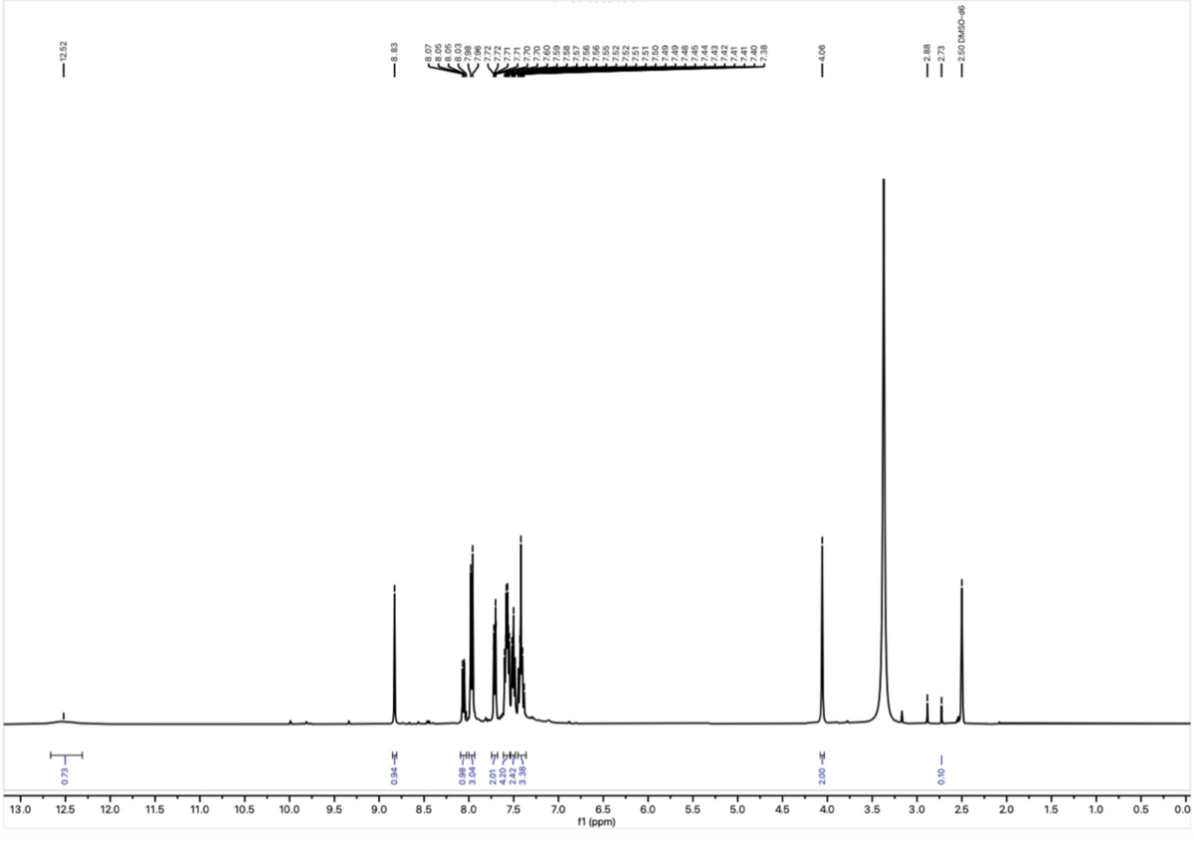

$^{1}H$ NMR spectrum of **14** in DMSO-$d_6$ at 400 MHz.

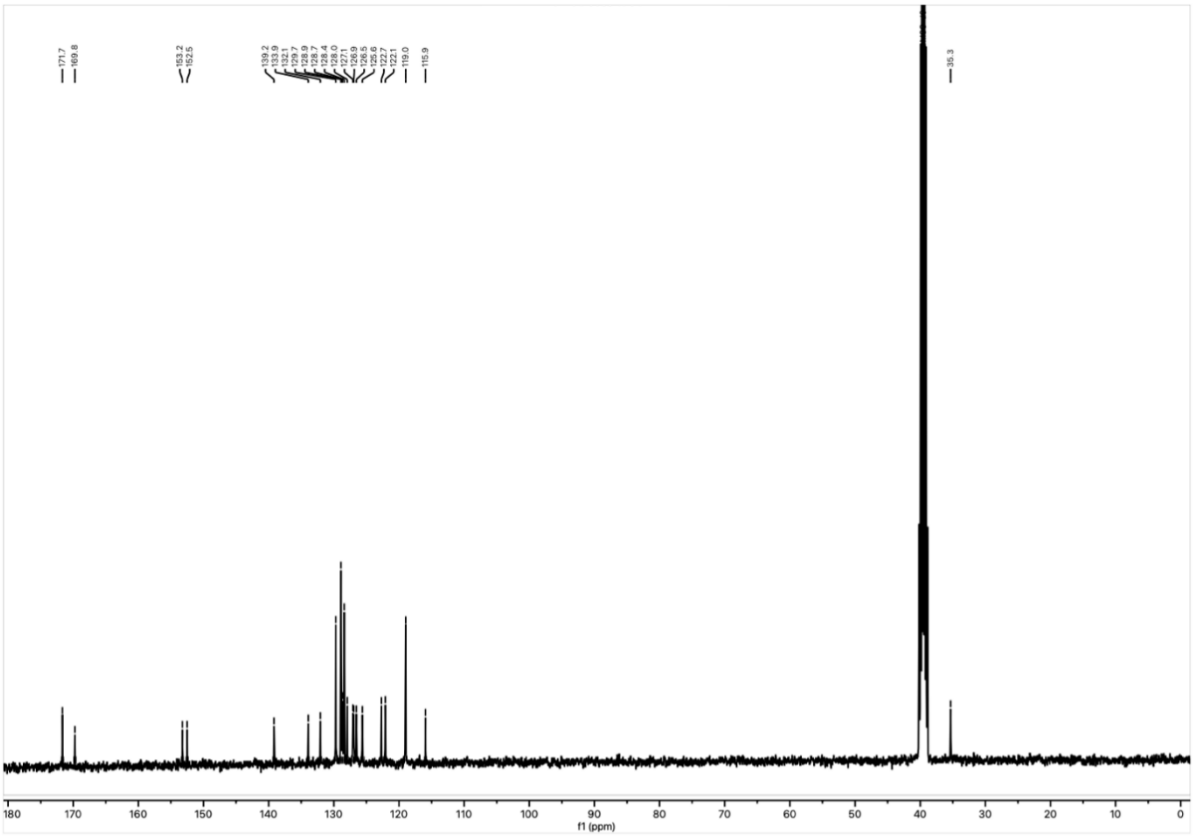


$^{13}C$ NMR spectrum of **14** in DMSO-$d_6$ at 101 MHz.

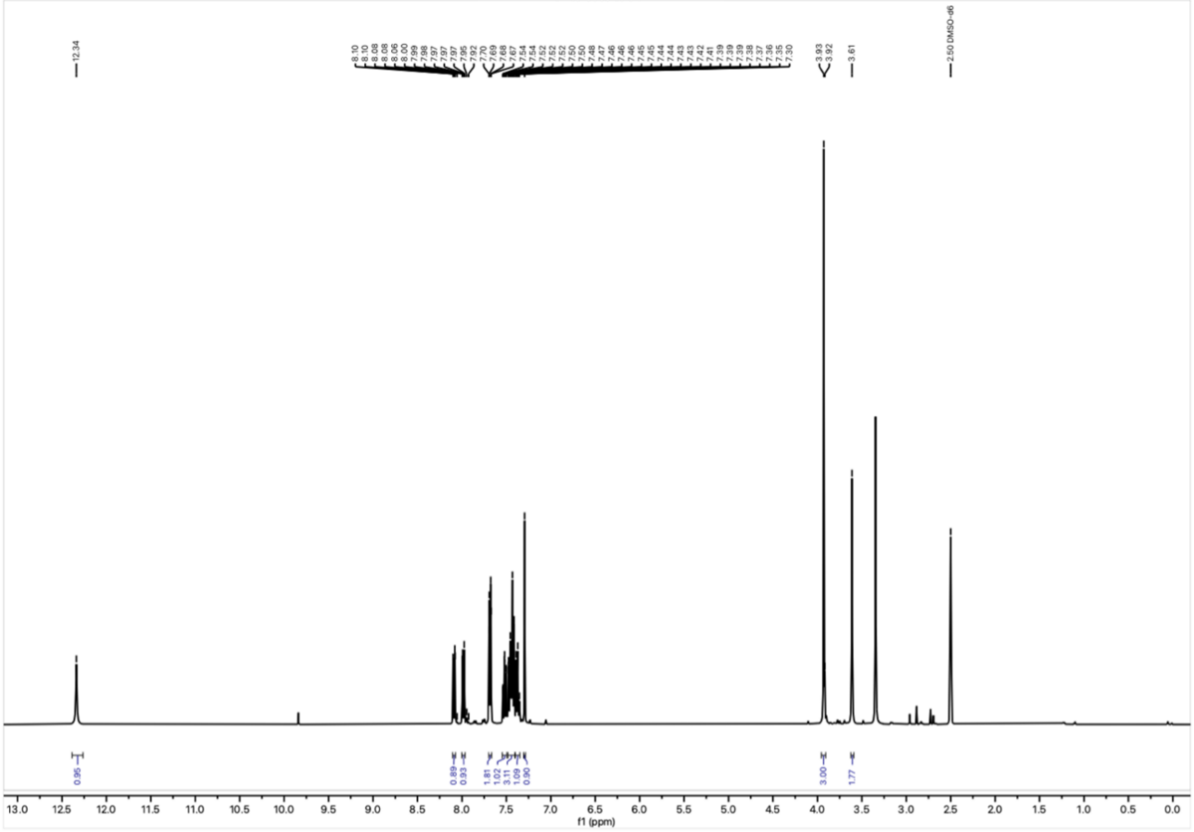


$^{1}H$ NMR spectrum of **15** in DMSO-$d_6$ at 400 MHz.

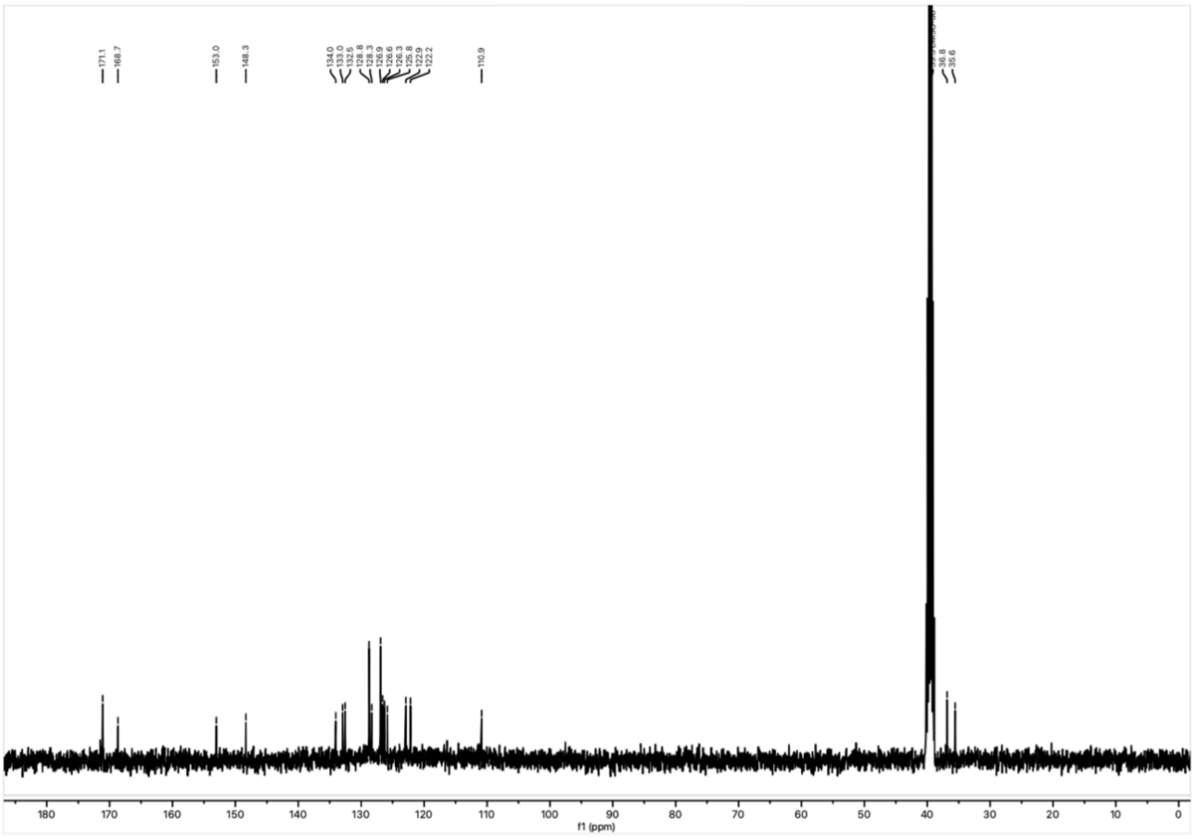

$^{13}$C NMR spectrum of **15** in DMSO-$d_6$ at 101 MHz.

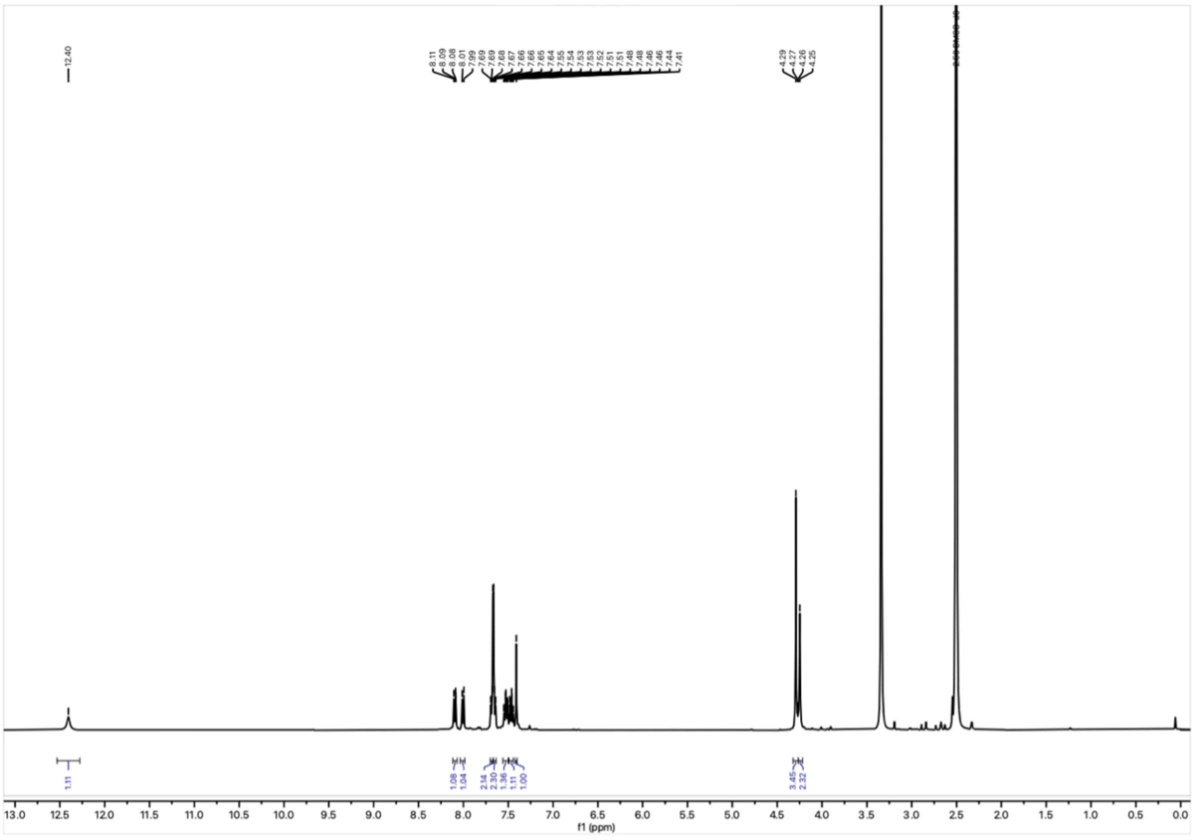

$^{1}$H NMR spectrum of **16** in DMSO-$d_6$ at 400 MHz.

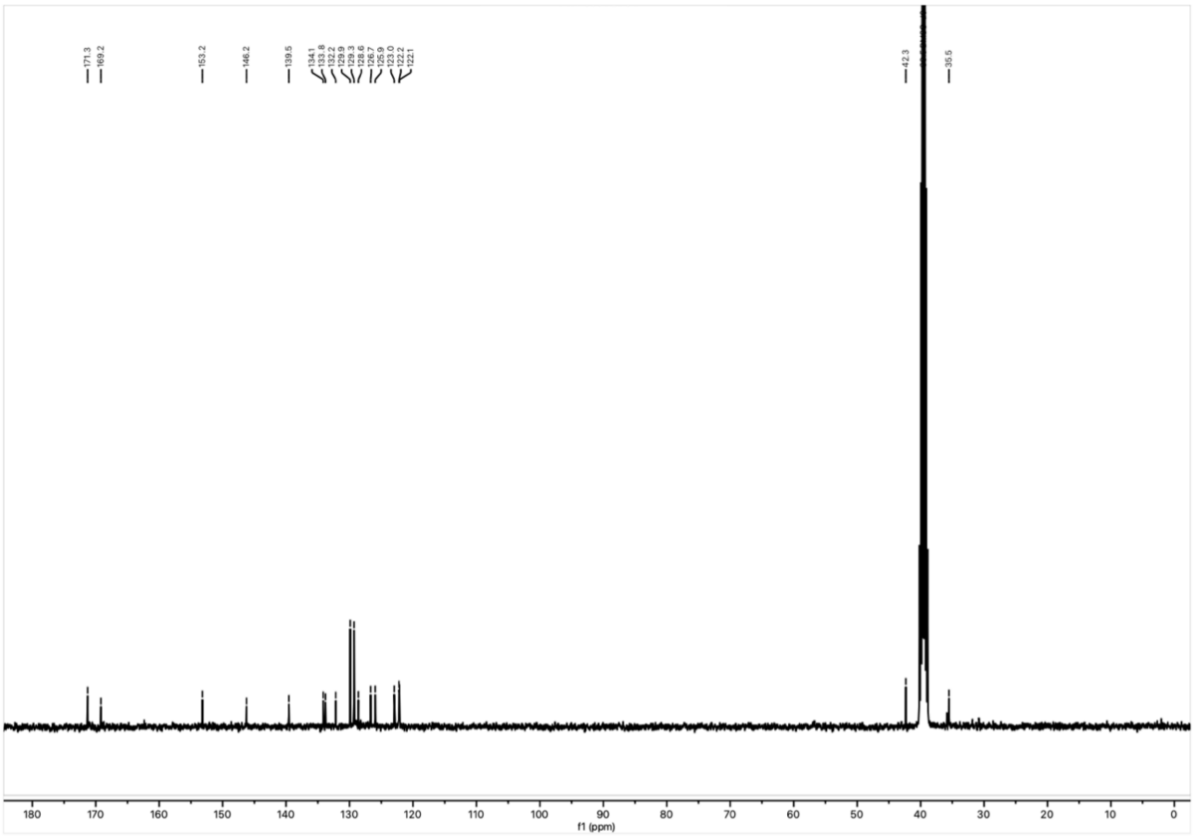


$^{13}C$ NMR spectrum of **16** in DMSO-$d_6$ at 101 MHz.

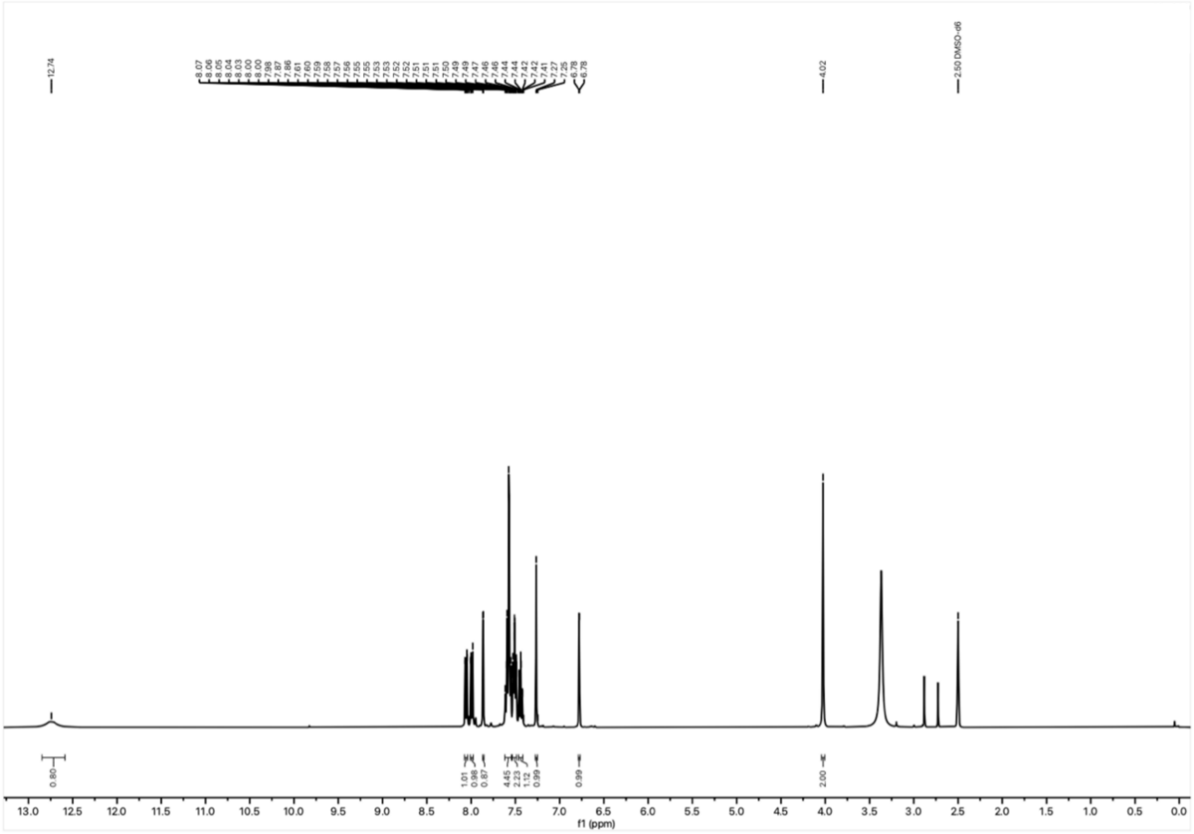


$^{1}H$ NMR spectrum of **17** in DMSO-$d_6$ at 400 MHz.

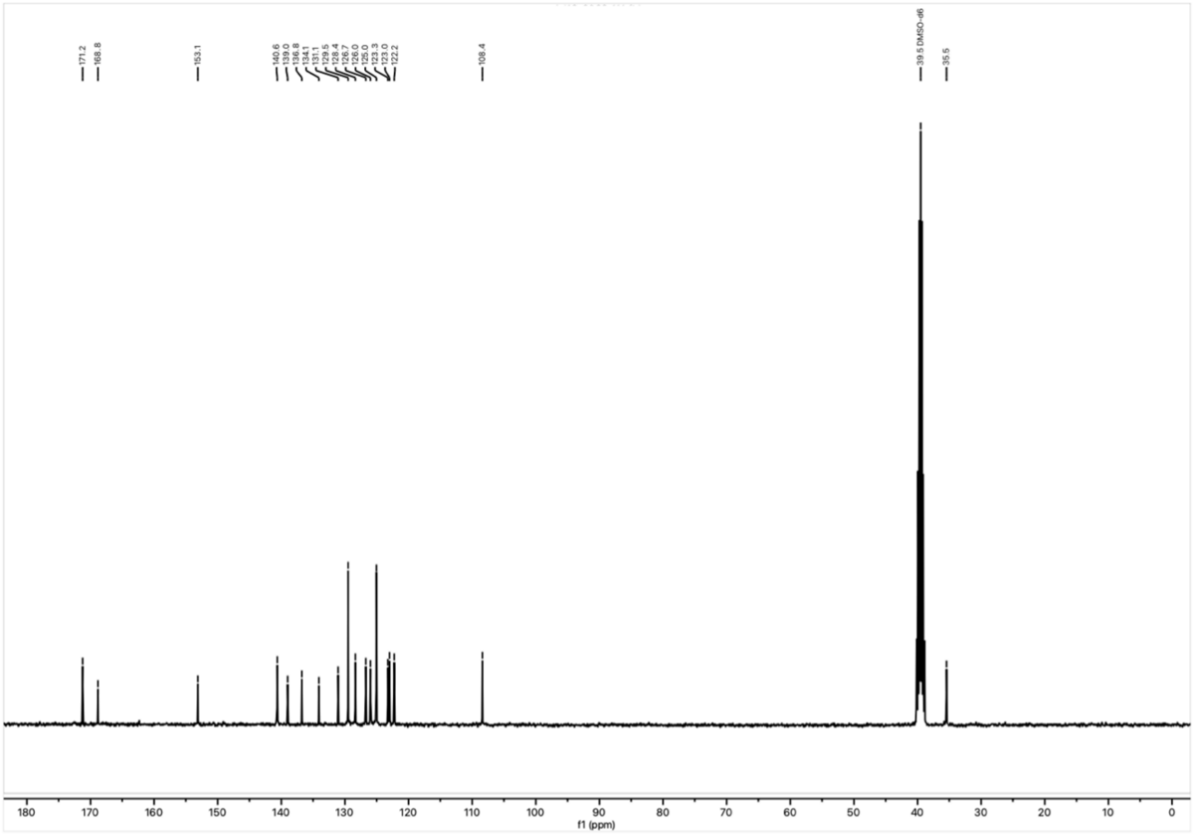


$^{13}C$ NMR spectrum of **17** in DMSO-$d_6$ at 101 MHz.

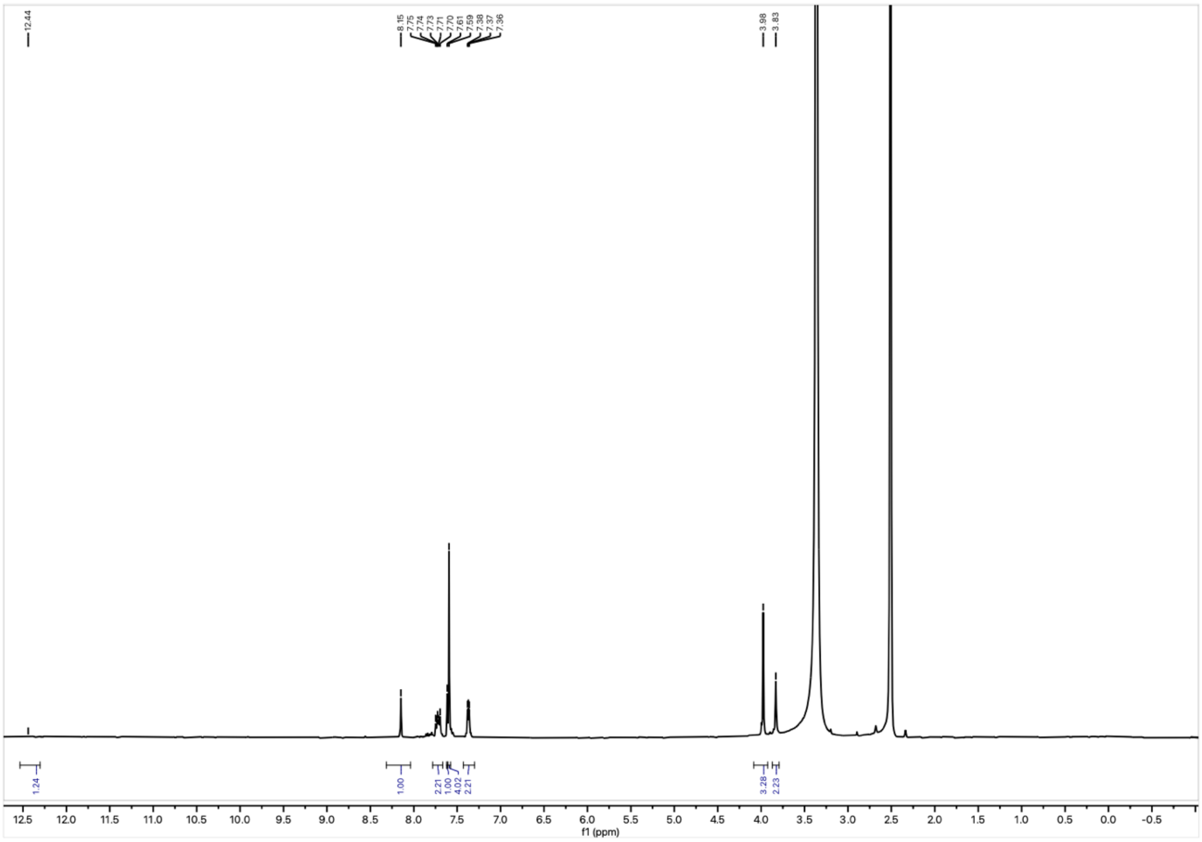


$^{1}H$ NMR spectrum of **18** in DMSO-$d_6$ at 400 MHz.

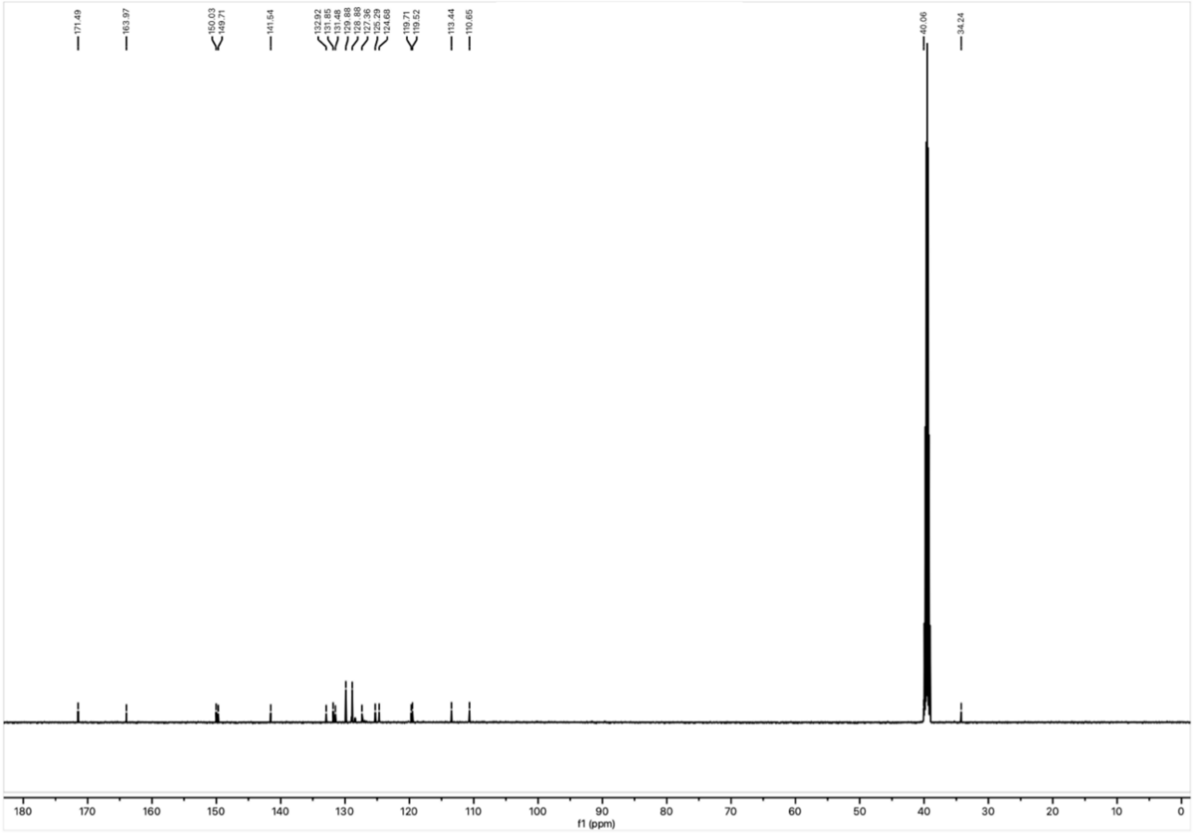

$^{13}C$ NMR spectrum of **18** in DMSO-$d_6$ at 151 MHz.

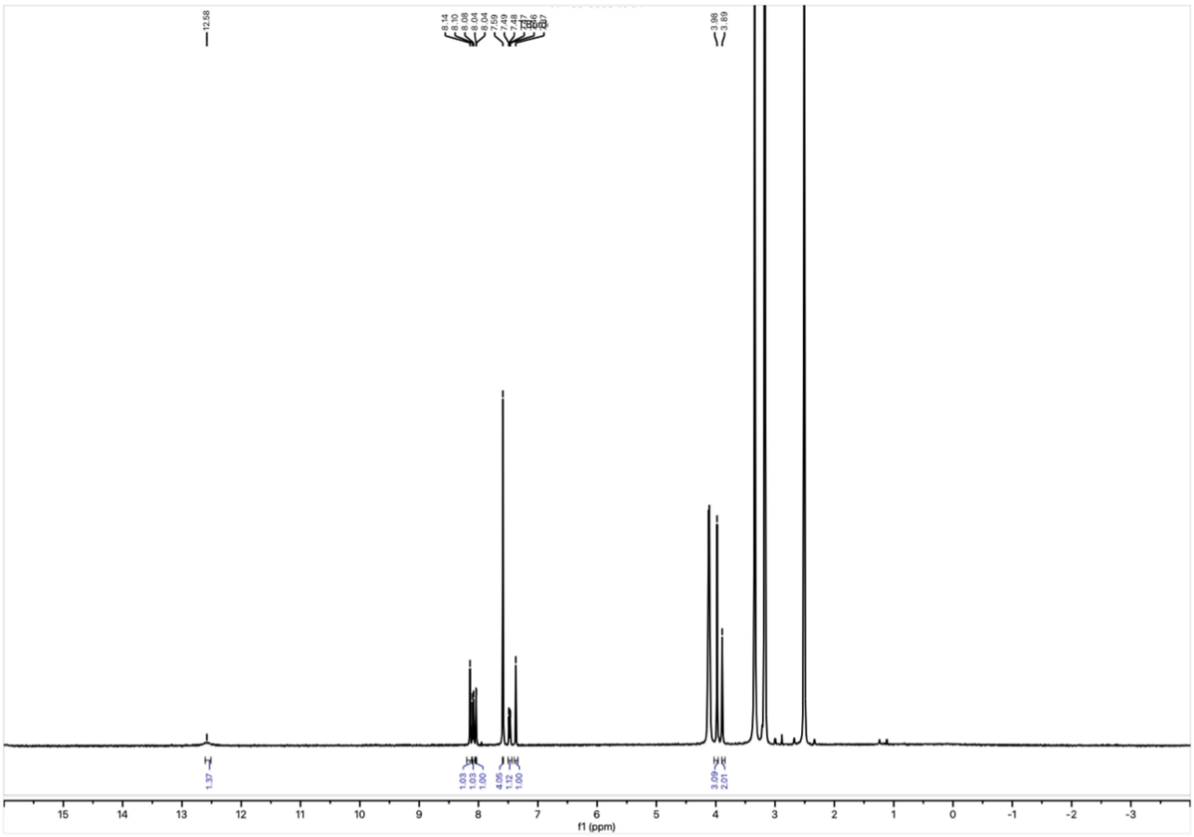

$^{1}H$ NMR spectrum of **19** in DMSO-$d_6$ at 400 MHz.

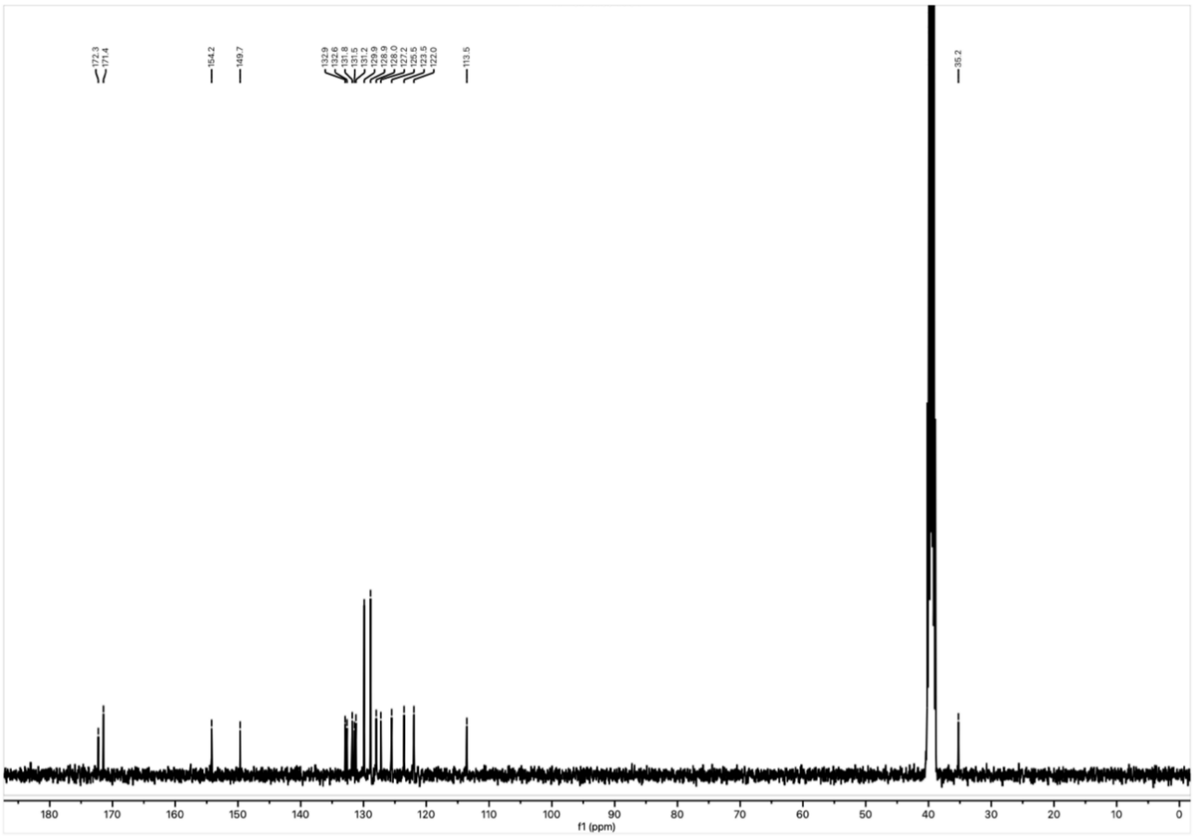

$^{13}C$ NMR spectrum of **19** in DMSO-$d_6$ at 101 MHz.

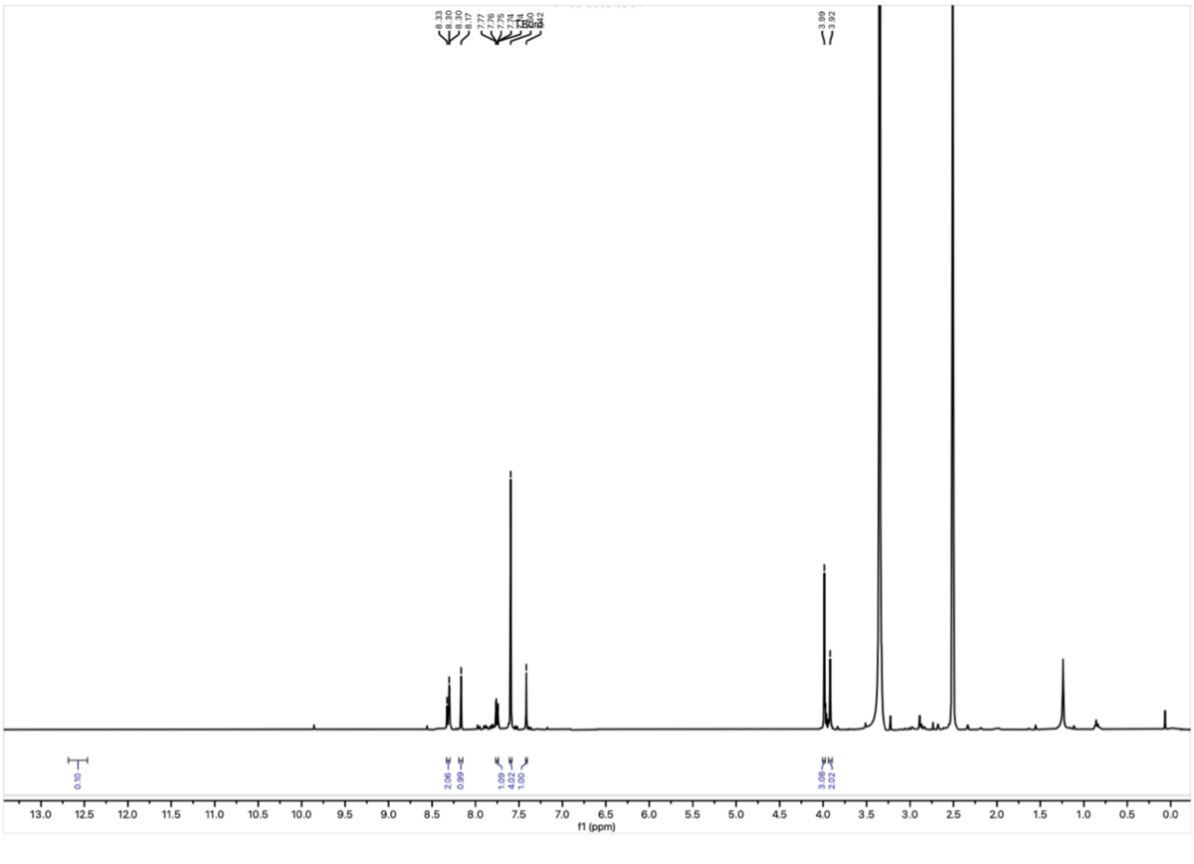

$^{1}H$ NMR spectrum of **20** in DMSO-$d_6$ at 400 MHz.

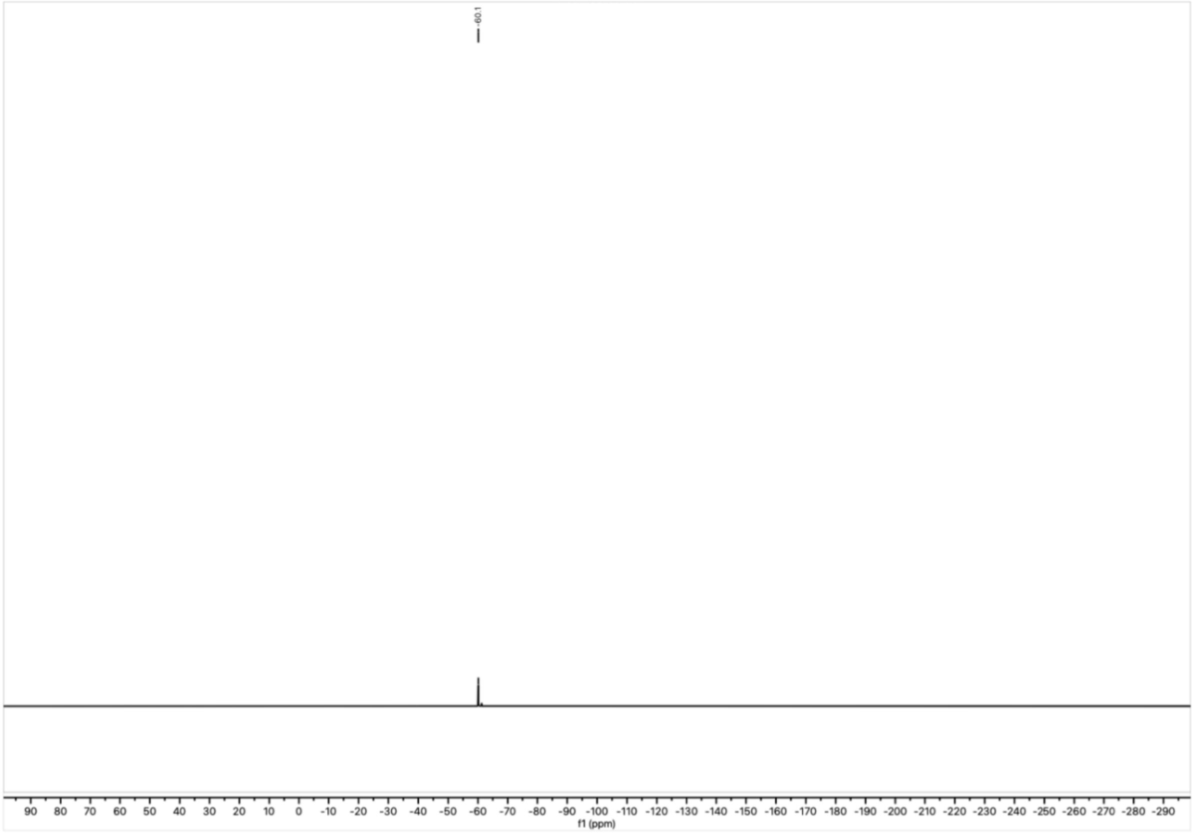


$^{19}$F NMR spectrum of **20** in DMSO-$d_6$ at 377 MHz.

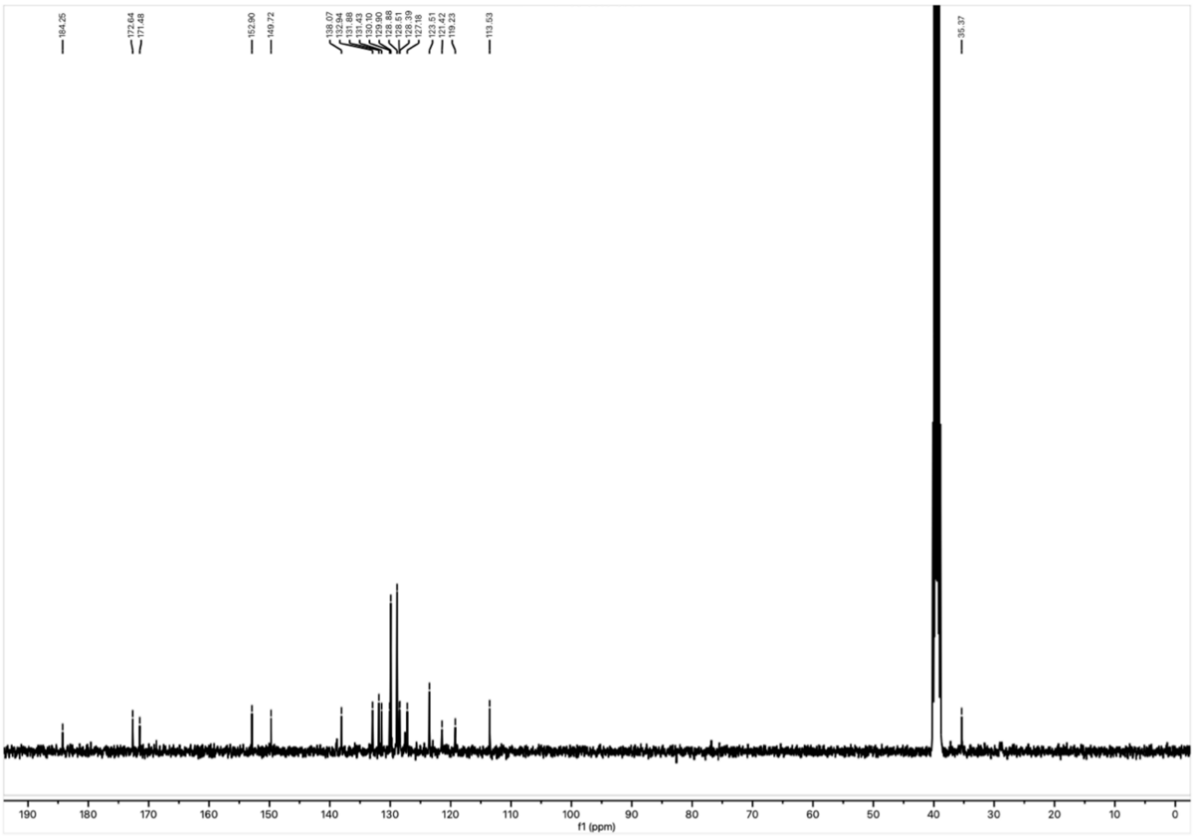


$^{13}$C NMR spectrum of **20** in DMSO-$d_6$ at 101 MHz.

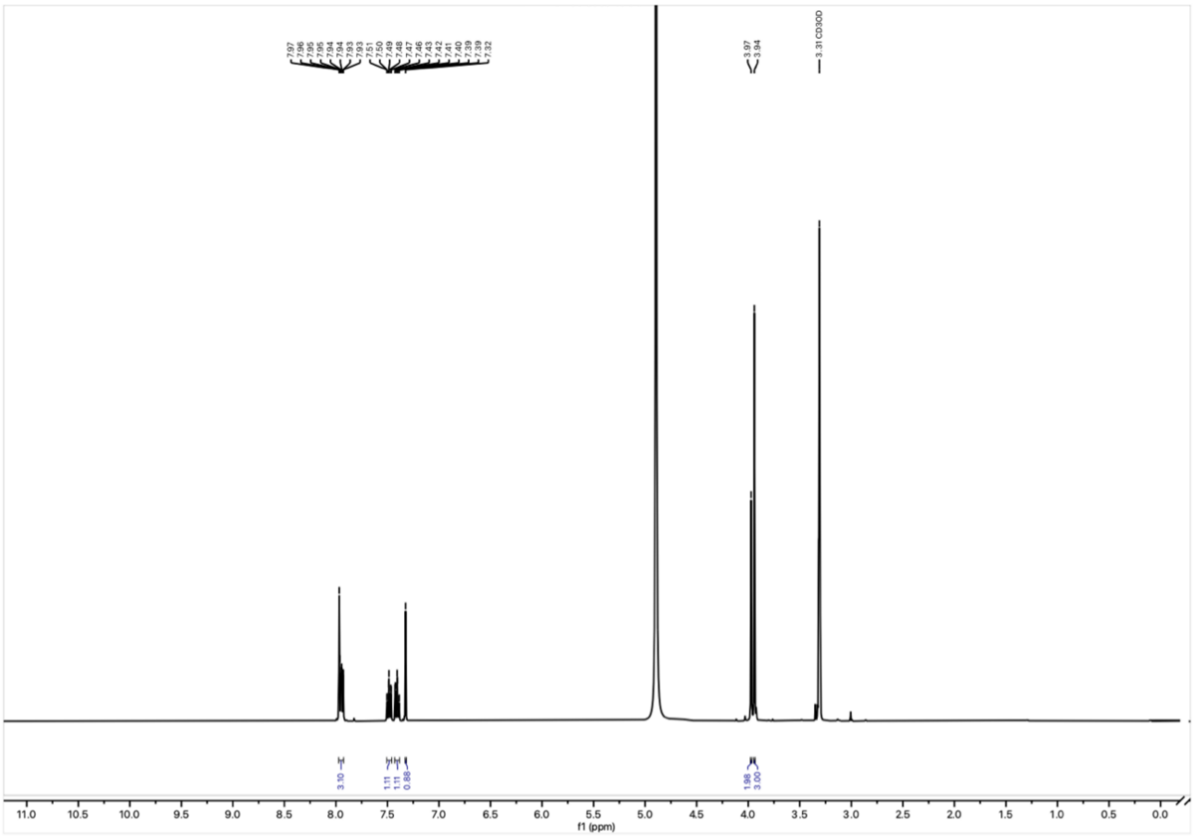


$^1$H NMR spectrum of **21** in MeOD-$d_4$ at 500 MHz.

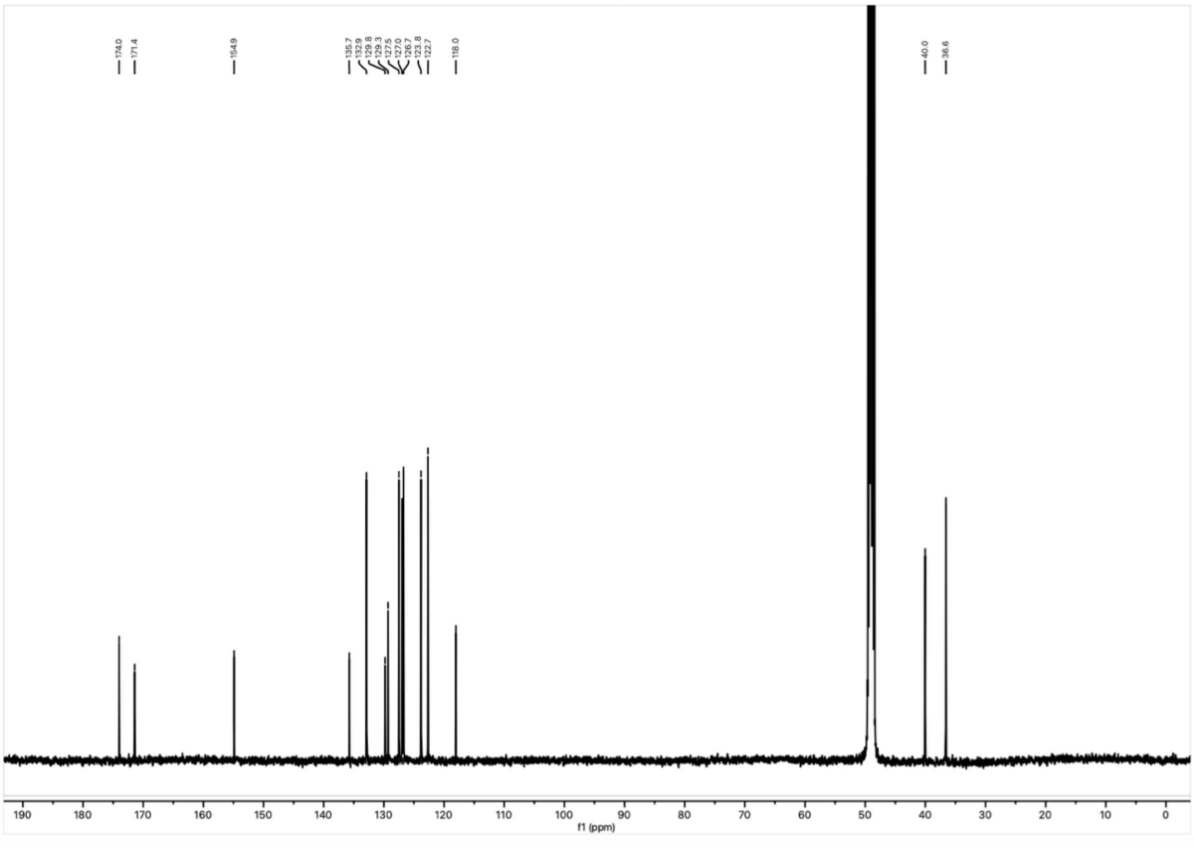


$^{13}$C NMR spectrum of **21** in MeOD-$d_4$ at 125 MHz.

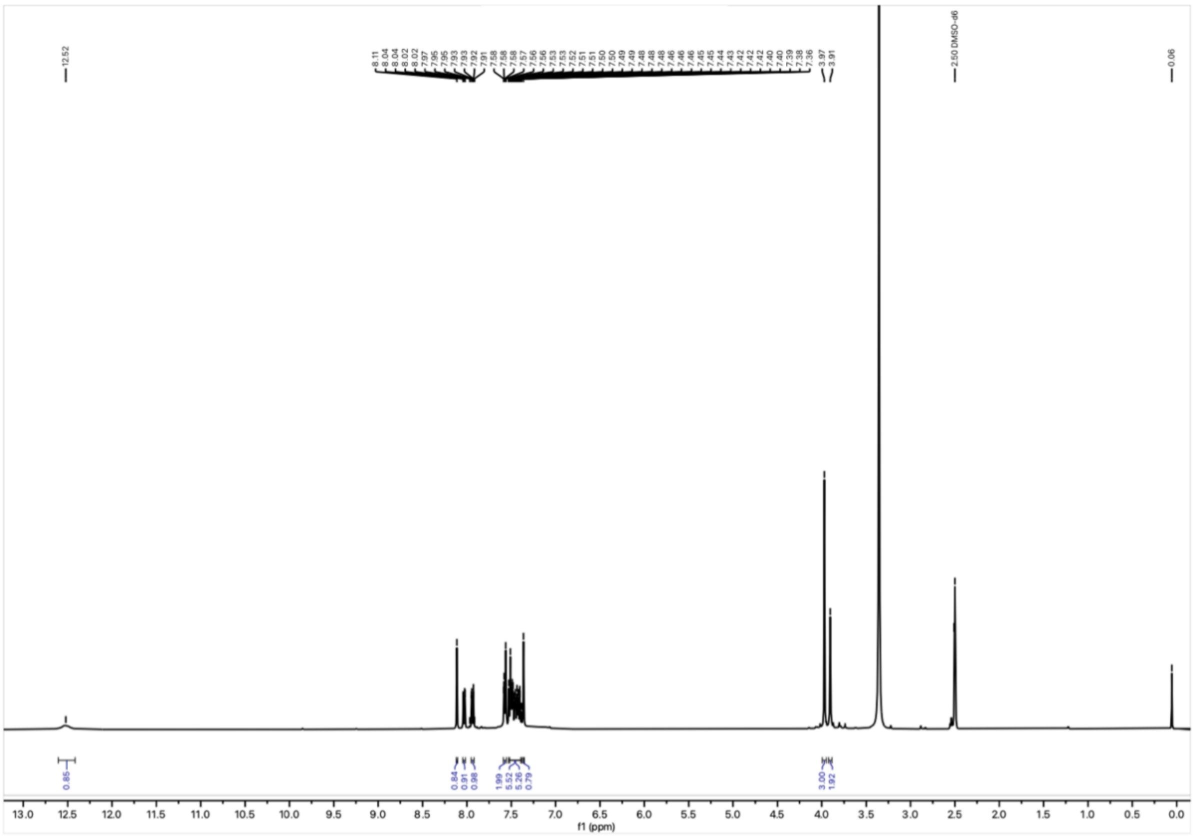


$^{1}$H NMR spectrum of **22** in DMSO-$d_6$ at 400 MHz.

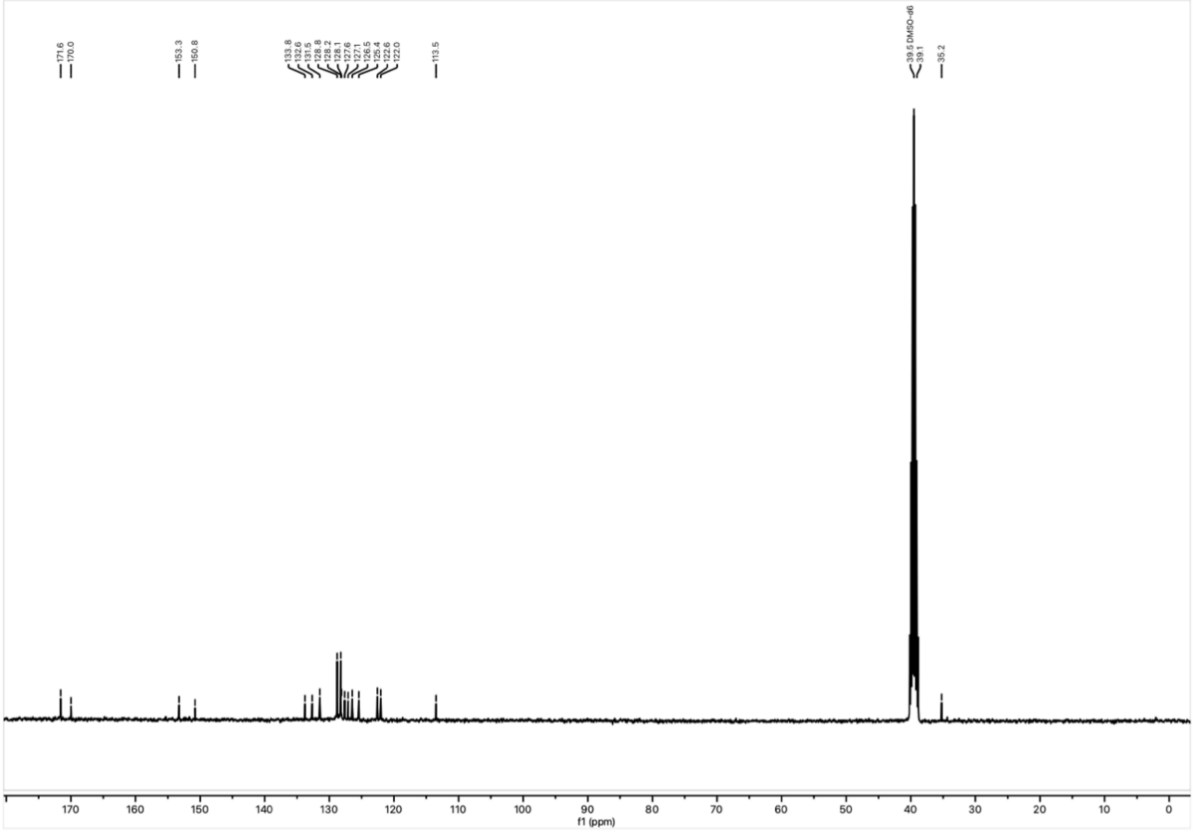


$^{13}$C NMR spectrum of **22** in DMSO-$d_6$ at 101 MHz.

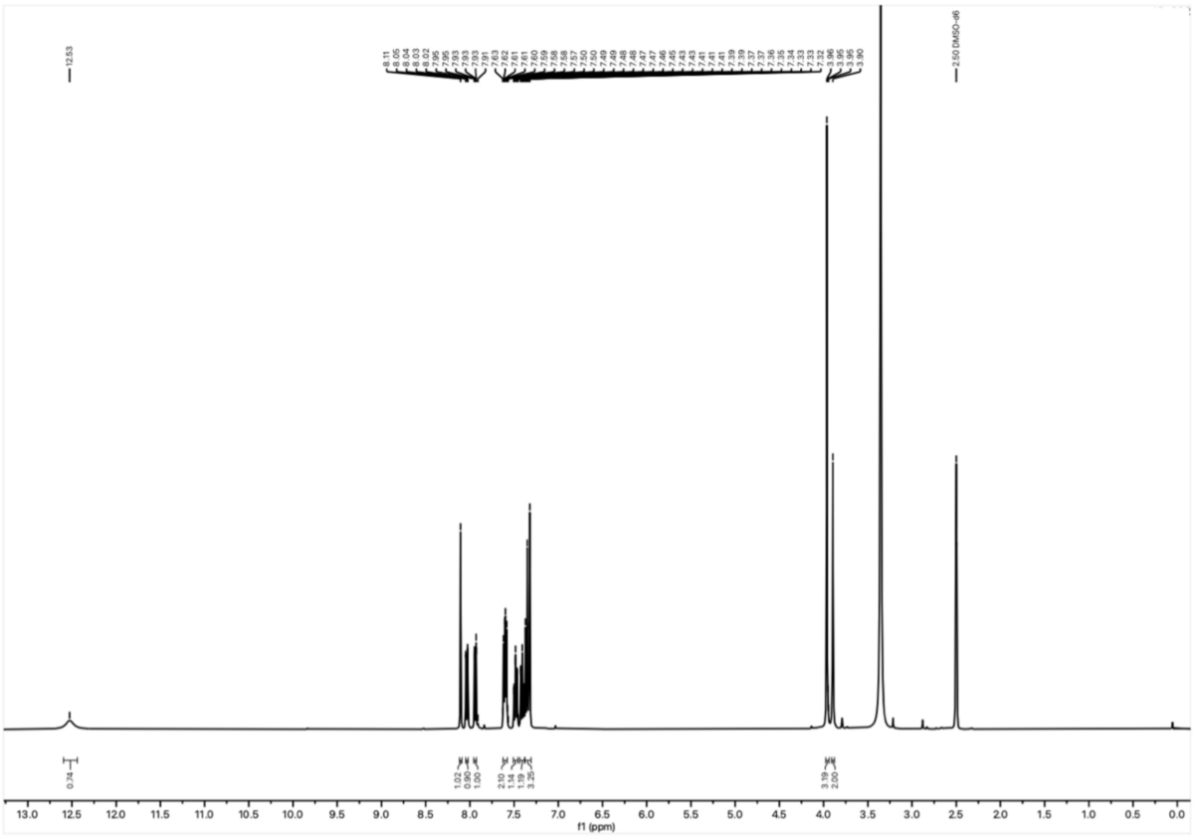

[1]H NMR spectrum of **23** in DMSO-$d_6$ at 500 MHz.

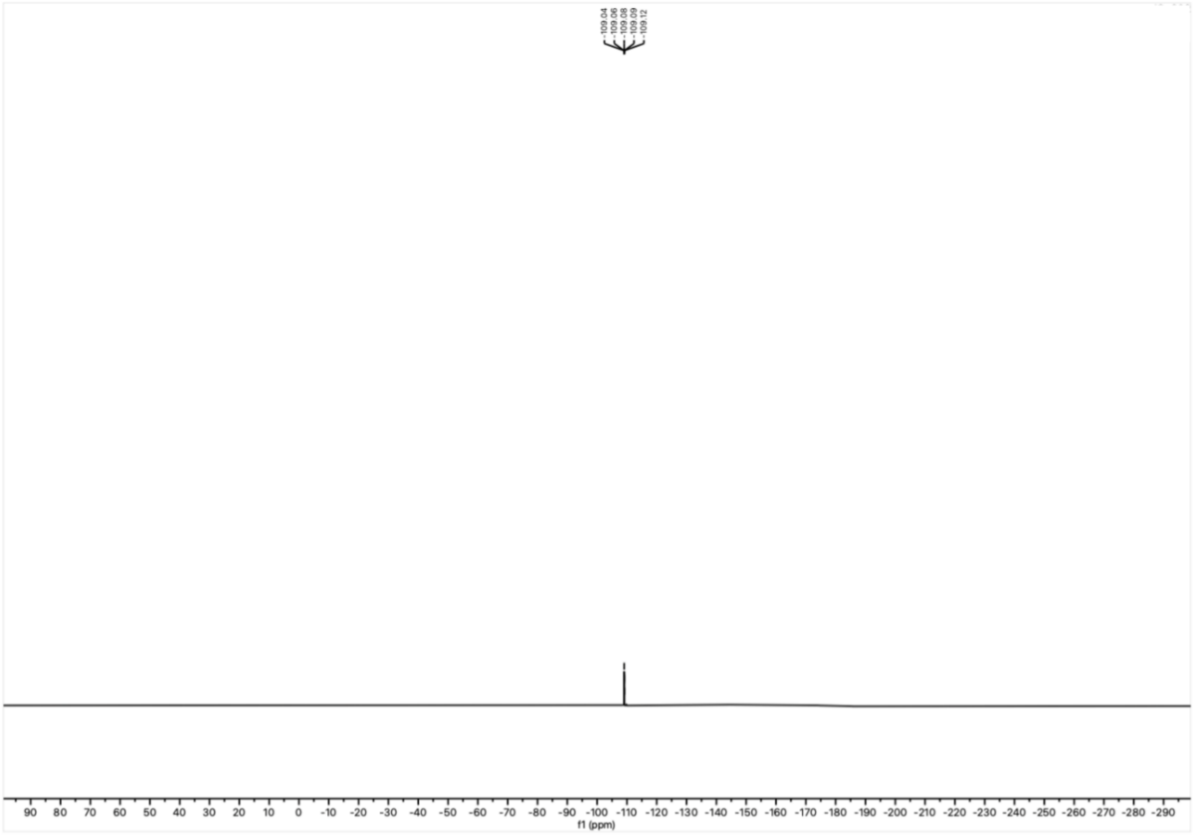

[19]F NMR spectrum of **23** in DMSO-$d_6$ at 377 MHz.

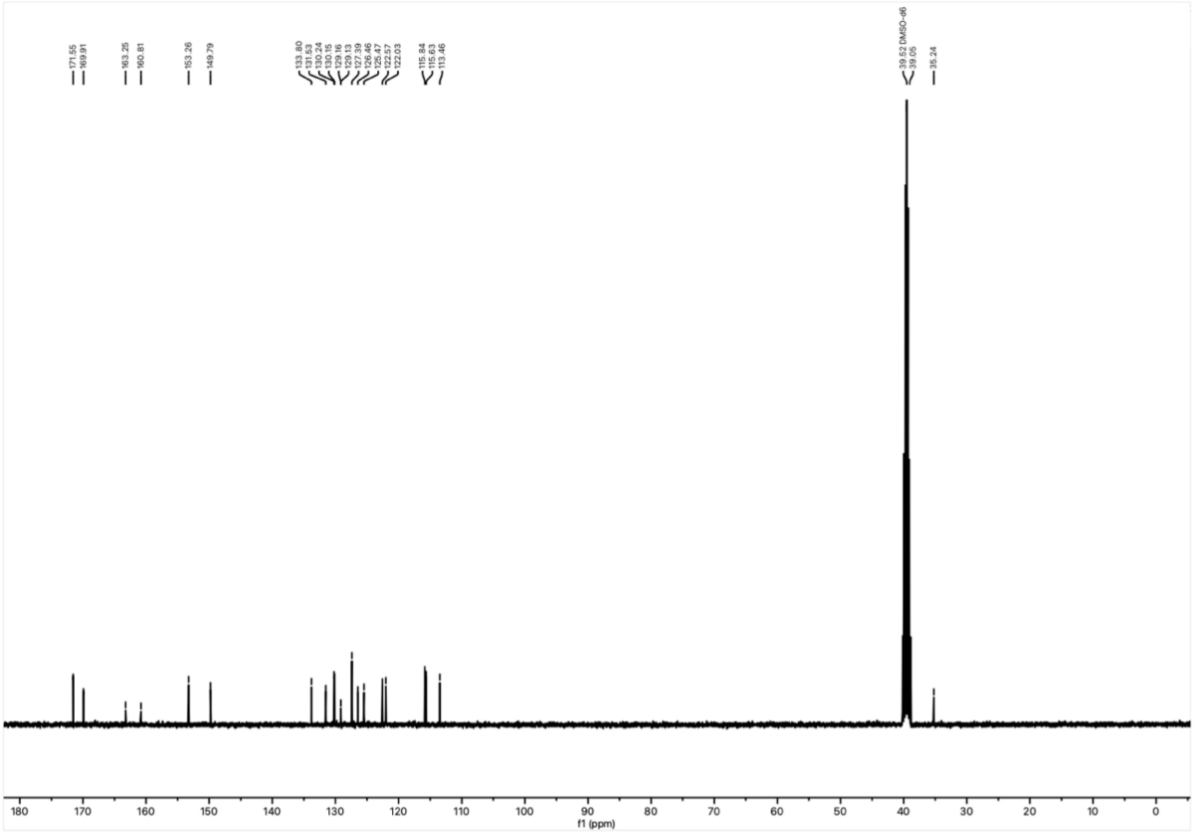

$^{13}C$ NMR spectrum of **23** in DMSO-$d_6$ at 125 MHz.

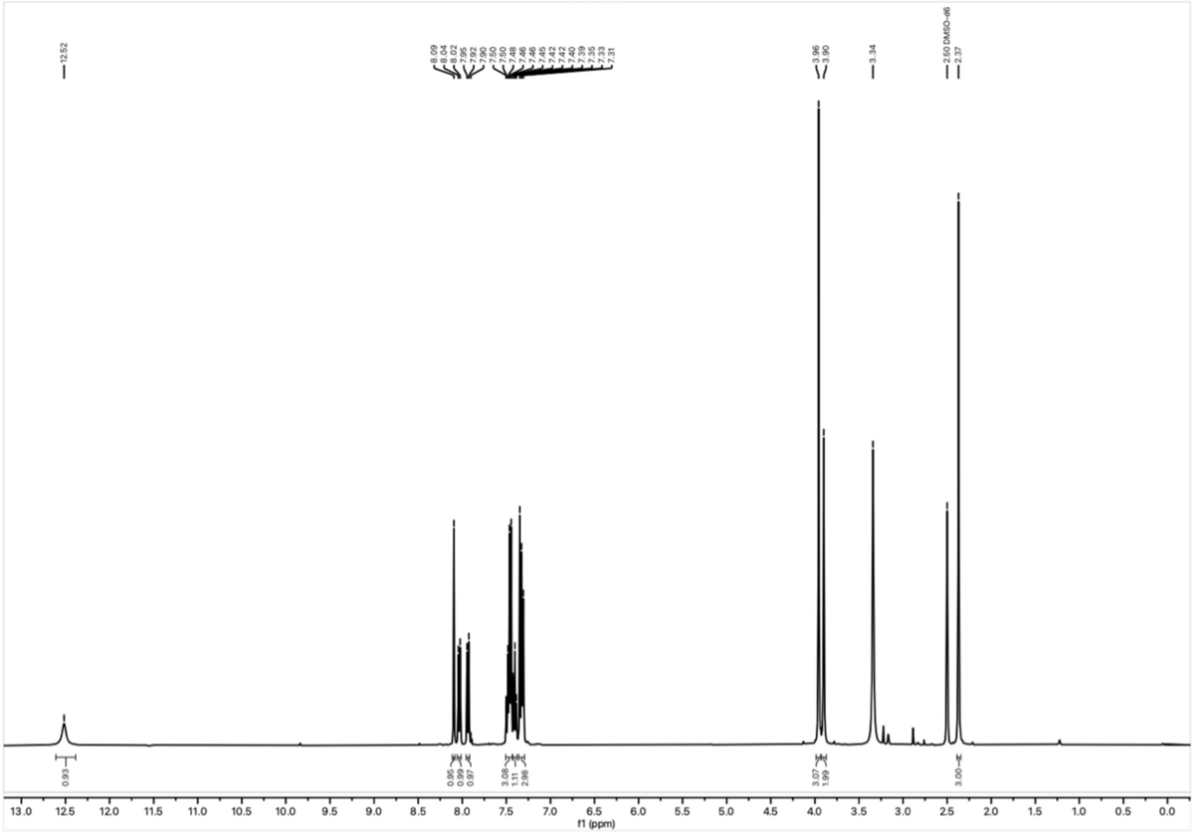

$^1H$ NMR spectrum of **24** in DMSO-$d_6$ at 500 MHz.

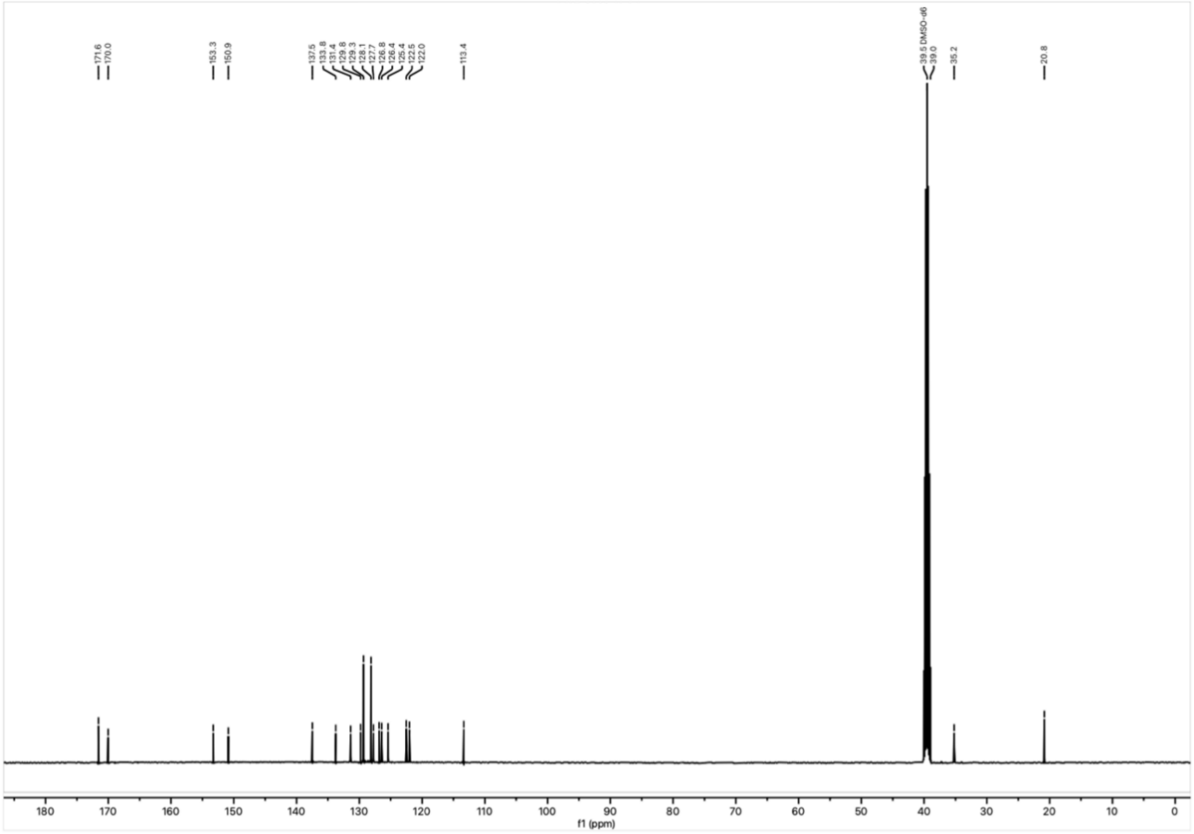

$^{13}C$ NMR spectrum of **24** in DMSO-$d_6$ at 125 MHz.

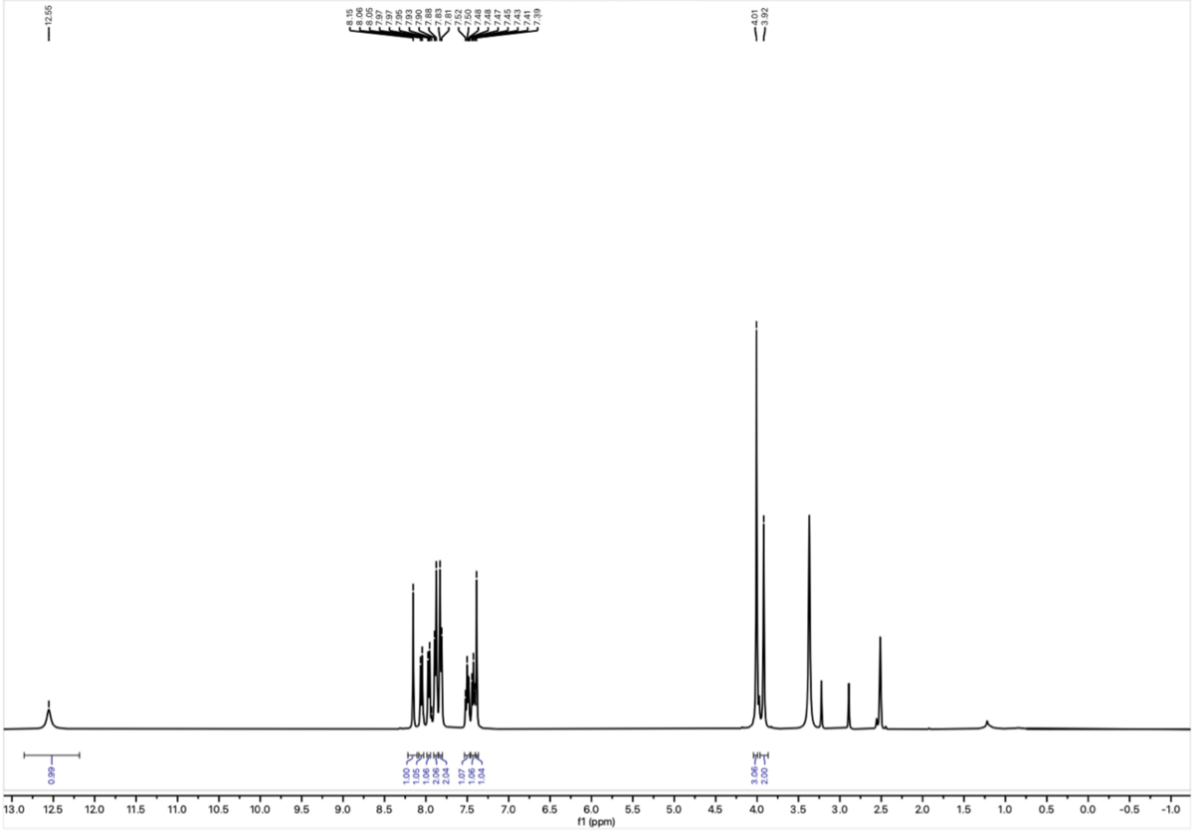

$^{1}H$ NMR spectrum of **25** in DMSO-$d_6$ at 400 MHz.

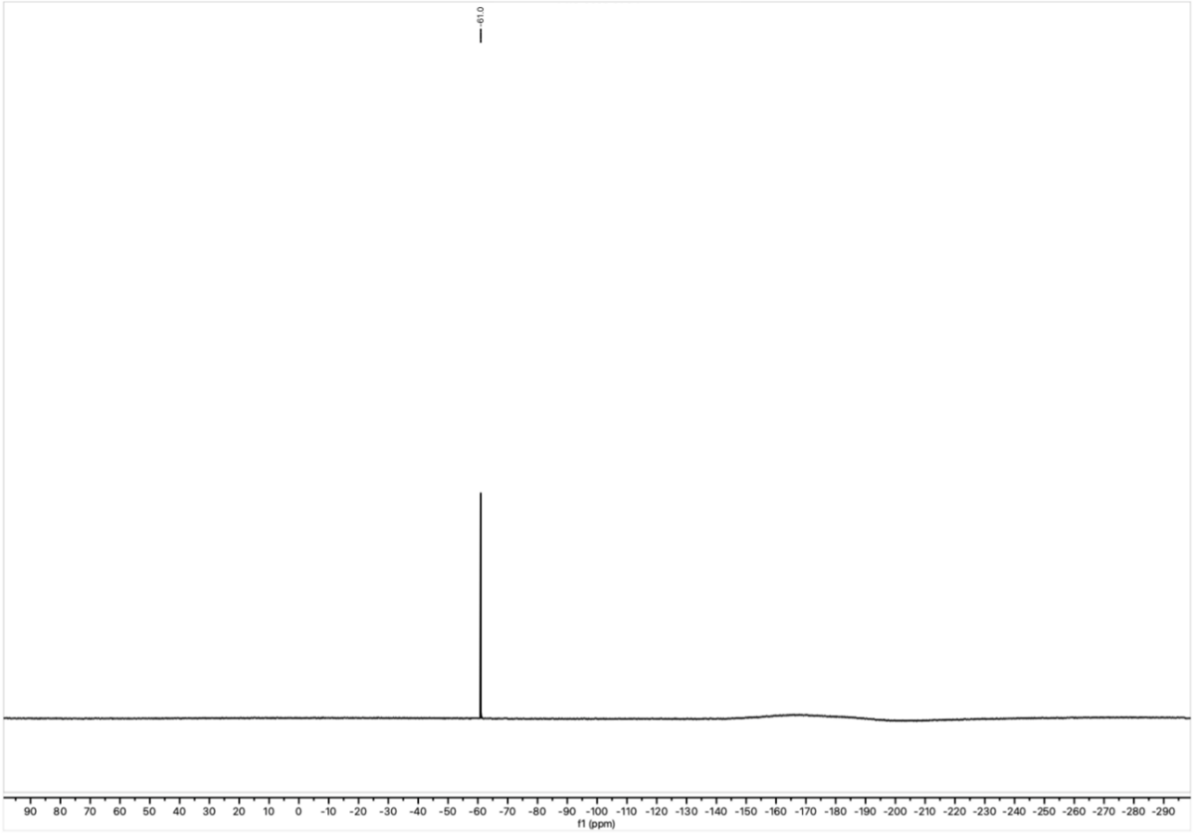


$^{19}$F NMR spectrum of **25** in DMSO-$d_6$ at 377 MHz.

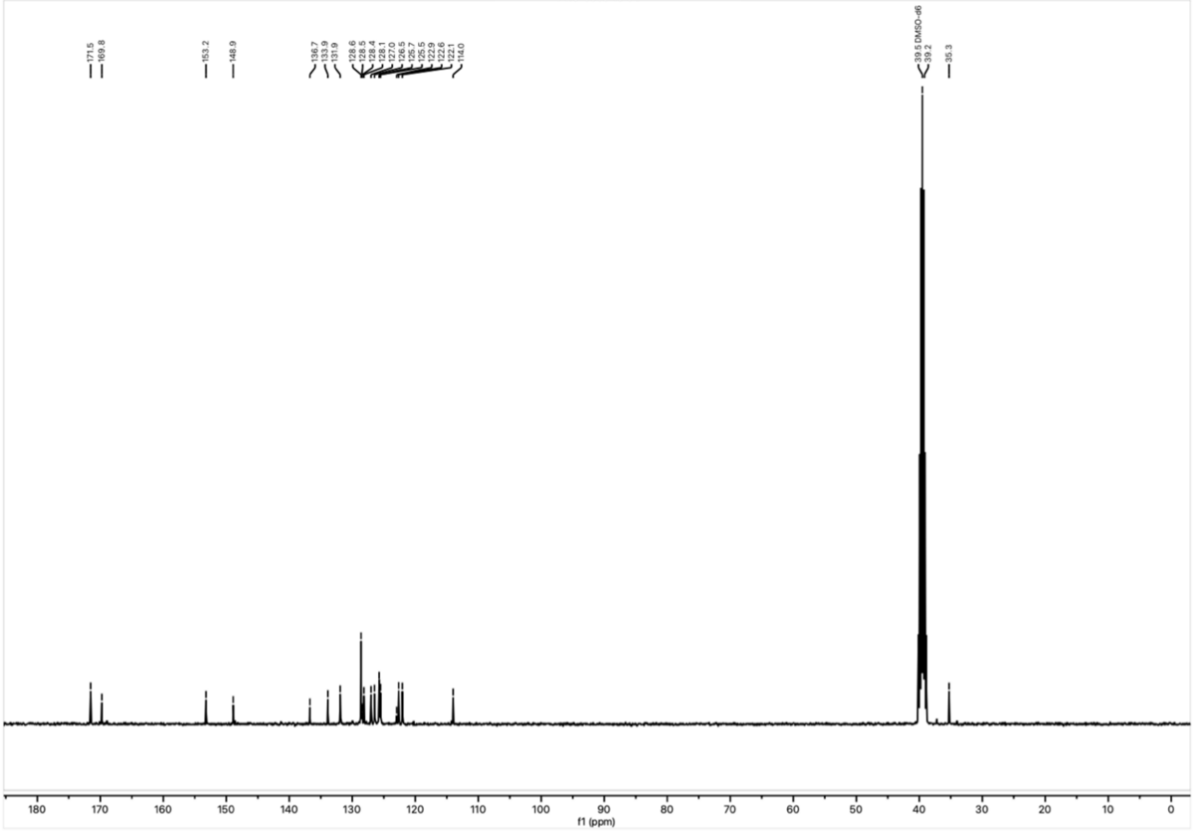


$^{13}$C NMR spectrum of **25** in DMSO-$d_6$ at 101 MHz.

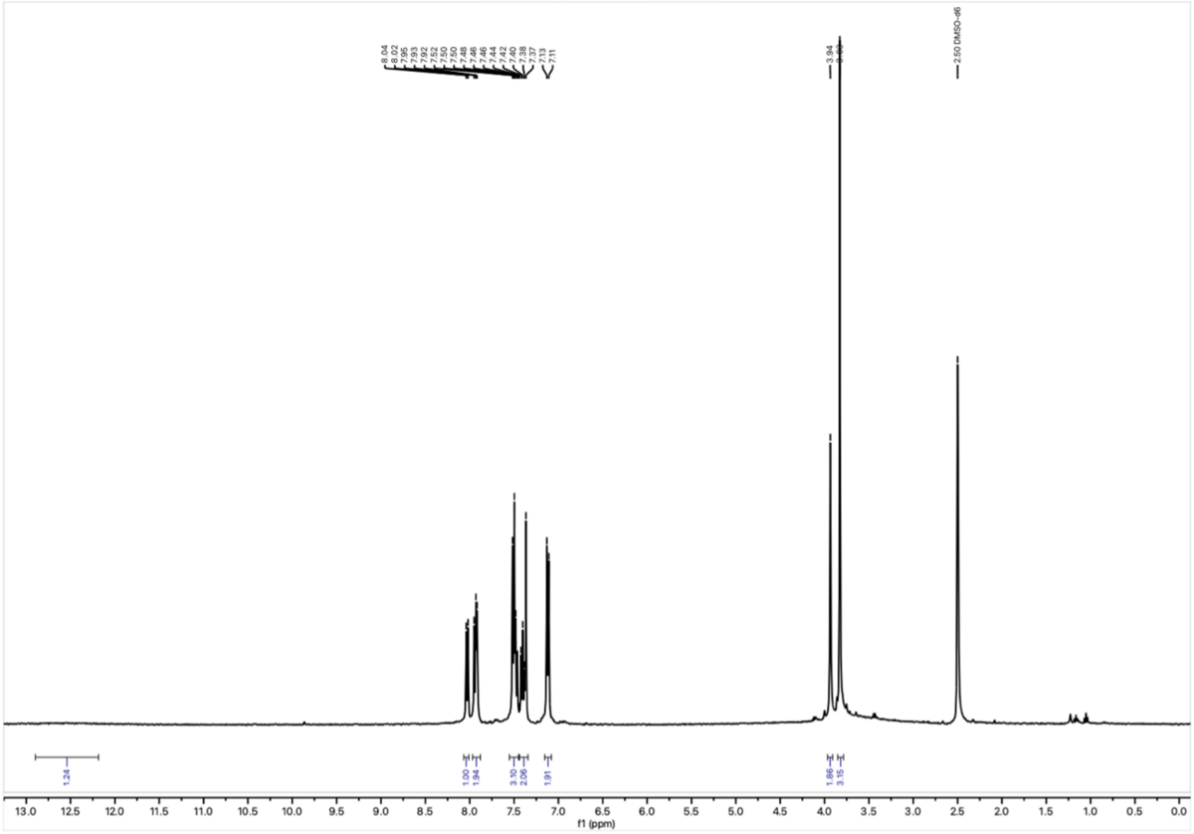


$^{1}H$ NMR spectrum of **26** in DMSO-$d_6$ at 400 MHz.

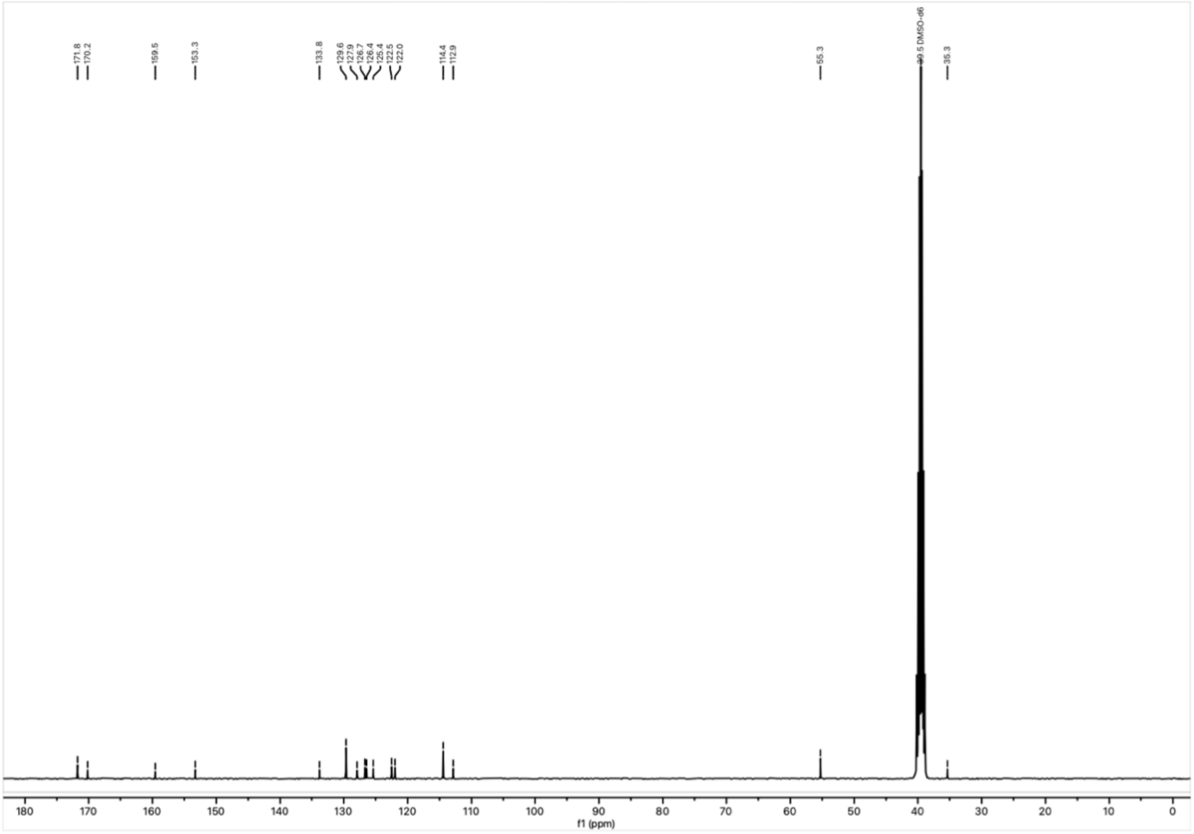


$^{13}C$ NMR spectrum of **26** in DMSO-$d_6$ at 101 MHz.

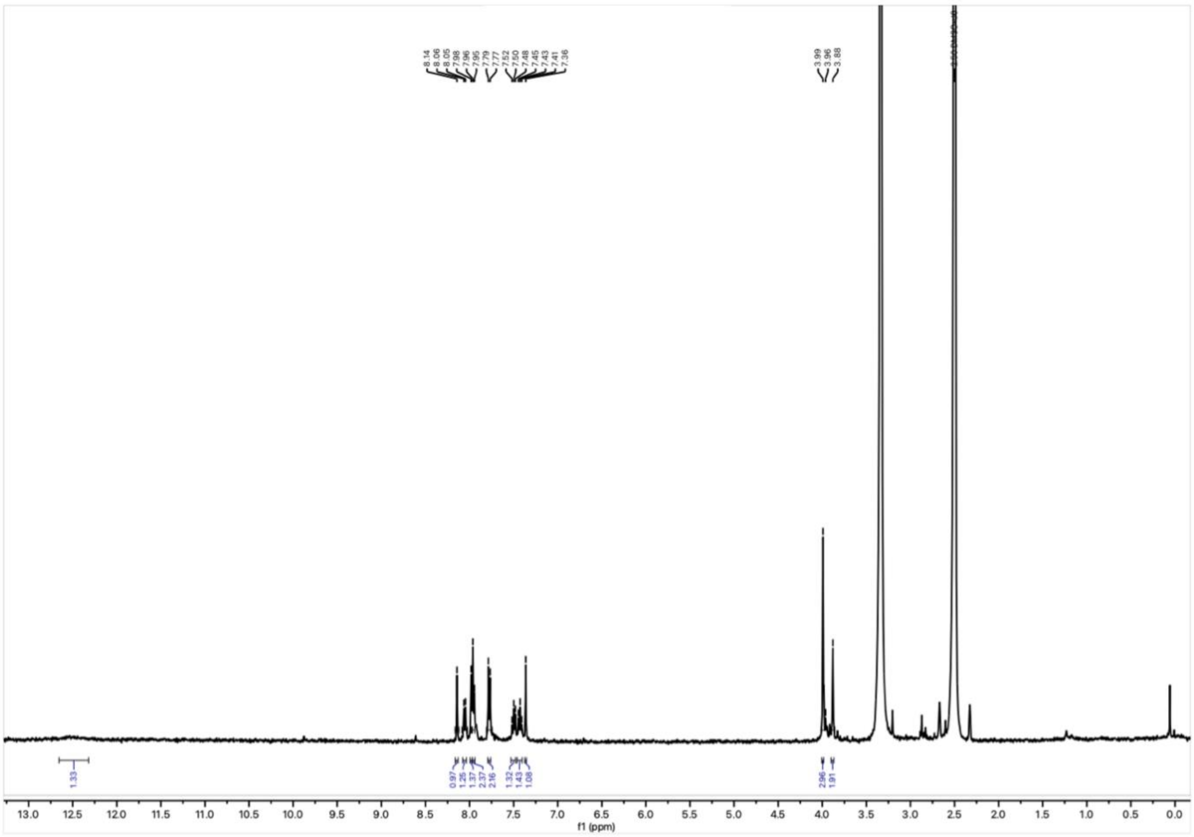

$^1H$ NMR spectrum of **27** in DMSO-$d_6$ at 400 MHz.

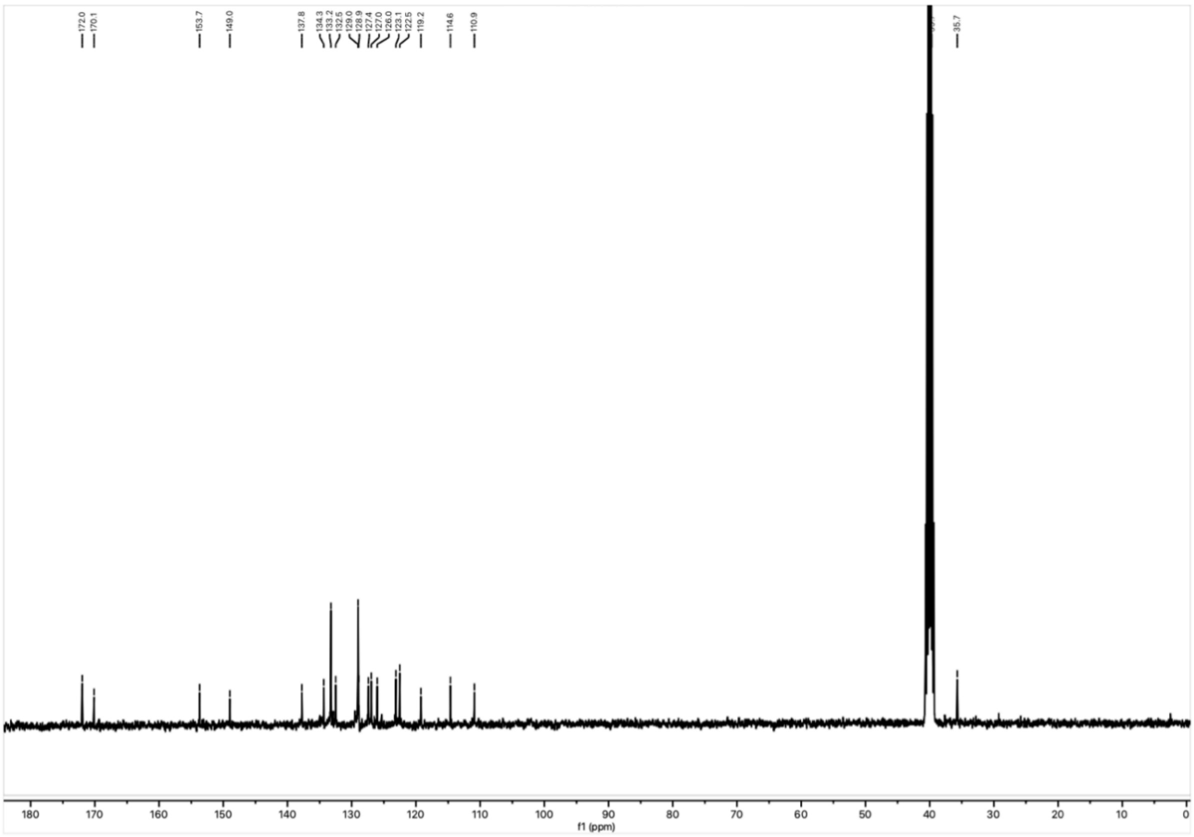

$^{13}C$ NMR spectrum of **27** in DMSO-$d_6$ at 101 MHz.

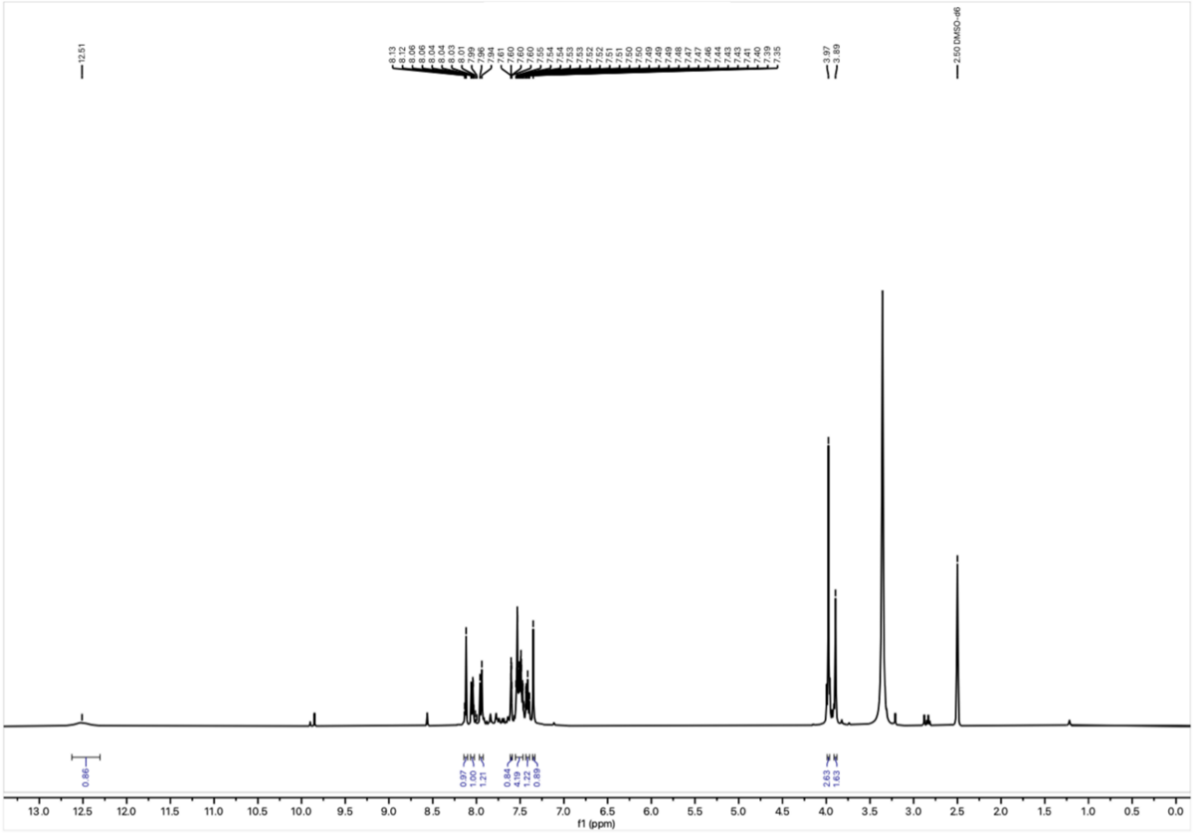

$^{1}H$ NMR spectrum of **28** in DMSO-$d_6$ at 400 MHz.

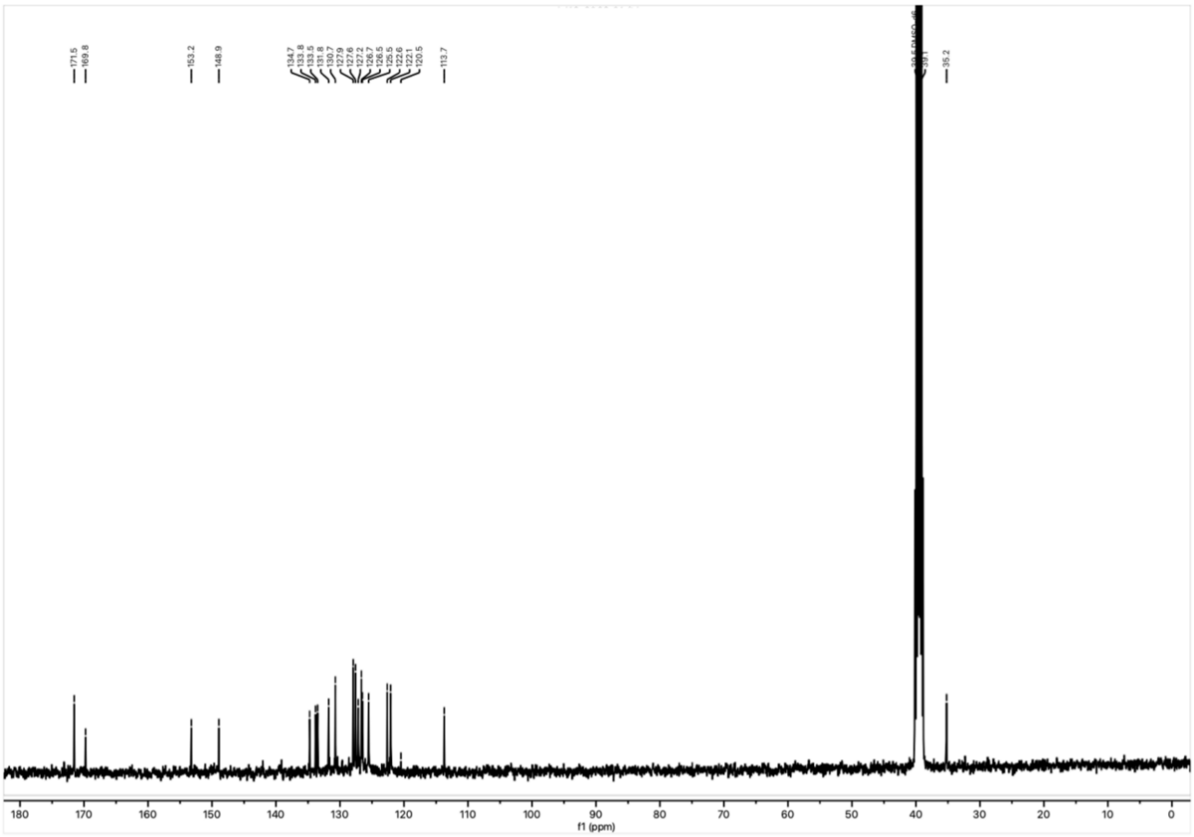

$^{13}C$ NMR spectrum of **28** in DMSO-$d_6$ at 101 MHz.

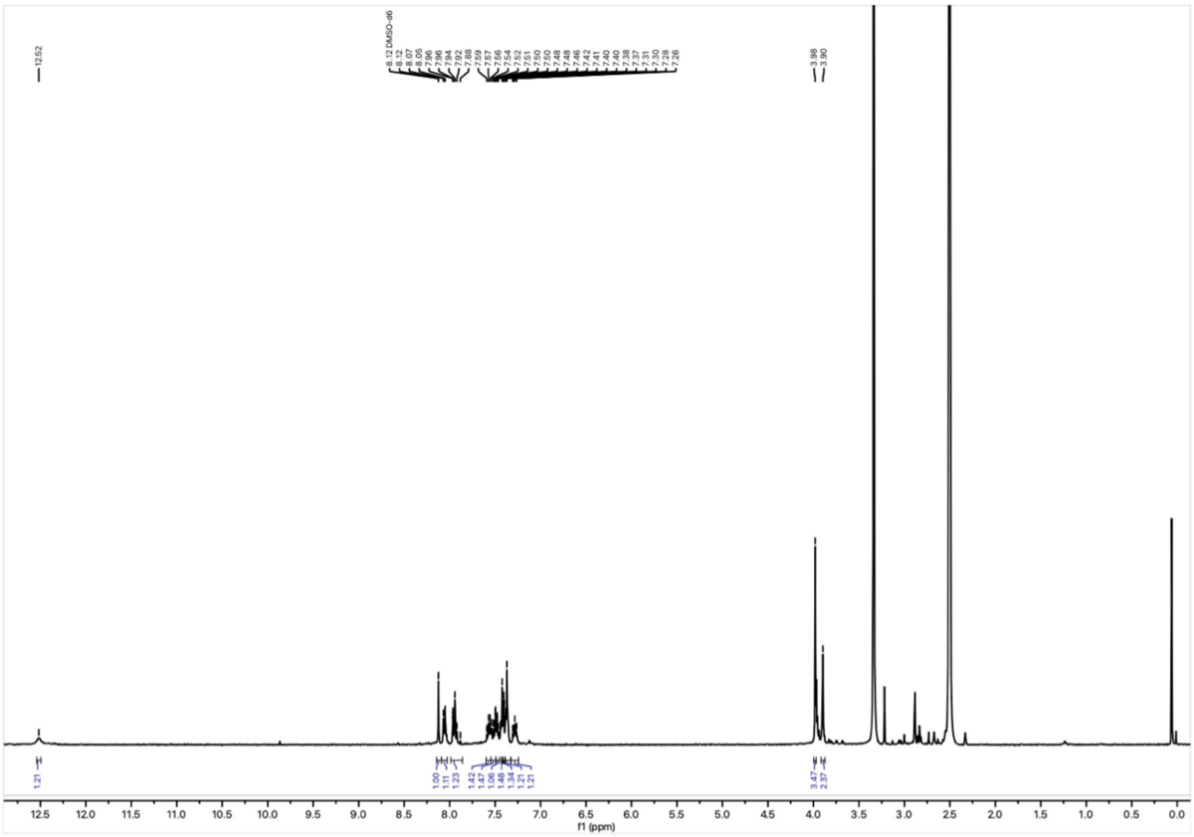

$^{1}H$

NMR spectrum of **29** in DMSO-$d_6$ at 400 MHz.

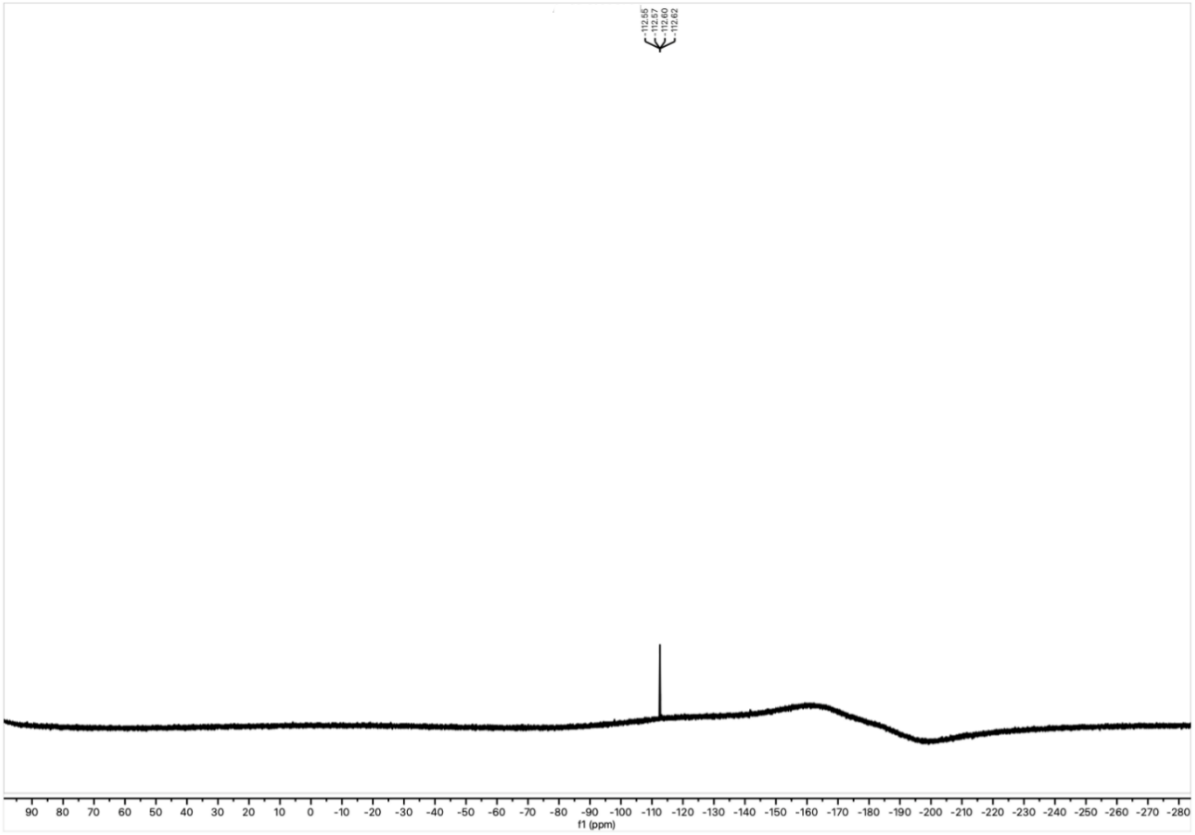

$^{19}F$

NMR spectrum of **29** in DMSO-$d_6$ at 377 MHz.

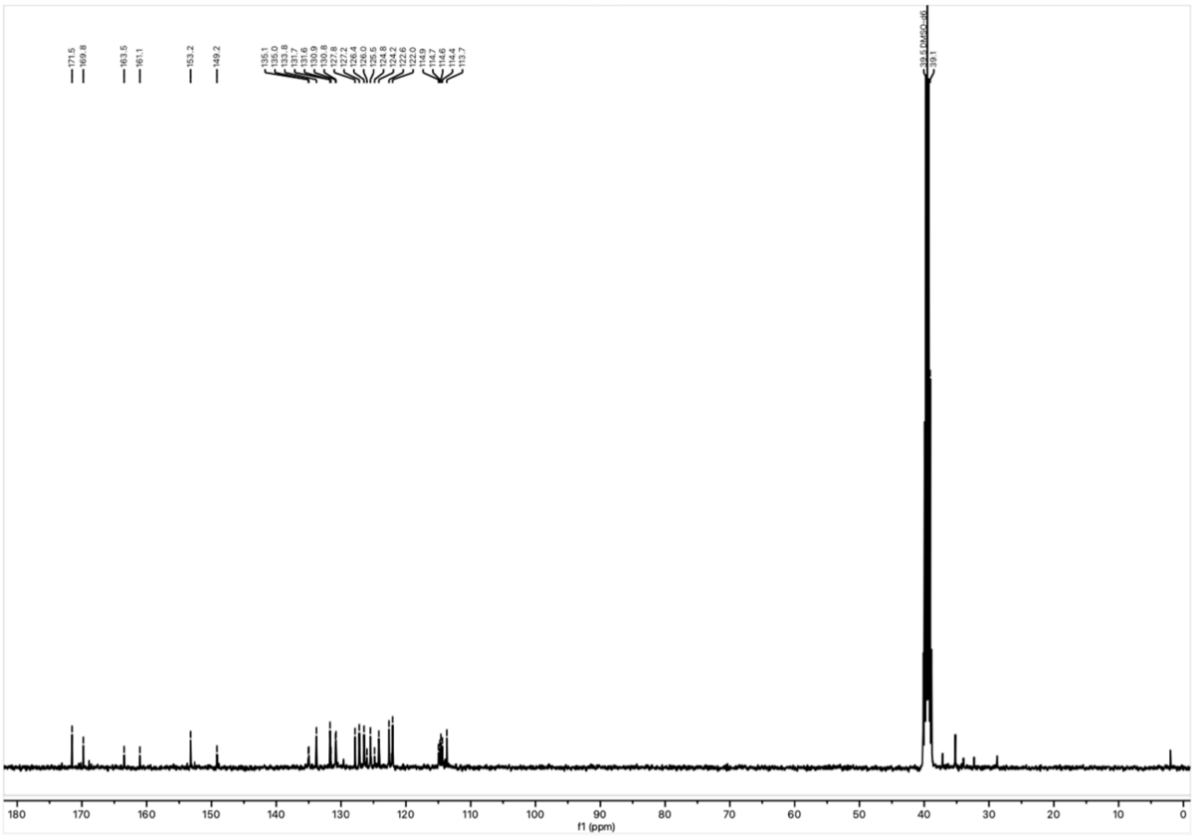


$^{13}C$ NMR spectrum of **29** in DMSO-$d_6$ at 101 MHz.

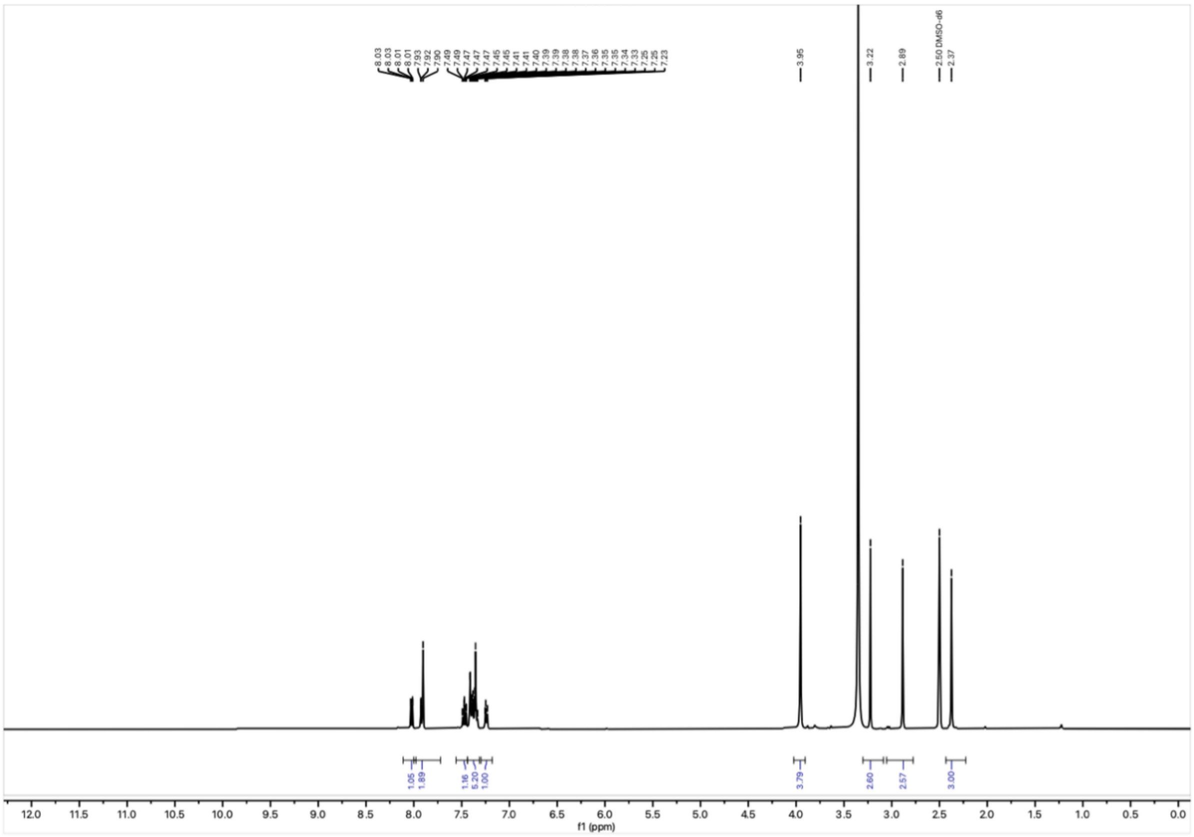


$^1H$ NMR spectrum of **30** in DMSO-$d_6$ at 400 MHz.

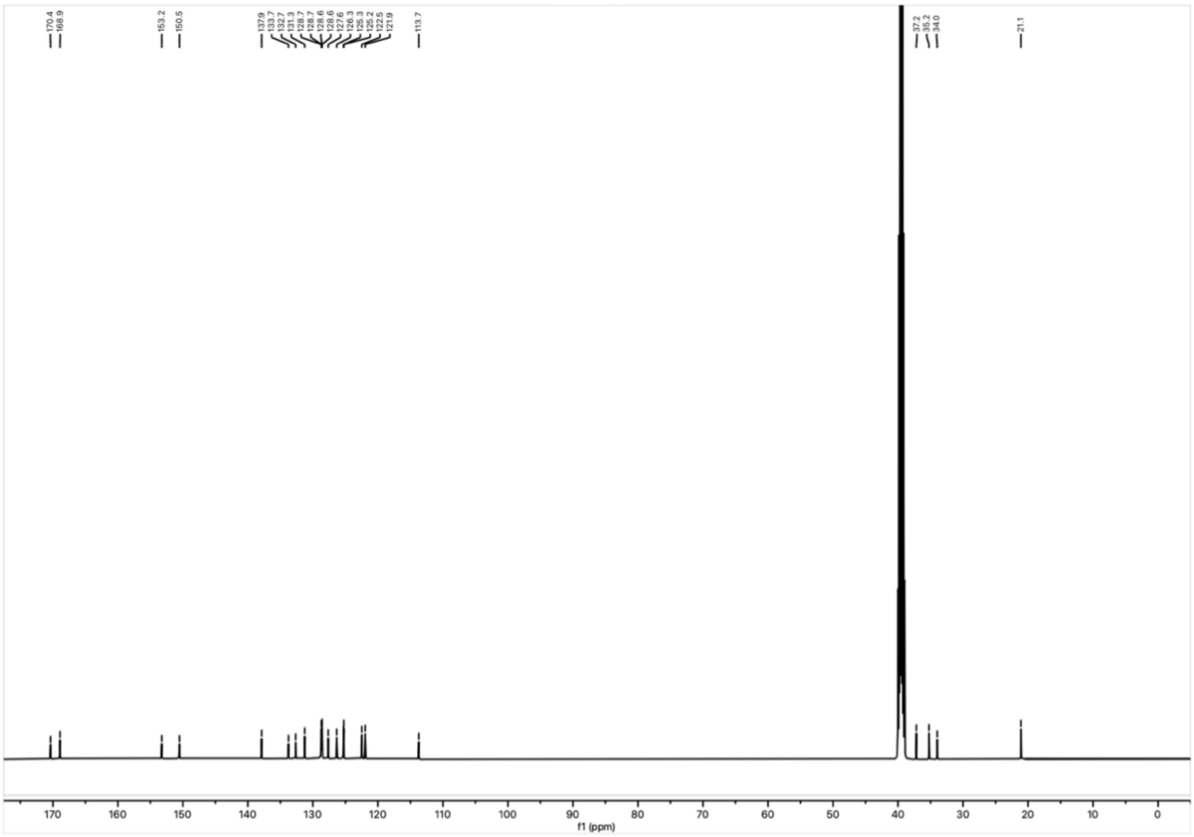

$^{13}$C NMR spectrum of **30** in DMSO-$d_6$ at 125 MHz.

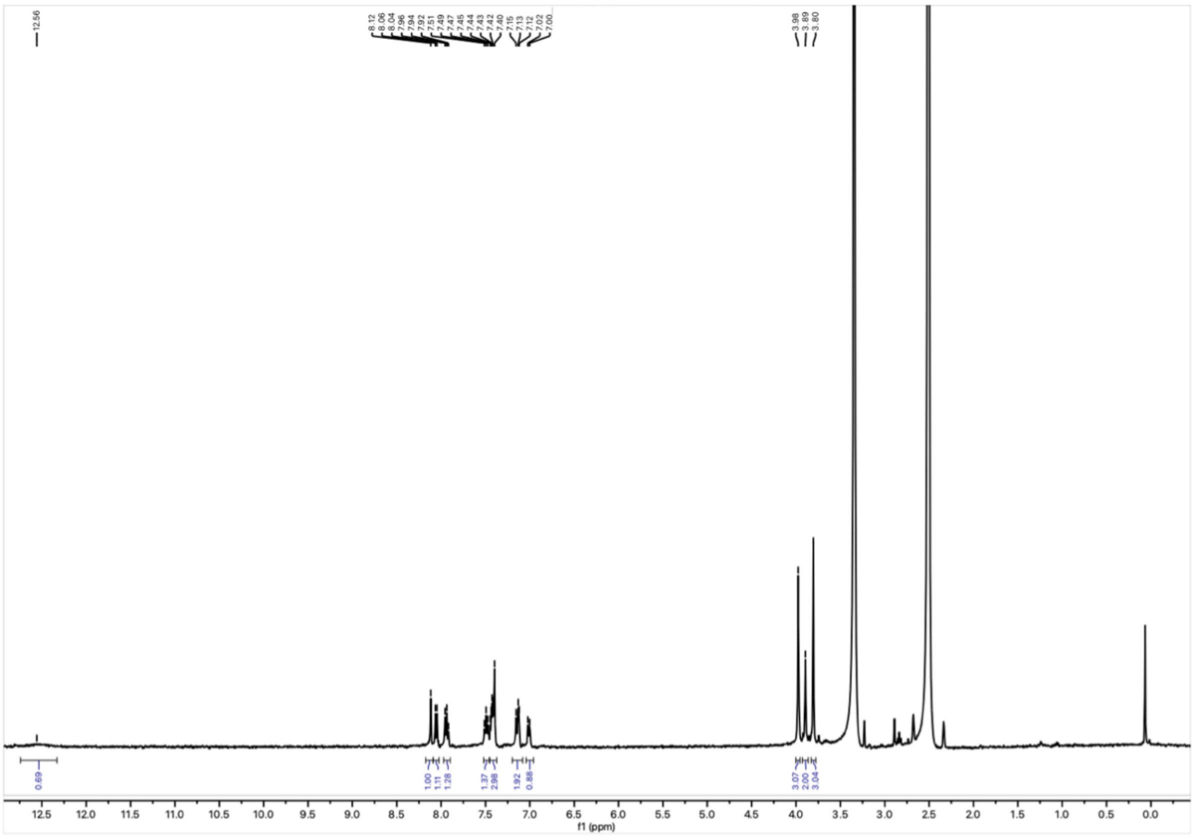

$^{1}$H NMR spectrum of **31** in DMSO-$d_6$ at 400 MHz.

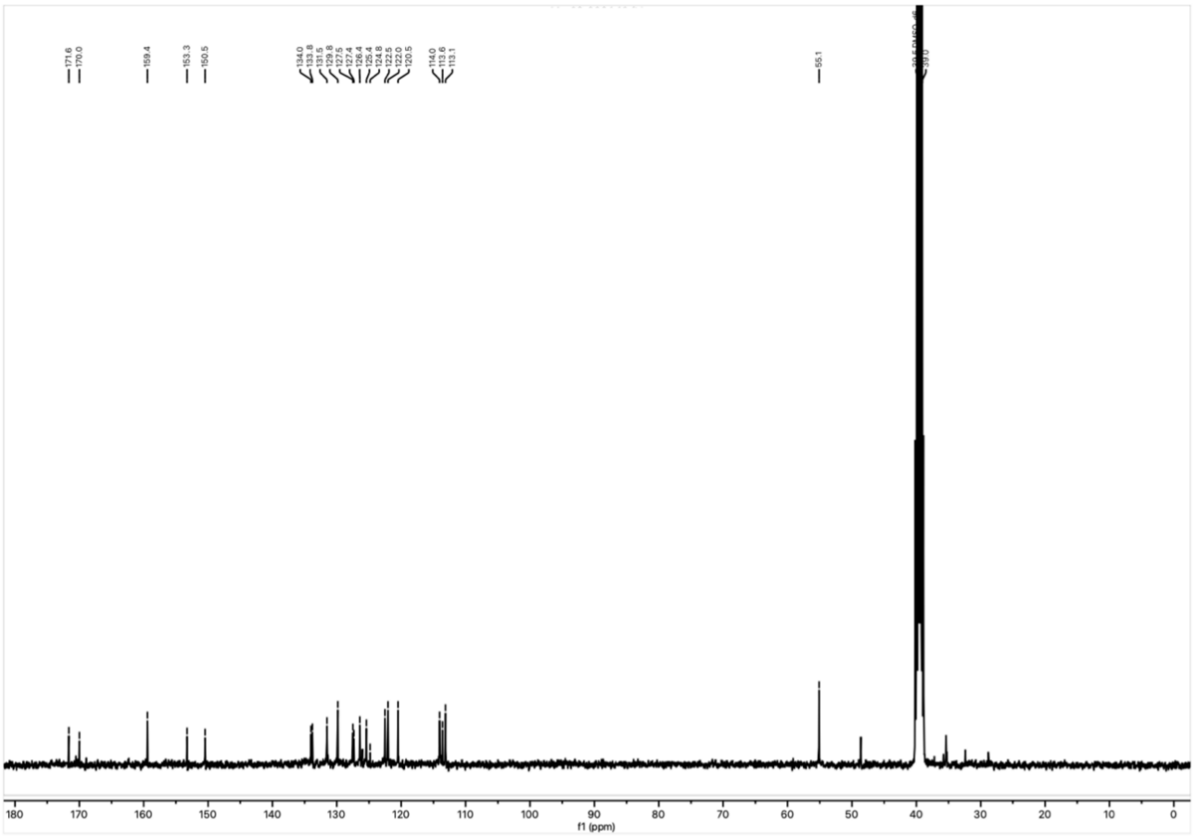

$^{13}C$ NMR spectrum of **31** in DMSO-$d_6$ at 101 MHz.

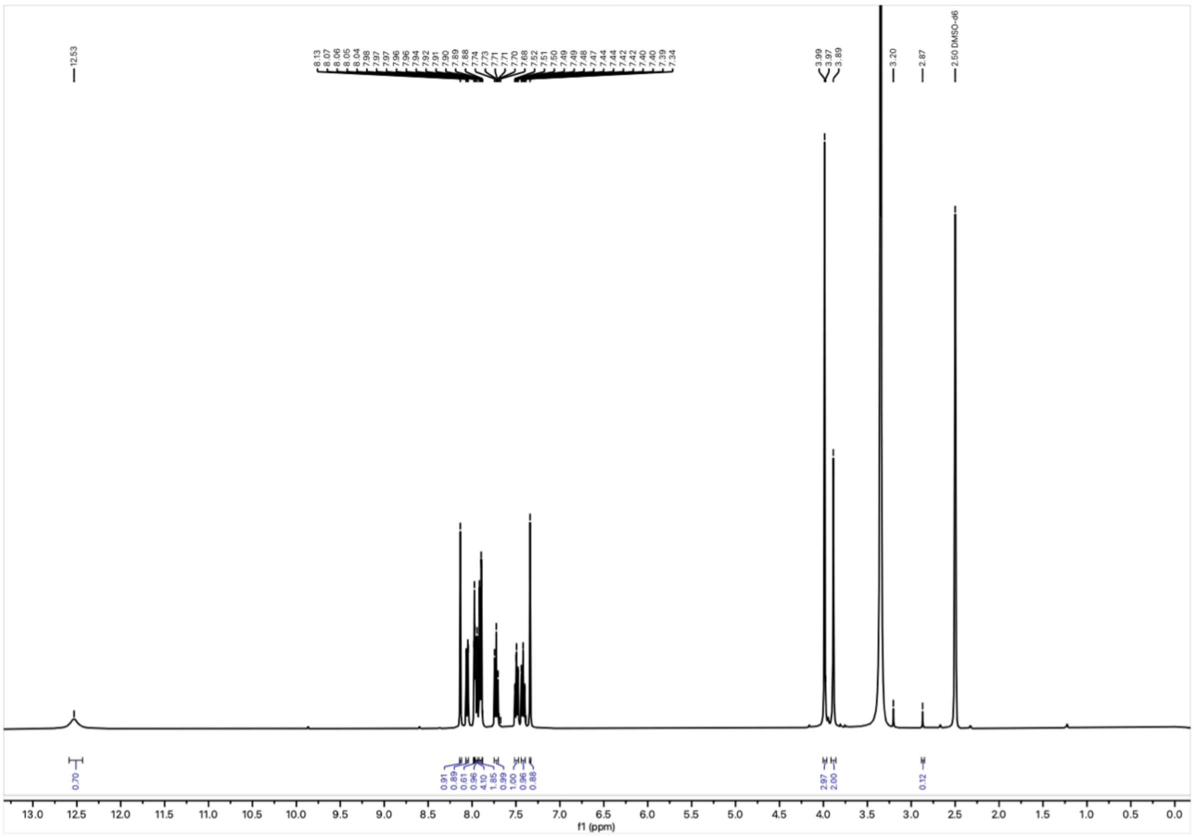

$^{1}H$ NMR spectrum of **32** in DMSO-$d_6$ at 400 MHz.

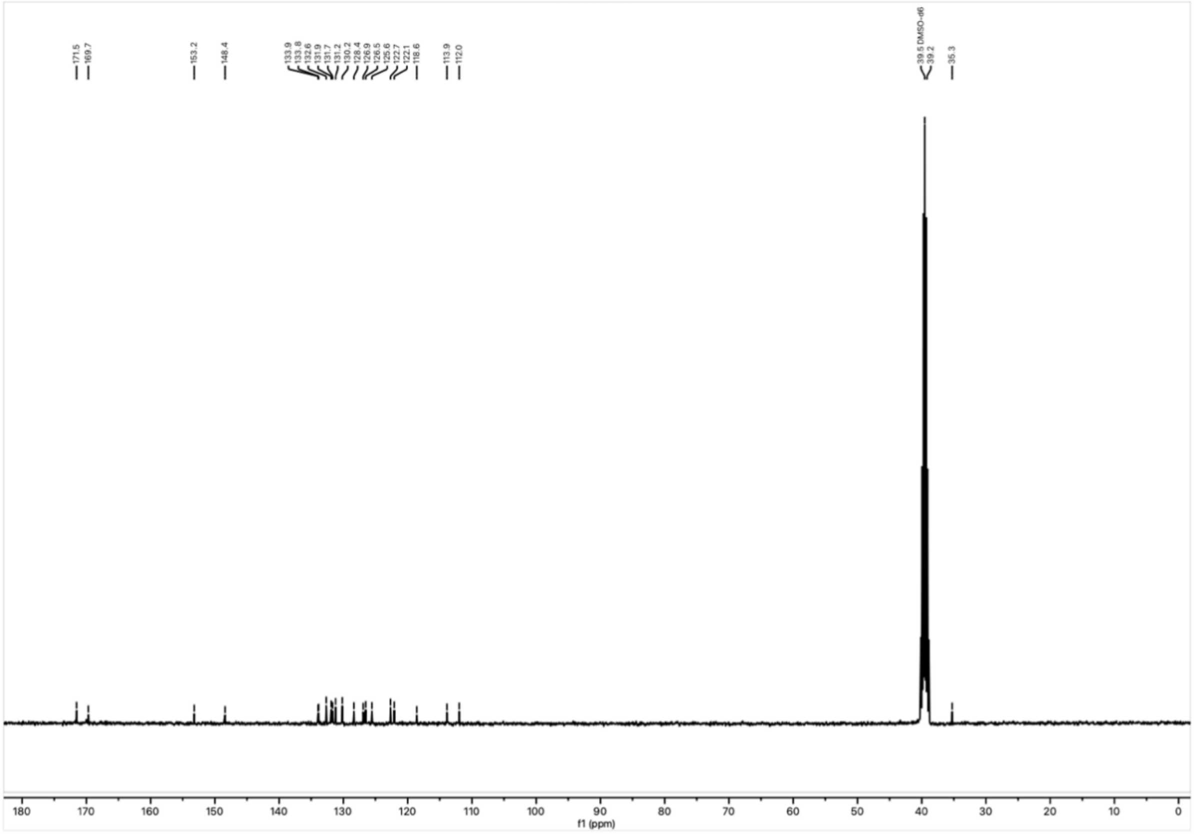

$^{13}C$ NMR spectrum of **32** in DMSO-$d_6$ at 101 MHz.

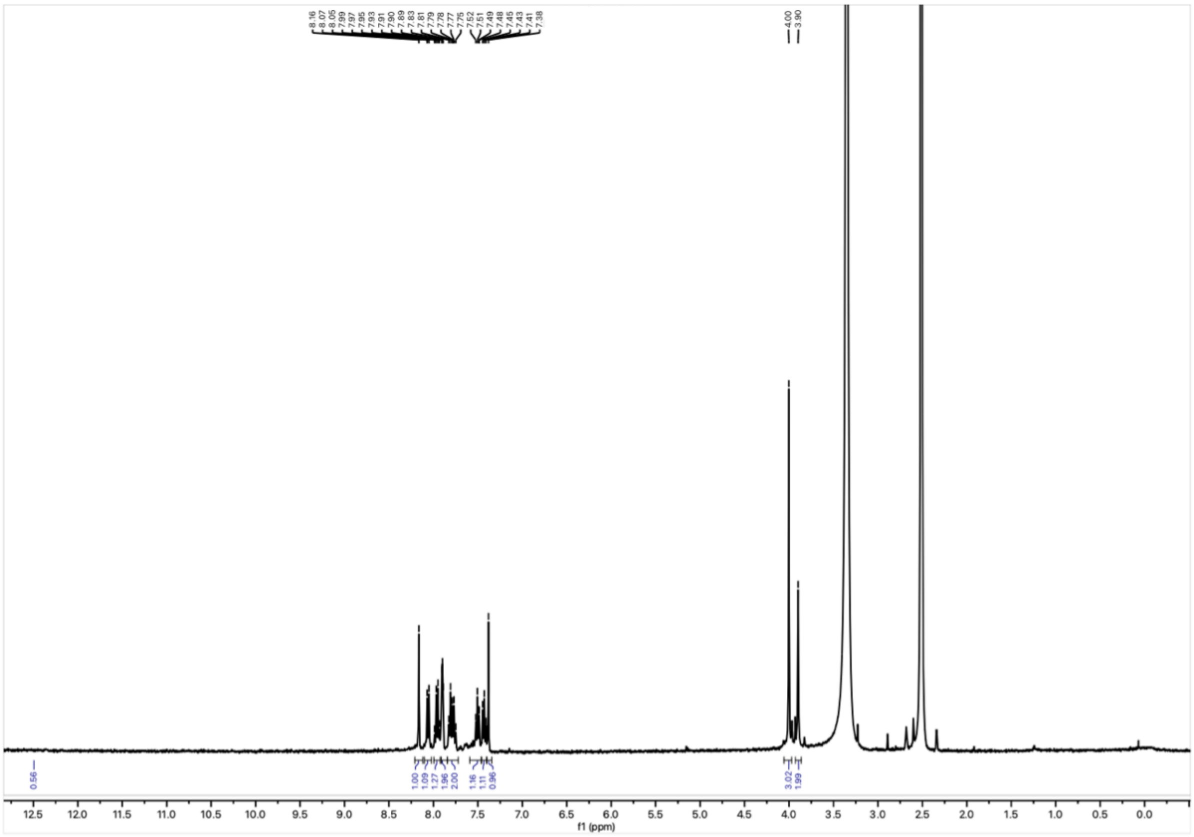

$^{1}H$ NMR spectrum of **33** in DMSO-$d_6$ at 400 MHz.

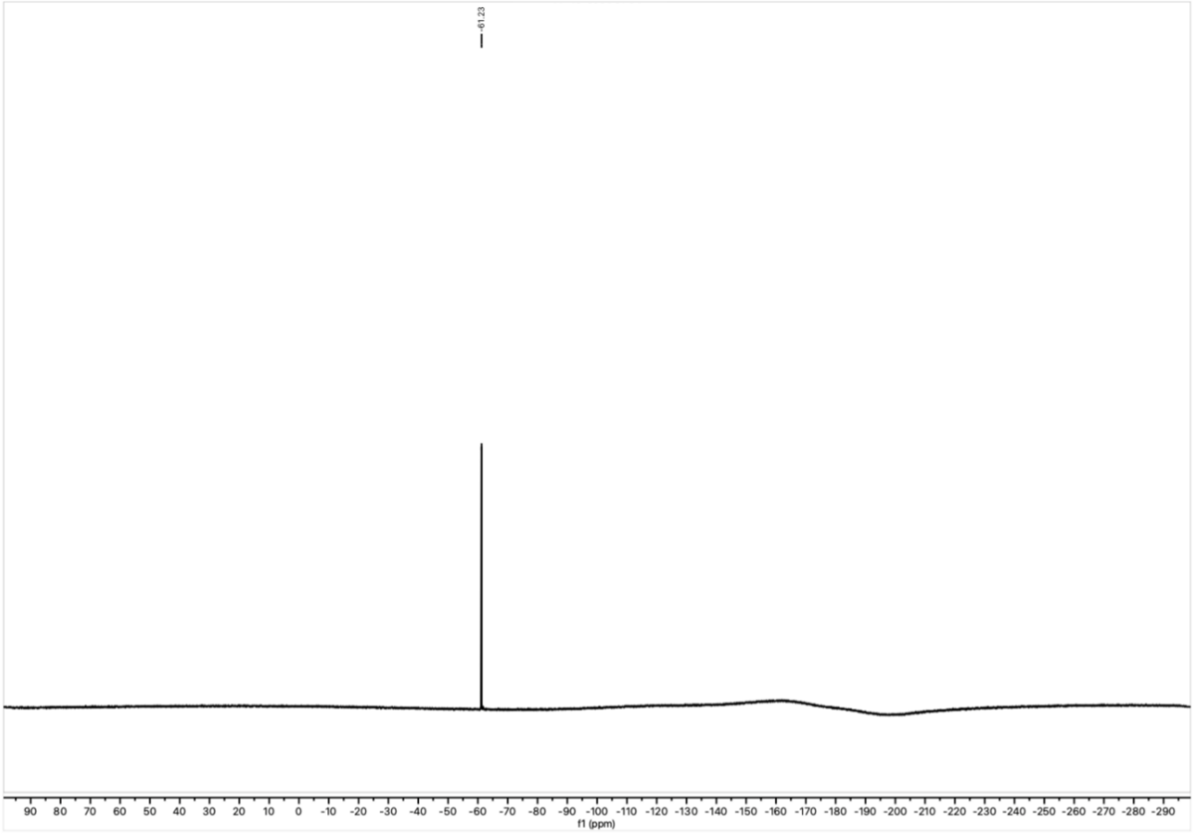


$^{19}F$ NMR spectrum of **33** in DMSO-$d_6$ at 377 MHz.

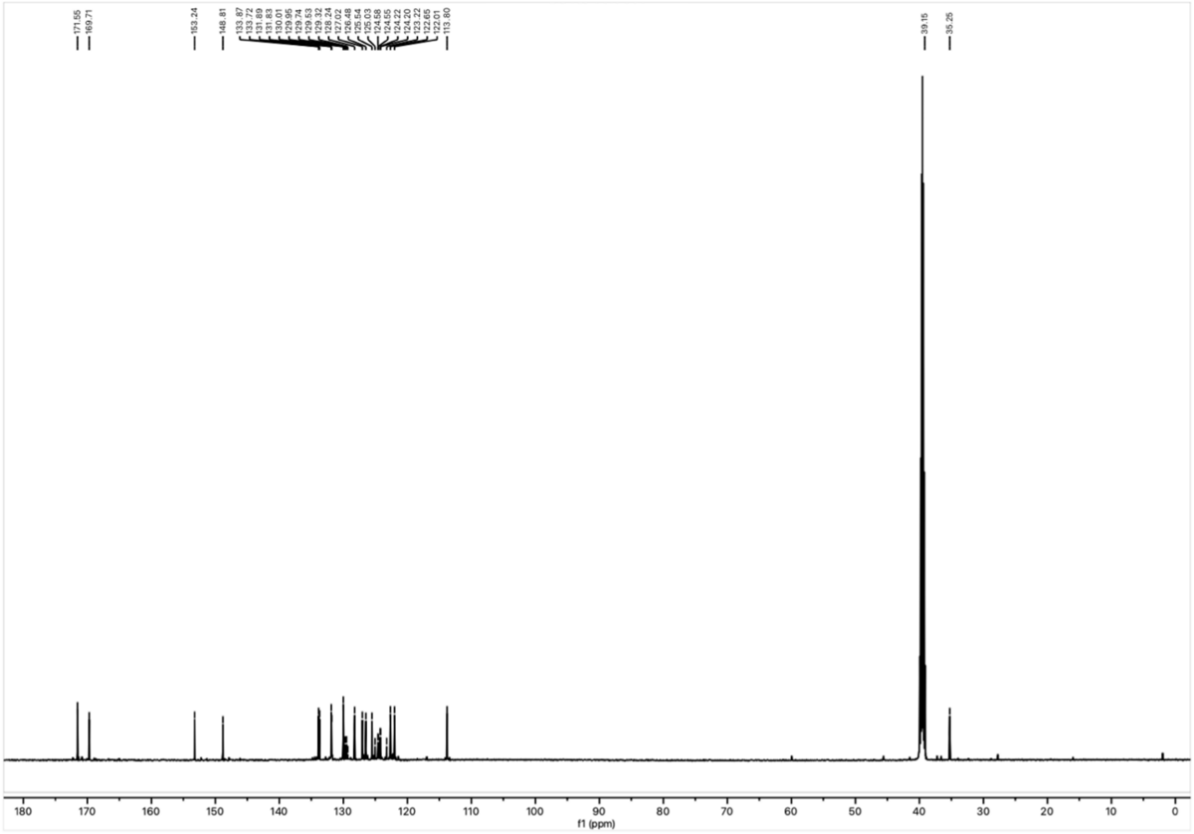


$^{13}C$ NMR spectrum of **33** in DMSO-$d_6$ at 101 MHz.

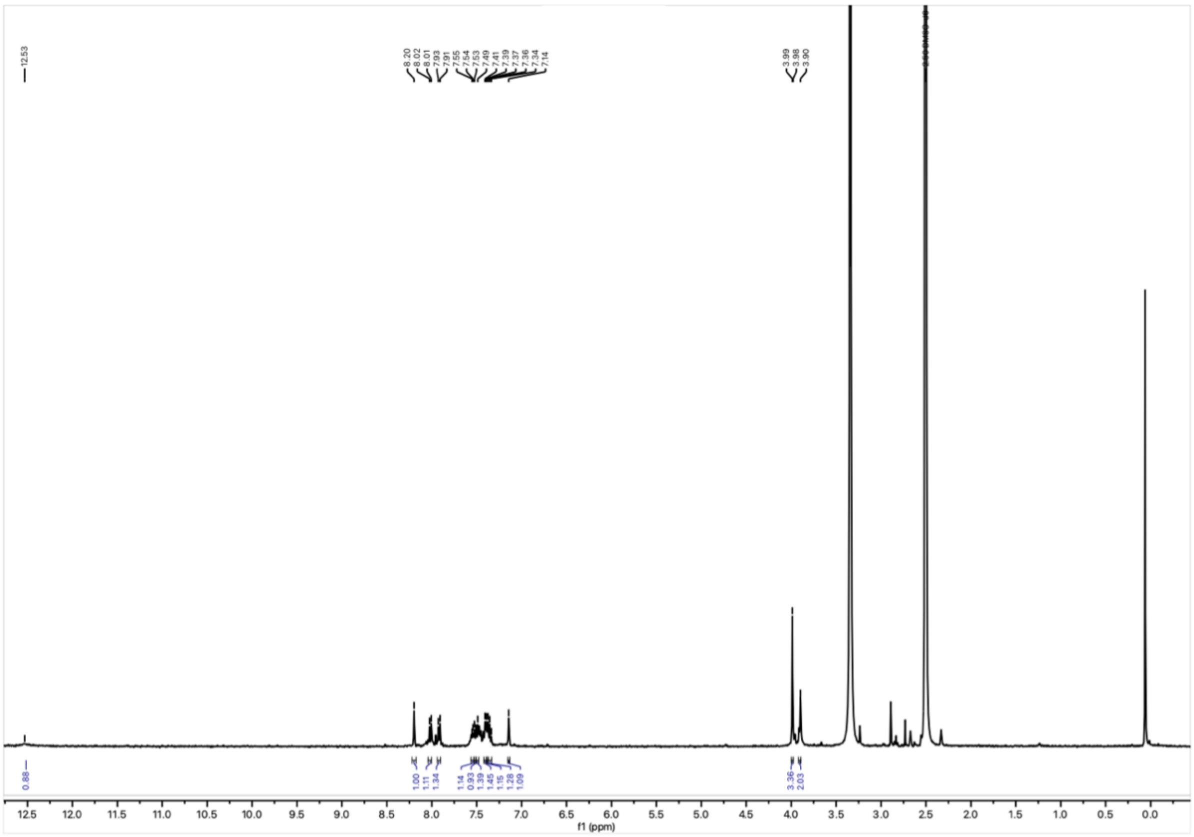

$^1H$

NMR spectrum of **34** in DMSO-$d_6$ at 400 MHz.

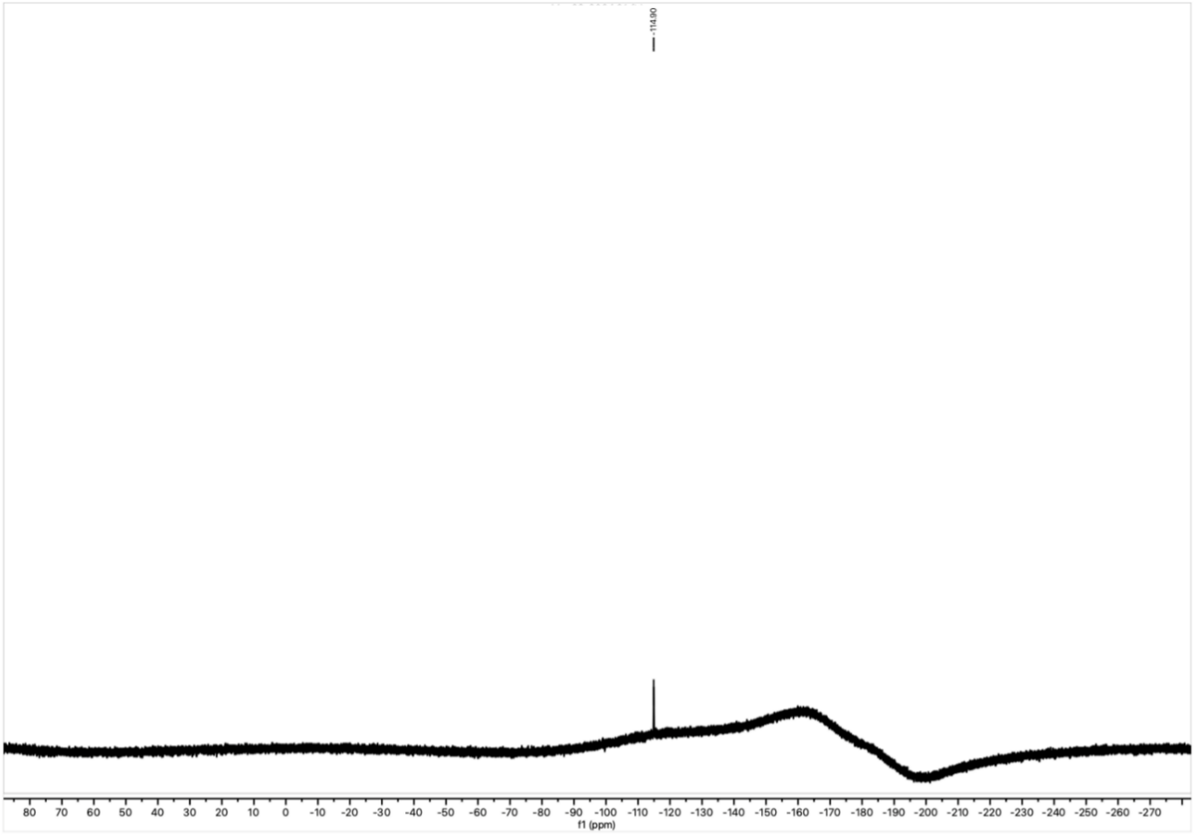

$^{19}F$

NMR spectrum of **34** in DMSO-$d_6$ at 377 MHz.

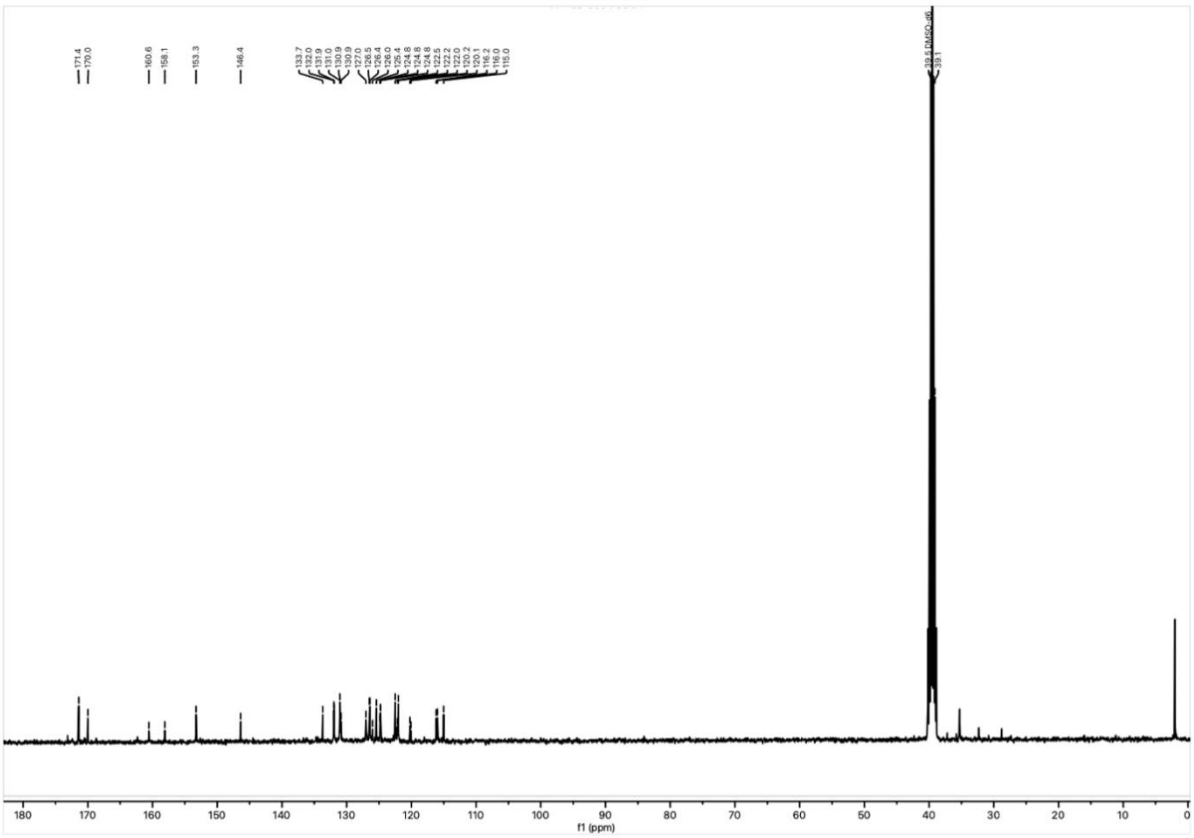


$^{13}C$ NMR spectrum of **34** in DMSO-$d_6$ at 101 MHz.

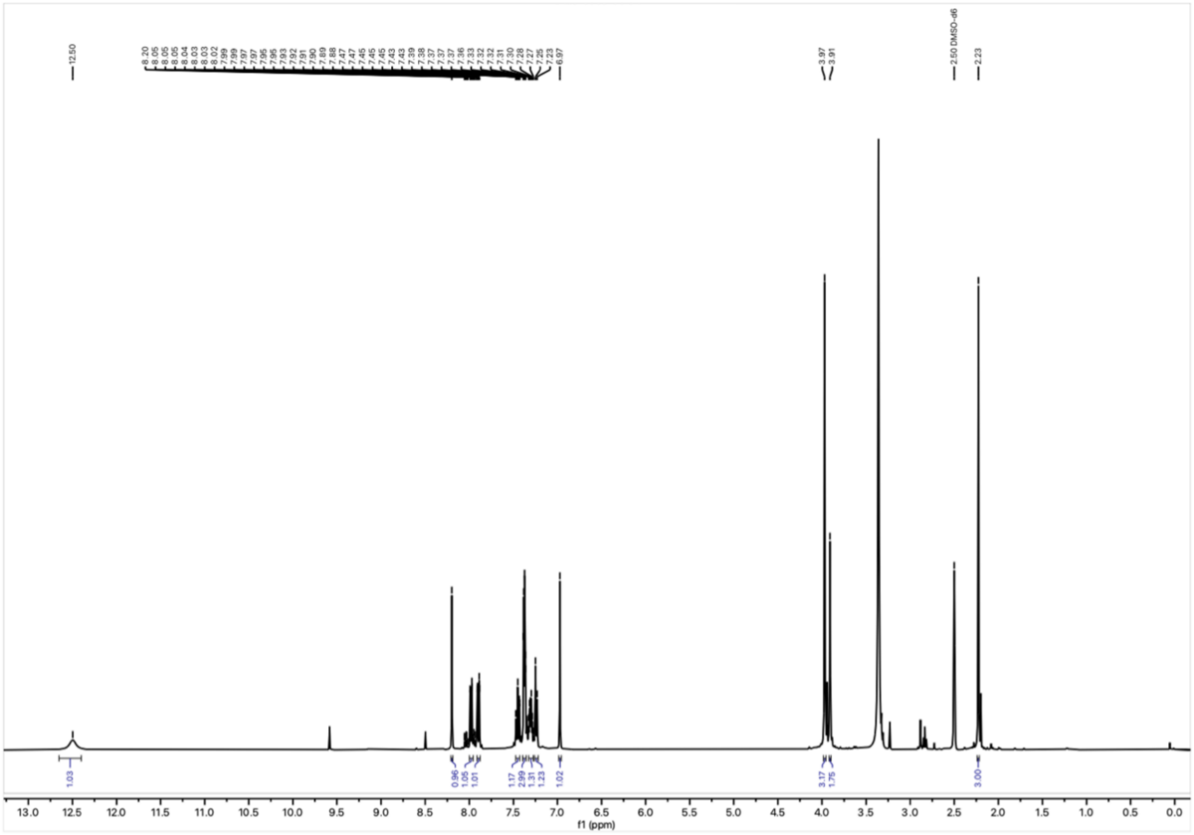


$^{1}H$ NMR spectrum of **35** in DMSO-$d_6$ at 400 MHz.

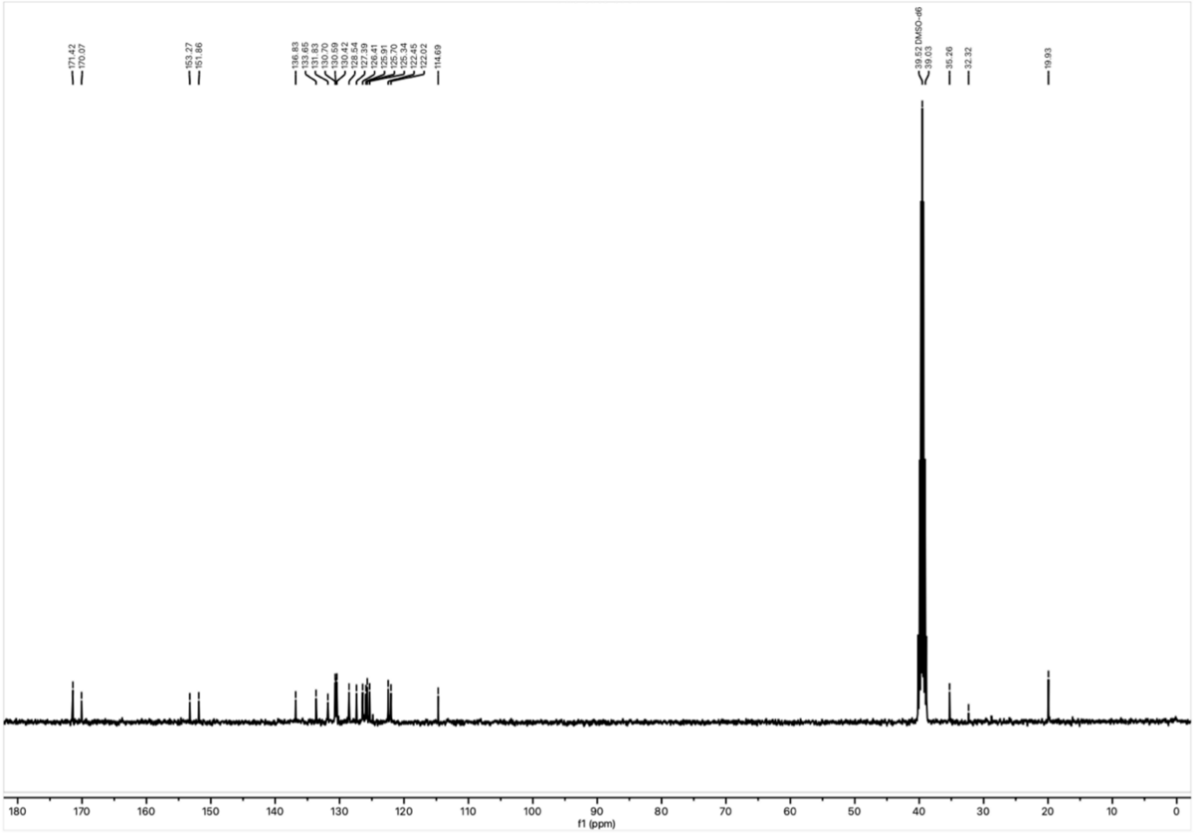


$^{13}C$ NMR spectrum of **35** in DMSO-$d_6$ at 101 MHz.

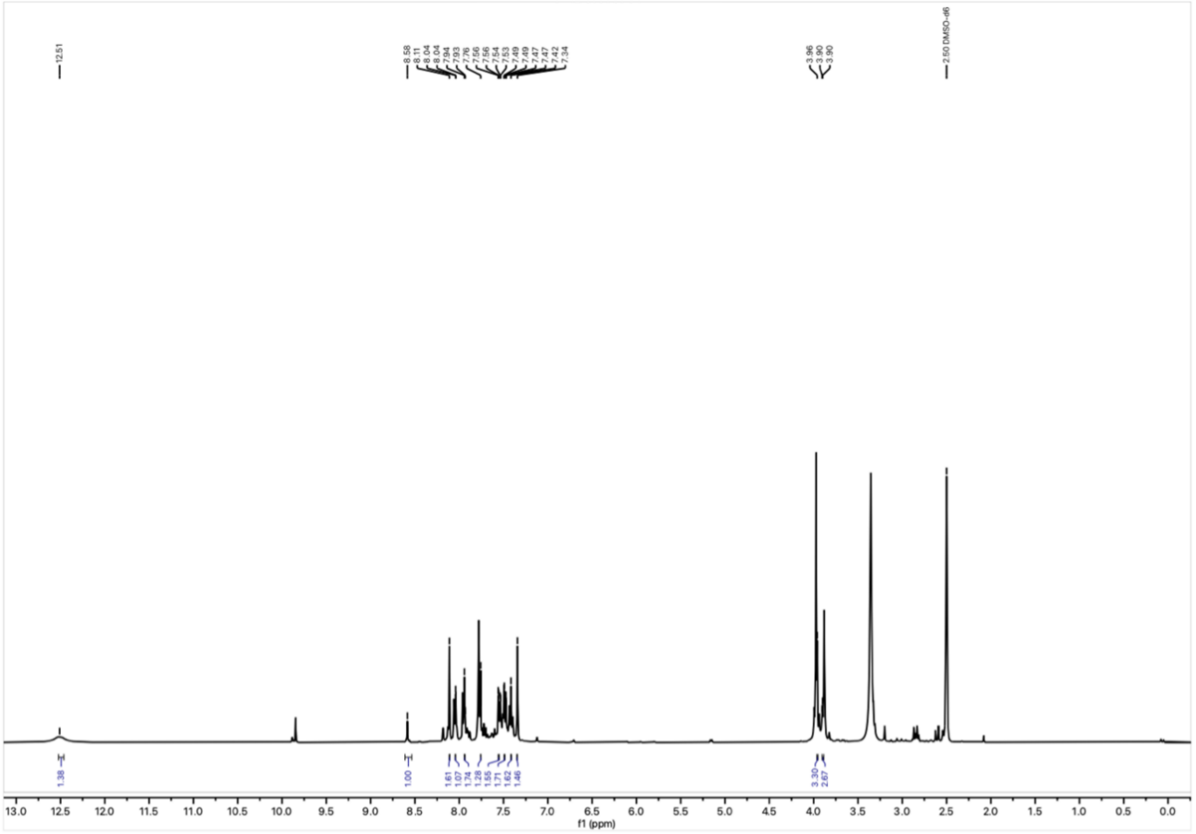


$^{1}H$ NMR spectrum of **36** in DMSO-$d_6$ at 400 MHz.

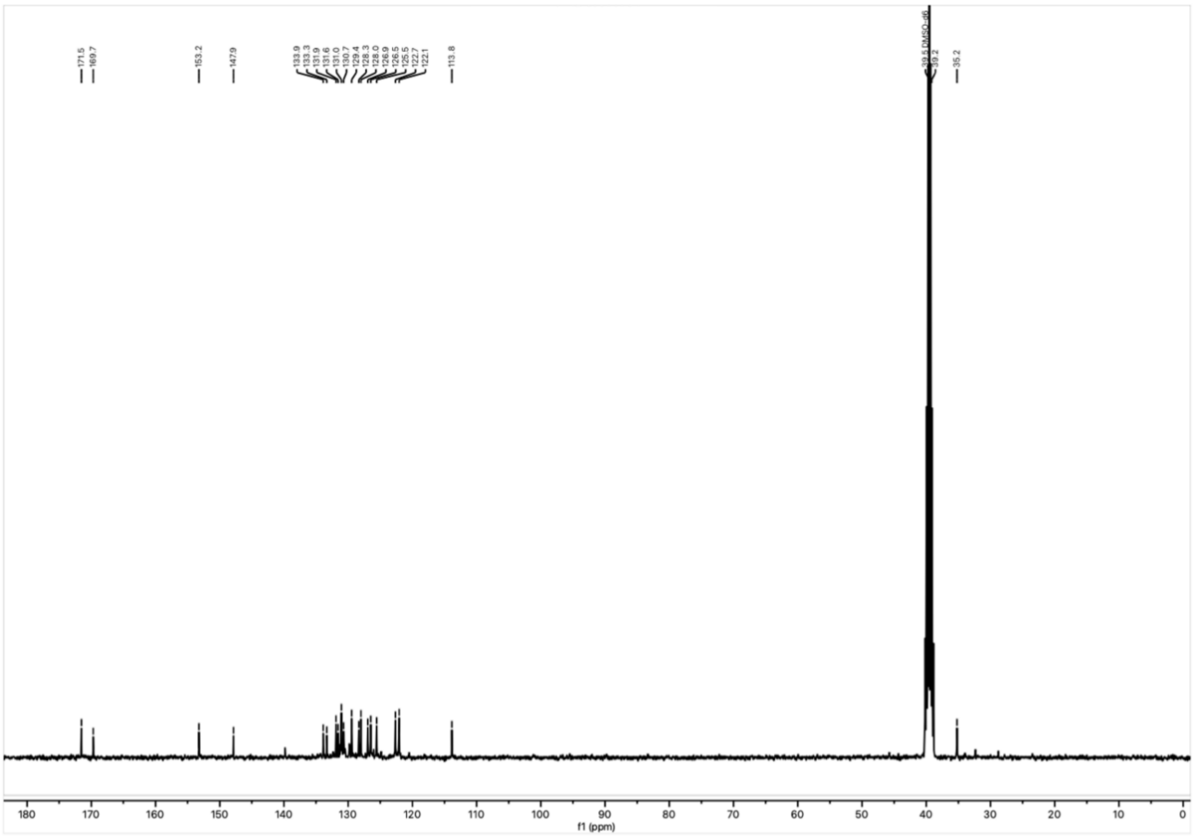


$^{13}C$

NMR spectrum of **36** in DMSO-$d_6$ at 101 MHz.

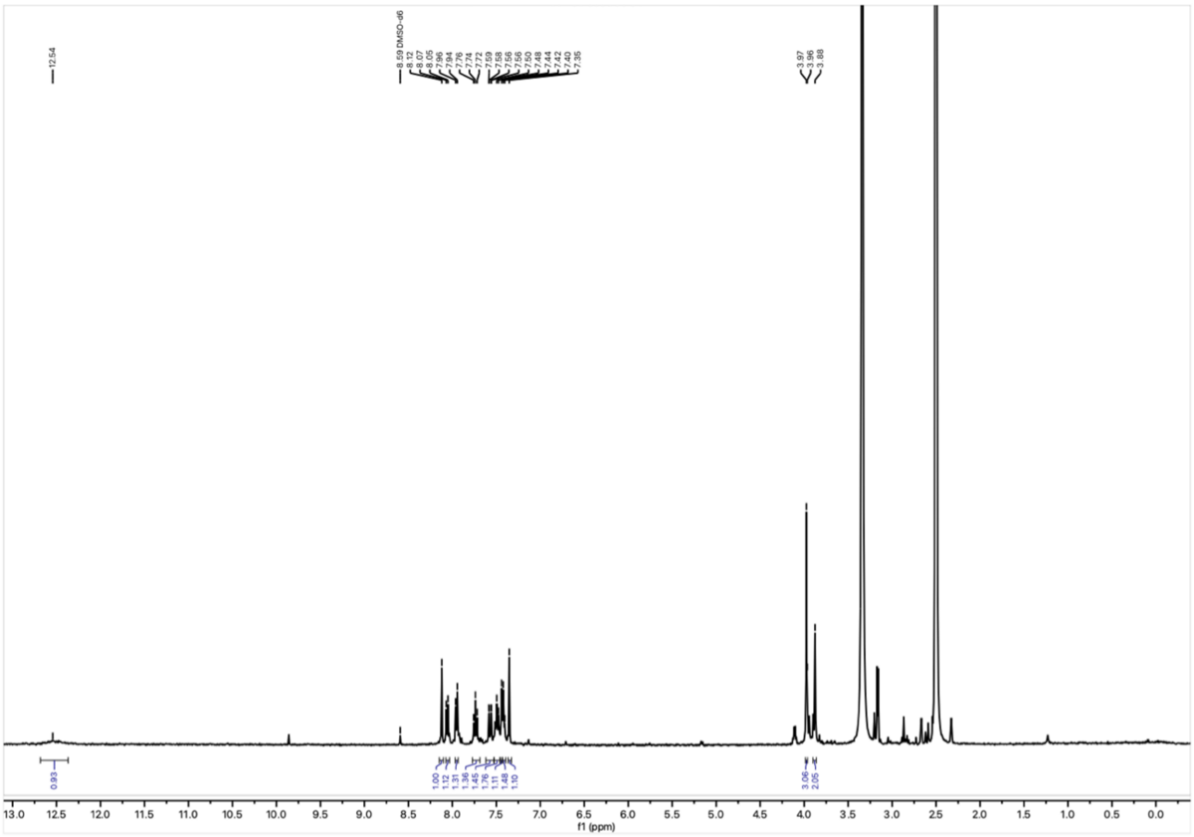


$^{1}H$

NMR spectrum of **37** in DMSO-$d_6$ at 400 MHz.

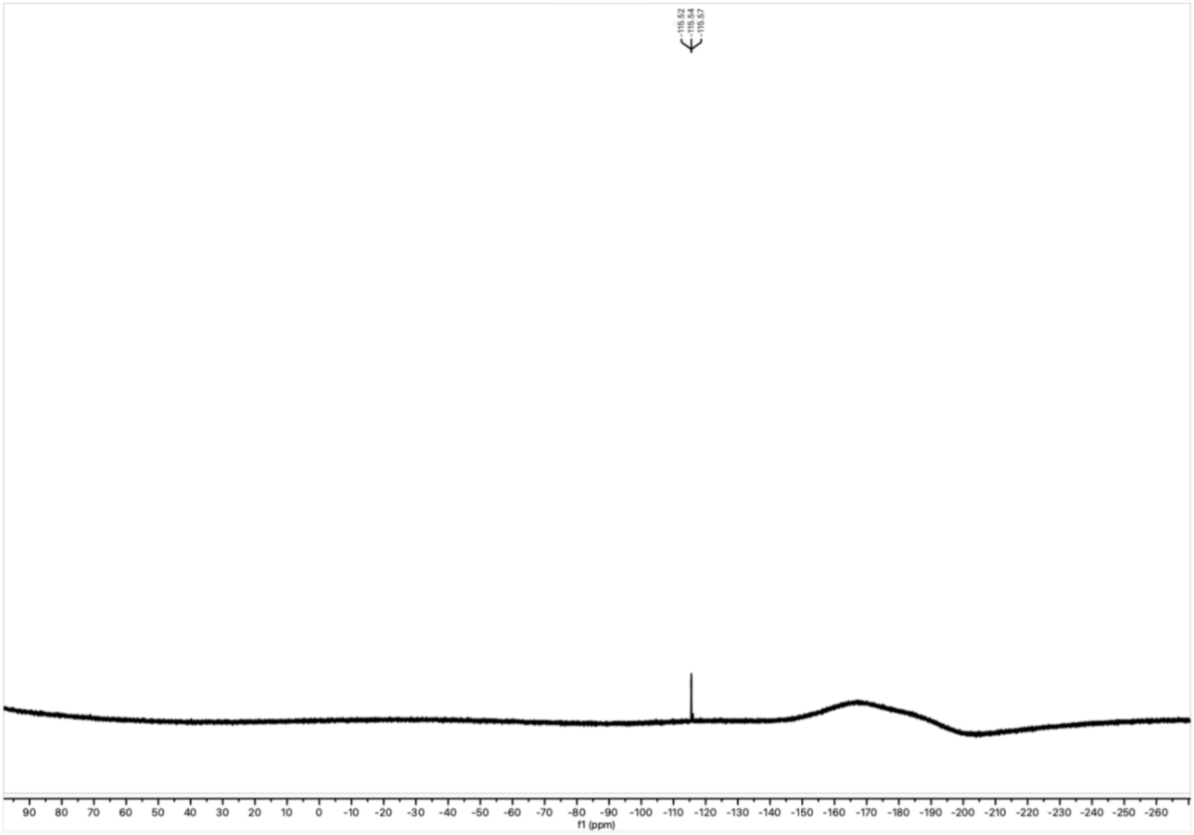

$^{19}F$ NMR spectrum of **37** in DMSO-$d_6$ at 377 MHz.

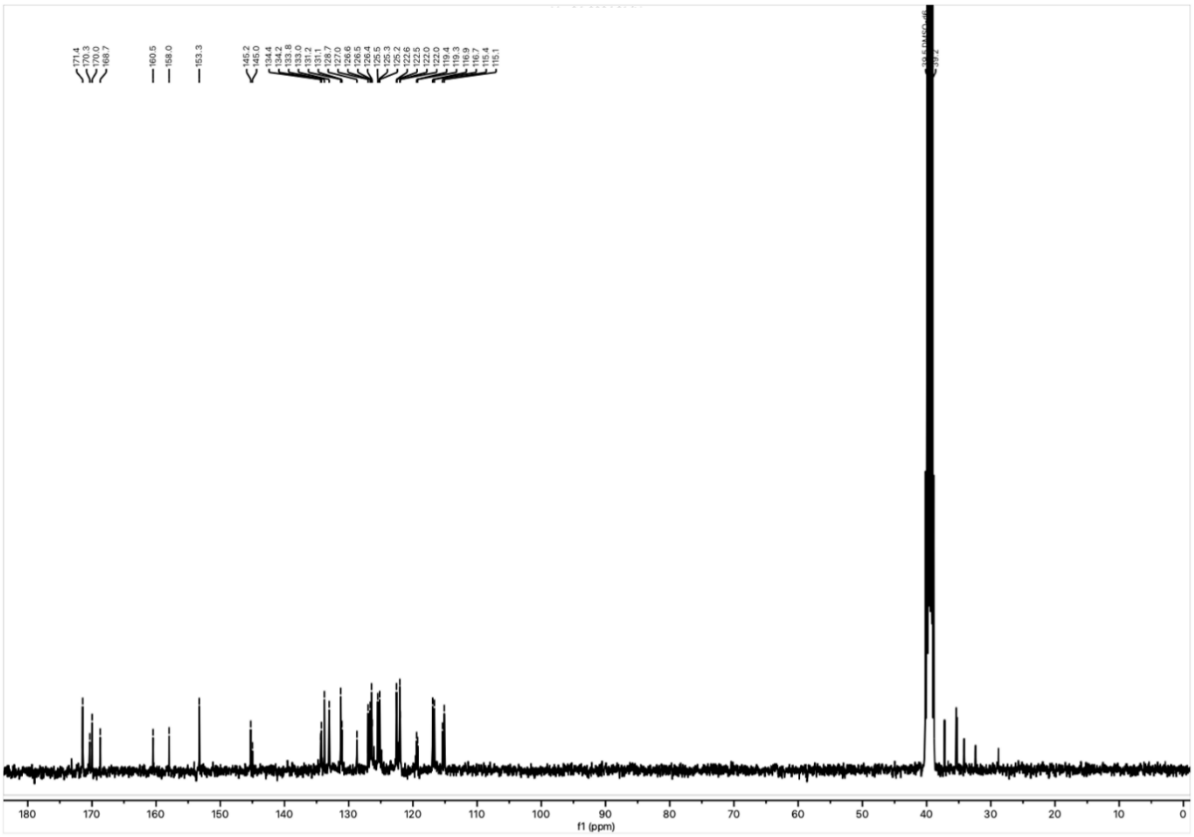

$^{13}C$ NMR spectrum of **37** in DMSO-$d_6$ at 101 MHz.

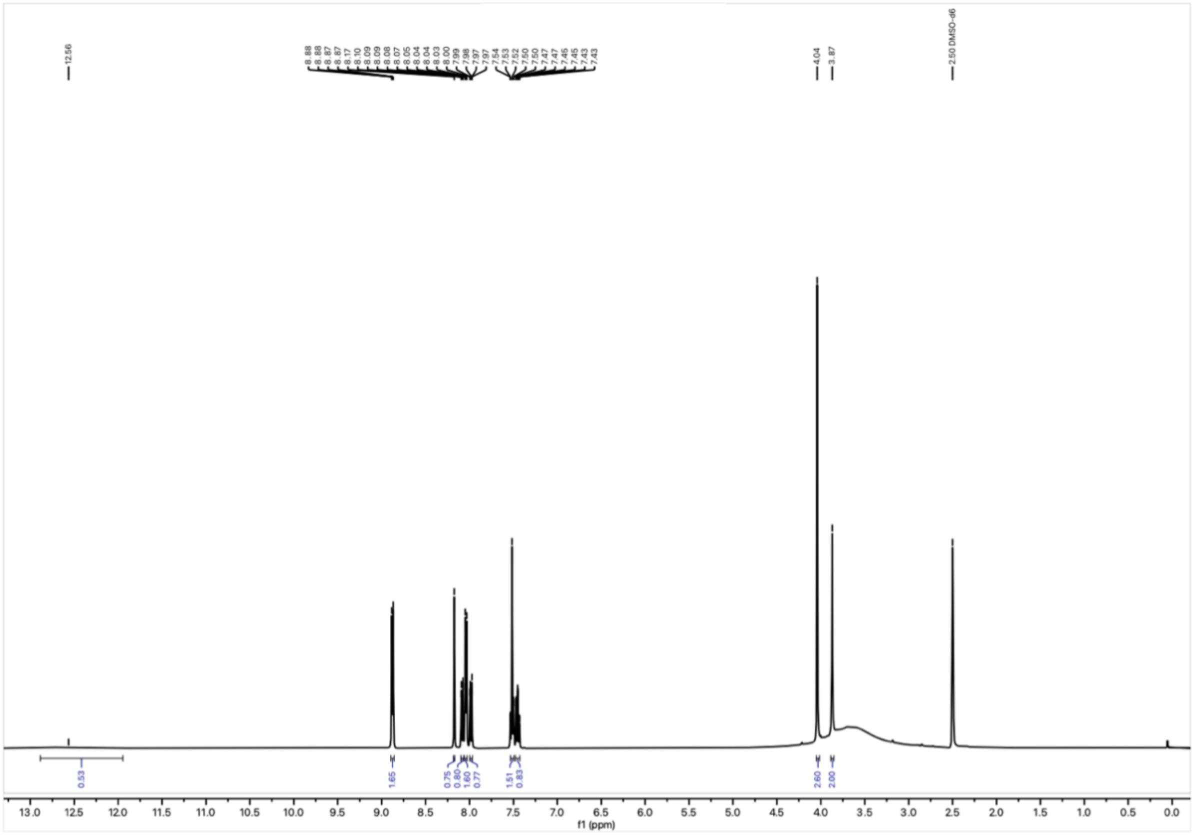

[1]H NMR spectrum of **38** in DMSO-$d_6$ at 400 MHz.

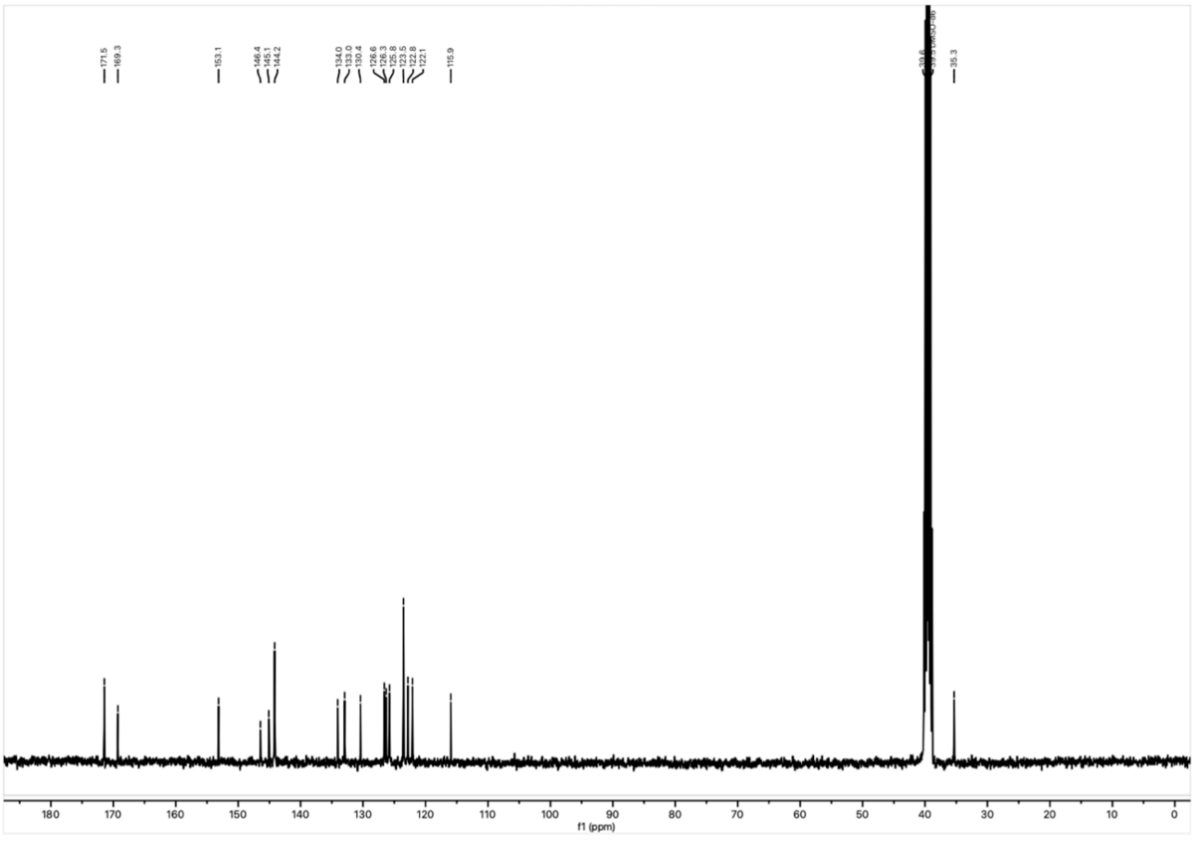

[13]C NMR spectrum of **38** in DMSO-$d_6$ at 101 MHz.

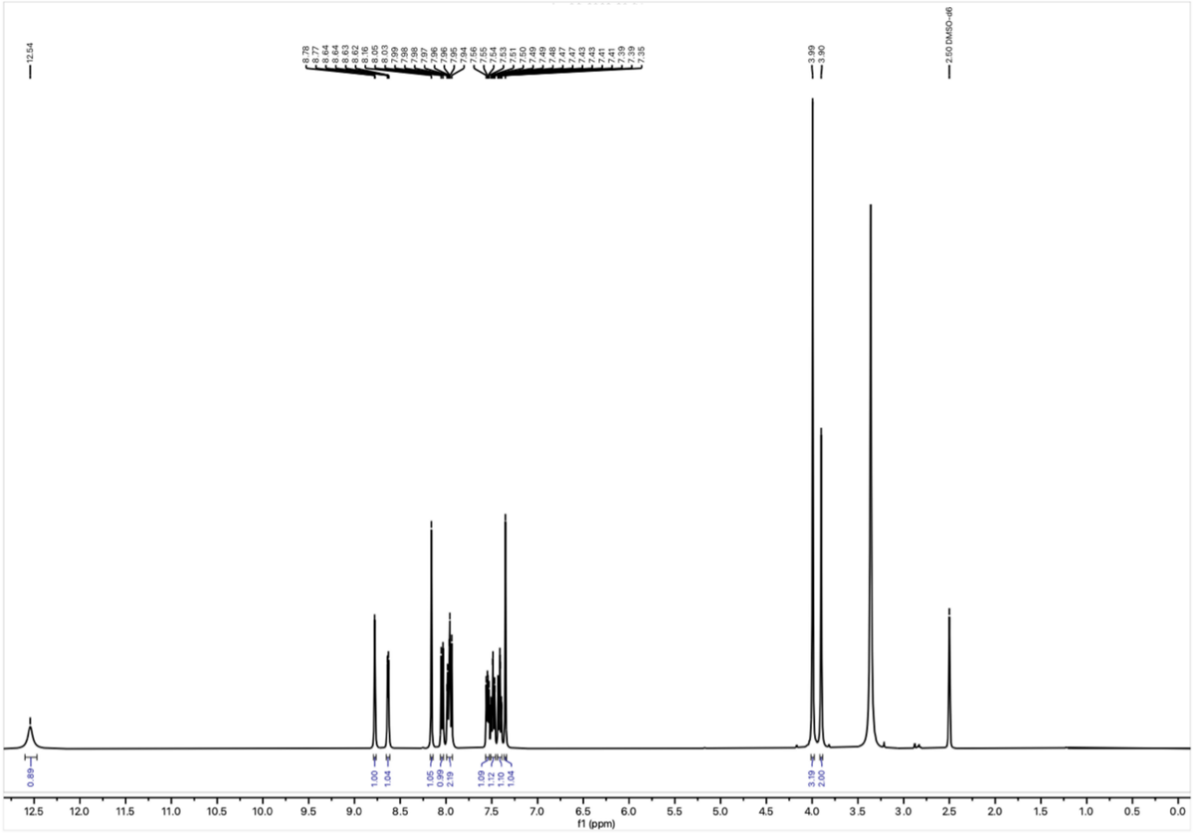

$^{1}H$ NMR spectrum of **39** in DMSO-$d_6$ at 400 MHz.

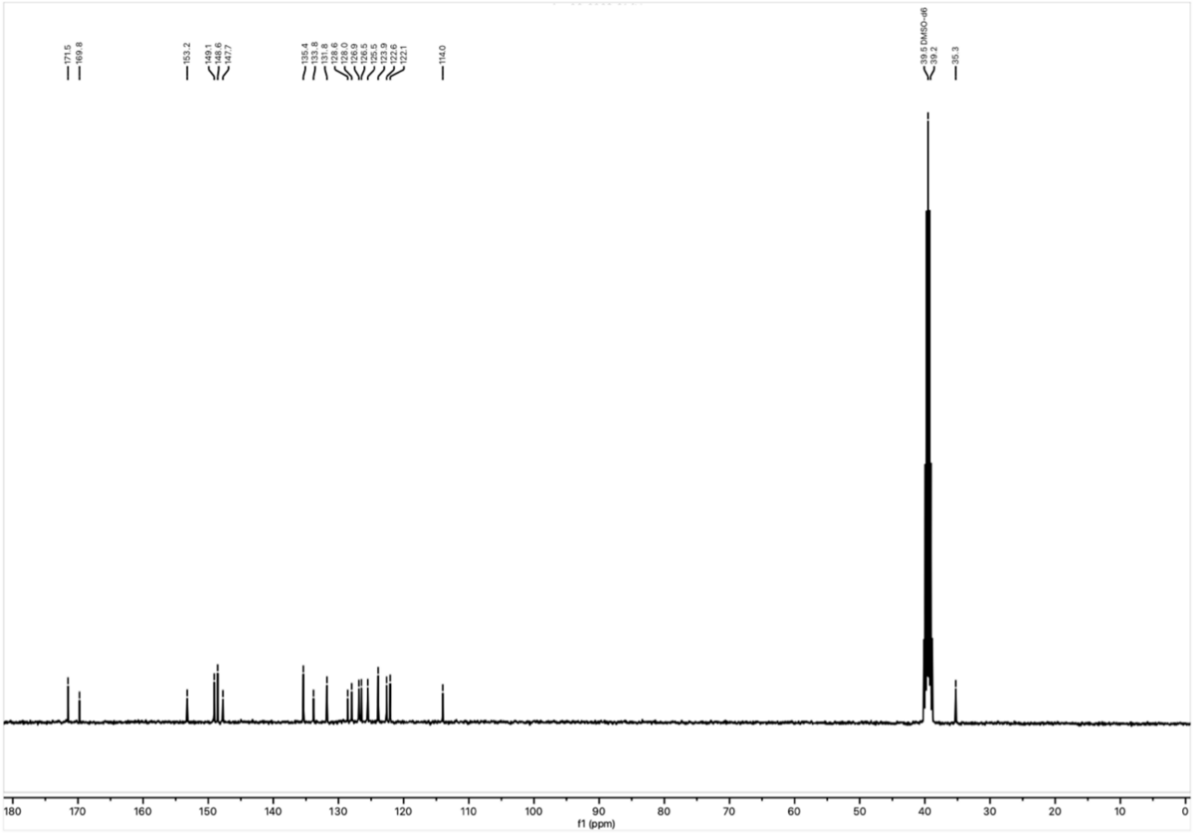

$^{13}C$ NMR spectrum of **39** in DMSO-$d_6$ at 101 MHz.

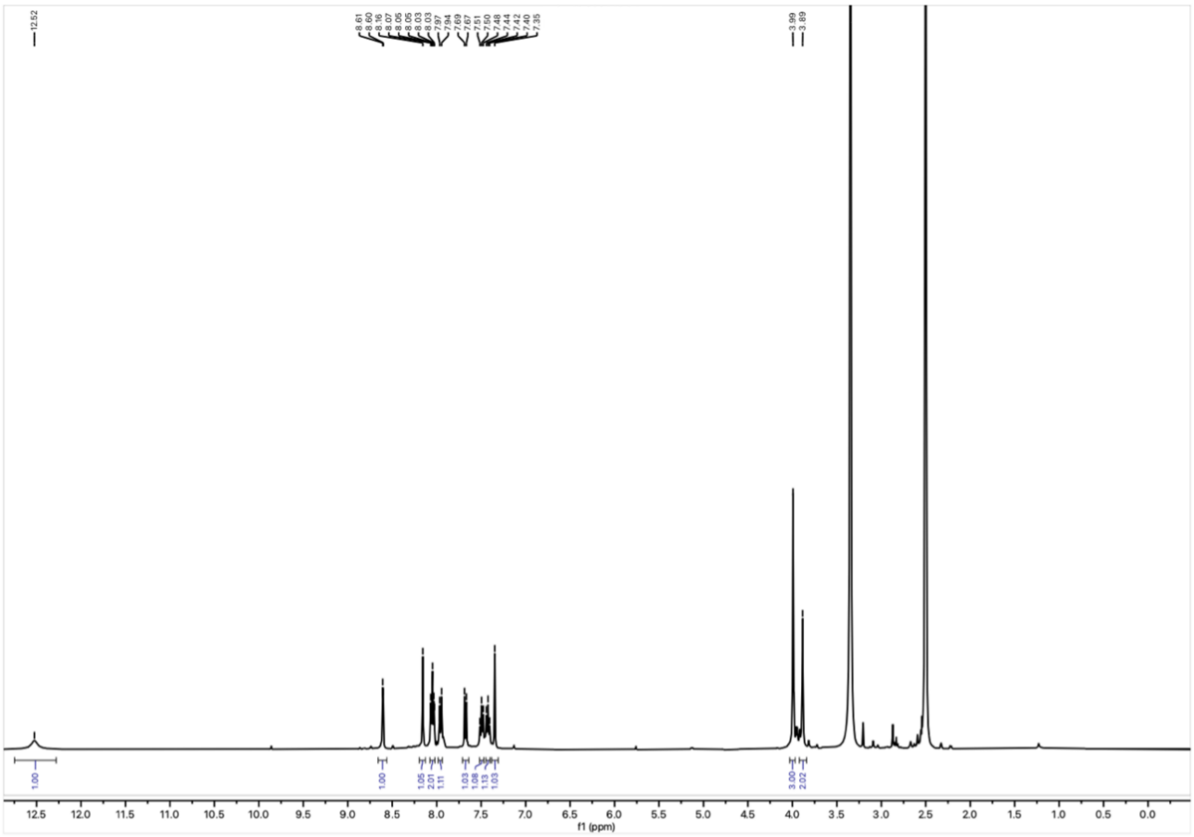

$^{1}H$ NMR spectrum of **40** in DMSO-$d_6$ at 400 MHz.

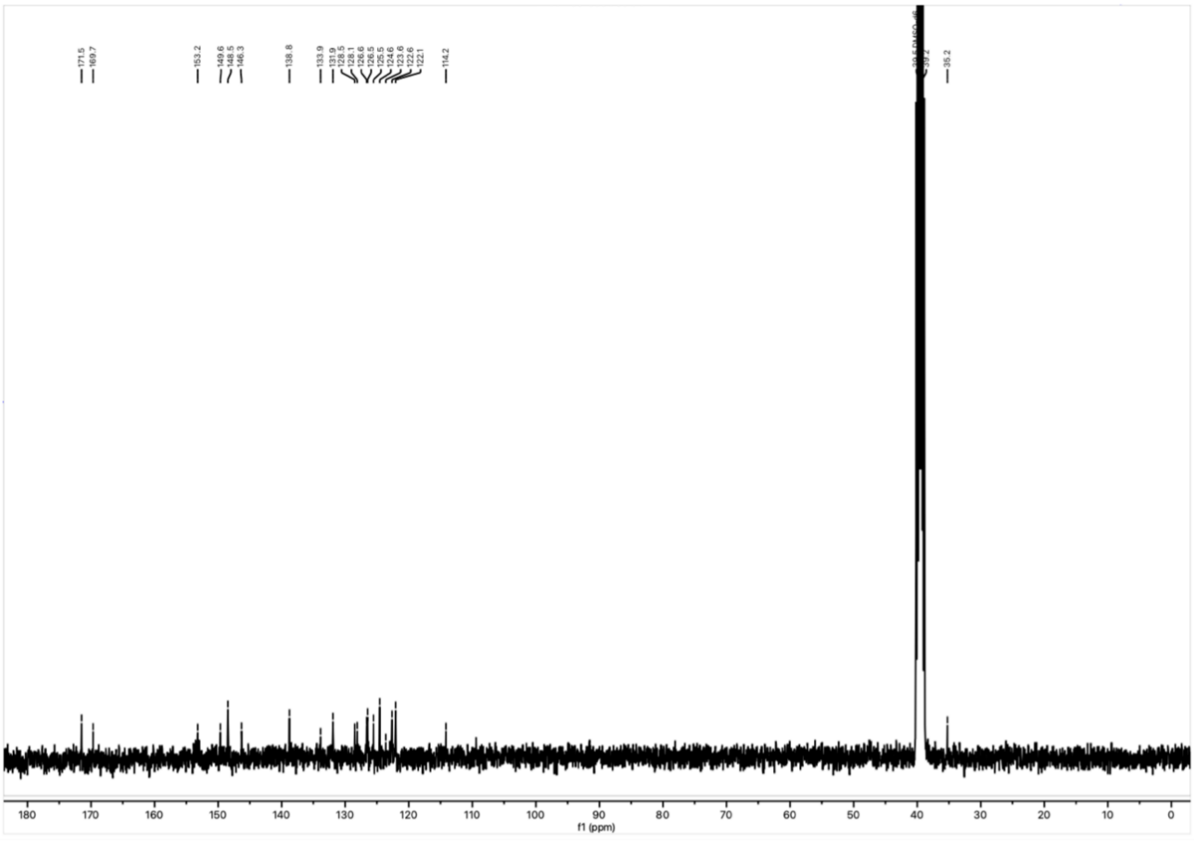

$^{13}C$ NMR spectrum of **40** in DMSO-$d_6$ at 101 MHz.

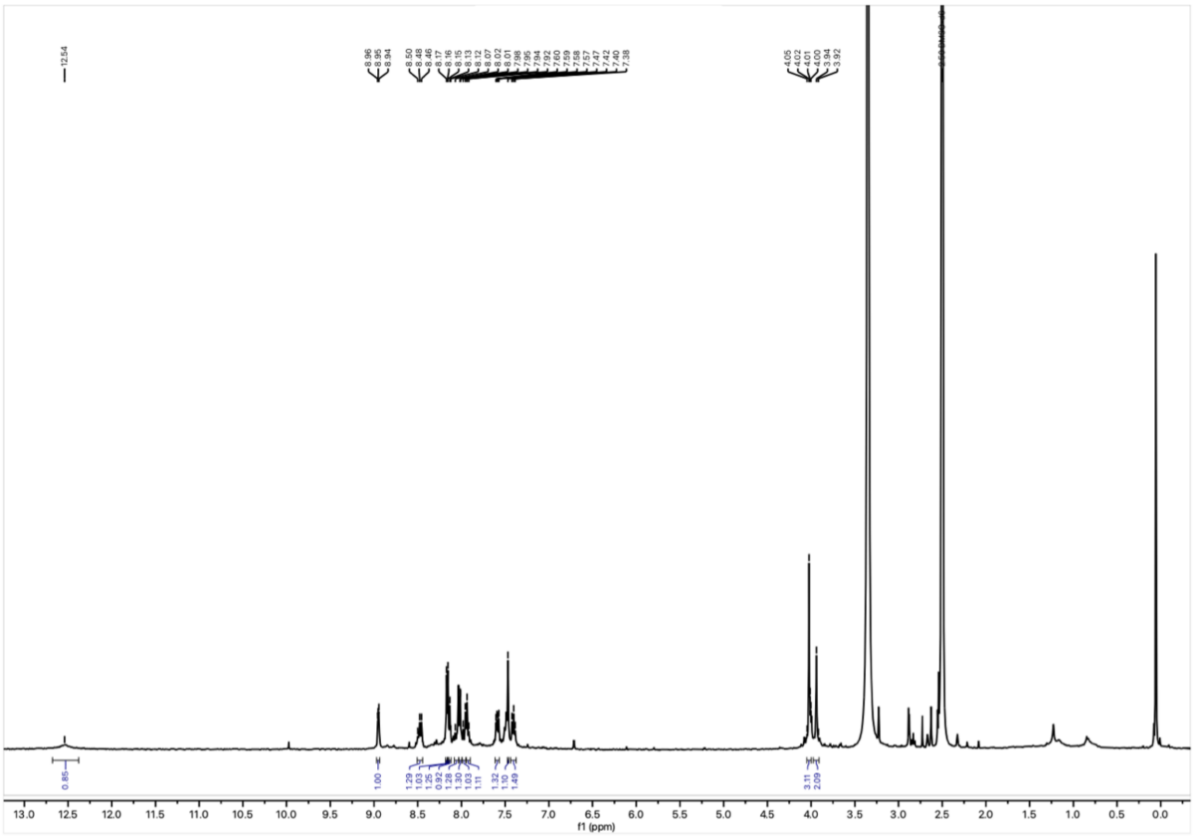

$^{1}H$

NMR spectrum of **41** in DMSO-$d_6$ at 400 MHz.

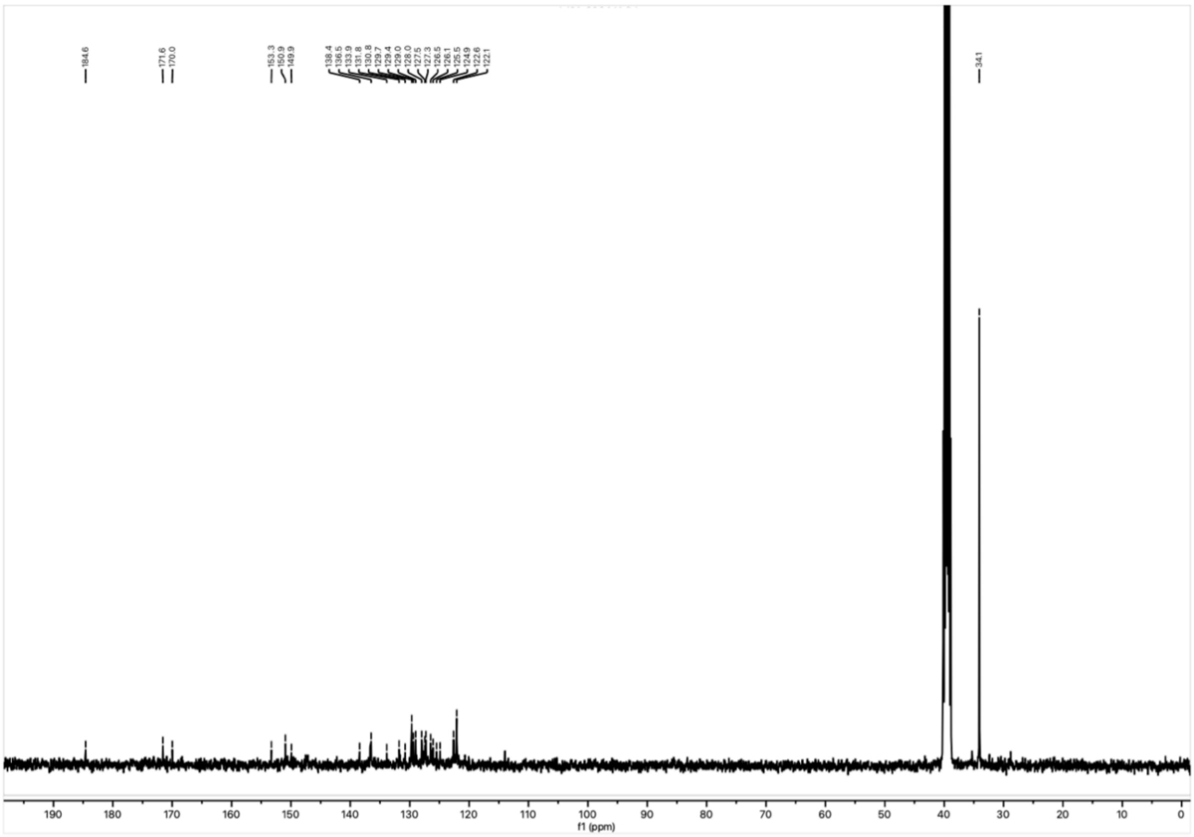

$^{13}C$

NMR spectrum of **41** in DMSO-$d_6$ at 101 MHz.

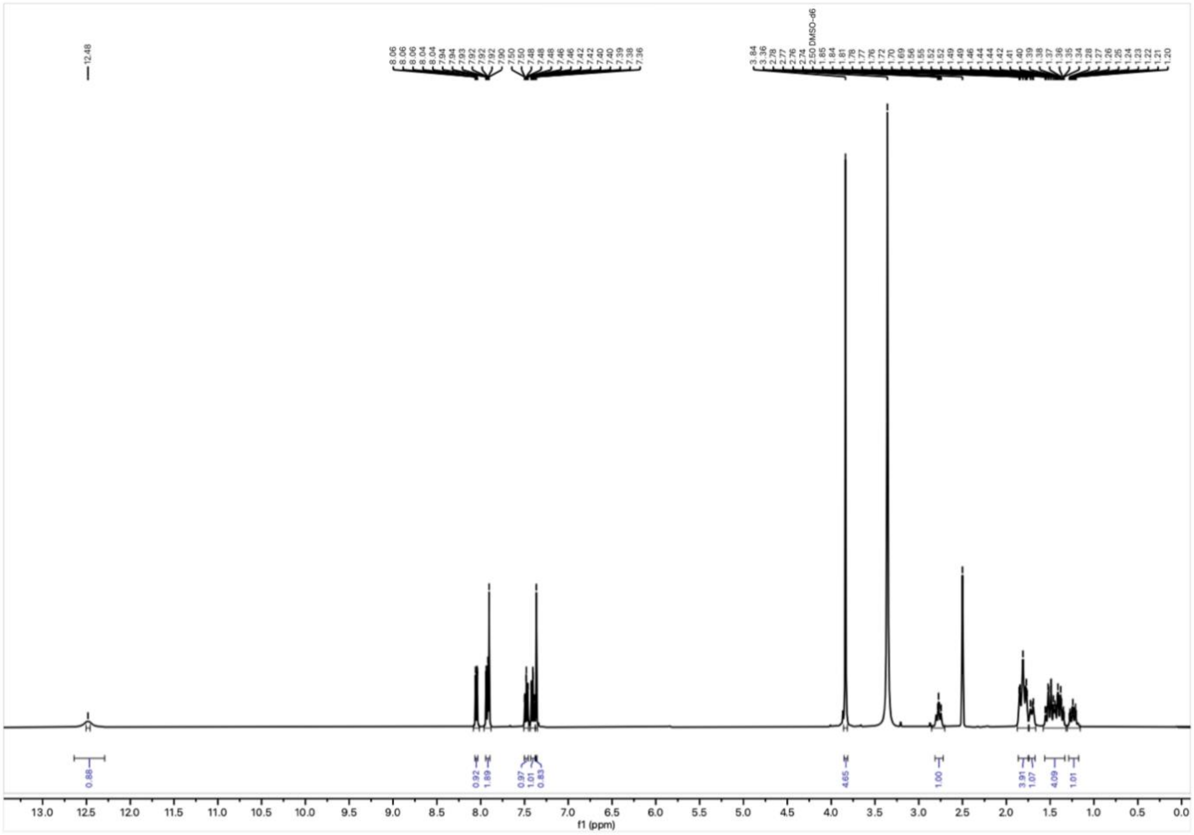


$^{1}$H NMR spectrum of **42** in DMSO-$d_6$ at 400 MHz.

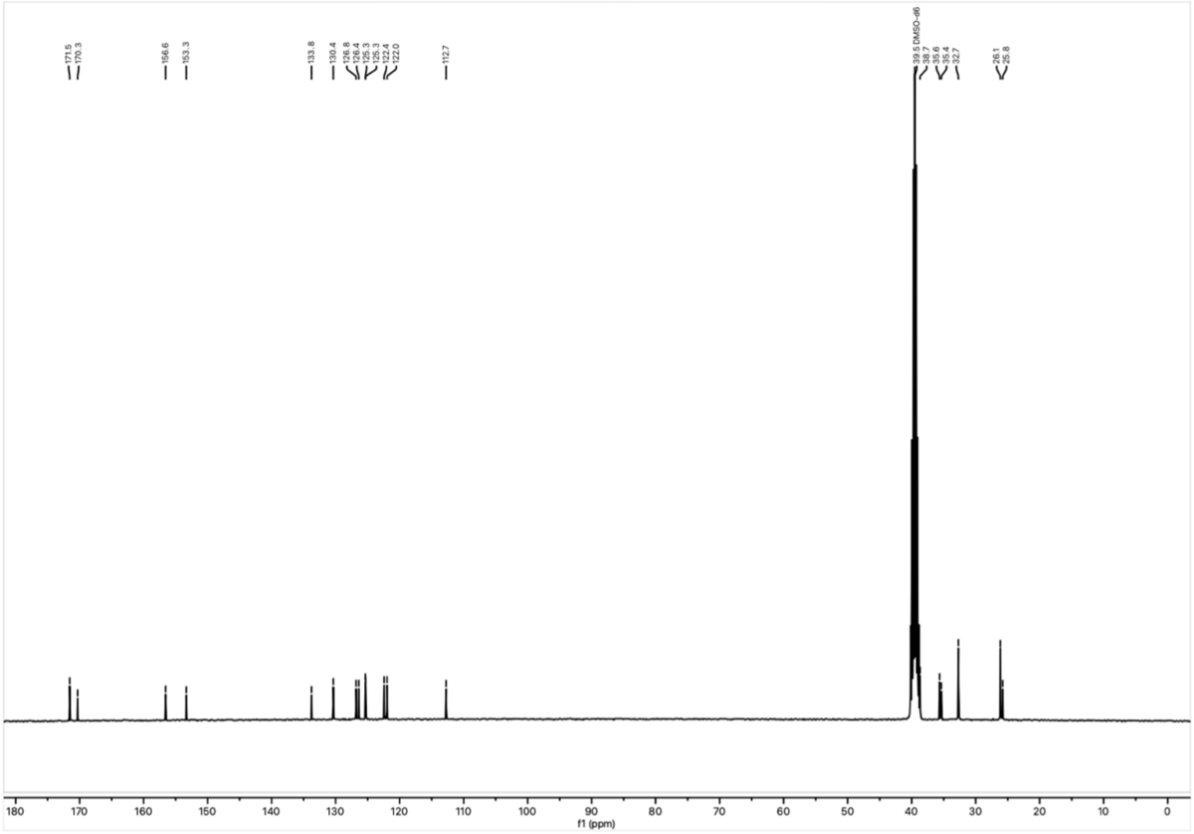


$^{13}$C NMR spectrum of **42** in DMSO-$d_6$ at 101 MHz.

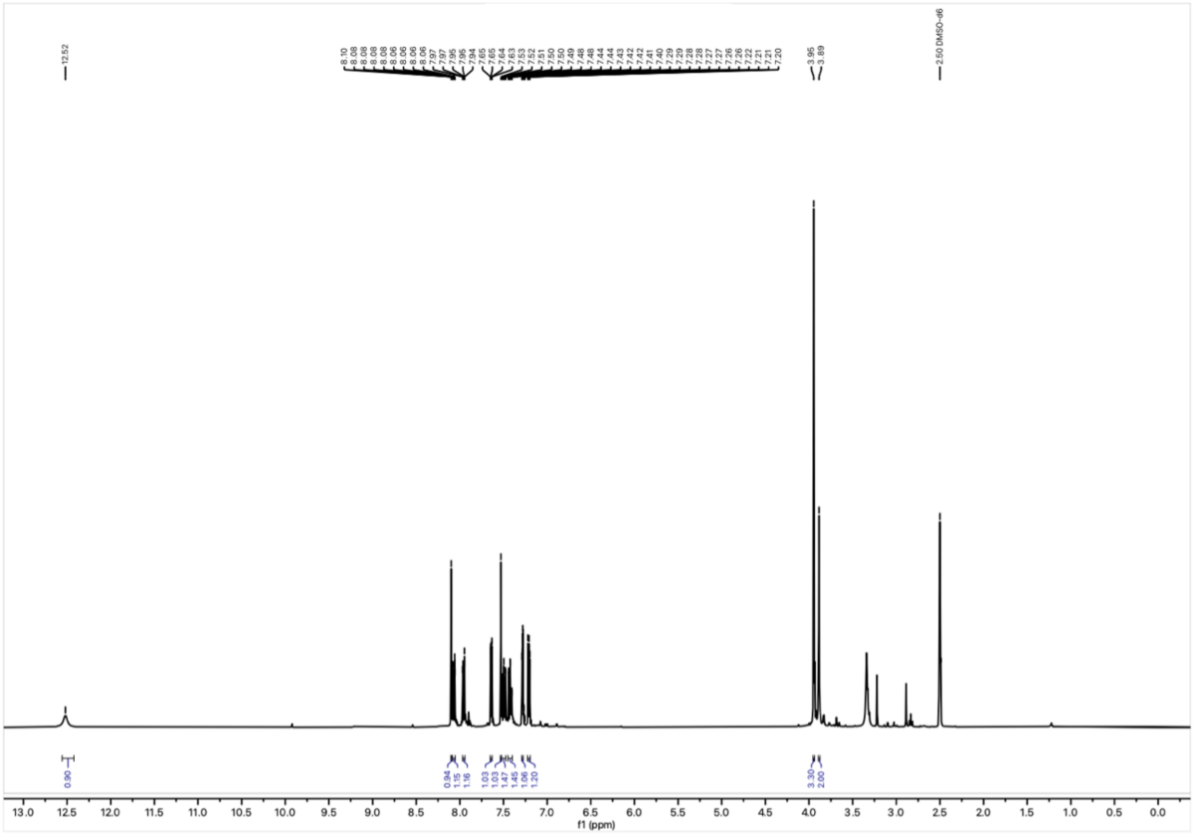

$^{1}H$ NMR spectrum of **43** in DMSO-$d_6$ at 400 MHz.

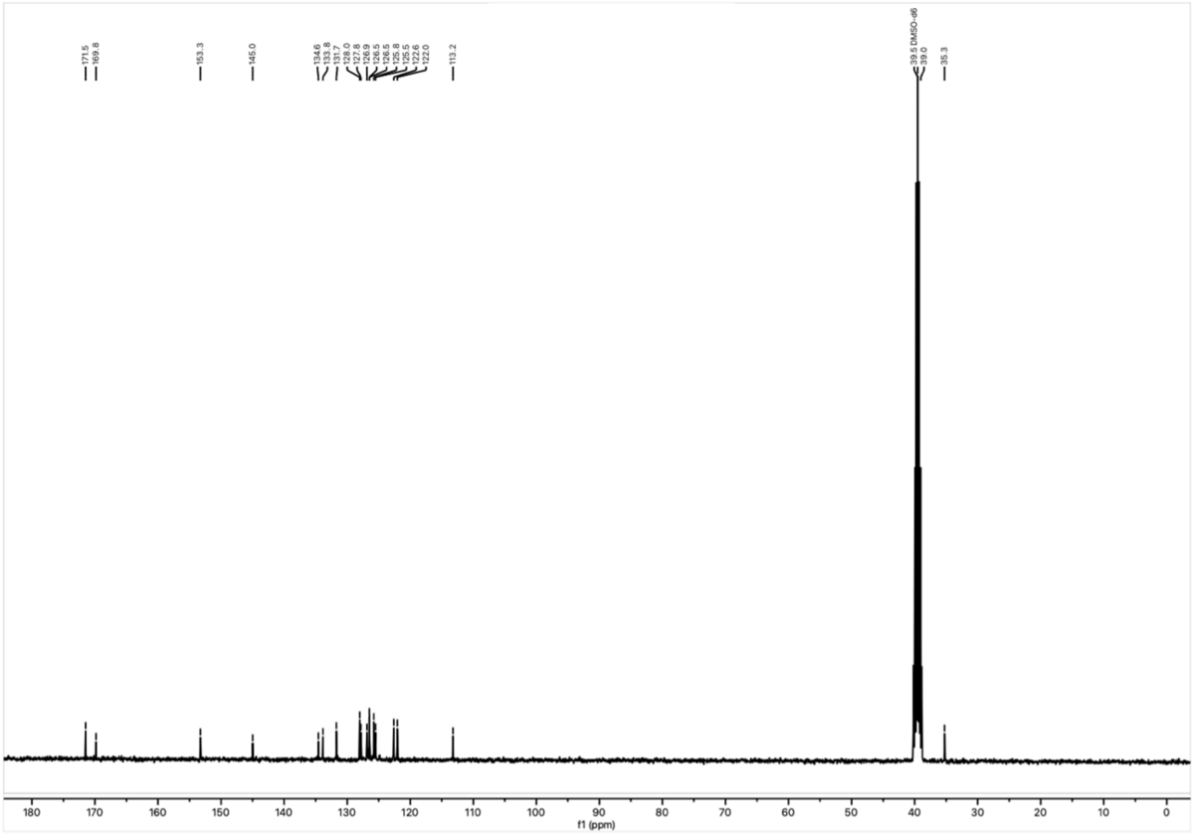

$^{13}C$ NMR spectrum of **43** in DMSO-$d_6$ at 101 MHz.

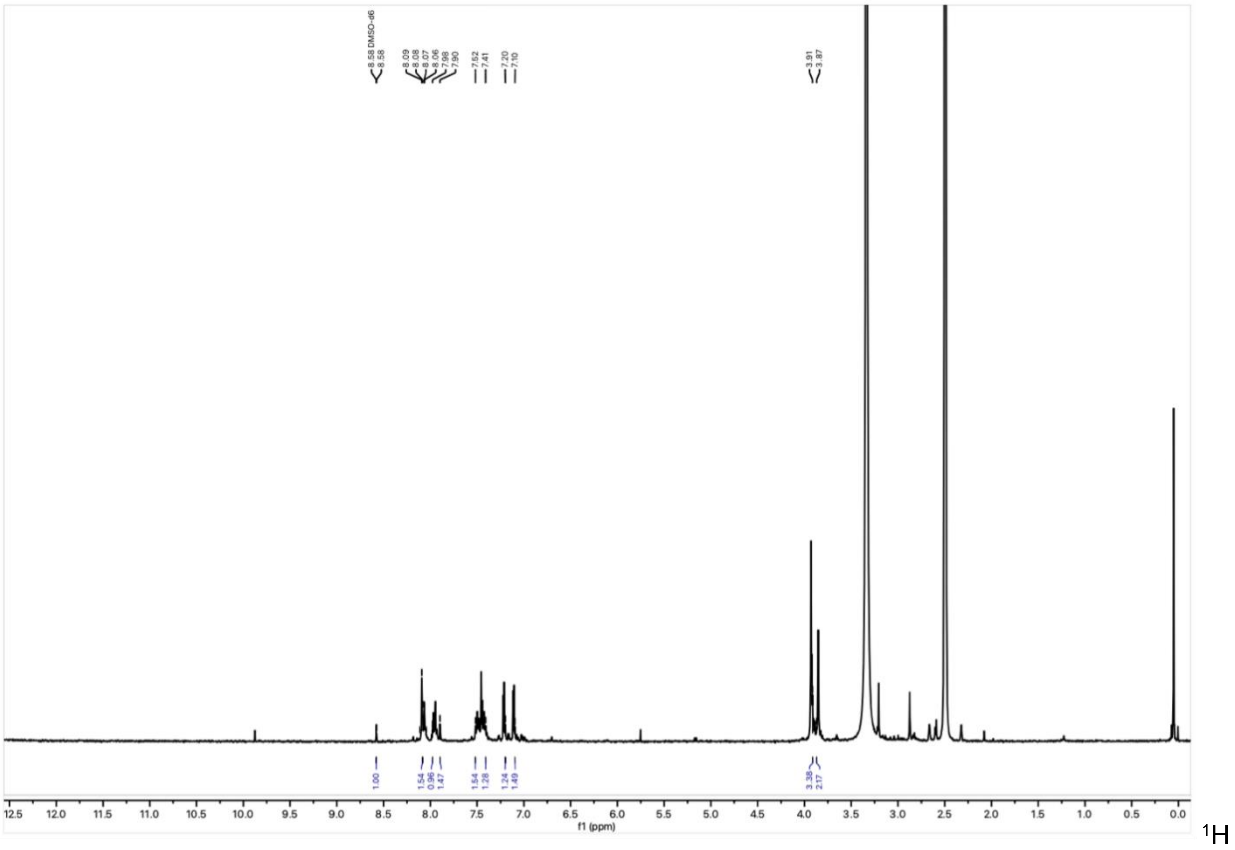


NMR spectrum of **44** in DMSO-$d_6$ at 400 MHz.

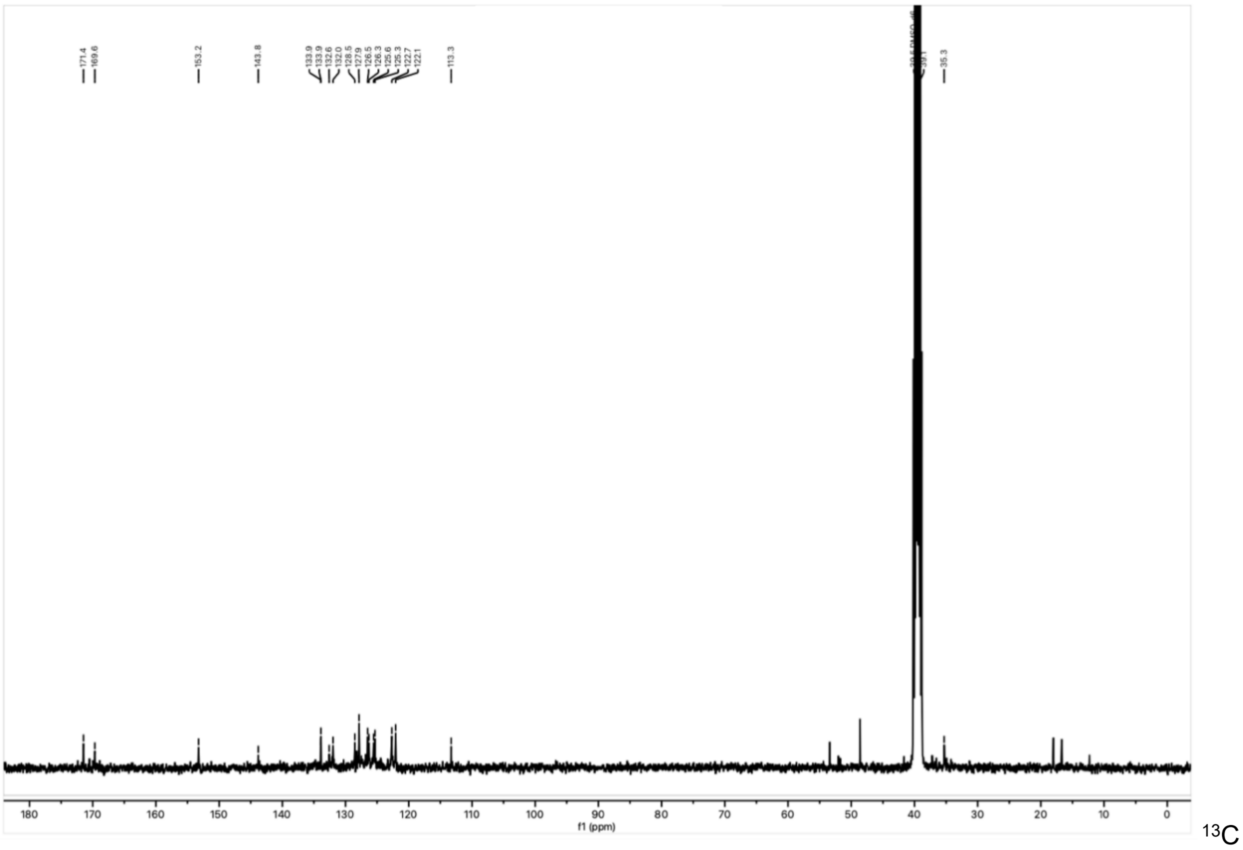


NMR spectrum of **44** in DMSO-$d_6$ at 101 MHz.
